# Using single-plant -omics in the field to link maize genes to functions and phenotypes

**DOI:** 10.1101/2020.04.06.027300

**Authors:** Daniel Felipe Cruz, Sam De Meyer, Joke Ampe, Heike Sprenger, Dorota Herman, Tom Van Hautegem, Jolien De Block, Dirk Inzé, Hilde Nelissen, Steven Maere

## Abstract

Most of our current knowledge on plant molecular biology is based on experiments in controlled lab environments. Over the years, lab experiments have generated substantial insights in the molecular wiring of plant developmental processes, stress responses and phenotypes. However, translating these insights from the lab to the field is often not straightforward, in part because field growth conditions are very different from lab conditions. Here, we test a new experimental design to unravel the molecular wiring of plants and study gene-phenotype relationships directly in the field. We molecularly profiled a set of individual maize plants of the same inbred background grown in the same field, and used the resulting data to predict the phenotypes of individual plants and the function of maize genes. We show that the field transcriptomes of individual plants contain as much information on maize gene function as traditional lab-generated transcriptomes of pooled plant samples subject to controlled perturbations. Moreover, we show that field-generated transcriptome and metabolome data can be used to quantitatively predict at least some individual plant phenotypes. Our results show that profiling individual plants in the field is a promising experimental design that could help narrow the lab-field gap.

## INTRODUCTION

Efforts to develop crops with higher yield and higher tolerance to environmental stress are more important than ever in the quest for global food security and sustainable agriculture. Crop improvement increasingly relies on the identification of genes and genetic variants that impact agronomically important traits, so that beneficial variants can be engineered into the crop or incorporated in breeding programs. Mapping of quantitative trait loci (QTLs), genome-wide association studies (GWAS) and genomic prediction techniques are some of the currently preferred means of identifying the genes and variants influencing a phenotypic trait (Korte and Farlow, 2013; Desta and Ortiz, 2014). All are based on associating genetic variants, mostly single-nucleotide polymorphisms (SNPs), to observed traits in a genetically diverse population of the targeted plant species, e.g. a panel of accessions or a panel of inbred crosses between two or more parental lines (recombinant inbred lines, RILs).

Although fairly successful in some plant species, e.g. maize, these techniques also have limitations. They can only detect loci that display genetic variation in the mapping population. In addition, their resolving power is limited by linkage disequilibrium (LD), i.e. the non-random association between markers due to genetic relatedness in the population (Brachi et al., 2011; Korte and Farlow, 2013; Huang and Han, 2014). As a consequence, loci can often not be resolved to the individual gene level. GWA studies also have low power for rare alleles and alleles with small effect sizes, which often account for a substantial proportion of phenotypic variation, in particular for complex traits such as yield. Moreover, when mapping genotypes straight to phenotypes, the many intermediate molecular layers that articulate the phenotype from the genotype, such as the transcriptome or metabolome, are ignored. Consequently, little mechanistic insight is gained from GWAS or genomic prediction studies into how a trait is established.

As many variants uncovered in GWA studies appear to be regulating gene expression (Li et al., 2012; Xiao et al., 2017), recent efforts have sought to complement GWAS with transcriptome-wide association studies (TWAS), i.e. mapping gene expression to phenotypes in a genetically diverse population (Harper et al., 2012; Koprivova et al., 2014; Pasaniuc and Price, 2017; Havlickova et al., 2018; Kremling et al., 2019). Similarly, several recent studies have used transcriptomic or metabolomic prediction in addition to genomic prediction to associate genes to plant traits, in particular in maize (Guo et al., 2016; Schrag et al., 2018; Azodi et al., 2020). Azodi et al. (2020) found that transcript levels and genetic marker data have comparable performance for predicting maize phenotypes, and that performance increased when combining both data layers in a joint model. However, the use of transcriptomes and other intermediate data layers to aid genotype-phenotype mapping generally remains underexplored (Baute et al., 2015, 2016; Kremling et al., 2019).

Whereas GWAS and related methods exploit the natural genetic variation within a species to associate genes with phenotypes, systems biology studies use controlled perturbations, either genetic, environmental or chemical, in a specific genetic background to unravel the molecular wiring of plant traits. Since the advent of high-throughput gene expression profiling platforms, massive amounts of data have been generated on the transcriptomic responses of e.g. *Arabidopsis thaliana* Col-0 to various mutations and environmental stresses, with the purpose of unraveling the molecular processes underlying a variety of traits. However, many independent perturbations are needed to accurately reconstruct the molecular network underlying a complex trait, and no datasets exist in which any particular complex plant trait is systematically assessed molecularly and phenotypically under a large-enough set of perturbations to unravel more than fragments of its molecular wiring.

The identification of a sufficient set of controlled perturbations informative of a process of interest is one of the major bottlenecks in present-day systems biology. It is often practically infeasible to identify, let alone implement, a large enough number of different controlled perturbations (mutants, stresses) relevant to a trait of interest in a single plant lineage (in contrast to GWA studies, where the genetic differences across lineages function as perturbations). Another issue is that such controlled perturbations are mostly applied in a lab environment, where apart from the imposed perturbation all other parameters are kept optimal and do not restrict plant growth and development. This situation does not reflect realistic field conditions, where at any given time plants are exposed to a combination of different environmental stressors with highly variable temporal and spatial patterns of occurrence (Mittler and Blumwald, 2010; Thoen et al., 2017). Increasing evidence is pointing towards the unique character of plant molecular responses to combinations of stresses, which often have non-additive effects on the molecular and phenotypic level (Atkinson and Urwin, 2012; Rasmussen et al., 2013; Cabello et al., 2014; Johnson et al., 2014; Suzuki et al., 2014; Barah et al., 2016; Davila Olivas et al., 2017; Thoen et al., 2017). As a result, perturbation studies performed under controlled laboratory conditions are often of limited predictive value for phenotypes in the field (Mittler, 2006; Oh et al., 2009; Atkinson and Urwin, 2012; Nelissen et al., 2014; Nelissen et al., 2019). It has been advocated that to close this lab-field gap, more -omics data and associated phenotypic data should be generated on field-grown plants (Alexandersson et al., 2014; Nelissen et al., 2019; Zaidem et al., 2019). Several pioneering studies have already investigated how gene expression is related to environmental stimuli in the field (Nagano et al., 2012; Richards et al., 2012; Plessis et al., 2015). Large-scale studies relating field-generated transcriptomes to field phenotypes are however still lacking.

Here, we propose a new strategy for studying the wiring of plant pathways and traits directly in the field, involving -omics and phenotype profiling of individual plants of the same genetic background grown in the same field. Uncontrolled variations in the micro-environment of the individual plants hereby serve as a perturbation mechanism. Our expectation is that, in addition to stochastic effects, the individual plants will be subject to subtly different sets of environmental cues, and will in response exhibit different molecular profiles and phenotypes. The aim of this study is to investigate to what extent we can use such individual plant differences in the field to link genes to biological processes and field phenotypes. Earlier, we found that gene expression variations among individual *Arabidopsis thaliana* plants grown under the same stringently controlled lab conditions contain a lot of information on the molecular wiring of the plants, on par with traditional expression profiles of pooled plant samples subject to controlled perturbations (Bhosale et al., 2013). If even gene expression variability among lab-grown plants contains functionally relevant information, the molecular and phenotypic variability among field-grown plants may contain a wealth of information on processes occurring in the field.

We profiled the ear leaf transcriptome, ear leaf metabolome and a number of phenotypes for individual field-grown maize plants of the same inbred line (*Zea mays* B104), and used the resulting data to predict the function of genes and to quantitatively predict individual plant phenotypes. We find that our single-plant transcriptome dataset can predict the function of maize genes as efficiently as traditional lab-based perturbational datasets. Furthermore, we show that some quantitative phenotypes, in particular leaf blade width and length, can be predicted fairly well from the leaf transcriptome and metabolome data generated for the individual plants. These results open perspectives for the further use of field-generated single-plant datasets to unravel the molecular networks underlying crop phenotypes and stress responses in the field.

## RESULTS

### Field trial design and exploratory data analysis

During the 2015 growth season, 560 maize plants of the B104 inbred line were grown in a field in Zwijnaarde, Belgium (see Methods and Figure 1). At tasseling (VT stage), the ear leaf and the growing ear were harvested for 200 non-border plants with a primary ear at leaf 16, and plant height, the number of leaves, the length and width of the ear leaf (leaf 16) blade, husk leaf length and ear length were measured (Supplemental Data Set 1). For 60 randomly chosen plants out of these 200, the transcriptome of mature ear leaf tissue was profiled using RNA-seq. Additionally, for 50 out of those 60 plants, metabolite profiles were generated on the same samples used for transcriptome profiling. After pre-processing and filtering (see Methods), data on the levels of 18,171 transcripts and 598 metabolites in mature ear leaf tissue were obtained for 60 and 50 plants, respectively (Supplemental Data Set 1).

**Figure 1.**
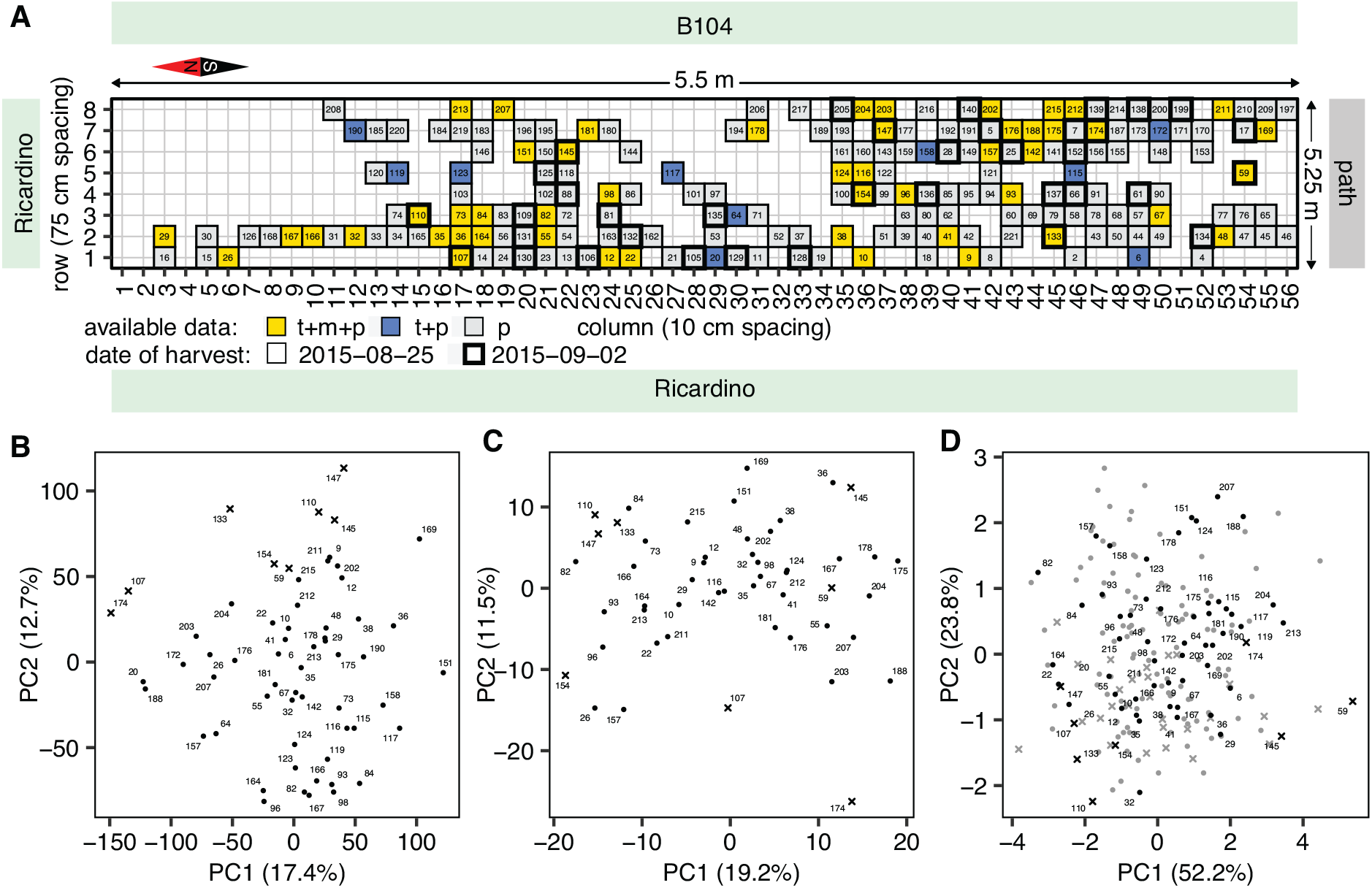
Field trial design and exploratory data analysis. **(A)** Layout of the field trial. A total of 560 *Zea mays* B104 plants were grown in a grid of 10 rows by 56 columns. Border rows 0 and 9 are not shown on the plot, and the dimensions on the figure are not to scale. Phenotypic data was measured for 200 plants (p). Transcriptome (RNA-seq) data was profiled for 60 out of those 200 plants (t+p). Metabolome data is available for 50 out of those 60 plants (t+m+p). Some plants were harvested later (see Methods), as indicated by a thicker cell border. **(B)** Plot showing the first two principal components (PCs) in a PCA of the 60 single-plant transcriptomes. **(C)** Plot showing the first two principal components (PCs) in a PCA of the 50 single-plant metabolomes. **(D)** Plot showing the first two principal components (PCs) in a PCA of the 200 plant phenomes. Light grey markers in panel **(D)** indicate plants for which only phenotype information is available. Only plants for which transcriptome data is available are numbered in plots **(B)**-**(D)**, according to the numbering in panel **(A)**. Crosses in panels **(B)**-**(D)** indicate plants harvested on the second harvest day.

As no differential treatments or control measures were applied to any plant subsets, no distinct sample groups are expected in our data, with the possible exception of subsets of plants harvested on different dates (because of developmental differences between plants, see Methods). Indeed, principal component analysis (PCA) on the gene expression, metabolite and phenotype data (Figure 1) did not reveal a clear group structure among the samples, although the date of harvest does have a clear effect along PC2 of the transcriptome and phenotype profiles of the plants. Despite the absence of designed major effects in our experimental setup, other than the harvesting date, we observed substantial variability in the transcriptome and metabolome profiles and the phenotypes of the individual plants (Figure 2). Transcript levels have on average a coefficient of variation (CV) of 0.3037 across plants, metabolite levels have a CV of 0.3128 on average, and all phenotypes have a CV ≥ 0.0521. This variability could either be caused by technical noise, inherent stochasticity of molecular processes within the plant, or external factors such as variability in the growth micro-environment of the individual plants. The last two processes are expected to generate biologically meaningful variation that may propagate from the molecular to the phenotypic level, or vice versa.

**Figure 2.**
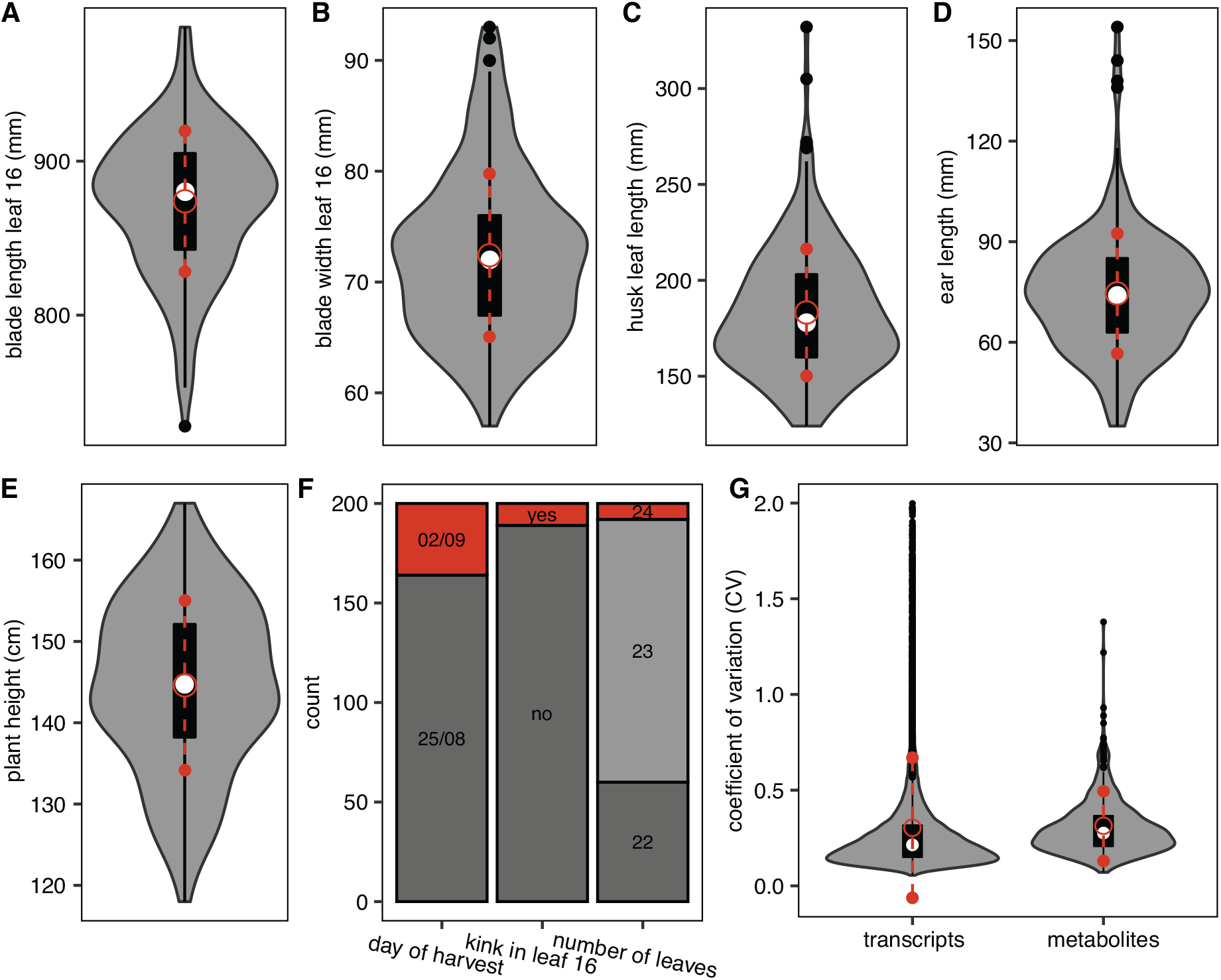
Transcriptomic, metabolic and phenotypic variability among individual field-grown maize plants. In panels **(A)** to **(E)**, violin plots show the variability in continuous leaf 16, ear and plant height phenotypes among individual plants. Panel **(F)** depicts the variability in harvesting date among plants, as well as the variability in two discrete phenotypes, namely the number of leaves at harvest and whether or not leaf 16 was kinked. Panel **(G)** shows violin plots for the distribution of the coefficient of variance (CV) across the sampled plants for the levels of individual transcripts and metabolites. For visualization purposes, the transcript CV was capped at 2.0. In all violin plots, the median is indicated by the white circle. The black box extends from the 25th to the 75th percentile, and black whiskers extend from each end of the box to the most extreme values within 1.5 times the interquartile range from the respective end. Data points beyond this range are shown as black dots. The red open circle indicates the mean of the distribution, with red whiskers extending to 1 standard deviation above and below the mean.

If the variability in the data is biological in nature and propagates through the molecular networks of the plant, plants with similar gene expression profiles may be expected to also have similar metabolite and phenotype profiles. Indeed, plant-to-plant distances in transcriptome, metabolome and phenotype space were found to be significantly positively correlated (Supplemental Figure 1). Interestingly, the phenotype distance between plants was also significantly positively correlated with the physical distance between plants in the field. All phenotypes were found to be spatially autocorrelated at *q*≤0.05 (see Methods, Supplemental Figure 2 and Supplemental Data Set 2). A weak but borderline significant positive correlation was also found between the metabolome distance and physical distance between plants, and 24 out of 592 metabolites exhibit spatial patterning at *q*≤0.01 (Supplemental Data Set 2). No significant correlation was found between the physical distance of plants and their overall distance in transcriptome space (Supplemental Figure 1), indicating that most genes do not exhibit spatially patterned gene expression. However, spatial autocorrelation analysis of the transcriptome data revealed that 1,134 out of 18,171 transcripts do exhibit spatial patterning at *q*≤0.01 (Supplemental Data Set 2). The spatially autocorrelated transcripts were grouped in 30 co-expression clusters plus one ‘noise’ cluster (see Methods and Supplemental Data Set 3, cluster 1 is the noise cluster). Significant GO enrichments were found in 17 of these autocorrelated transcript clusters, e.g. cluster 3 was found enriched in genes involved in the response to chitin, cluster 16 in reproductive system development genes, and cluster 31 in chloroplast-associated genes (Supplemental Data Set 3). This indicates that the activity of several biological processes varied across the field in a spatially patterned way. Eleven of the 30 autocorrelated transcript clusters correlated with at least one measured phenotype at *q*≤0.05 (Supplemental Data Sets 3 and 4). The average gene expression profile of cluster 29 for instance correlates significantly with ear length (Figure 3). Interestingly, two of the 35 genes in cluster 29 are homeotic transcription factors, and both have previously been associated with ear development: GRMZM2G171365 (*SUPPRESSOR OF OVEREXPRESSION OF CONSTANS 1*, *ZmSOC1*, *ZmMADS1*), a MADS-box transcription factor known to promote flowering (Zhao et al., 2014; Alter et al., 2016) and also known to be upregulated in leaves during the floral transition (Alter et al., 2016), and GRMZM2G034113 (*hb126*), a homeobox transcription factor previously found in a GWAS study as a candidate gene for ear height (Li et al., 2016). Overall, the presence of spatially autocorrelated patterns in the transcriptome, metabolome and phenotype data indicate that at least part of the variability observed among the individual plants is due to micro-environmental factors that have a spatial structure. Correlations between the molecular and phenotypic data layers indicate that this variability propagates from one layer to another.

**Figure 3.**
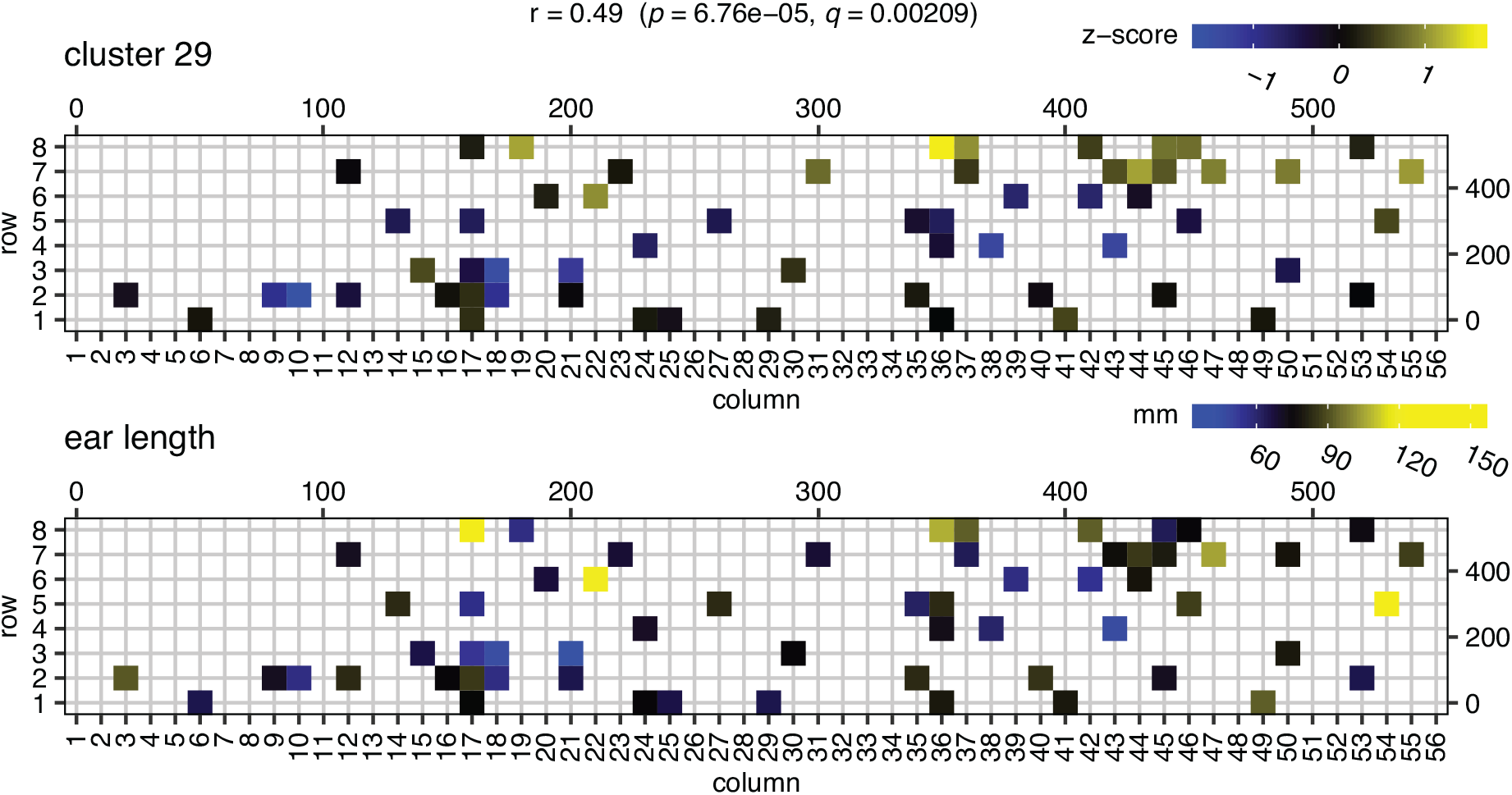
Gene expression patterns in cluster 29 correlate with ear length. The top panel displays the average *z*-scored gene expression profile of spatially autocorrelated gene cluster 29 (35 genes), mapped to the field. The bottom panel displays the ear length phenotype on the field (only for plants that were transcriptome profiled). Shown on top are the Pearson’s correlation (*r*) between the cluster 29 expression profile and ear length, the corresponding *p*-value (computed using cor.test in R) and the corresponding *q*-value (computed using the Benjamini-Hochberg method on all comparisons of cluster gene expression profiles with the ear length profile). The scales on the top and to the right of the field maps give field plot dimensions in cm.

### Variability of gene expression across plants gives insight into biological processes active in the field

We investigated which genes have highly variable expression levels in the field setting used, and which ones are stably expressed across the field. We ranked genes based on the coefficient of variation (CV) of their gene expression profile across the field (Supplemental Data Set 5), excluding the 5% lowest expressed genes. We found that stably expressed genes have on average longer coding sequences than variably expressed genes and have on average more introns and exons (Supplemental Table 1). Similar results were previously obtained in a study in which individual lab-grown *Arabidopsis thaliana* plants were expression profiled (Cortijo et al., 2019), and the authors showed that their observations could not be accounted for by technical artefacts related to differences in the average RNA-seq coverage of longer versus shorter genes. Similar to Cortijo et al. (2019), we also found that variably expressed genes are on average connected to 6.54 times more transcription factors than stably expressed genes in a coexpression network constructed from the single-plant transcriptome data (see Methods, one-tailed Mann– Whitney U (MWU) test, *q* = 5.92E-59). This again suggests that at least part of the observed variability in gene expression levels across plants is biological in nature.

Mann–Whitney U tests (Mann and Whitney, 1947) were performed to determine which Gene Ontology (GO) biological processes are represented more at the top or bottom of the CV-ranked gene list than expected by chance (Supplemental Data Set 6). Genes related to cell wall organization, biotic stresses impacting the cell wall (herbivores, chitin), secondary metabolism, photosynthesis, abscisic acid transport, brassinosteroid and trehalose metabolism and gibberilic acid signaling were found to be among the more variably expressed genes across the field, suggesting that the harvested leaves were differentially impacted by biotic and possibly abiotic stress factors. The processes that are most stably expressed across the field are mainly housekeeping processes related to e.g. the metabolism and transport of proteins and mRNAs, and chromatin organization (Supplemental Data Set 6). However, not all genes annotated to ‘stable’ GO processes are stably expressed. The top-10 of most variably expressed genes for instance includes eight genes involved in chromatin organization or DNA replication, among which five histones (Supplemental Data Set 5). Interestingly, the GO enrichments obtained for variably and stably expressed genes in the field-grown maize plant dataset are largely in line with the results reported by Cortijo et al. (2019) on the variability of gene expression in individual lab-grown *A. thaliana* plants. Photosynthesis, secondary metabolism, cell wall organization and defense response genes for instance were also found enriched by Cortijo et al. (2019) in several of the highly variable gene sets they compiled for different sampling time points in a 24h time span, while RNA and protein metabolism genes feature prominently in some of their lowly variable gene lists.

Hierarchical clustering of the transcriptome and metabolome data offers an overall view of the molecular variability across the plants profiled (Supplemental Figure 3). Several clusters were found to be significantly enriched in genes involved in particular biological processes, further confirming that the single-plant dataset contains biologically meaningful information (Supplemental Data Set 7). Also the biclustering approaches ISA (Bergmann et al., 2003), SAMBA (Tanay et al., 2002) and ENIGMA (Maere et al., 2008) yielded a variety of modules enriched for genes involved in processes such as photosynthesis, cell wall organization, response to chitin and others (Supplemental Data Set 7). An example ENIGMA module, enriched for known reproductive development genes, is shown in Figure 4. In this module and many others (see e.g. the photosynthesis and response to chitin clusters in Supplemental Figure 3), different subgroups of plants show clearly different expression profiles, highlighting that many processes are not homogeneously active across the field.

**Figure 4.**
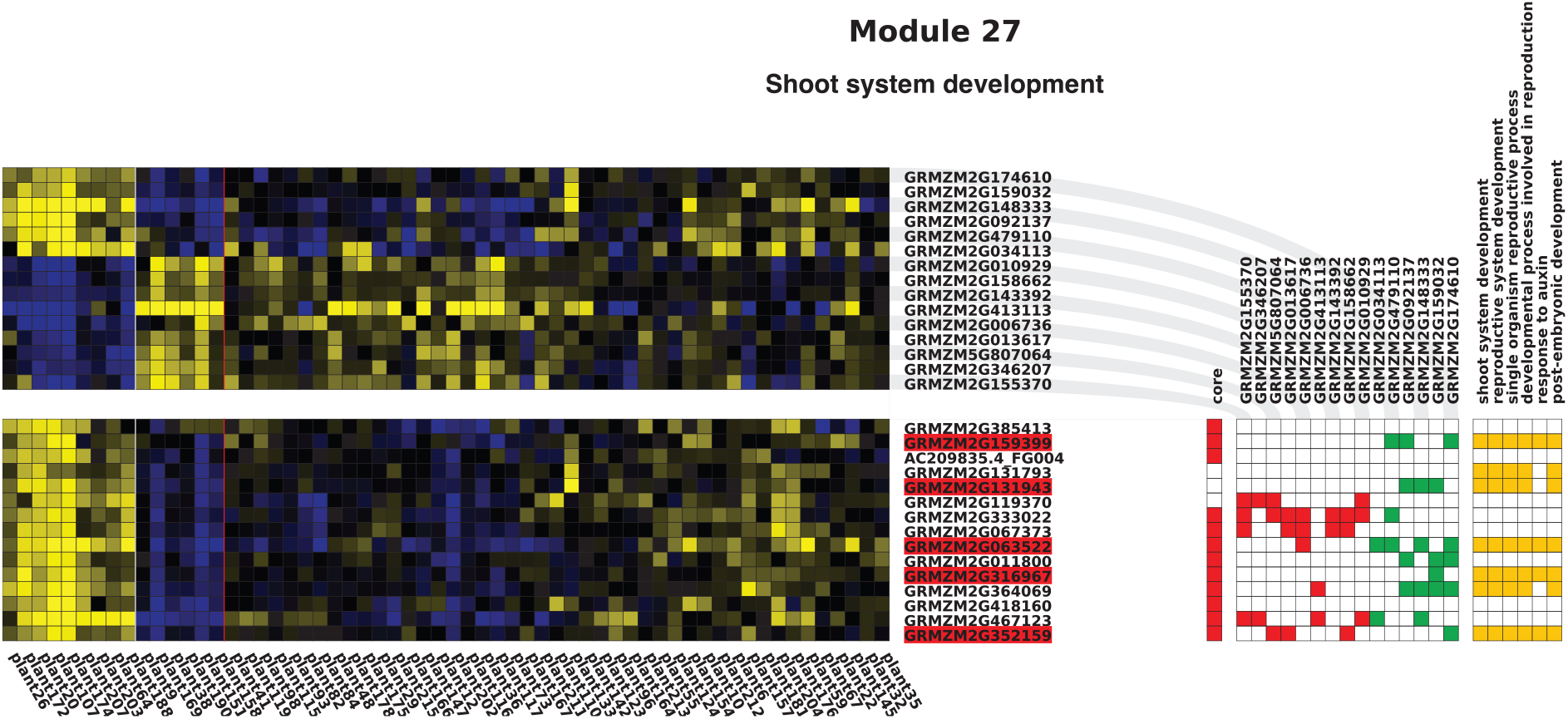
Example ENIGMA module learned from the single-plant transcriptome dataset. Yellow/blue squares indicate higher/lower gene expression with respect to the average expression of a gene across plants. The bottom grid shows the expression profiles of the module genes, while the top grid contains the expression profiles of predicted regulators of the module. Significant co-differential expression links between the regulators and the module genes are indicated in the red/green matrix to the right (green = positively correlated, red = negatively correlated). Gene names highlighted in red indicate regulators that are part of the module. Genes indicated as core genes belong to the original module seed, other genes were accreted by the seed in the course of module formation (Maere et al., 2008). Enriched GO categories in the module gene set are displayed on the right, with orange squares depicting which module genes are annotated to these GO categories. This particular module is significantly enriched (*q* ≤ 0.01) in known reproductive system development genes, mostly regulators.

### Gene function prediction from single-plant transcriptome data

In previous work, we showed that expression variations among individual *Arabidopsis thaliana* plants, all grown under the same stringently controlled conditions, can efficiently predict gene functions (Bhosale et al., 2013). To investigate whether expression variations among maize plants grown under uncontrolled field conditions can similarly be used to predict gene functions, we constructed a network of significantly coexpressed genes from the transcriptome data, using spatially adjusted Pearson correlation coefficients between the log2-transformed gene expression profiles (see Methods). Accounting for the spatial autocorrelation structure of our field-generated data is necessary to avoid inflation of the false positive rate (Lennon, 2000). The function of any given gene in this coexpression network was predicted based on the annotated functions of the gene’s network neighbors (see Methods). To compare the function prediction performance of our single-plant dataset with that of traditional gene expression datasets on pooled samples of plants grown under controlled conditions, we ran the same function prediction pipeline on 500 networks constructed from gene expression datasets on maize leaves available from the Short Read Archive (SRA) transcriptome database (see Methods and Supplemental Data Set 8). Each of these 500 networks was inferred from a dataset of the same size as the single-plant dataset, containing 60 transcriptome profiles sampled from the SRA. The number of significant edges (Bonferroni-corrected *p* ≤0.01) inferred from these sampled datasets was systematically higher than the number of edges inferred from the single-plant dataset. One factor causing this is that the SRA transcriptome data exhibits clear groups of experimental conditions for which expression profiles are more similar within groups than between groups (Supplemental Figure 4), more so than the single-plant data. This group structure causes inflated correlation *p*-values in the sampled networks. Since correlation networks with more edges are biased towards better function prediction performance (Supplemental Figure 5), the number of edges included in each sampled network was fixed to the number of significant edges observed in the single-plant network (771,610 edges). Other network properties such as the number of nodes, network density, average clustering coefficient and unannotated gene fraction were not significantly different between the resulting sampled networks and the single-plant network (Table 1).

**Table 1.**
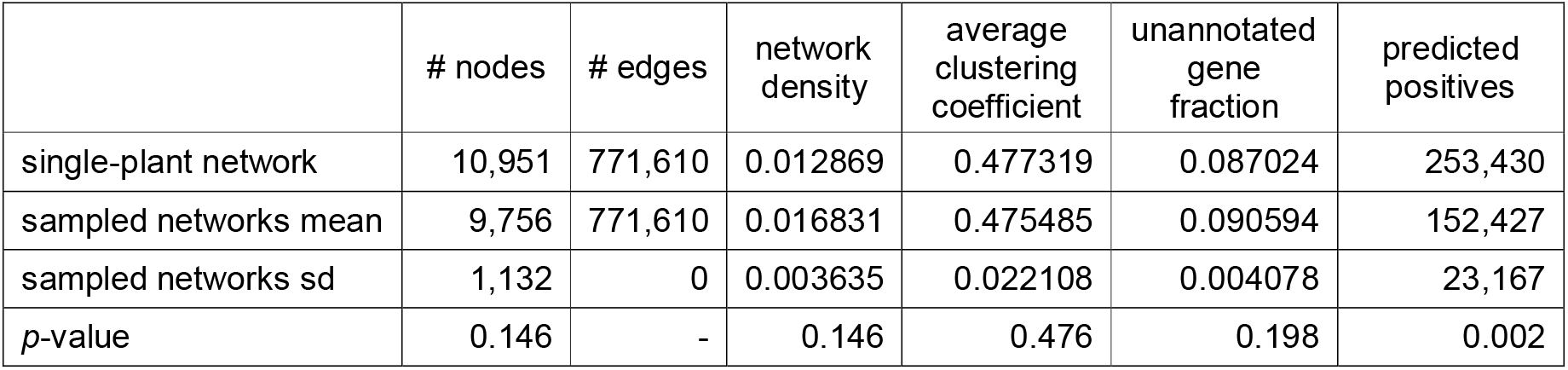
Topological parameters for the single-plant and sampled expression correlation networks. The ‘predicted positives’ column indicates the amount of true positive plus false positive predictions made by each type of network at *q* ≤ 0.01.

The overall gene function prediction performance of all networks was scored using known GO annotations for maize as the gold standard (see Methods). For each network, we calculated the fraction of known gene function annotations recovered by the predictions (recall), the fraction of gene function predictions supported by the gold standard (precision) and the F-measure (harmonic mean of precision and recall) at different false discovery rate (FDR) levels, ranging from *q* = 0.01 to 10^−11^ (Figure 5). Except at the highest-confidence prediction thresholds (*q* ≤ 10^−9^), the recall of the single-plant network was higher than the 75^th^ percentile of the recall values for the sampled networks, indicating that the single-plant network predictions generally recover more known gene functions than the sampled network predictions. On the other hand, the predictions of the single-plant network are generally less precise than those of most sampled networks, except at lower-confidence prediction thresholds (*q* ≥ 10^−4^). As a result, the overall function prediction performance of the single-plant network (as measured by the F-measure) is higher than that of the majority of sampled networks for *q* ≥ 10^−6^, but lower for *q* ≤ 10^−7^. This is mostly due to the lower precision of the single-plant network predictions at higher confidence levels : compared to the sampled networks, a bigger proportion of the high-confidence function predictions made by the single-plant network is not supported by the gold standard.

**Figure 5.**
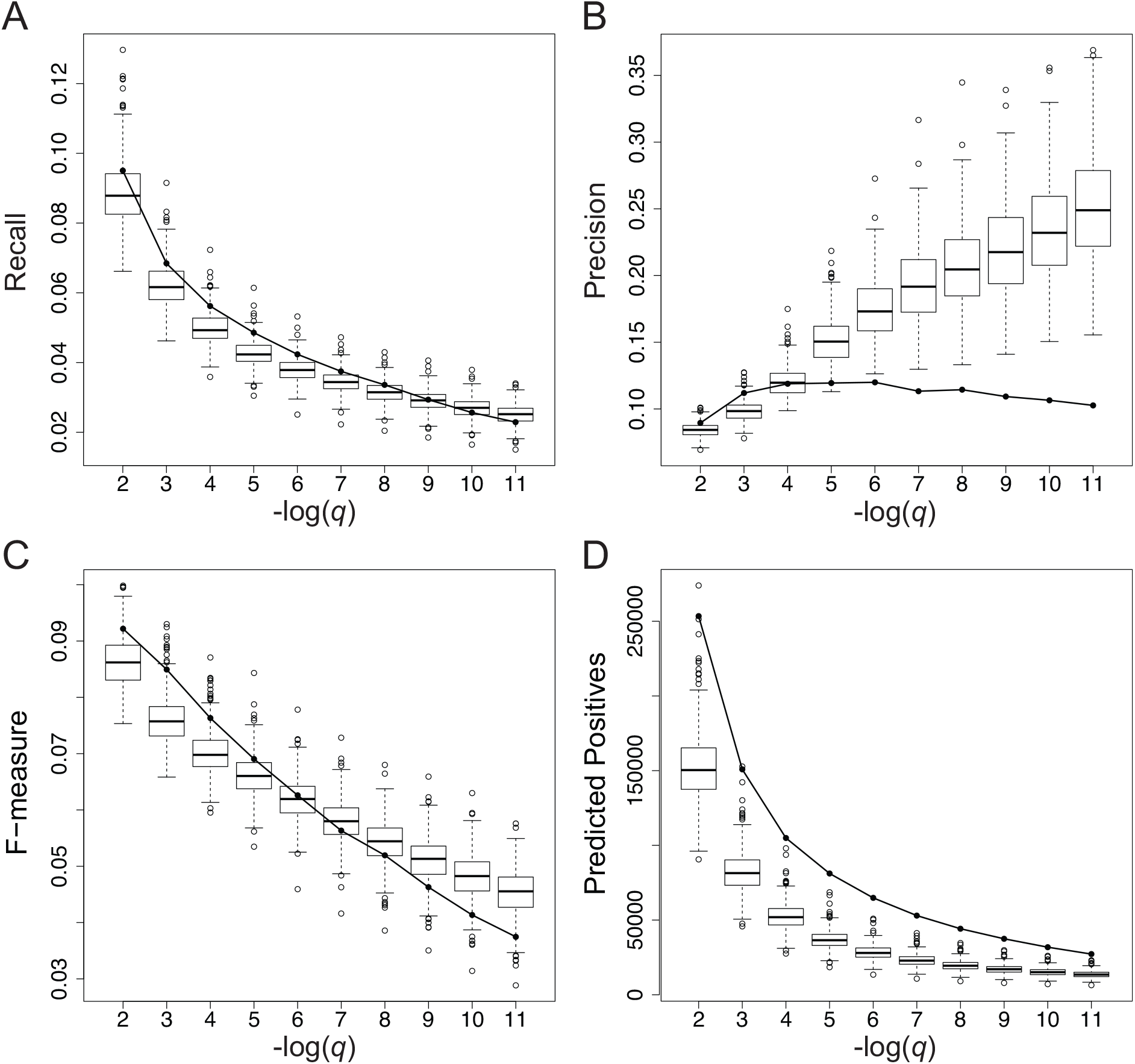
Global gene function prediction performance. Panels **(A)** to **(D)** depict the gene function prediction performance of the single-plant network (solid line) and 500 sampled networks (box-and-whisker plots) averaged across all genes in a given network. Boxes extend from the 25th to the 75th percentile of the sampled networks, with the median indicated by the central black line. Whiskers extend from each end of the box to the most extreme values within 1.5 times the interquartile range from the respective end. Data points beyond this range are displayed as open black circles. Panels **(A)**, **(B)** and **(C)** respectively represent the recall, precision and F-measure of the network-based gene function predictions as a function of the prediction FDR threshold (*q*). Panel **(D)** depicts the number of gene functions predicted from each network (predicted positives = true positives + false positives) as a function of the prediction FDR threshold. As multiple gene functions can be predicted per gene, the number of predicted positives is generally higher than the number of genes.

There are reasons to believe that not all of these excess false positive predictions made by the single-plant network at high confidence levels are truly wrong. First, the GO annotation for maize, used here as the gold standard, is incomplete. Of the 39,479 genes in the maize genome (version V3 5b+), 9,884 have no biological process assignments in the GO annotation file we compiled (see Methods), and many others likely have incomplete or faulty annotations (Rhee and Mutwil, 2014; Wimalanathan et al., 2018). High-confidence gene function predictions labeled as false positives may therefore be regarded rather as new gene function predictions to be tested. By itself however, the incompleteness of the gold standard should not lead to a specific disadvantage for the single-plant network, as all networks are compared on the same footing. More importantly, the current annotations in GO are mostly derived from traditional lab-based perturbation experiments on pooled plant samples, akin to the ones used to construct the sampled networks. This may create a bias in favor of the sampled networks, in particular for the precision measurements (see also Discussion). The recall measure should therefore probably get a higher weight when comparing the gene function prediction performance of the single-plant and sampled networks.

### Single-plant dataset contains information on biological processes that are active and varying between plants in the field context

To assess whether the single-plant dataset contains more information on some biological processes than on others, we investigated how well the gene function predictions on the single-plant network and sampled networks could recover the genes involved in specific biological processes (see Methods). The function prediction performance of all networks was scored for 207 different GO categories, including the categories investigated in (Bhosale et al., 2013) and 56 GO categories that were found enriched in one or more of the (bi)clusters obtained from the single-plant dataset (Supplemental Data Set 9). Figure 6 shows the relative performance of the single-plant network for a selection of GO categories related to abiotic and biotic stress responses, hormonal responses and development (see Supplemental Data Set 9 and 10 for results on other GO categories).

**Figure 6.**
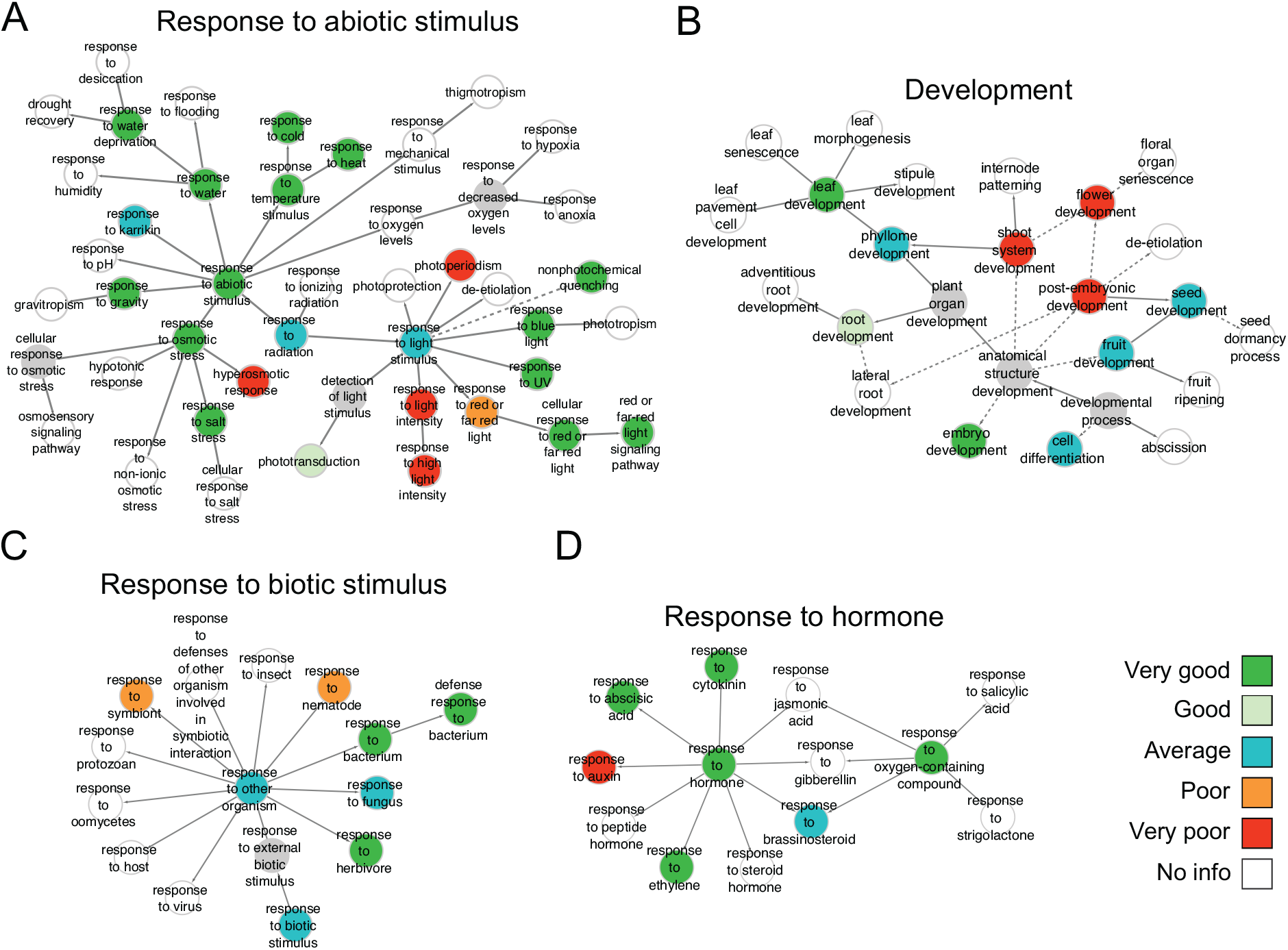
Gene function prediction performance for specific GO categories. Panels **(A)** to **(D)** show the gene function prediction performance of the single-plant network versus sampled networks for GO categories related to abiotic stimulus responses, development, biotic stimulus responses and hormone responses, respectively. Categories are shown in the context of the GO hierarchy and colored according to how well the single-plant network performs in comparison with 500 sampled networks (see Methods). Solid arrows represent direct parent-child relationships in GO, dashed arrows represent indirect relationships. Grey nodes depict untested GO categories. White nodes depict GO categories for which there was insufficient information to score the performance of the single-plant network versus the sampled networks, i.e. categories for which the single-plant network and more than half of the sampled networks did not give rise to any predictions at *q* ≤ 10e−2.

For abiotic stresses, the single-plant network scores very well compared to the sampled networks for responses to cold and heat, salt stress and drought (water deprivation), all of which are relevant from a field perspective. For light responses, the picture is more nuanced, with very good performance for responses to blue light and UV light, ambiguous performance for categories related to ‘*response to red- and far-red light*’ and very poor performance for ‘*response to light intensity*’ and ‘*photoperiodism*’. The overall very good function prediction performance for ‘*response to abiotic stimulus*’ indicates that there is considerable variation across the field in the transcriptional activity of the genes concerned, which suggests that the individual plants were subject to multiple abiotic environmental cues that varied in intensity across the field.

Concerning responses to biotic stimuli, the single-plant predictions score very well for the ‘*response to herbivore*’ and ‘*response to bacterium*’ categories, average for ‘*response to fungus*’, and poor for ‘*response to nematode*’ and ‘*response to symbiont*’ (Figure 6 and Supplemental Dataset 9). This indicates that the individual plants may have been variably exposed to biotic stresses, in particular bacteria and fungi. The single-plant network also scored very well for some GO categories related to biotic stimulus responses that are not shown in Figure 6, such as ‘*defense response’* and ‘*response to chitin’* (Supplemental Data Set 9). The function prediction performance for other biotic stress categories such as ‘*response to insect*’ or ‘*response to oomycetes*’ could not be assessed because both the sampled and single-plant datasets did not yield enough predictions (see Methods).

Similarly, both the sampled and single-plant datasets failed to deliver sufficient predictions to score the function prediction performance for responses to jasmonic acid, gibberellins, salicylic acid and strigolactones. Among the hormone responses for which the gene function prediction performance of the single-plant dataset could be scored, the responses to abscisic acid (ABA), cytokinin and ethylene score very well, ‘*response to brassinosteroids*’ scores average and ‘*response to auxin*’ scores very poorly. The very poor function prediction performance for auxin response genes is consistent with the fact that only mature leaf tissue was profiled in the single-plant experiment, where auxin signaling is less active (Brumos et al., 2018). In contrast, the sampled datasets also contain experiments on entire leaves, leaf primordia and leaf zones such as the division and elongation zone where auxin signaling is more active (Supplemental Data Set 8).

Regarding developmental processes, the single-plant dataset scores very well for predicting genes involved in leaf development and embryo development, well for root development, average for seed and fruit development and very poor for flower development. The (very) good prediction performances for root and embryo development may come as a surprise given that only leaf material was profiled, but one needs to keep in mind that all performances are scored relative to the performance of the sampled datasets, which also exclusively profiled leaves. Even then, it may be considered surprising that leaf expression profiles contain any information at all on developmental processes occurring in other tissues such as roots, flowers or fruits. However, many genes influencing e.g. root development may also function in some capacity in leaves (Taniguchi et al., 2017; Yang et al., 2019). More genuinely surprising is that the single-plant dataset outperforms more than 75% of the sampled datasets for predicting genes involved in leaf development, both in terms of precision and recall, despite only profiling mature leaf tissue of ear leaves.

### Exploration of new maize genes predicted to be involved in biotic and abiotic stress responses

In total, 1,620,503 novel gene function predictions (i.e. predictions not matching GO annotations) were obtained from the single-plant dataset at *q* ≤ 0.01 (Supplemental Data set 11). To assess the quality of these predictions, we performed a literature screen to search for evidence supporting the top-10 regulator predictions for the GO categories ‘*response to chitin*’, ‘*response to water deprivation*’ and ‘*C_4_ photosynthesis*’. The first two are categories for which the single-plant dataset exhibited very good gene function prediction performance compared to the sampled datasets. ‘*C_4_ photosynthesis*’ on the other hand scored very poorly in the single-plant dataset (Supplemental Data Set 9-10). We included this category in the literature validation effort to assess whether poor gene function prediction performance for a biological process, as scored based on which genes are already annotated to the process in GO, also entails that newly predicted links between genes and the process under study are of poor quality.

‘*Response to chitin*’ was among the best-scoring GO categories in our assessment of the gene function prediction performance of the single-plant dataset. Chitin is a main component of fungal cell walls and insect exoskeletons (Fleet, 1991; Latgé, 2007), and the response to chitin is therefore closely related to the responses to fungi and insects. For three out of the top-10 novel transcriptional regulators predicted to be involved in the response to chitin (Supplemental Table 2), we found indirect evidence in literature in support of the predictions. *ZmWRKY53* (GRMZM2G012724), on the 3^rd^ position in the ranking, was previously found to be involved in the response of maize to *Aspergillus flavus,* a fungal pathogen that affects maize kernel tissues and produces mycotoxins that are harmful for humans and animals (Fountain et al., 2015). *ZmWRKY53* was found to be strongly upregulated in both a susceptible and a resistant maize line upon inoculation of kernels with *Aspergillus flavus* (Fountain et al., 2015). Its putative functional ortholog in *Arabidopsis thaliana*, *AtWRKY33*, is known to regulate defense response genes (Zheng et al., 2006; Birkenbihl et al., 2012), and its putative functional orthologs in *Triticum aestivum* (*TaWRKY53*) and *Oryza sativa* (*OsWRKY53*) have previously been suggested to regulate several biotic and abiotic stress response genes, including chitinases (Van Eck et al., 2014). Overexpression of *OsWRKY53* was also shown to increase the resistance of *O. sativa* to herbivory by the brown planthopper *Nilaparvata lugens* (Hu et al., 2016). Another WRKY TF in the top-10 list, *ZmWRKY92* (GRMZM2G449681, rank 5), was previously found to be induced upon *Fusarium verticillioides* inoculation of kernels in the ear rot-resistant maize inbred line BT-1 (Wang et al., 2016). The 8^th^ gene in the top-10 list, GRMZM2G042756 (AP2-EREBP-transcription factor 105), was previously found to be upregulated upon infection of a maize line with *Ustilago maydis*, a basidiomycete fungus that causes common smut in maize (Donaldson et al., 2013).

The second GO category for which we screened literature is ‘*response to water deprivation*’. Four of the top-10 transcriptional regulators predicted to be involved in drought stress responses, but not annotated as such in GO, have previously been linked to drought stress in other studies (Supplemental Table 3). *ZmWRKY40* (GRMZM2G120320, rank 9) was shown to confer drought resistance when it was overexpressed in *A. thaliana* (Wang et al., 2018b). *ZmXLG3b* (GRMZM2G429113, rank 1), encoding a guanine nucleotide-binding protein predicted to be involved in the response to desiccation, was found to be downregulated in the drought-tolerant H082183 line but upregulated in the drought-susceptible maize line Lv28 under severe drought stress versus control conditions (Zhang et al., 2017). Moreover, *ZmXLG3b* was identified as a candidate drought stress response gene in a GWAS study on 300 inbred maize lines, and its expression level was found to anticorrelate with drought stress tolerance levels in four tested maize lines (Yuan et al., 2019). *ZmMPK3-1* (GRMZM2G053987, rank 4), a mitogen-activated protein kinase (MAPK), was previously found to be upregulated in leaf and stem tissue upon drought stress in maize (Liu et al., 2015b). *ZmTPS13.1* (GRMZM2G416836, rank 3), predicted to be involved in drought recovery in our analysis, encodes a putative trehalose-phosphate synthase functioning in the trehalose biosynthesis pathway. The trehalose precursor trehalose-6-phosphate (T6P) is known to function as a signaling molecule coordinating carbohydrate metabolism and developmental processes in plants (Ponnu et al., 2011). Trehalose and T6P have also been implicated in protecting plants from various stresses, including drought stress, but the mechanisms involved are still unclear (Fernandez et al., 2010; Lunn et al., 2014; Nuccio et al., 2015).

Finally, we screened literature for the top-10 regulators predicted to be involved in C_4_ photosynthesis (Supplemental Table 4). Surprisingly, the single-plant dataset performed very poorly for the light-associated GO categories ‘*photosynthesis*’ and ‘*C_4_ photosynthesis*’ (Supplemental Data Set 9-10), even though several ‘*response to light stimulus*’ subcategories scored very well (Figure 6) and though our clustering analyses revealed several (bi)clusters heavily enriched in photosynthesis genes (see Supplemental Data Set 7). The performance plots show that the very poor function prediction performance for photosynthesis categories is due to the single-plant predictions having a very low precision compared to the predictions from the sampled datasets, while the number of predictions made by the single-plant data and their recall are comparatively very high (Supplemental Data Set 10). As argued above, recall values may be more indicative for the quality of gene function predictions than precision values, given the incompleteness of the maize GO annotation we use as a reference. If this is the case, genes that are predicted with high confidence to be involved in C_4_ photosynthesis but were scored as false positives by GO may still offer valuable leads. Indeed, we found evidence in literature linking three of the top-10 predicted regulators to C_4_ photosynthesis. *ZmCSP41A* (GRMZM2G111216, rank 1), a highly conserved sequence-specific chloroplast mRNA binding protein and unspecific endoribonuclease, was previously found to be more highly expressed in bundle sheet chloroplasts than in mesophyll chloroplasts (Friso et al., 2010). In the genus *Flaveria*, which contains C_3_ and C_4_ species as well as intermediates, a homolog of *ZmCSP41A* was found to be downregulated in leaves of C_4_ species compared to C_3_ species (Gowik et al., 2011). Transcripts of *ZmCRB* (GRMZM2G165655, rank 2), also accumulate preferentially in bundle sheet cells and are known to stabilize several chloroplast transcripts, e.g. for photosystem I and II components (John et al., 2014). *ZmSIG5* (GRMZM2G543629, rank 4) encodes a plastid sigma factor. Several homologous sigma factors in the *Flaveria* and *Cleome* genera were found to be upregulated in leaves of C_4_ species compared to C_3_ species (Gowik et al., 2011). Furthermore, six of the top-10 genes are known to be chloroplast-localized (GRMZM2G111216, GRMZM2G165655, GRMZM2G543629, GRMZM2G140288, GRMZM2G010929) or light-responsive (GRMZM2G158662), increasing the likelihood that they are involved in processes related to photosynthesis.

### Predicting phenotypic traits of individual plants from leaf transcriptome and metabolome data

We investigated to what extent the transcriptome and metabolome data generated on the individual plants can predict individual plant phenotypes. First, we performed spatially corrected correlation analyses (see Methods) to identify transcripts that show a significant linear association with a given phenotype (Supplemental Data Set 12). 1,677 genes exhibit an expression profile that is significantly correlated with leaf 16 blade length, and 411 gene expression profiles are significantly correlated (q≤0.05) with leaf 16 blade width. Notably, both for blade length and blade width, the set of significantly negatively correlated genes with *R*^2^ > 0.2 is enriched in known leaf and flower development genes (*q*<0.01, Supplemental Data Set 12). 273 genes exhibit an expression profile in mature leaf 16 tissue that is significantly correlated with ear length at *q*≤0.05 (Supplemental Data Set 12). Among those, the set of genes negatively correlated to ear length with *R*^2^ > 0.2 contains 3 genes known to be involved in cellular iron ion homeostasis (enrichment *q* = 8.56e−3), but no other significant GO enrichments were found. 241 genes have an expression profile that correlates significantly with husk leaf length (Supplemental Data Set 12). The set of genes whose expression in mature leaf 16 tissue positively correlates to husk leaf length (*q*≤0.05, *R*^2^ > 0.2) is enriched in genes involved in e.g. the response to oxidative stress, osmotic stress, UV stress and cell growth (*q*<0.01, Supplemental Data Set 12). Only 35 genes exhibited an expression profile in leaf 16 that is significantly correlated with plant height at *q*≤0.05, among which only 4 with an *R*^2^ value > 0.2, making plant height the phenotype that is least easily connected to the expression of individual genes in the leaf 16 blade.

The phenotypes of the individual plants can be predicted by the expression patterns of single genes in the leaf 16 blade with maximum *R*^2^ scores ranging from 0.509 (for blade length) to 0.291 (for plant height, Supplemental Data Set 12). We investigated whether combinations of genes could lead to a better prediction performance. Elastic net and random forest techniques were used to construct models predicting the phenotypes of individual plants as a function of the transcript and metabolite levels in the harvested leaf samples (see Methods). Elastic net (e-net) regression is a shrinkage method that is generally well-suited for use on high-dimensional datasets (Zou and Hastie, 2005). Its combination of the L1 and L2 penalties of its relatives lasso and ridge regression, respectively, makes e-net regression capable of selecting groups of correlated features (transcripts, metabolites) as predictors. Rather than selecting one representative feature from each group (as in lasso regression), e-nets can select multiple correlated features (as in ridge regression) while still setting the regression coefficients of irrelevant features to zero. This makes the resulting models more biologically interpretable. Random forest regression (Breiman, 2001) was used in addition because this technique can account for some types of interaction effects between features and is fairly robust to overfitting.

Both types of models were learned for each phenotype using either the transcript levels, the metabolite levels or both as features (see Table 2), each time using a 10-fold nested cross-validation strategy (see Methods). Transcript-based models were additionally run with either all transcripts or a pre-defined selection of regulatory transcripts as features (see Methods). The performance of the models was evaluated in two ways: by pooling the predictions for the test sets in each of the 10 folds into one dataset and computing the combined ‘out-of-bag’ (oob) *R*^2^ (pooled *R*^2^), and by computing the oob *R*^2^ on each test fold individually and taking the median (median *R*^2^, see Methods). For each model with a positive pooled or median *R*^2^ score, 500 datasets with permuted phenotype data were used to compute an empirical *p*-value that reflects whether the *R*^2^ score of the model is significantly higher than the *R*^2^ scores of models learned on randomized data (see Methods and Table 2).

**Table 2.** Performance of e-net and random forest models for phenotype prediction. Three different sections of the table show the pooled *R*^2^, median *R*^2^ and Pearson correlation (PCC) measures for the prediction performance of the models learned for all phenotypes using all transcripts (Transcripts), only regulatory transcripts (Regulators), all metabolites (Metabolites), and both transcripts and metabolites (Both) as features. Numbers between parentheses indicate *p*-values for the oob *R*^2^ values obtained, derived from permutation tests. No permutation tests were performed for the ear length and plant height phenotypes, given the poor oob *R*^2^ values of the models concerned.

| Pooled $R^2$ | | | | | |
| --- | --- | --- | --- | --- | --- |
| Trait |  | Transcripts | Regulators | Metabolites | Both |
| Blade 16 length | Elastic Net | 0.345 (0.002) | 0.409 (0.002) | 0.412 (0.002) | 0.226 (0.004) |
|  | Random Forest | 0.567 (0.002) | 0.609 (0.002) | 0.323 (0.004) | 0.474 (0.002) |
| Blade 16 width | Elastic Net | 0.659 (0.002) | 0.582 (0.002) | 0.611 (0.002) | 0.622 (0.002) |
|  | Random Forest | 0.274 (0.002) | 0.312 (0.002) | 0.388 (0.002) | 0.229 (0.004) |
| Husk leaf length | Elastic Net | 0.438 (0.002) | 0.445 (0.002) | 0.519 (0.002) | 0.412 (0.002) |
|  | Random Forest | 0.274 (0.002) | 0.337 (0.002) | 0.461 (0.002) | 0.315 (0.002) |
| Ear length | Elastic Net | 0.057 | 0.084 | 0.082 | 0.129 |
|  | Random Forest | 0.115 | 0.091 | 0.162 | 0.137 |
| Plant height | Elastic Net | -0.058 | -0.045 | -0.036 | -0.043 |
|  | Random Forest | -0.061 | -0.059 | 0.008 | -0.096 |
| Median $R^2$ | | | | | |
| Trait |  | Transcripts | Regulators | Metabolites | Both |
| Blade 16 length | Elastic Net | 0.234 (0.002) | 0.318 (0.002) | 0.314 (0.002) | -0.199 (0.471) |
|  | Random Forest | 0.531 (0.002) | 0.534 (0.002) | 0.325 (0.004) | 0.379 (0.002) |
| Blade 16 width | Elastic Net | 0.726 (0.002) | 0.582 (0.002) | 0.481 (0.002) | 0.641 (0.002) |
|  | Random Forest | 0.279 (0.002) | 0.322 (0.002) | 0.521 (0.002) | 0.192 (0.004) |
| Husk leaf length | Elastic Net | 0.501 (0.002) | 0.471 (0.002) | 0.295 (0.002) | 0.392 (0.002) |
|  | Random Forest | 0.280 (0.002) | 0.267 (0.002) | 0.431 (0.002) | 0.197 (0.002) |
| Ear length | Elastic Net | 0.149 | 0.129 | -0.045 | -0.054 |
|  | Random Forest | 0.122 | 0.029 | -0.047 | -0.154 |
| Plant height | Elastic Net | -0.205 | -0.169 | -0.433 | -0.336 |
|  | Random Forest | -0.213 | -0.261 | -0.291 | -0.375 |
| PCC |  |  |  |  |  |
| Trait |  | Transcripts | Regulators | Metabolites | Both |
| Blade 16 length | Elastic Net | 0.592 | 0.641 | 0.645 | 0.477 |
|  | Random Forest | 0.782 | 0.789 | 0.586 | 0.726 |
| Blade 16 width | Elastic Net | 0.821 | 0.765 | 0.784 | 0.808 |
|  | Random Forest | 0.560 | 0.573 | 0.664 | 0.542 |
| Husk leaf length | Elastic Net | 0.670 | 0.669 | 0.734 | 0.665 |
|  | Random Forest | 0.605 | 0.630 | 0.704 | 0.638 |
| Ear length | Elastic Net | 0.295 | 0.304 | 0.299 | 0.361 |
|  | Random Forest | 0.341 | 0.305 | 0.404 | 0.376 |
| Plant height | Elastic Net | -0.168 | -0.161 | 0.106 | -0.091 |
|  | Random Forest | 0.021 | 0.053 | 0.160 | -0.105 |

The blade length and blade width of leaf 16 (the ear leaf) are the phenotypes that are best predictable from both the transcriptome and metabolome data (Table 2 and Supplemental Data Sets 13-14). This is not surprising, as these phenotypes are most closely related to the plant material that was profiled (mature leaf 16 blade tissue). The whole-transcriptome e-net model for leaf 16 blade width reached a pooled *R*^2^ score of 0.659, whereas the ordinary least squares (OLS) *R*^2^ value for the best-correlated single gene is only 0.463 (Supplemental Data Set 12). This indicates that the multi-gene model for blade width performs substantially better than single-gene models. The performance difference is likely even higher than suggested by the *R*^2^ difference, as single gene models have an advantage in this comparison: multi-gene model *R*^2^ values are based on test data while single-gene model *R*^2^ values are based on training data.

The best-performing whole-transcriptome model for leaf 16 blade length on the other hand has a pooled *R*^2^ score that is only marginally higher than the OLS *R*^2^ value for the best-correlated single gene (pooled *R*^2^ = 0.567 for the whole-transcriptome random forest model versus OLS *R*^2^ = 0.509 for the gene GRMZM2G553379, *ZMM15*, Supplemental Data Set 12). This suggests that maybe only few genes contribute substantially to the random forest model performance. Indeed, next to the aforementioned gene *ZMM15*, only one other gene, *ZAP1* (GRMZM2G148693), has a median importance score above 0.05 in the random forest model for leaf 16 blade length (Supplemental Data Set 13). Like *ZMM15*, *ZAP1* is found in the top-10 of genes that are most significantly anticorrelated with blade length (Supplemental Data Set 12, see below for model interpretation).

The models for ear length and plant height have considerably lower oob *R*² scores than for the leaf 16-related phenotypes, and for plant height even negative *R*² scores were obtained (Table 2). This suggests that the transcriptome of the sampled leaves may not contain sufficient information to accurately predict phenotypes measured on other organs at the time of sampling (see also Discussion). Tellingly, the multi-gene model oob *R*² scores for both ear length and plant height are much lower than the best single-gene OLS *R*² scores, suggesting that the multi-gene models severely overfit the training data (Supplemental Data Set 12). Husk leaf length on the other hand is predicted almost equally well as the leaf 16 phenotypes (whole-transcriptome e-net model, pooled *R*^2^ = 0.438, Table 2). This may be due to the phenotype being closer to the material that was molecularly profiled, in terms of tissue type or spatial proximity, than ear length and plant height. However, the best multi-gene model for husk leaf length merely performs on par with the best single-gene model (OLS *R*^2^ = 0.460, Supplemental Data Set 12). In contrast to what was found for leaf 16 blade length, this is not because only a few genes contribute to the e-net model performance for husk leaf length (Supplemental Data Set 15).

In general, the models learned on transcriptome and metabolome data have similar performance for most phenotypes (Table 2). This suggests that both datasets contain roughly the same amount of information on the phenotypes, despite the fact that there are many more transcripts (18,171) than metabolites (592) in the data. Surprisingly, the models learned on both data sources combined did not outperform the models learned on the transcriptome or metabolome data separately. This suggests that most of the relevant phenotype information is redundantly present in both data types. Interestingly, the models learned on the transcriptome data using only the transcript levels of regulatory genes as features performed generally on par with the overall transcriptome models (Table 2). This indicates that using the expression levels of regulatory genes as features may be sufficient to obtain adequate phenotype predictors, with the advantage that the predictors obtained may be more interpretable from a mechanistic perspective.

We took a closer look at the best-performing transcriptome models for the blade length and blade width phenotypes. For blade length, the best-performing model is the random forest model with only regulators as predictors, with a median *R*^2^ score of 0.534 and a pooled *R*^2^ score of 0.609 (Figure 7). The two regulators with the highest variable importance in this model are the same as the two most important genes in the whole-transcriptome model, GRMZM2G148693 (*ZAP1*) and GRMZM2G553379 (*ZMM15*) (Supplemental Data Set 13). Both are MADS-box transcription factors homologous to the *A. thaliana* gene *APETALA1*, and they exhibit a Pearson expression correlation of 0.79, which explains why one of the two was given a higher importance score (the second one contains largely redundant information). Their correlation with blade length is negative and strong. The heavy reliance of both the whole transcriptome and regulator random forest models on either of these two genes also helps explain why the predicted blade length values in Figure 7 exhibit a distinctly bimodal distribution. Interestingly, *ZAP1* was previously found in QTL and GWA studies as a candidate gene associated with ear length (Xue et al., 2016), ear height (Vanous et al., 2018), tassel length (Wang et al., 2018a) and flowering time (Wallace et al., 2016), and it has been implicated in maize domestication, in particular for temperate maize lines, in which its expression is downregulated (Liu et al., 2015a).

**Figure 7.**
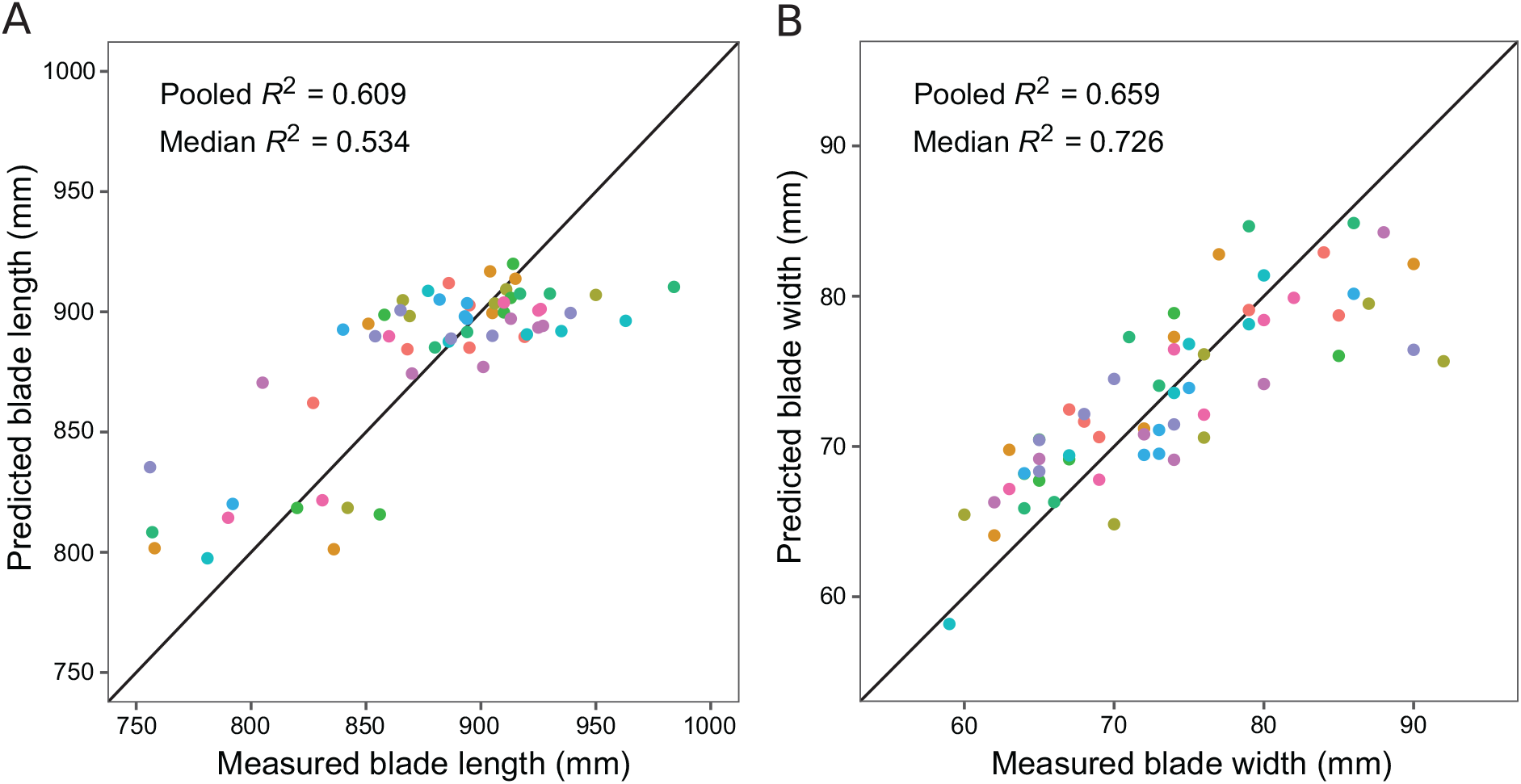
Predictive models for leaf 16 blade length and width. Graphs plotting predicted versus measured phenotypes are shown for **(A)** the random forest model for leaf 16 blade length using only transcript levels of regulators as predictors, and **(B)** the e-net model for leaf 16 blade width using only transcript levels of regulators as predictors. The dot colors represent different outer cross-validation folds. Perfect predictions are located on the diagonal line in each panel.

For blade width, the e-net model built on all transcripts performed best (median *R*² = 0.726, pooled *R*^2^ = 0.659). 235 transcripts have a median coefficient >0.01 in this elastic net model (Supplemental Data Set 14), but no significant GO enrichments were found in the corresponding gene set. In the e-net model for blade width run with only regulators as predictors (Figure 7), 178 transcripts have a median coefficient above 0.01 (Supplemental Data Set 14). The regulators with the strongest negative influence are GRMZM2G023625, a putative HIRA histone chaperone, and GRMZM2G377311, a putative cyclin T. The only *A. thaliana* homolog of GRMZM2G023625, *AT3G44530* (*HIRA*), is known to be involved in *knox* gene silencing during leaf development, and reduced *HIRA* expression levels give rise to transversally curled (pinched) leaves with shorter petioles and often lobes in the proximal region of the blade (Phelps-Durr et al., 2005). The *A. thaliana* homologs of GRMZM2G377311 with the highest sequence similarity, *AT4G19600* (*CYCT1;4*) and *AT5G45190* (*CYCT1;5*), have been implicated previously in the regulation of leaf and flower development (Cui et al., 2007). The two regulators with the strongest positive predicted influence on blade width in the e-net model were GRMZM2G062914 (*MAP KINASE 14*, *MPK14*) and GRMZM2G430780, a putative serine/threonine protein kinase.

## DISCUSSION

In this study, we molecularly and phenotypically profiled 60 individual maize plants of the same inbred line (B104) grown in the same field. Our purpose was to investigate how much information can be extracted from this simple experimental design on the function of genes, and on how gene and metabolite expression relates to plant phenotypes. Although one may expect that this design should yield datasets with a low information content, due to the very limited genetic and environmental variability employed, substantial variability was found in the transcriptomes, metabolomes and phenotypes of the individual plants. Standard deviations on the transcript and metabolite levels across the field were found to be generally in the order of 10-50% of the mean. The average transcript level CV of ~0.3 is about three times higher than the transcript level CV of lab-grown *A. thaliana* plants in a recent study (Cortijo et al., 2019). Genes involved in processes such as photosynthesis and herbivory responses were found to be more variably expressed across the field than housekeeping genes involved in e.g. RNA and protein metabolism, and the expression patterns of 12.1% of the transcripts and 7.1% of the metabolites profiled exhibited significant spatial patterning, indicating that the variability uncovered is not merely random noise.

We used the single-plant dataset to predict the function of maize genes from the function of their coexpression network neighbors (‘guilt-by-association’), and found that field-grown single-plant transcriptomes overall have similar gene function prediction power as traditional transcriptome datasets profiling pooled plant responses to controlled perturbations in a lab. Furthermore, the single-plant dataset was found to outperform the controlled perturbation datasets for several processes that were likely variably active in the field setting used, in particular abiotic stress responses. This suggests that datasets in which processes are perturbed more subtly around a common baseline may hold an advantage for unraveling gene functions. One of the issues with harsher perturbations is that their effects may propagate further in the cellular networks, and essentially swamp more subtle variations in other, sideways associated processes, decreasing the information content of the resulting data. Pooling samples, although enhancing experimental repeatability, may similarly decrease the data information content by smoothing out subtle variations across samples.

Comparable results were obtained in an earlier study on individual lab-grown *A. thaliana* plants (Bhosale et al., 2013). One notable difference with the Arabidopsis results however is that the maize single-plant dataset performs better at predicting gene functions than most of the traditional transcriptome datasets it is compared to at higher (less stringent) *q*-value thresholds, whereas it performs worse at lower *q*-value thresholds. The opposite trend was observed in Arabidopsis (Bhosale et al., 2013). This is because, taking the precision of predictions from the traditional datasets as a baseline for both species, a disproportionately large fraction of the high-confidence predictions emerging from the maize single-plant dataset are not supported by existing maize gene function annotations. One potential reason for this surplus of high-confidence ‘false positives’ is that the maize single-plant dataset, in contrast to the other maize and Arabidopsis datasets, was generated in a field setting. It is not unthinkable that lab and field experiments may profile different aspects of gene function, and therefore lead to complementary predictions. This may help explain why the lab-generated datasets lead to high-confidence predictions that are more closely aligned with known gene function annotations, as most of these were also derived directly or indirectly (in the case of annotations transferred by orthology from other plant species) from lab experiments. If lab and field experiments indeed profile complementary aspects of gene function, the novel gene function predictions obtained from field-generated data could be as valuable as those from lab-generated data. Confirming the potential value of the novel predictions generated by our field dataset, we found indirect evidence in literature in support of more than 30% of the top-10 novel regulator predictions obtained for C_4_ photosynthesis, the response to chitin and the response to water deprivation.

Our results indicate that profiling individual plants in the field may also be useful to identify genes that influence plant phenotypes under field conditions. We used machine learning models to quantitatively predict phenotypes of individual plants based on leaf gene expression and metabolome data, and found that leaf phenotypes could be predicted reasonably well, in particular the blade width of leaf 16 (max. median oob *R*^2^ score = 0.726, max. Pearson correlation (PCC) between predicted and observed values = 0.821). This is fairly remarkable given that the models were learned on data for only 60 plants. For comparison, a recent study in which maize phenotypes were predicted from genetic marker and transcriptome data for 388 different maize lines reported PCC values of 0.56 to 0.66 between predicted and measured phenotypes when using both genetic markers and transcript levels as features, and PCC values of 0.51 to 0.61 when using only transcript levels as features (Azodi et al., 2020). An important difference however is that the (Azodi et al., 2020) study predicted mature plant phenotypes (final plant height, final yield, flowering time) from seedling data, whereas we predicted actively developing phenotypes from contemporarily profiled leaf transcriptome data. Whereas we could generate decent predictive models for phenotypes that were closely related to the plant material that was molecularly profiled (length and width of the ear leaf blade, and to a lesser extent the length of the developing husk leaf), models learned for more distant phenotypes such as plant height and ear length at sampling time did not perform well. This discrepancy between the (Azodi et al., 2020) study and ours suggests that intermediate phenotypes may be inherently less predictable than final phenotypes, unless the plant material profiled is directly associated with the phenotype under study. Follow-up experiments will be necessary to assess whether individual plant datasets can be used as efficiently as genomic prediction datasets (Azodi et al., 2020) for predicting final plant phenotypes from molecular data profiled at an earlier developmental stage.

Together, our results show that profiling individual plants in the field is a promising addition to the toolbox we have at our disposal to study the molecular wiring of plants and relationships between genes and phenotypes, in particular in a field context. More steps will have to be taken however to realize the full potential of this new experimental design. A major bottleneck in all transcriptome profiling-based strategies to associate genes with phenotypes, not only the single-plant setup but also TWAS and classical systems biology strategies, is that the models they produce are correlational rather than causal in nature. A shift to more causal modeling approaches is direly needed, but not straightforward, as causal inference from the high-dimensional datasets generated by transcriptome profiling, which are frequently observational in nature and contain lots of hidden variables and confounders, is notoriously difficult. Profiling additional data layers in the single-plant setup, such as micro-environmental variables, may further improve modeling performance and enhance causal interpretability.

Up to now, we only profiled a limited amount of plants of one cultivar in one season and field environment. It remains to be seen to what extent the resulting models can be generalized to other cultivars and growth environments. The fact that the single-plant setup only profiles one specific cultivar at a time may be seen as a disadvantage with respect to the classical TWAS setup, in which multiple cultivars are modeled simultaneously. On the other hand, as the phenotypic effects of expression variants often depend on the genetic background (epistasis) and environment in which they are introduced, it might in fact make sense to study the molecular wiring of a trait in a specific cultivar and environment before attempting generalizations to other cultivars or growth environments, in particular for plant species with large pan-genomes such as maize (Gore et al., 2009; Hirsch et al., 2014; Lu et al., 2015). The single-plant setup might for instance be used for studying an elite cultivar directly in a target field environment in which yield or stress tolerance improvements are desired.

## METHODS

### Field trial setup, sampling and phenotyping

During the summer of 2015, 560 B104 maize inbred plants were grown under ‘uncontrolled’ field conditions at a site in Zwijnaarde, Belgium (51°00’35.2”N, 3°42’56.5”E) with a sowing density of approximately 177,778 plants per hectare. Plants were sown by hand in ten adjacent rows of 5.5 m length, 75 cm apart and each containing 56 maize B104 plants. To the North and West of the B104 plants the commercial hybrid ‘Ricardino’ was sown, while to the East more B104 plants were grown and to the South other hybrids and recombinant inbred lines were grown, separated from the B104 plants by a 2.5 m-wide path (Figure 1A).

In total, 200 non-border plants that exhibited a primary ear at leaf 16 were harvested at the VT (tasseling) stage. Since not all plants reached this stage at the same time, plants were harvested on two different dates, 2015-08-25 (164 plants) and 2015-09-02 (36 plants). On each of these days, harvesting and sampling occurred from 10 am until noon. Damaged plants were discarded to avoid outliers in the data. The position in the field was recorded for the harvested plants, and plant height was measured from the plant base to the collar of the top leaf. The primary ear leaf (leaf 16) of each selected plant was cut off at the ligule. Leaf 16 blade length was measured from the ligule to the tip of the leaf while leaf 16 blade width was measured in the middle between the ligule and the leaf tip. For molecular data generation, a 10 cm-long part of the leaf was cut from the middle of the leaf 16 blade, the midrib was removed (to avoid detection of exogenous metabolites during untargeted metabolite profiling) and the resulting mature leaf samples were stored in liquid nitrogen on the field. Primary ears were also cut off from the plants, and the length of the ears and husk leaves (from base to tip) was measured on the field.

### RNA sequencing

Sixty of the 200 leaf samples for individual plants were randomly selected for RNA sequencing. Total RNA was isolated with the guanidinium thiocyanate-phenol-chloroform extraction method using TRI-reagent (Sigma-Aldrich). Total RNA was sent to GATC Biotech for RNA-sequencing. Library preparation was done using the NEBNext Kit (Illumina). In brief, purified poly(A)-containing mRNA molecules were fragmented, randomly primed strand-specific cDNA was generated and adapters were ligated. After quality control using an Advanced Analytical Technologies Fragment Analyzer, clusters were generated through amplification using cBOT (Cluster Kit v4, Illumina), followed by sequencing on an Illumina HiSeq2500 with the TruSeq SBS Kit v3 (Illumina). Sequencing was performed in paired-end mode with a read length of 125◻bp.

The raw RNA-seq data was processed using a custom Galaxy pipeline (Goecks et al., 2010) implementing the following steps. First, the fastq files were quality-checked using FastQC (v:0.5.1) (Andrews, 2010). Next, Trimmomatic (v:0.32.1) (Bolger et al., 2014) was used to remove adapters, read fragments with average quality below 10 and trimmed reads shorter than 20 base pairs. The trimmed and filtered reads were mapped against the *Zea mays* AGP genome annotation v:3.23 (Schnable et al., 2009) using GSNAP v:2013-06-27 (Wu and Nacu, 2010). A k-mer size of 12 was used, the ‘local novel splicing event’ parameter was set to 50,000, and default values were used for the rest of the parameters. The option for splitting the bam files into unique and multiple alignments was activated, and only the uniquely mapping reads were kept for the following analyses. The mapping files were quantified using HTSeq v:0.6.1p1 (Anders et al., 2015) with the option ‘Intersection-strict’ and using the *Zea mays* AGP genome annotation v:3.23 (Schnable et al., 2009). The resulting raw counts were filtered to only keep genes with at least 5 counts per million in at least 1 sample. Then, raw counts were divided by size factors calculated by DEseq2 (v:1.14.1) (Love et al., 2014), resulting in library size-corrected gene expression values for 18,171 genes across 60 plants. Pseudocounts of 0.5δ, with δ the smallest non-zero value in the normalized expression matrix, were added to all gene expression values. For all downstream analyses except coefficient of variation (CV) calculations, the resulting expression matrix was log_2_-transformed.

### Metabolome Profiling

Fifty of the 60 leaf samples selected for RNA sequencing were additionally metabolome-profiled. For metabolome analysis, 100 mg of frozen, grinded mature leaf 16 material for the selected maize plants was sent to Metabolon Inc. (Durham, NC, USA). Sample extracts were prepared using the automated MicroLab STAR® system from Hamilton Company and divided into five fractions. Samples were normalized based on dry weight and further processed and analyzed by Metabolon for untargeted metabolic profiling involving a combination of four independent approaches: two separate reverse phase (RP)/UPLC-MS/MS analyses with positive ion mode electrospray ionization (ESI), RP/UPLC-MS/MS analysis with negative ion mode ESI and HILIC/UPLC-MS/MS analysis with negative ion mode ESI. All methods utilized a Waters ACQUITY ultra-performance liquid chromatographer (UPLC) and a Thermo Scientific Q-Exactive high resolution/accuracy mass spectrometer interfaced with a heated electrospray ionization (HESI-II) source and an Orbitrap mass analyzer operated at a mass resolution of 35,000. Sample extracts were dried and then reconstituted in solvents compatible to each of the four methods. Each reconstitution solvent contained a series of standards at fixed concentrations to ensure injection and chromatographic consistency. One aliquot was analyzed using acidic positive ion conditions, chromatographically optimized for more hydrophilic compounds. In this method, the extract was gradient eluted from a C18 column (Waters UPLC BEH C18-2.1×100 mm, 1.7 µm) using water and methanol, containing 0.05% perfluoropentanoic acid (PFPA) and 0.1% formic acid (FA). Another aliquot was analyzed using acidic positive ion conditions, chromatographically optimized for more hydrophobic compounds. In this method, the extract was gradient eluted from the same aforementioned C18 column using methanol, acetonitrile, water, 0.05% PFPA and 0.01% FA and was operated at an overall higher organic content. Another aliquot was analyzed using basic negative ion optimized conditions using a separate dedicated C18 column. The basic extracts were gradient eluted from the column using methanol and water, however with 6.5mM Ammonium Bicarbonate at pH 8. The fourth aliquot was analyzed via negative ionization following elution from a HILIC column (Waters UPLC BEH Amide 2.1×150 mm, 1.7 µm) using a gradient consisting of water and acetonitrile with 10mM Ammonium Formate, pH 10.8. The MS analyses alternated between MS and data-dependent MS scans using dynamic exclusion. The scan range varied slighted between methods but covered 70-1,000 m/z. Raw data was extracted, peak-identified and QC processed using Metabolon’s hardware and software. Compounds were identified by comparison to library entries of more than 3,300 purified standards or recurrent unknown entities. Metabolon’s library was based on authenticated standards that contain the retention time/index (RI), mass to charge ratio (m/z), and chromatographic data (including MS/MS spectral data) of all molecules present in the library.

The metabolite profiles used in the downstream analyses were obtained from the raw data delivered by Metabolon Inc. as follows. Log_2_ transformation was applied to the initial matrix containing the levels of 601 metabolites across 50 samples. Outliers were identified iteratively using two-tailed Grubbs tests (threshold for outlier detection was *p* = 0.01) and converted to missing values (NA). Metabolites with missing values for more than half of the samples were removed, resulting in a matrix containing the levels of 592 metabolites across 50 samples. To deal with residual missing values, imputation was performed using Bayesian principal component analysis (BPCA) with 48 components (using the pca function of the pcaMethods R package, v:1.76.0 with method=”bpca”, scaling=”uv” (unit variance), npcs=48). Finally, quantile normalization was applied to give each sample the same data distribution. This matrix was used for downstream analysis, except for CV calculations where the raw metabolite values were used instead.

### Clustering analyses

The transcriptome and metabolome datasets were *z*-scored and jointly clustered using the ward.D2 hierarchical clustering method (Murtagh and Legendre, 2014) included in the R stats package (v:3.6.0), and using squared Euclidean distance as the distance measure. The same protocol was used for clustering the RNA-seq datasets sampled from the Short Read Archive v. 2018/01/30 (Leinonen et al., 2011) (see further). Additionally, the single-plant transcriptome dataset was analyzed using the biclustering algorithms ISA (Bergmann et al., 2003), SAMBA (Tanay et al., 2002), both part of EXPANDER v:7.1 (Hait et al., 2019), and ENIGMA v:1.1 (Maere et al., 2008). For biclustering, the log_2_ expression values were transformed to log_2_ fold changes with respect to the mean log_2_ gene expression across the individual plants. Default parameters were used for running ISA. For SAMBA, default parameter settings were used except for the setting ‘*use option files of type*’ = valsp_3ap. For ENIGMA, default parameters were used, except for ‘*fdr*’=0.001, ‘*fdrBiNGO*’=0.01, ‘*namespaces*’=biological_process and ‘*pvalThreshold*’ = 0.6296976. The latter threshold is the standard deviation of the log_2_ fold changes across the entire RNA-seq dataset, which, by lack of differential expression *p*-values for the single plants, is used by ENIGMA as a threshold for discretizing transcript log_2_ fold changes into the categories ‘upregulated’, ‘downregulated’ and ‘unchanged’.

### Gene Ontology (GO) enrichment analyses

The gene ontology file used for GO enrichment analyses was downloaded on 30^th^ August 2016 from the Gene Ontology website (The Gene Ontology Consortium, 2017). A GO annotation file for AGP maize genome version 3.23 was parsed from the functional annotations provided by PLAZA (Proost et al., 2015), development version cnb 02, on 27^th^ November 2017. To ensure that all the functional annotations found for the genes in the AGP maize genome version 2 were included in our analyses, we also included the maize gene functional annotations provided by the older PLAZA 3.0 platform (Proost et al., 2015), taking into account gene identifier changes from maize genome version 2 to version 3 as recorded in MaizeGDB (Portwood et al., 2018). Given the lack of maize genes annotated to the C_4_ photosynthesis category in GO, we manually added annotations to this category for 78 genes identified as C_4_ genes by Li et al. (2010). In all GO enrichment analyses, enrichment *p*-values were calculated using hypergeometric tests and adjusted for multiple testing (*q*-values) using the Benjamini-Hochberg (BH) procedure (Benjamini and Hochberg, 1995), which controls the false discovery rate (FDR). For GO enrichment analyses on (bi)clustering results, multiple testing correction was done for each cluster separately. Genes annotated to the categories ‘*DNA binding transcription factor activity’* (GO:0003700), ‘*signal transducer activity*’ (GO:0004871) and ‘*regulation of transcription – DNA-templated*’ (GO:0006355) were combined in a list of potential regulators (Supplemental Data Set 16), for use in the ENIGMA analysis, the literature screen for evidence supporting our gene function predictions, and some of the phenotype prediction models, namely those that use a predefined list of regulators as potential predictors (see further).

### Spatial autocorrelation analyses and correlation network generation

Spatially autocorrelated transcripts, metabolites and phenotypes were detected using Moran’s I with an inverse distance-weighted matrix in the Ape package (v:5.2) in R (v:3.6.0) (Paradis and Schliep, 2018). The *p*-values computed by the Ape package were adjusted for multiple testing with the BH method. The *z*-scored profiles of all transcripts with *q* ≤ 0.01 were assigned to clusters using the Tight Clustering algorithm (Tseng and Wong, 2005) (parameters: seed = 1, kmin = 35, nstart = 50, resamp = 10). Associations between a given spatially autocorrelated transcript cluster and any phenotypes were assessed by testing for Pearson correlation between the average *z*-scored gene expression profile of the cluster and the phenotype profiles. The resulting *p*-values were corrected per phenotype using the BH method.

For each pair of genes *x* and *y* in the single-plant transcriptome dataset, a ‘spatially adjusted Pearson correlation’ was computed by *z*-scoring the log_2_ gene expression profiles of both genes and fitting the following model to the data:

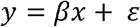

with *β* the correlation coefficient and *ɛ* an error term with a spherical covariance structure. That is, *ɛ* is assumed to follow a 60-dimensional (= number of plant samples) multivariate normal distribution with mean zero and a covariance matrix given by:

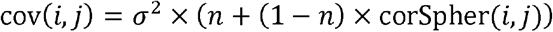

where *σ* is the magnitude of the noise (comparable to the standard deviation of an independent normal distribution), the nugget *n* determines which proportion of the residuals is governed by spatial auto-covariance, and is given by:

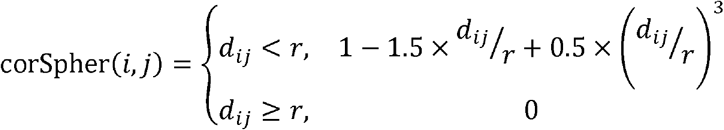

with *d_ij_* the physical distance between plants *i* an *j* in the field. The range parameter *r* is related to the distance at which two plants become independent of one another. The spherical covariance structure was chosen as it gave the most meaningful range estimates (within bounds of the field when *n* ≠ 1) and the best overall performance as measured by the Bayesian Information Criterion (BIC). All four parameters (*β*, *r*, *n*, *σ*) were optimized with restricted maximum likelihood optimization using the nlme package (Linear and Nonlinear Mixed Effects Models, v:3.1-140) (Pinheiro et al., 2019) in R (v:3.6.0). Although there is an asymmetry in the regression equation, swapping *x* and *y* for gene pairs with a range estimate *r* above zero gave parameter estimates that were not meaningfully different.

For most gene pairs *r* converged to zero or *n* converged to 1, which means the best-fit model is one without spatial covariance, yielding the exact same correlation coefficient *β* and corresponding *p*-value as a normal ordinary least-squares (OLS) regression or Pearson correlation on the *z*-scored variables (up to rounding errors). Only for about 10% of the gene pairs, *r* converged to a non-zero distance. This means that for about 10% of gene pairs, there would be spatial structure left in the residuals of an OLS regression, violating the assumption of independence in OLS regression. All *p*-values were Bonferroni-corrected, and correlations with corrected *p*-values ≤ 0.01 were included as edges in the correlation network.

The correlation network obtained from the single-plant datasets was compared with networks obtained from traditional RNA-seq datasets sampled from the Short Read Archive v. 2018/01/30 (Leinonen et al., 2011). The raw RNA-seq data downloaded from the SRA in first instance involved all transcriptome data on *Zea mays* profiled with Illumina sequencing platforms. Only runs profiling mRNA (as opposed to e.g. small RNAs) with an average read length > 30 bp and ≥ 4.10^6^ reads were retained. In many cases, the meta-information obtained from SRA did not specify the genotype and tissue profiled in the RNA-seq experiments. We therefore used information from the BioSample database (https://www.ebi.ac.uk/biosamples/, v. 2018/02/28) to select only RNA-seq datasets produced on leaves of the maize inbred line B73, discarding crosses, mutants and NILS. Only samples with a unique BioSample ID were retained to avoid data replication. This led to a compendium of 470 unique RNA-seq samples (Supplemental Data Set 8), which were preprocessed and normalized in the same way as the single-plant samples. As an additional data quality filtering step, samples with <80% uniquely mapping reads, samples with a clearly divergent data distribution and samples with less than 20,000 expressed genes were discarded. This resulted in a compendium of 407 RNA-seq samples, which we randomly sampled without replacement to extract 500 compendia of 60 samples. For each of these randomly sampled compendia, a correlation network was built using Pearson correlation. Note that in contrast to the single-plant dataset, spatial autocorrelation correction is not necessary for the datasets sampled from SRA. Every sampled network was thresholded to obtain the same number of edges as obtained for the single-plant network.

### Gene function prediction

Gene functions (GO Biological Process annotations) were predicted from the single-plant correlation network and all 500 sampled networks using a command-line version of PiNGO (v:1.11) (Smoot et al., 2011). PiNGO predicts the function of a given gene based on the GO annotations of its neighbors in a given network, using hypergeometric GO enrichment tests on the gene’s network neighborhood. The resulting *p*-values were adjusted for multiple testing (for each input network separately) using the BH method. The overall function prediction performance of the single-plant and sampled networks was calculated as in (Bhosale et al., 2013). Recall and precision of the functional predictions for a given gene in a given network were calculated as described by (Deng et al., 2004) using the known maize GO annotations as gold standard, and the overall recall and precision values for the given network were obtained by averaging across all genes in the network. Next to this overall analysis of gene function prediction performance, we also assessed how accurately the networks predicted genes involved in specific GO Biological Processes. For these analyses, recall (*R*) and precision (*P*) were calculated in the traditional way as *R* = *tp*(*tp* + *fn*) and *P* = *tp*(*tp* + *fp*) with *tp* the number of true positives, *fp* the number of false positives and *fn* the number of false negatives identified.

For every GO category and overall, the recall, precision, and *F*-measure (harmonic mean of recall and precision) of the predictions were calculated for every network at prediction *q*-value thresholds ranging from 10^−2^ to 10^−11^. Undefined precisions and F-measures, resulting from a network not producing any predictions at a given *q*-value threshold, were set to 0 in order to reflect poor performance of the network at the *q*-value concerned. The relative prediction performance of the single-plant network with respect to the sampled networks was classified as very good, good, average, poor, or very poor based on the root mean square deviation of the single-plant network *F*-measures from the 25th, 50th, and 75th percentiles of the sampled network *F*-measures over the FDR subrange in which either the single-plant network or at least 250 of the 500 sampled networks, or both, exhibited non-zero *F*-measures.

### Predictive models for phenotypes

Phenotypes were regressed on the expression of single genes using a mixed model with the following formulation:

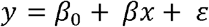

with *x* the log_2_ expression of a given gene and *y* the phenotype value. The error *ɛ* is assumed to follow a multivariate normal distribution with a rational quadratic distance-based covariance function. That is, the covariance of *ɛ* is described by:

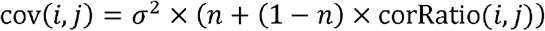

Where *σ* is the magnitude of the noise and *n* determines which proportion of the residuals is governed by spatial auto-covariance. The correlation function *corRatio*(*i, j*) between two samples *i* and *j* is given by:

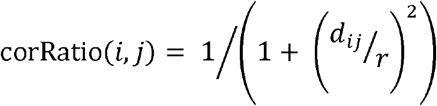

with *d_ij_* the physical distance between plants *i* an *j* in the field. The range parameter *r* is related to the distance at which two plants become independent of one another. The ratio kernel was chosen because it gave meaningful range estimates (Supplemental Figure 6) and the best overall performance as measured by BIC. Regression analyses were performed using the nlme package (v:3.1-140)(Pinheiro et al., 2019) in R (v:3.6.0). *p*-values were adjusted for each phenotype separately using the BH method.

Elastic net and random forest methods were used to learn multi-feature predictive models for the phenotypes using transcript levels, metabolite levels or both as features. Elastic net and random forest models were also built using as features only the transcript levels of a predefined set of regulators (Supplemental Data Set 16). Both types of models were built with the scikit-learn package (v:0.21.0) (Pedregosa et al., 2011) in Python. For elastic net models, the maximum number of iterations (parameter ‘*max_iter*’) was set to 10^6^. For random forest models, the number of estimators, i.e. the number of averaged trees, was set to 500, the ‘*criterion*’ parameter was set to ‘mse’ and the ‘*bootstrap*’ parameter was set to ‘True’. For each phenotype, models were built with each method on each feature set using 10-fold nested cross-validation. For each of the 10 outer folds, 4 inner folds were used to tune the model hyperparameters (the shrinkage parameter α and the L1-ratio ρ for elastic nets; the ‘*max_features*’ parameter with possible values ‘sqrt’, 0.33, ‘log2’ and ‘None’ and the ‘*min_samples_split*’ parameter with possible value 2, 3, 4 and 5 for random forests). After completing the inner cross-validation, the combination of hyperparameters that scored best on test data across the 4 folds were used to retrain the model on all 4 folds combined, yielding 10 trained models with optimized hyperparameters per phenotype (GridSearchCV function in scikit-learn). Each of the 10 models was used to predict the phenotypes of the 6 hold-out samples for the fold it was trained on, yielding 60 ‘test data’ predictions in total, one for each sample.

The ‘out-of-bag’ (oob) *R*^2^ score, defined as 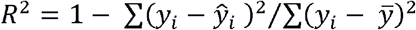 where 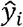 and *y_i_* are the predicted and observed phenotypes for sample *i*, respectively, and where 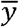 is the mean of the observed phenotypes, was used to measure how well the predictions align with the true phenotypes. Note that the meaning of this oob *R*^2^ is different from the classical meaning of *R*^2^, which is the percentage of variance explained by a linear model. As opposed to the classical *R*^2^, the oob *R*^2^ can become negative when the sum of squared errors (numerator) is larger than the variance of the data (denominator). When all predictions 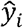 equal the mean 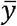, the oob *R*^2^ equals zero. A negative oob *R*^2^ score indicates that the model does worse than assigning the mean phenotype value of the training samples to the unseen samples. Positive oob *R*^2^ scores indicate that the model does better than predicting the mean, and a model that perfectly predicts the unseen phenotypes has an oob *R*^2^ score of one. We report two oob *R*^2^ scores for each model, the ‘pooled *R*^2^’ score and the ‘median *R*^2^’ score. For calculating the pooled *R*^2^, the test set predictions of all folds were taken into account together to calculate one oob *R*^2^ value that summarizes all folds. The ‘median *R*^2^’ score is the median of the oob *R*^2^ scores calculated for each fold independently.

For modeling methods that use built-in feature selection/reduction techniques, such as elastic nets and random forests, an analytical statistical framework to assess whether models perform better than expected by chance is lacking. A typical solution used is to compute empirical *p*-values by applying the same data analysis to a large number of datasets that follow the null hypothesis of no relation between the dependent and independent variables, and comparing the parameter values and performance measures of the model to their empirical null distributions (Ojala and Garriga, 2010; Steinfath et al., 2010; Riedelsheimer et al., 2012). 500 datasets following the null hypothesis of no relation between gene expression and phenotypes were generated by randomly permuting the phenotypes among the 60 plants. The following formula (Ojala and Garriga, 2010) was used to calculate *p*-values for the original oob *R*^2^ scores:

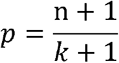

Where *n* is the number of times that a permuted model gave an equal or better *R*^2^ score than the ‘true’ model. Following (Ojala and Garriga, 2010), the standard deviation on the empirical *p*-value can be calculated as 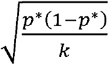, where *k* is the number of permutations and *p\** is the true *p*-value. This underlying true *p*-value is unknown, but at the critical *p*\* = 0.05, the calculated standard deviation on the empirical *p*-value when using 500 permutations is 0.0097, which is sufficiently low for our purposes.

### Accession Numbers

RNA-seq data have been deposited in the ArrayExpress database at EMBL-EBI (www.ebi.ac.uk/ arrayexpress) under accession number E-MTAB-8944. Sequence data from this article can be found in the Maize Genetics and Genomics Database (MaizeGDB) or GenBank/ EMBL databases under the following accession numbers: MADS1 (GRMZM2G171365), hb126 (GRMZM2G034113), WRKY53 (GRMZM2G012724), WRKY92 (GRMZM2G449681), AP2-EREBP (GRMZM2G042756), WRKY40 (GRMZM2G120320), XLG3b (GRMZM2G429113), MPK3-1 (GRMZM2G053987), TPS13.1 (GRMZM2G416836), CSP41A (GRMZM2G111216), CRB (GRMZM2G165655), SIG5 (GRMZM2G543629), prh2 (GRMZM2G140288), hb26 (GRMZM2G010929), PHR2 (GRMZM2G158662), ZMM15 (GRMZM2G553379), ZAP1 (GRMZM2G148693), unknown (GRMZM2G023625), unknown (GRMZM2G377311), MPK14 (GRMZM2G062914), unknown (GRMZM2G430780)

## Supporting information

Supplemental Data Set 1

Supplemental Data Set 2

Supplemental Data Set 3

Supplemental Data Set 4

Supplemental Data Set 5

Supplemental Data Set 6

Supplemental Data Set 7

Supplemental Data Set 8

Supplemental Data Set 9

Supplemental Data Set 10

Supplemental Data Set 11

Supplemental Data Set 12

Supplemental Data Set 13

Supplemental Data Set 14

Supplemental Data Set 15

Supplemental Data Set 16

## Supplemental Data

**Supplemental Data Set 1.** Transcriptome, metabolome, field position and phenotype data for the individual plants profiled in this study.

**Supplemental Data Set 2.** Spatially autocorrelated transcripts, metabolites and phenotypes in the single-plant dataset.

**Supplemental Data Set 3.** Spatially autocorrelated gene clusters in the single-plant dataset.

**Supplemental Data Set 4.** Significant correlations between the average expression profiles of spatially autocorrelated gene clusters and phenotypes.

**Supplemental Data Set 5.** Gene expression statistics for single-plant dataset.

**Supplemental Data Set 6.** Functional enrichment analysis of variably and stably expressed genes.

**Supplemental Data Set 7.** GO enrichment analysis of clusters and biclusters obtained from the single-plant transcriptome data.

**Supplemental Data Set 8.** List of Sequence Read Archive (SRA) samples used to calculate sampled networks.

**Supplemental Data Set 9.** List of target GO terms used for category-specific gene function predictions.

**Supplemental Data Set 10.** Gene function prediction performance plots for the GO categories listed in Supplemental Data Set 9.

**Supplemental Data Set 11.** Novel gene function predictions based on the single-plant co-expression network.

**Supplemental Data Set 12.** Transcripts significantly correlated with plant phenotypes.

**Supplemental Data Set 13.** Elastic net and random forest feature importance scores for blade length predictive models.

**Supplemental Data Set 14.** Elastic net and random forest feature importance scores for blade width predictive models.

**Supplemental Data Set 15.** Elastic net and random forest feature importance scores for husk leaf length predictive models.

**Supplemental Data Set 16.** List of regulatory genes annotated to the GO categories GO:0003700, GO:0006355 or GO:0004871.

## ACKNOWLEDGEMENTS

The authors thank Alex de Vliegher and his team from the Flanders Research Institute for Agricultural, Fisheries and Food (ILVO) for field trial management. Metabolomics data generation, funding for the RNA-seq experiments and funding for DH and TVH were provided by Syngenta Crop Protection, LLC. SDM is a fellow of the Research Foundation – Flanders (FWO, grant 1146319N).

## AUTHOR CONTRIBUTIONS

SM designed the study. SM and DI supervised the study. TVH, JDB, HN, DH and SM performed the field trial and generated data. DFC, SDM, JA, HS, DH and SM analyzed data. SM, DFC and SDM wrote the paper with input from the other authors.

## Notes

https://www.ebi.ac.uk/arrayexpress/experiments/E-MTAB-8944

