## Supplemental Figures and Tables for "Using single-plant -omics in the field to link maize genes to functions and phenotypes"

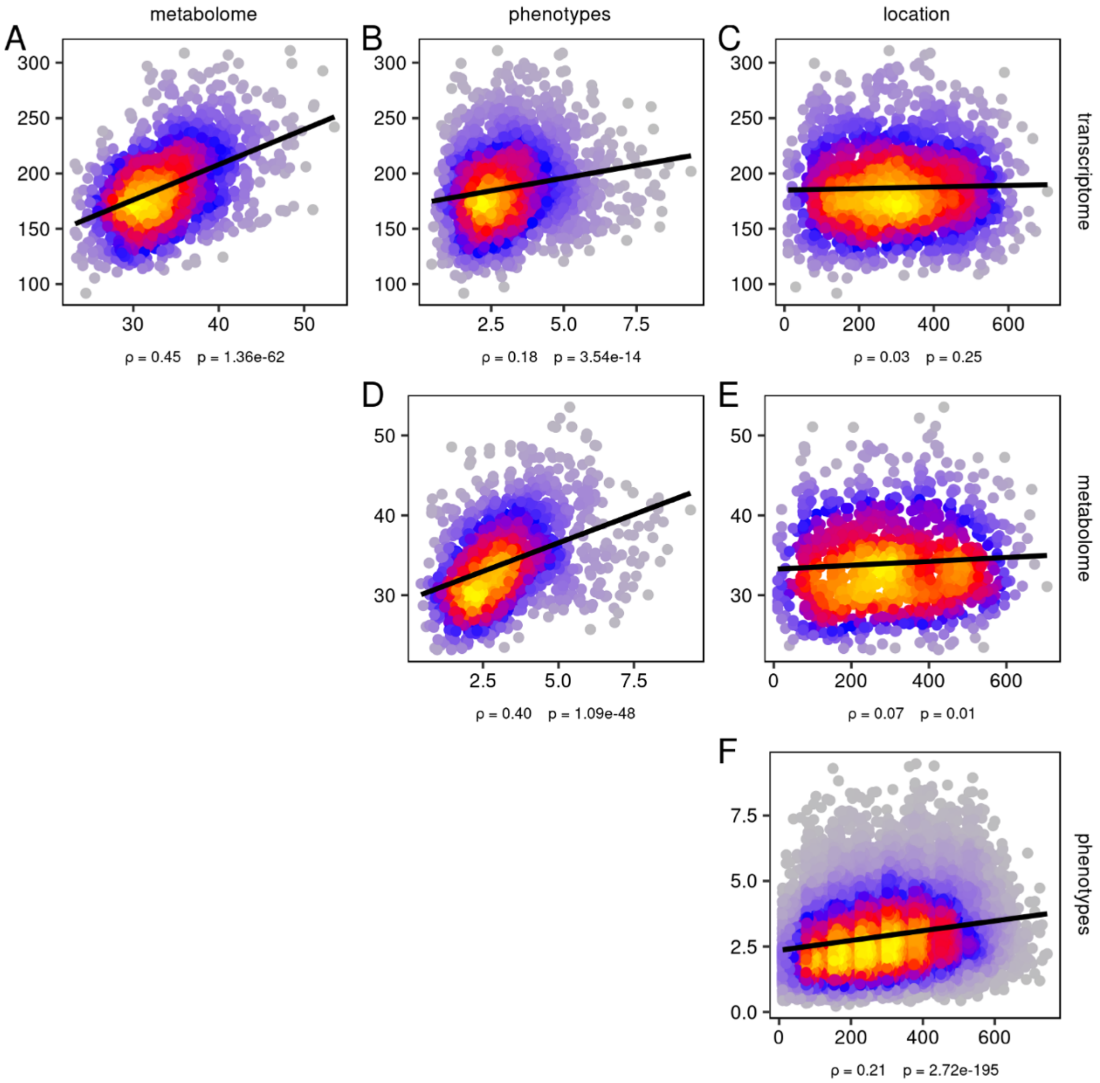


**Supplemental Figure 1. Correlations between pairwise distance profiles of different data types.** For each pair of plants, the Euclidean distance was calculated between their gene expression profiles, metabolite profiles, phenotype profiles and their locations in the field. Each point represents two distances for two different data types (on the *x* and *y* axis) for the same pair of plants. The color gradient visualizes the density of overlapping points in the graph. Linear regression lines are added in black, the corresponding regression coefficients and their significance are indicated below the plots. For example in **(A)**, pairs of plants that have more similar gene expression levels also tend to have more similar metabolite levels. The vertical bands in **(F)** arise from having only 8 widely spaced rows in the field as opposed to 56 narrowly spaced columns (Figure 1A). To prevent highly expressed genes from dominating the Euclidean distance calculation, all transcript profiles were z-scaled prior to calculating distances. Metabolite levels and phenotypes have similarly been z-scaled. Field distances (*x*-axis on subplots **C**, **E**, **F**) are given in centimeters.

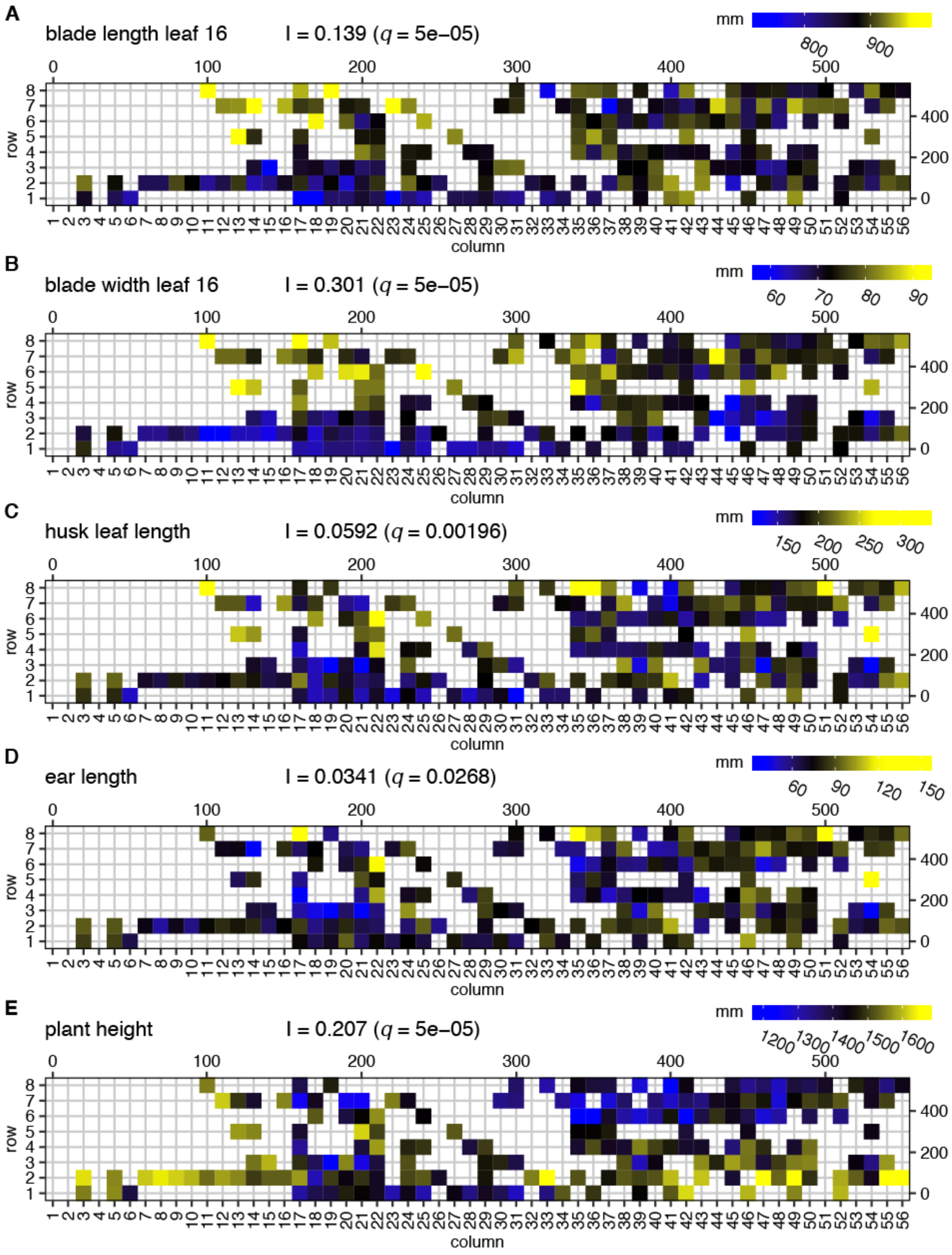


**Supplemental Figure 2. Spatial autocorrelation of phenotypes.** Each panel displays a phenotype mapped to the field. Moran's I values for spatial autocorrelation and the corresponding *q*-values after multiple testing correction (BH) are shown on top of the panels. The scales on the top and to the right of the field maps give field plot dimensions in cm.

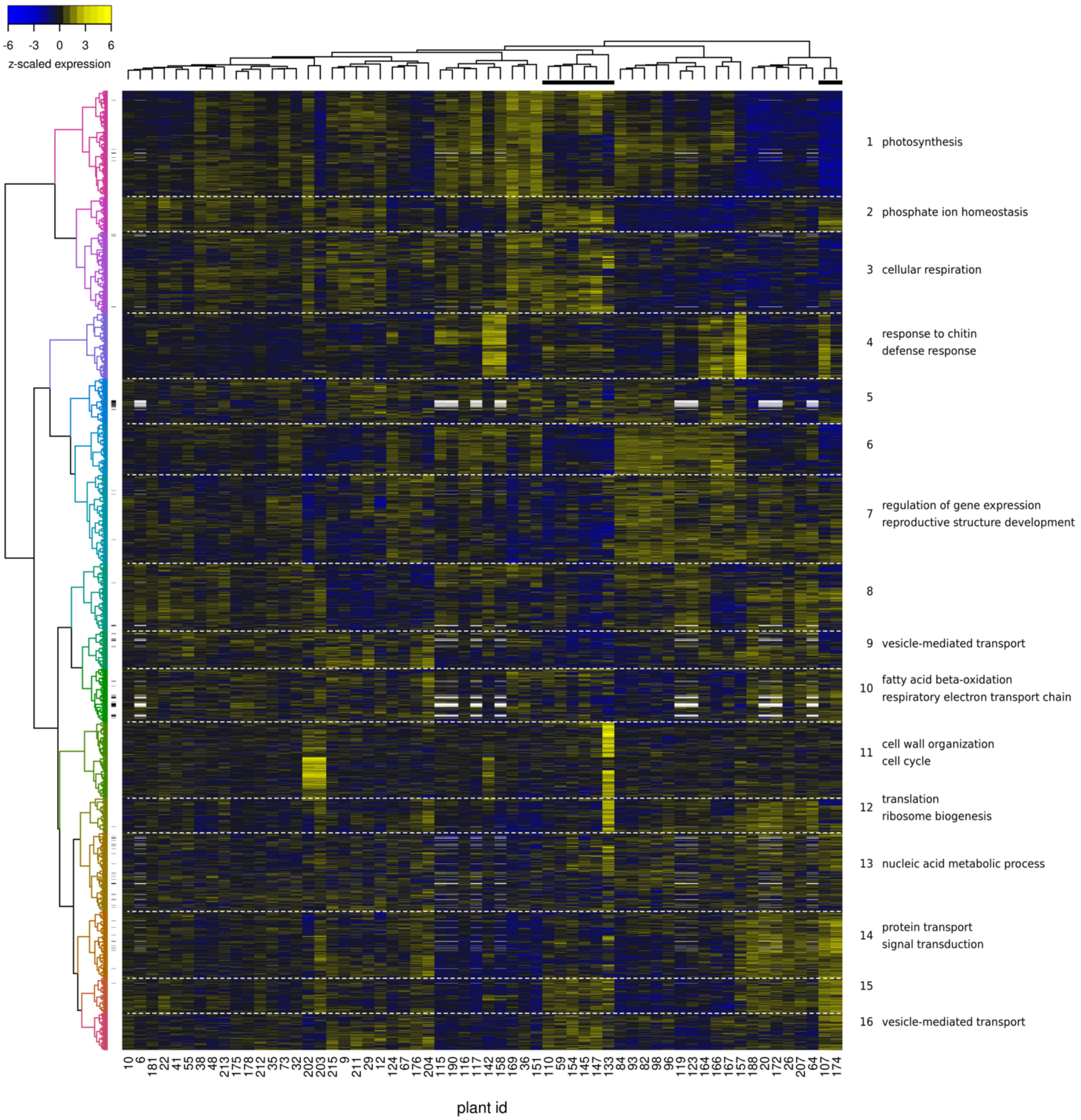


**Supplemental Figure 3. Hierarchical clustering of the transcriptome and metabolome datasets.** Rows are transcript/metabolite profiles and columns are plant expression profiles. Transcript/metabolite profiles were *z*-scaled to make them comparable. Metabolites are indicated with small dashes at the right of the dendrogram on the left. Gene/metabolite clusters are separated by horizontal white dashed lines. Representative significant GO enrichments for each cluster ($q < 0.01$) are indicated on the right. White rectangles are missing data (the metabolome was only profiled for 50 out of 60 plants). The black bars under the plant dendrogram indicate plants harvested on the second harvest date.

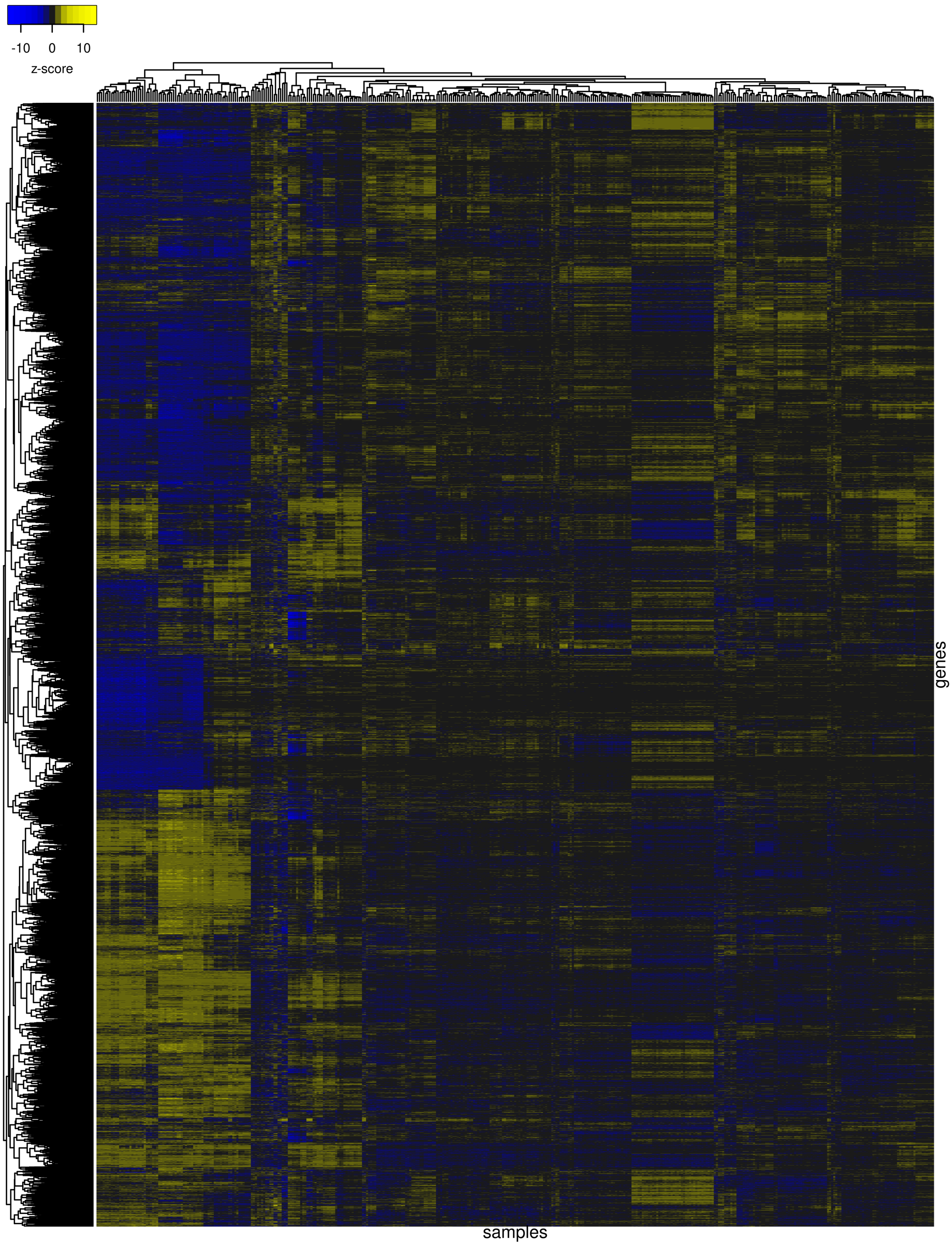


**Supplemental Figure 4. Hierarchical clustering of the maize B73 leaf transcriptome datasets obtained from the SRA database.** Rows are gene expression profiles and columns are SRA sample expression profiles. Transcript profiles were *z*-scaled to make them comparable.

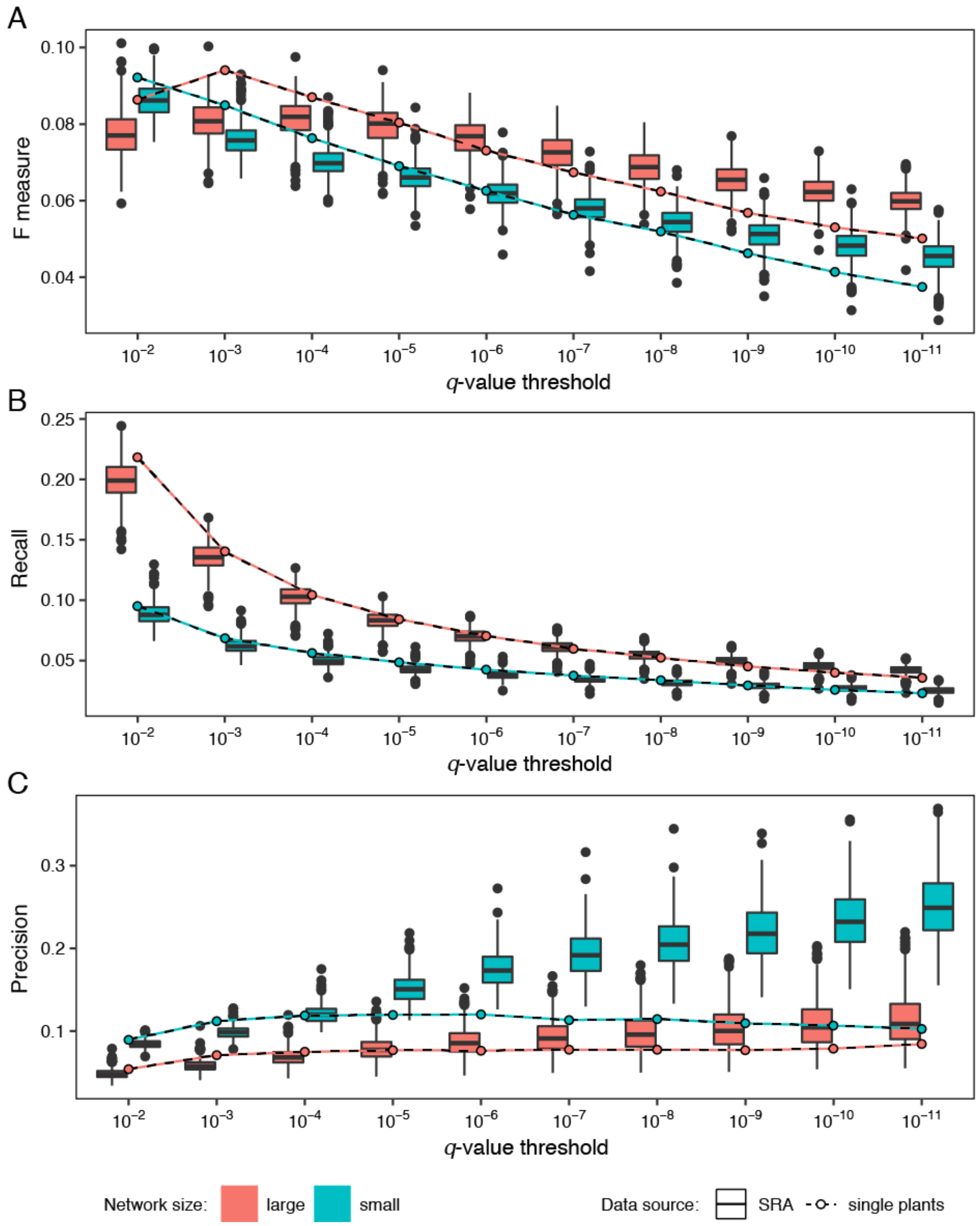


**Supplemental Figure 5. Gene function prediction performance of small versus large coexpression networks.** Gene function prediction was done on networks learned from the single-plant data and from sampled SRA datasets (see Methods). For the small network performance stats (depicted in blue), both the single-plant network and the 500 sampled networks were thresholded to contain 771,610 edges (the same number as in the single-plant network at a Bonferroni-corrected *p*-value cutoff of 0.01 for edge significance). For the large network performance stats, the sampled networks were generated using a Bonferroni corrected *p*-value cutoff of 0.01 for edge significance, giving rise to networks with 1-10 million edges. The large single plant network was thresholded to contain 4,680,819 edges, the average amount of edges obtained for the large sampled networks. Panels **(A)** to **(C)** depict F-measure, recall and precision curves for the resulting networks at various FDR thresholds, as in Figure 5. Except at FDR = 0.01 (-log(*q*) = 2), larger networks generally have higher gene function prediction performance, as measured by the F-measure.

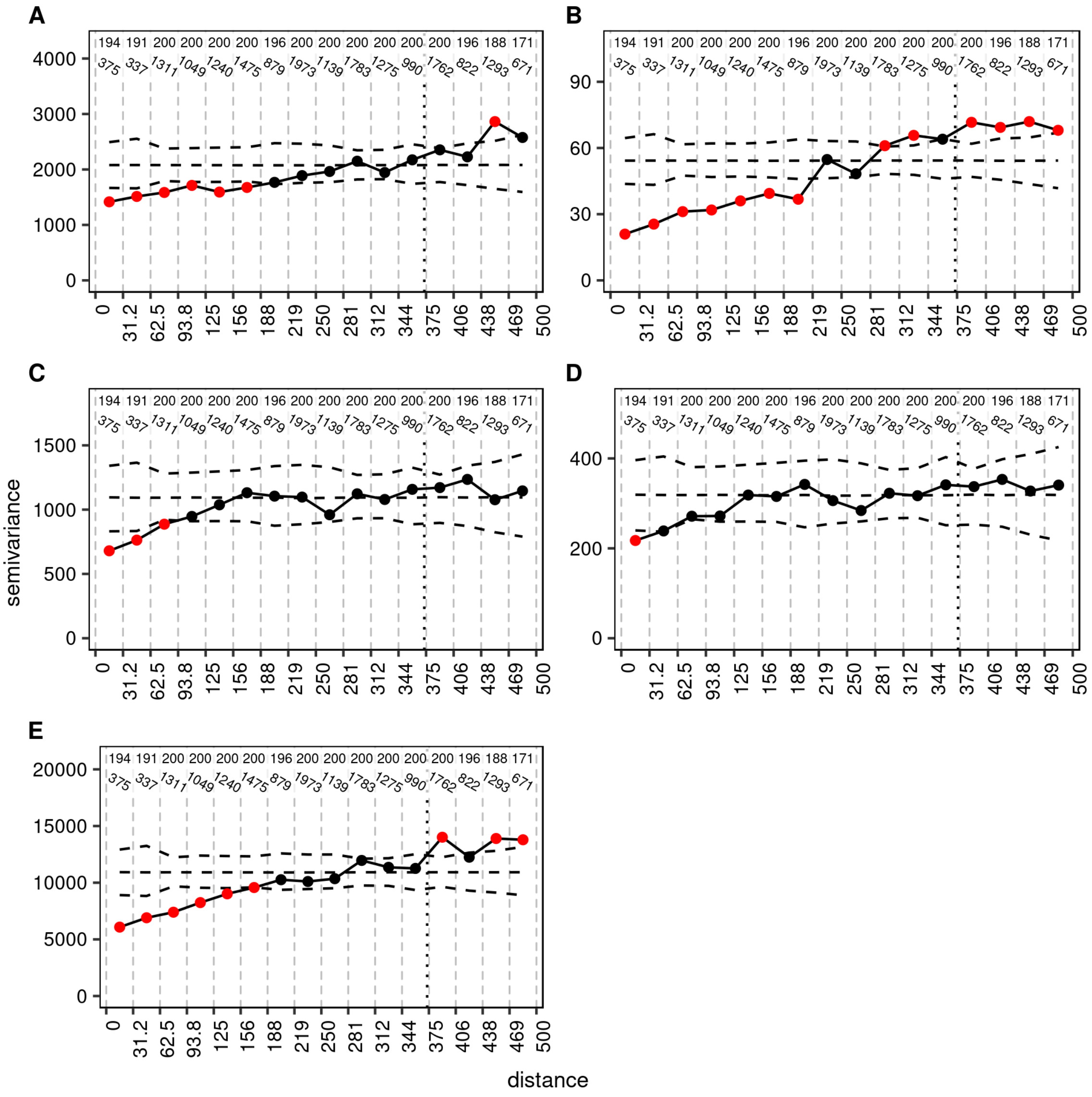


**Supplemental Figure 6. Range estimates for each phenotype.** Semivariance for each continuous phenotype in the study : **(A)** leaf 16 blade length, **(B)** leaf 16 blade width, **(C)** husk leaf length, **(D)** ear length, and **(E)** plant height. Semivariance (*y*-axis) is plotted with a solid black line for each distance bin (*x*-axis). The number of samples involved in the semivariance calculation is shown in the top row of numbers in each panel, the number of pairs in each bin is shown just below (slanted numbers). The semivariance was recalculated 10,000 times after permuting the locations of each plant to get an empirical null distribution. Dashed lines show the mean and the $0.15625^{th}$ (= 2.5/16) and $99.84375^{th}$ (= 1 - 2.5/16) percentiles of the empirical semivariance null distribution in each of the 16 bins. Red dots indicate distance bins in which the observed semivariance is significantly different from null expectations (Bonferroni-corrected $p < 0.05$). The vertical black dotted line in each panel indicates half the maximum distance in the field, after which the semivariance becomes hard to interpret.

**Supplemental Table 1. Genomic characteristics of highly variable genes and lowly variable genes in the single plant transcriptome dataset.** The highly variable gene (HVG) and lowly variable gene (LVG) sets are defined as the sets of genes with the 10% highest and lowest transcript CV, respectively, after removing the 5% most lowly expressed genes. The Q1 and Q3 columns give the first and third quartiles, respectively, of the genomic attribute distributions. The *q*-value column contains BH-corrected *p*-values of one-tailed Mann–Whitney U (MWU) tests to assess whether genomic attribute medians are significantly different between HVG and LVG. sd = standard deviation.

| | **min** | **Q1** | **median** | **Q3** | **max** | **mean** | **sd** | ***q*** |
|---|---|---|---|---|---|---|---|---|
| **Gene length (bp)** | | | | | | | | |
| HVG | 269 | 1,418.5 | 2,201.5 | 3,812 | 153,142 | 3,221.3 | 5,923.1 | 3.11E-289 |
| LVG | 1,062 | 4,104.3 | 5,844 | 9,044.5 | 136,766 | 7,878.7 | 7,556.3 | |
| **CDS length (bp)** | | | | | | | | |
| HVG | 74 | 614.3 | 1,036 | 1,531.8 | 7,194 | 1,188.2 | 788.7 | 1.66E-27 |
| LVG | 77 | 819.3 | 1,305.5 | 1,929.5 | 10,913 | 1,527.9 | 1,055.3 | |
| **# Introns** | | | | | | | | |
| HVG | 0 | 0 | 2 | 6 | 121 | 5.4 | 10.2 | 6.66E-245 |
| LVG | 0 | 6 | 13 | 21.8 | 307 | 16.9 | 17.5 | |
| **# Exons** | | | | | | | | |
| HVG | 1 | 1 | 3 | 8 | 126 | 7.1 | 11.0 | 1.11E-237 |
| LVG | 1 | 8 | 15 | 25 | 314 | 19.3 | 18.3 | |
| **# Correlated TFs** | | | | | | | | |
| HVG | 1 | 3 | 13 | 27 | 63 | 17.0 | 15.1 | 5.92E-59 |
| LVG | 1 | 1 | 2 | 3 | 13 | 2.6 | 2.4 | |

**Supplemental Table 2: Top-10 novel regulators predicted to be involved in the response to chitin based on the single plant data.** Predictions supported by literature evidence are highlighted in yellow.

| Rank | Gene ID | Predicted GO ID | Predicted GO Name | *q*-value | Gene Description |
|---|---|---|---|---|---|
| 1 | GRMZM2G301089 | 10200 | response to chitin | 1.0506E-17 | Transcription factor bHLH92 |
| 2 | GRMZM2G174347 | 10200 | response to chitin | 1.3968E-17 | ethylene-responsive transcription factor 4 |
| 3 | GRMZM2G012724-ZmWRKY53 | 10200 | response to chitin | 5.3193E-17 | Superfamily of TFs having WRKY and zinc finger domains |
| 4 | GRMZM2G174558 | 10200 | response to chitin | 5.3678E-17 | DNA-binding WRKY |
| 5 | GRMZM2G449681-FPKM53 | 10200 | response to chitin | 8.0247E-16 | WRKY53 - superfamily of TFs having WRKY and zinc finger domains |
| 6 | GRMZM2G107031 | 10200 | response to chitin | 2.8395E-15 | Transcription factor PCF2 |
| 7 | GRMZM2G379005 | 10200 | response to chitin | 3.2908E-15 | C2C2-Putative GATA transcription factor 13 |
| 8 | GRMZM2G042756 | 10200 | response to chitin | 5.2456E-15 | AP2-EREBP, dehydration-responsive element-binding protein 1D |
| 9 | GRMZM2G137341 | 10200 | response to chitin | 1.6835E-14 | dehydration-responsive element-binding protein 1A |
| 10 | GRMZM2G120320 | 10200 | response to chitin | 1.8174E-14 | WRKY transcription factor 40 |
| 11 | GRMZM2G153206 | 10200 | response to chitin | 2.084E-14 | Rapid alkalinization factor |

**Supplemental Table 3: Top-10 novel regulators predicted to be involved in the response to water deprivation based on the single plant data.** Predictions supported by literature evidence are highlighted in yellow.

| Rank | Gene ID | Predicted GO ID | Predicted GO Name | *q*-value | Gene Description |
|---|---|---|---|---|---|
| 1 | GRMZM2G429113 | 9269 | response to desiccation | 0.00266142 | Extra-large guanine nucleotide-binding protein 3 |
| 2 | GRMZM2G059428 | 9414 | response to water deprivation | 0.00283171 | NAC transcription factor NAM |
| 3 | GRMZM2G416836 | 9819 | drought recovery | 0.00291509 | alpha-trehalose-phosphate synthase [UDP-forming] 11 |
| 4 | GRMZM2G053987-ZmMPK3-1 | 9414 | response to water deprivation | 0.00316057 | activated protein kinase |
| 5 | GRMZM2G083759 | 42631 | cellular response to water deprivation | 0.00318067 | signal transducer (LOC100281544)POZ domain-containing protein NPY1 |
| 6 | GRMZM2G171569 | 42631 | cellular response to water deprivation | 0.00401808 | Ethylene-responsive transcription factor |
| 7 | GRMZM2G135300 | 9819 | drought recovery | 0.00402637 | TAF5-like RNA polymerase II p300/CBP-associated factor-associated |
| 8 | GRMZM2G000520 | 9414 | response to water deprivation | 0.00452256 | responsive transcription factor ERF025 |
| 9 | GRMZM2G120320-ZmWRKY40 | 9414 | response to water deprivation | 0.00513427 | WRKY transcription factor 40 |
| 10 | GRMZM2G176489 | 9414 | response to water deprivation | 0.00526752 | WRKY transcription factor 72 |

**Supplemental Table 4: Top-10 novel regulators predicted to be involved in $C_4$ photosynthesis based on the single plant data.** Predictions supported by literature evidence are highlighted in yellow.

| Rank | Gene ID | Predicted GO ID | Predicted GO Name | *q*-value | Gene Description |
|---|---|---|---|---|---|
| 1 | GRMZM2G111216-CSP41A | 9760 | $C_4$ photosynthesis | 2.71E-23 | |
| 2 | GRMZM2G165655-CRB | 9760 | $C_4$ photosynthesis | 2.01E-17 | Chloroplast stem-loop binding protein of 41 kDa b |
| 3 | GRMZM2G158662 | 9760 | $C_4$ photosynthesis | 1.57E-15 | Blue-light photoreceptor PHR2 |
| 4 | GRMZM2G543629-SIG5 | 9760 | $C_4$ photosynthesis | 2.48E-15 | sigma factor SigA, Sigma70-like |
| 5 | GRMZM2G438438 | 9760 | $C_4$ photosynthesis | 6.97E-14 | zf-HD, Transcription factor HB29 |
| 6 | GRMZM2G049672 | 9760 | $C_4$ photosynthesis | 2.83E-13 | protein ligase CHFR |
| 7 | GRMZM2G010929 | 9760 | $C_4$ photosynthesis | 1.29E-12 | lumenal 15 kDa protein 1, chloroplastic |
| 8 | GRMZM2G007063 | 9760 | $C_4$ photosynthesis | 2.69E-12 | bZIP:PlantTFDB~opaque2 heterodimerizing protein2 |
| 9 | GRMZM5G854901 | 9760 | $C_4$ photosynthesis | 7.84E-12 | cytidine(34)-2'-O)-methyltransferase |
| 10 | GRMZM2G140288 | 9760 | $C_4$ photosynthesis | 2.34E-11 | MYST-like histone acetyltransferase 1 |
