## Supplemental Data Set 4 for "Using single-plant -omics in the field to link maize genes to functions and phenotypes"

**Supplemental Data Set 4. Significant correlations between the average expression profiles of spatially autocorrelated gene clusters and phenotypes.**

Each figure contains two panels. The top panel displays the average z-scored gene expression profile of a spatially autocorrelated gene cluster mapped to the field. The bottom panel displays a phenotype that correlates significantly with the average cluster expression profile concerned ($q \leq 0.05$). Shown on top of each figure are the Pearson's correlation (r) between the cluster expression profile and the phenotype, the corresponding p-value (computed using `cor.test` in R) and the corresponding q-value (computed using the Benjamini-Hochberg method on all comparisons per phenotype). The scales on the top and to the right of the field maps give field plot dimensions in cm.

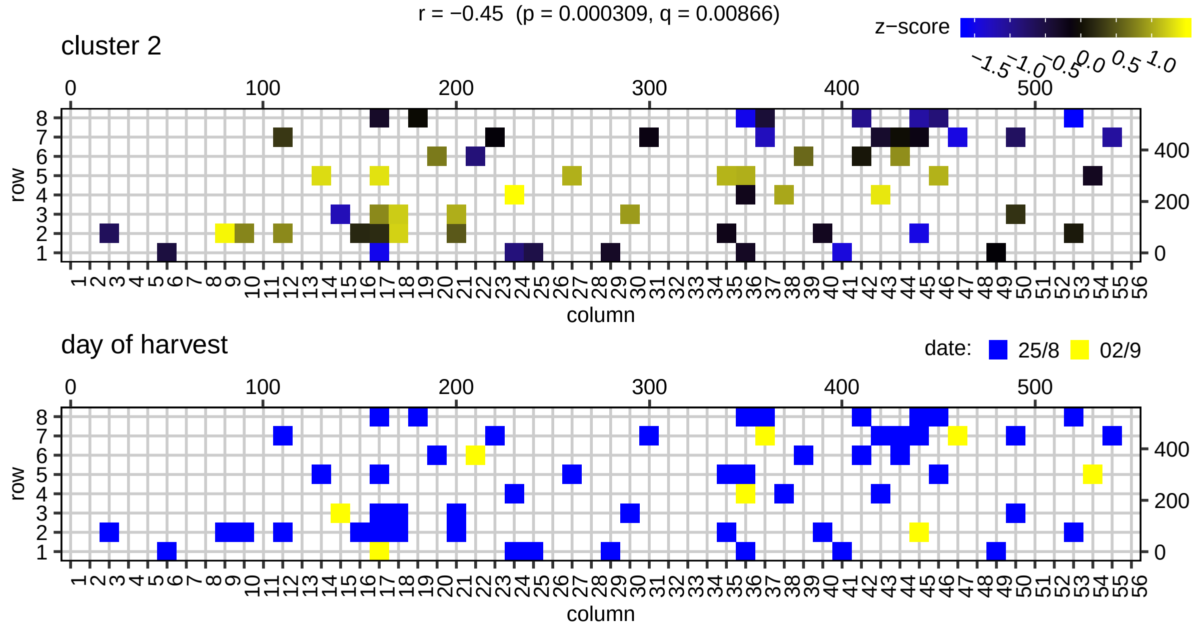


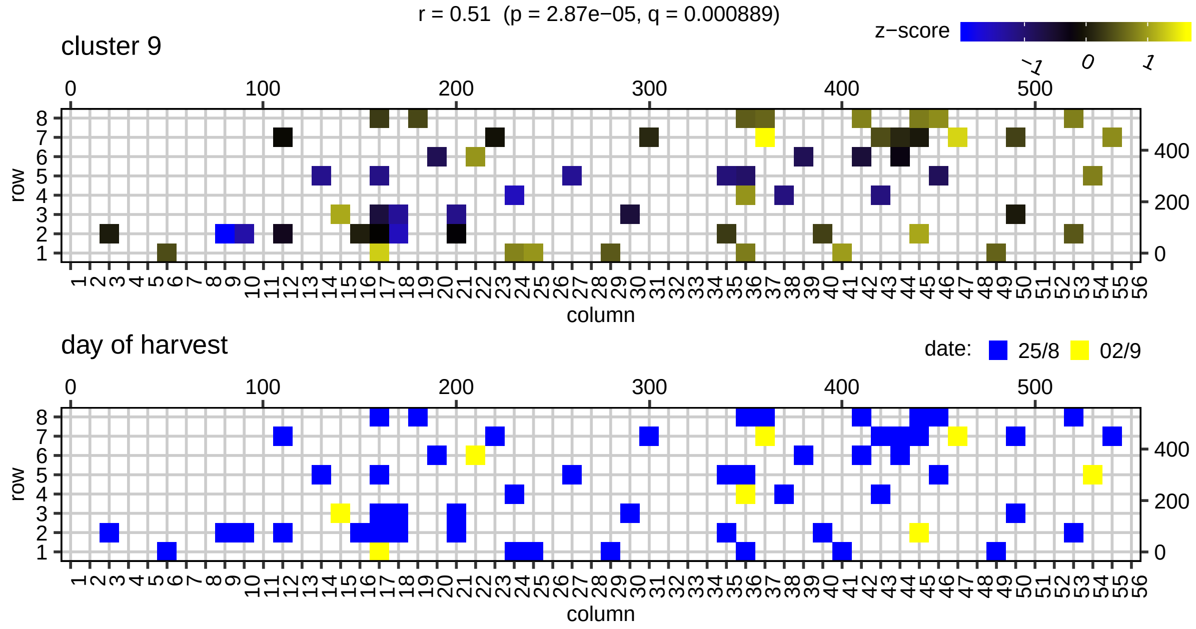

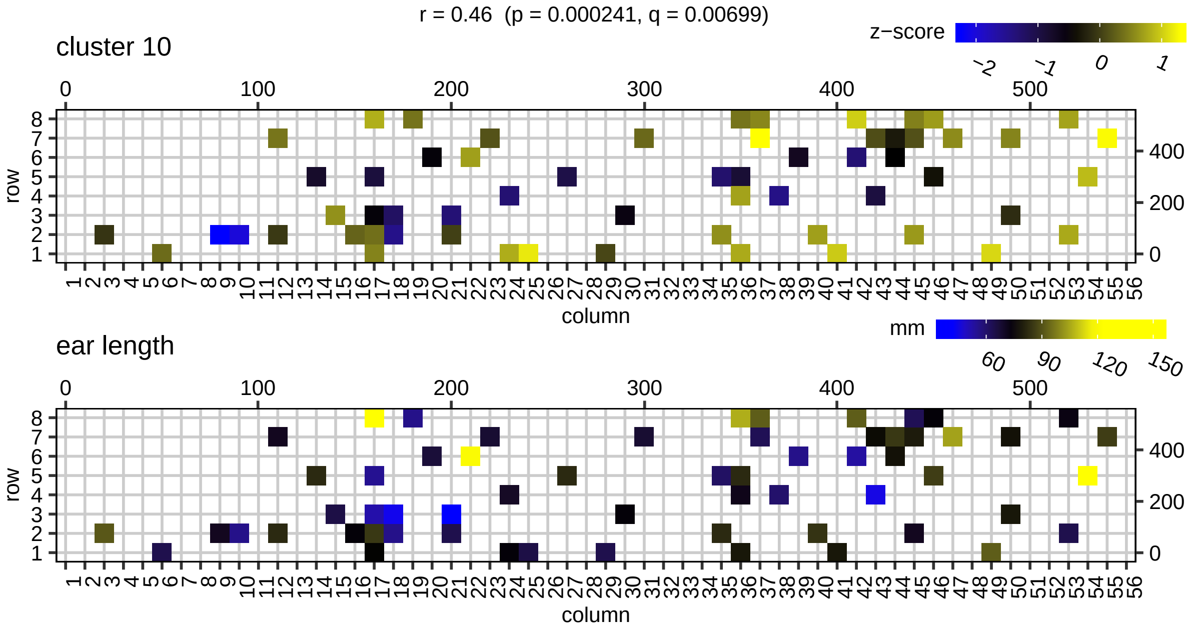
r = 0.46 (p = 0.000241, q = 0.00699)
cluster 10
z−score
−2 −1 0 1
row
column
ear length
mm
60 90 120 150
row
column

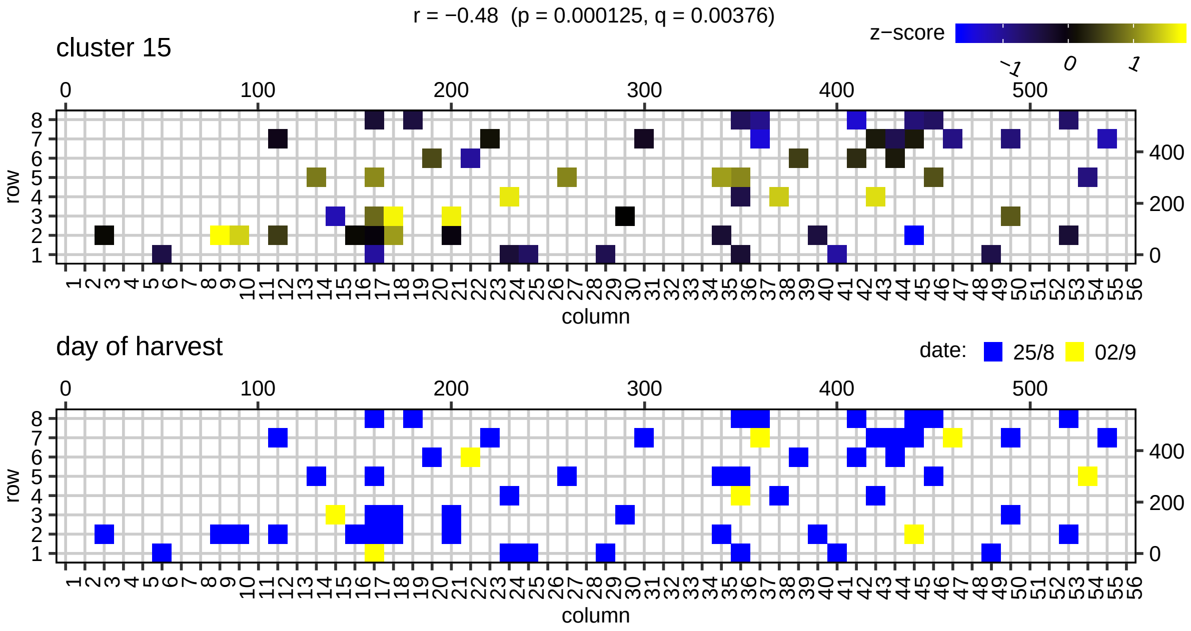
r = −0.48 (p = 0.000125, q = 0.00376)
cluster 15
z−score
−1 0 1
row
column
day of harvest
date: 25/8 02/9
row
column

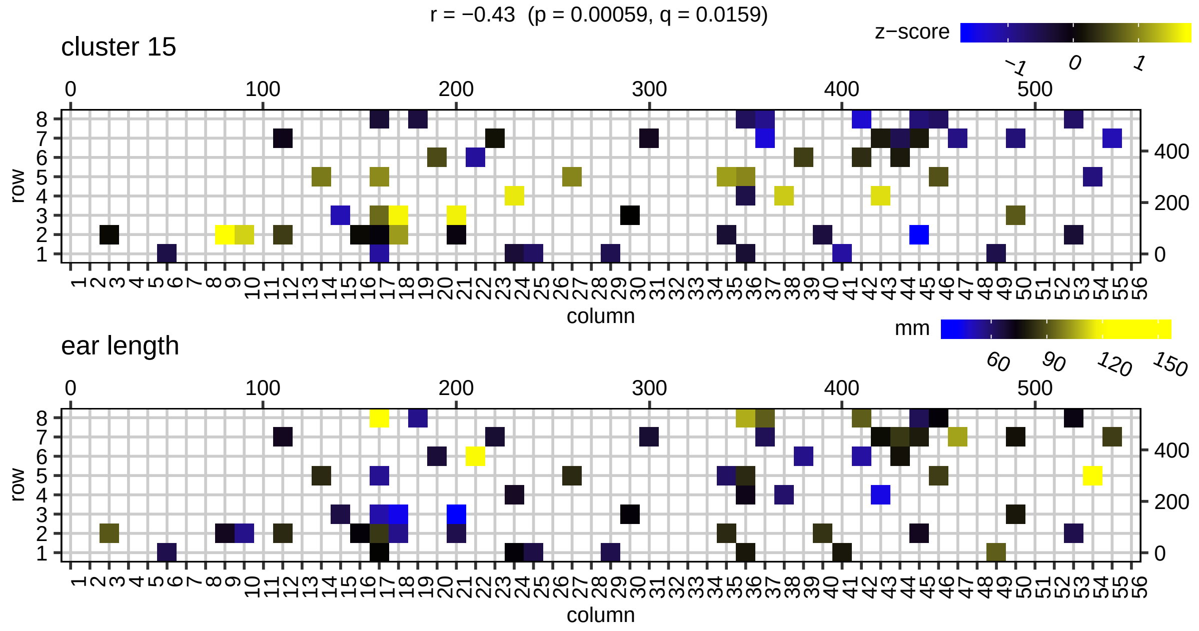
r = −0.43 (p = 0.00059, q = 0.0159)
cluster 15
z−score
−1 0 1
ear length
mm
60 90 120 150
row
column

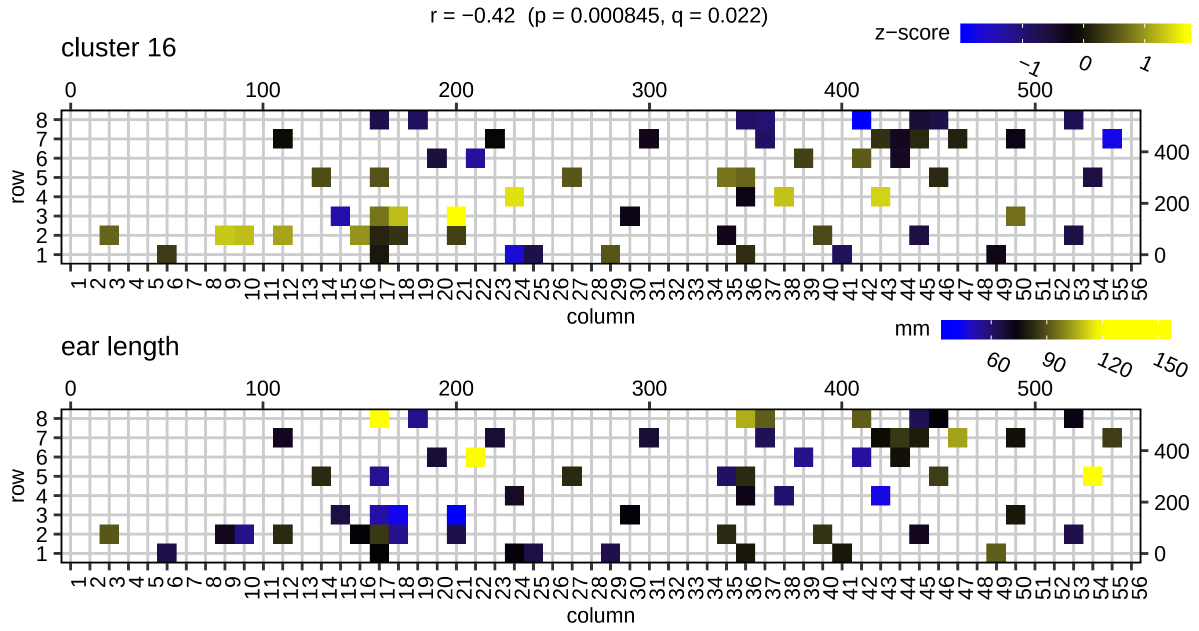
r = −0.42 (p = 0.000845, q = 0.022)
cluster 16
z−score
−1 0 1
ear length
mm
60 90 120 150
row
column

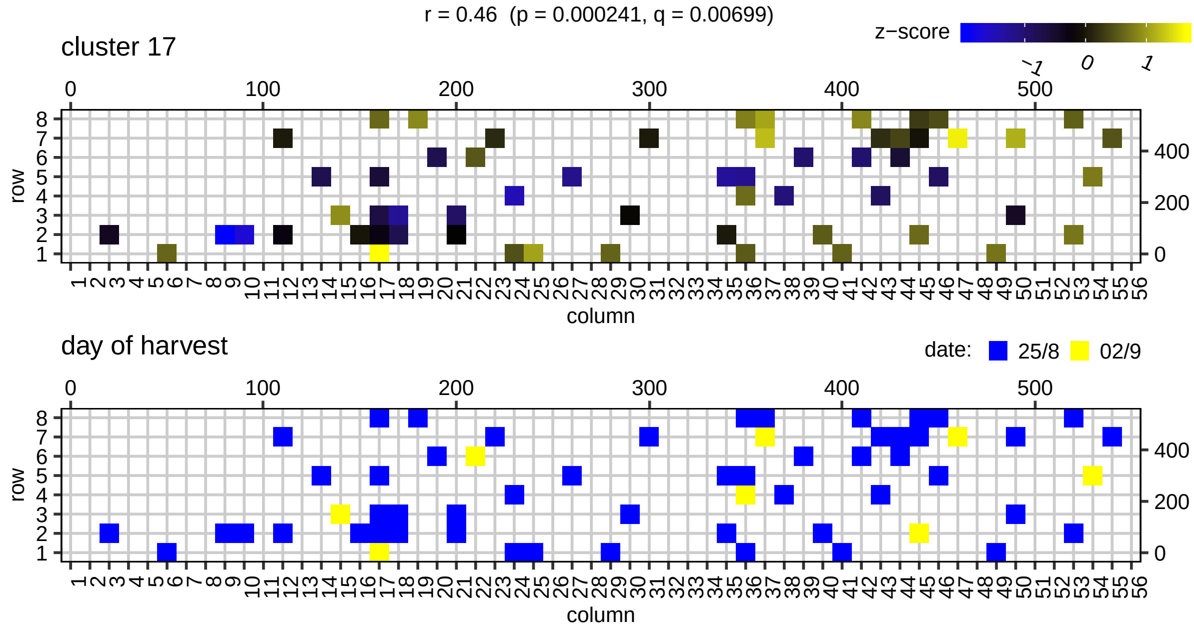
r = 0.46 (p = 0.000241, q = 0.00699)
cluster 17
z−score
-1 0 1
day of harvest
date: 25/8 02/9
row
column

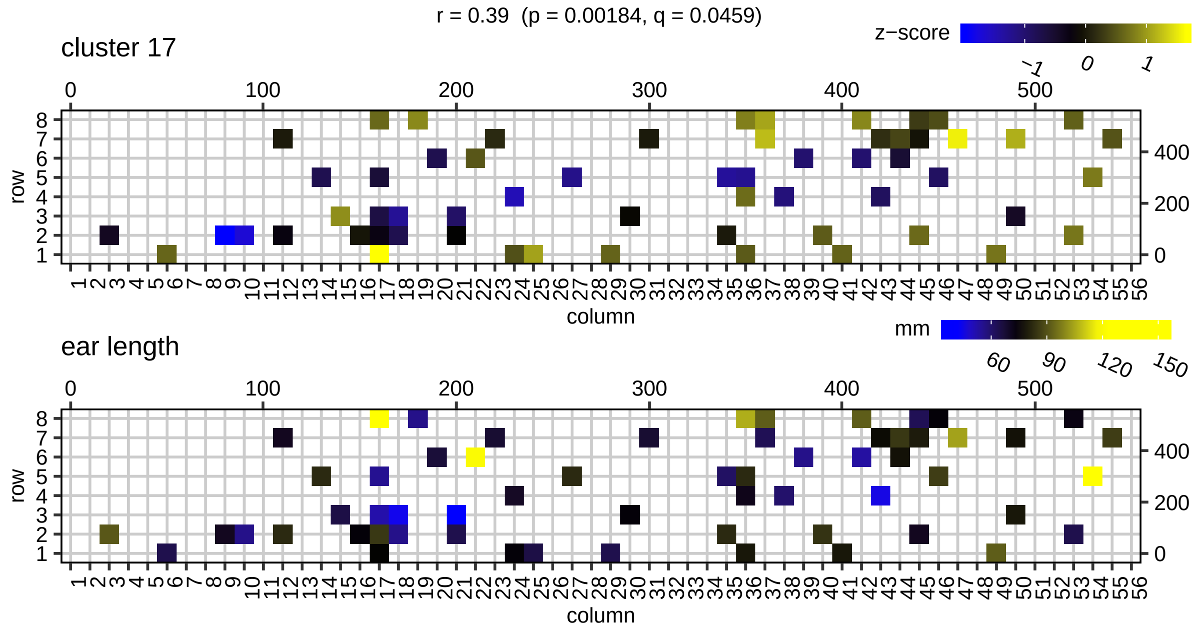
r = 0.39 (p = 0.00184, q = 0.0459)
cluster 17
z−score
-1 0 1
ear length
mm
60 90 120 150
row
column

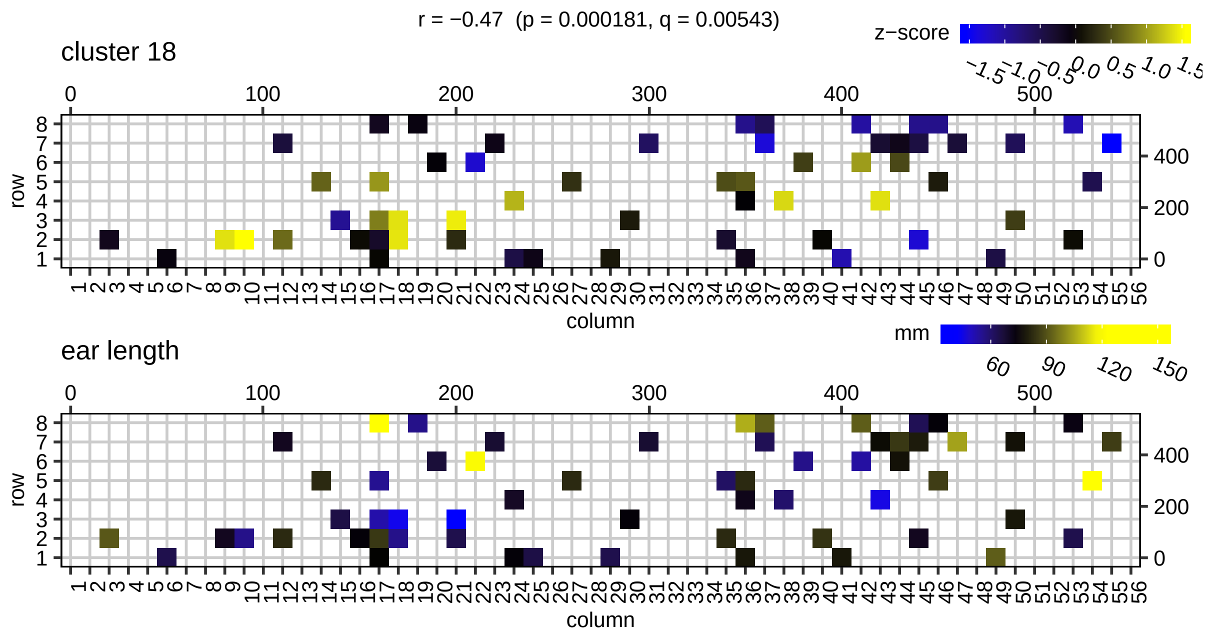
r = −0.47 (p = 0.000181, q = 0.00543)
cluster 18
z−score
ear length
mm
row
column

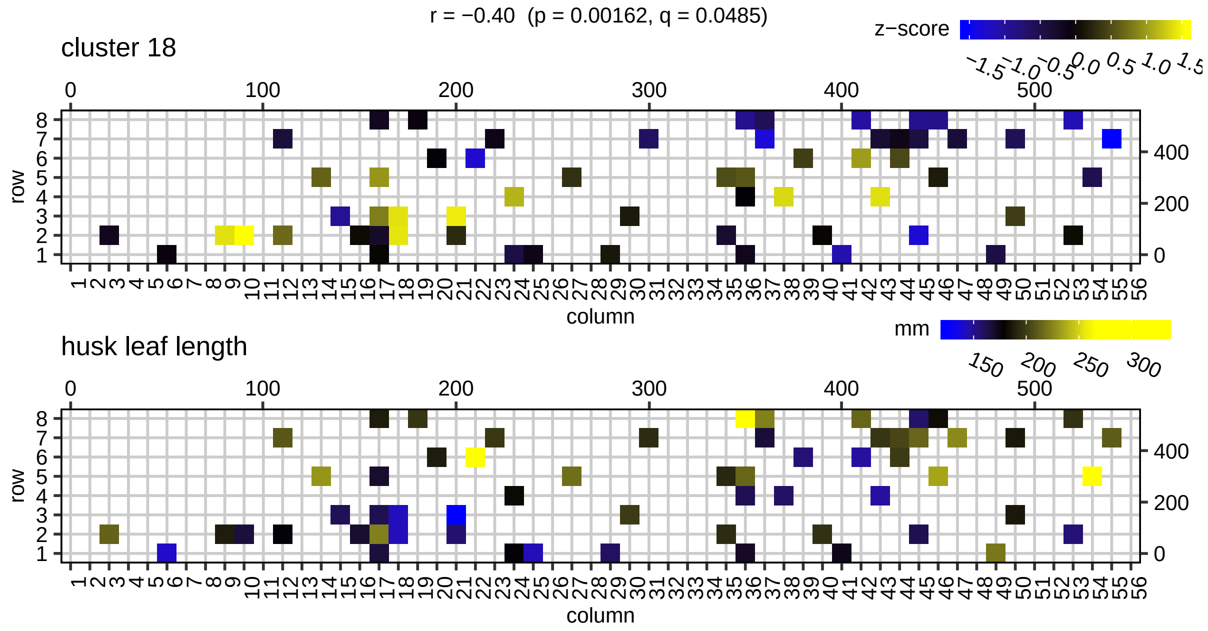
r = −0.40 (p = 0.00162, q = 0.0485)
cluster 18
z−score
husk leaf length
mm
row
column

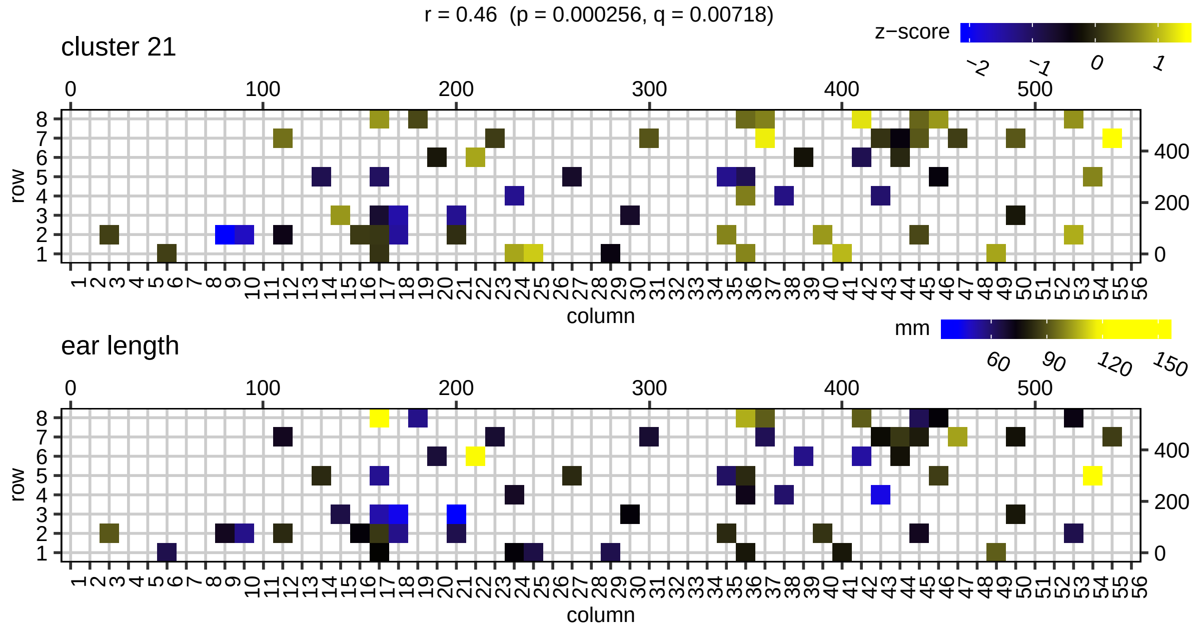
r = 0.46 (p = 0.000256, q = 0.00718)
cluster 21
z−score
ear length
mm
row
column

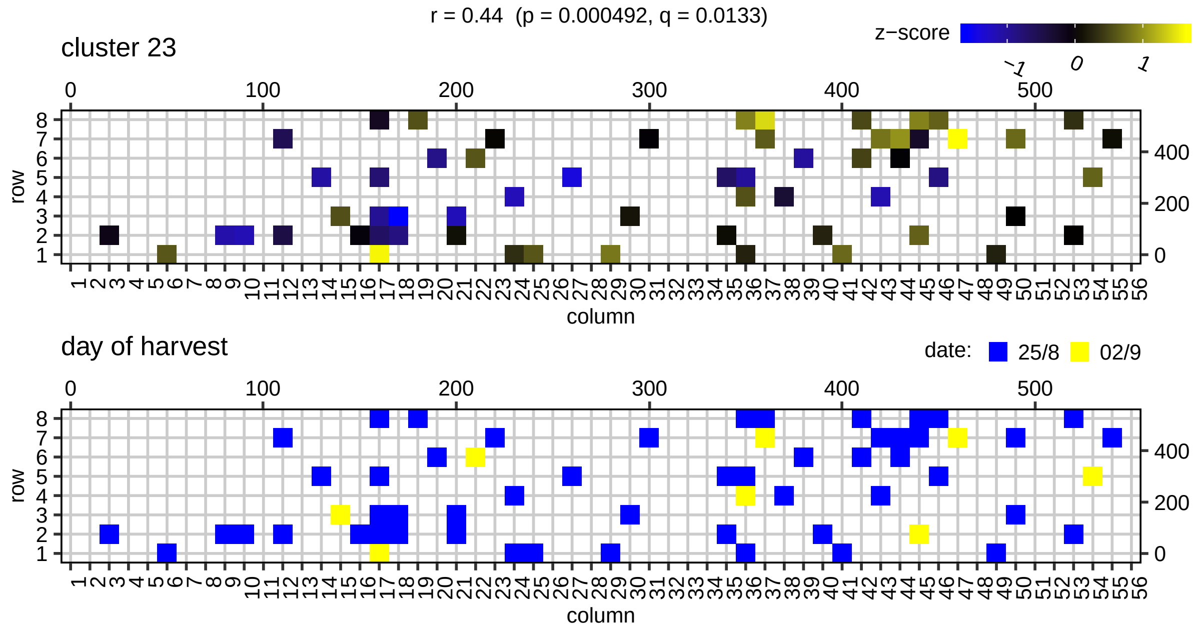
r = 0.44 (p = 0.000492, q = 0.0133)
cluster 23
z−score
day of harvest
date: 25/8 02/9
row
column

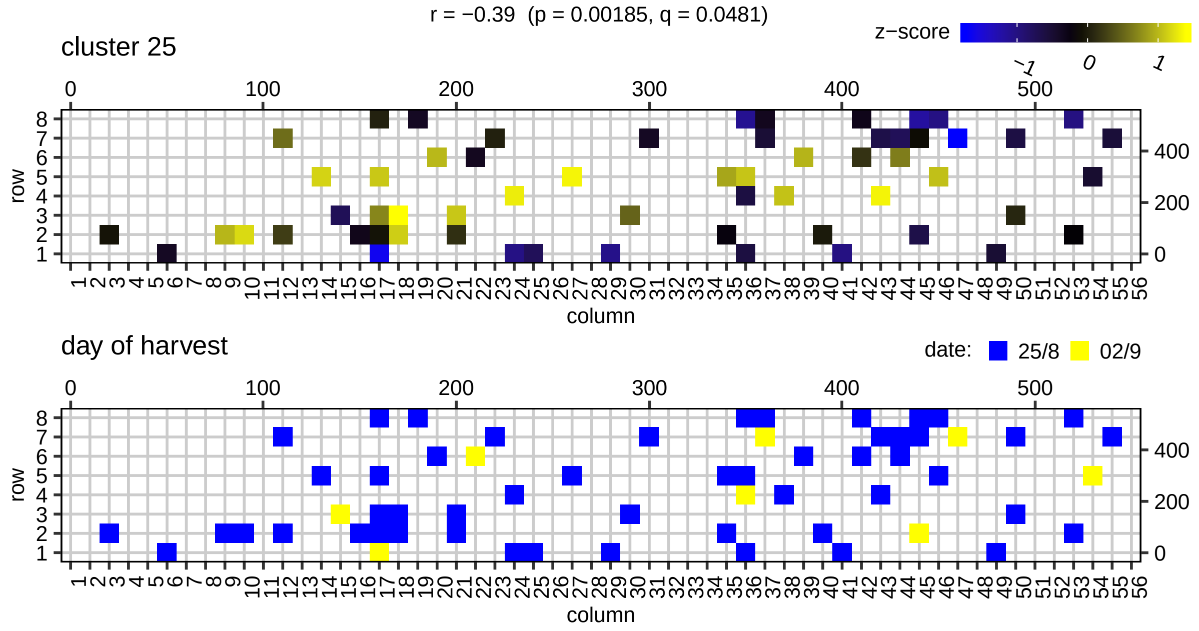
r = −0.39 (p = 0.00185, q = 0.0481)
cluster 25
z−score
−1
0
1
row
column
day of harvest
date: 25/8 02/9

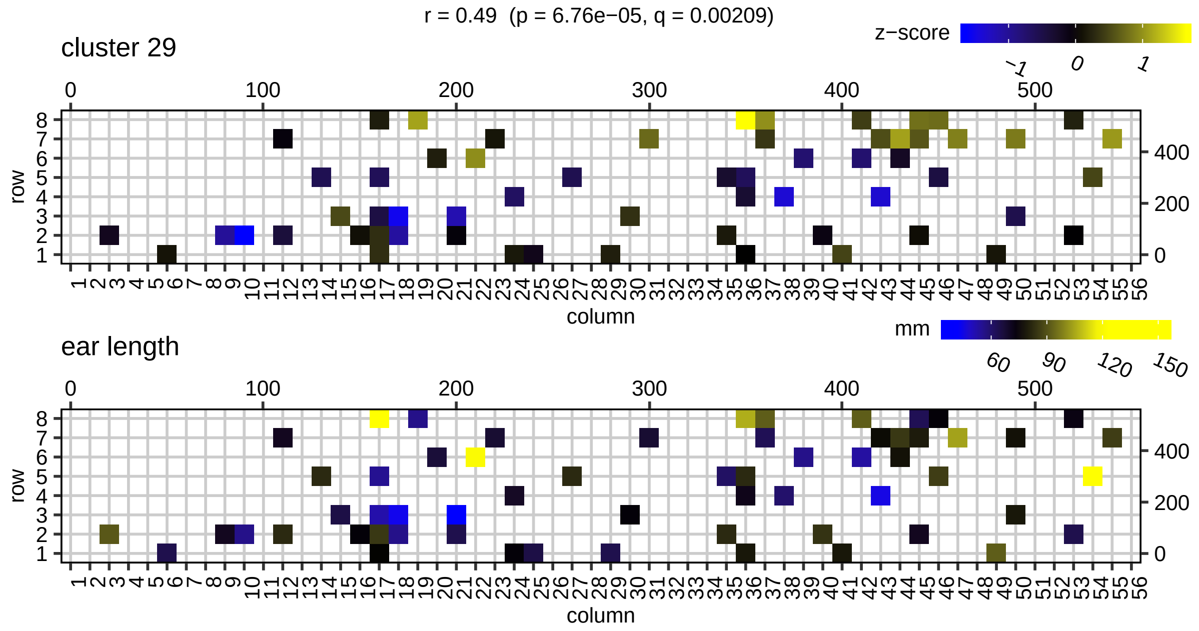
r = 0.49 (p = 6.76e−05, q = 0.00209)
cluster 29
z−score
−1
0
1
row
column
ear length
mm
60
90
120
150

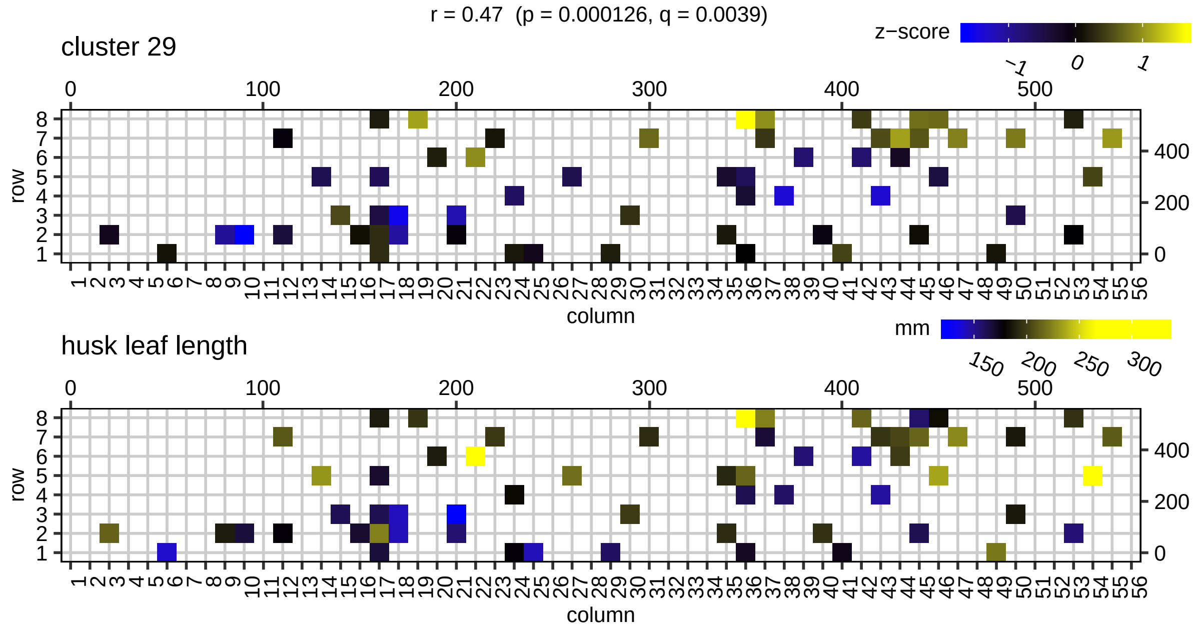
r = 0.47 (p = 0.000126, q = 0.0039)
cluster 29
z−score
−1
0
1
row
column
husk leaf length
mm
150
200
250
300
