## Supplemental Data Set 10 for "Using single-plant -omics in the field to link maize genes to functions and phenotypes"

**Supplemental Data Set 10. Gene function prediction performance plots for the GO categories listed in Supplemental Data Set 9.**

For each GO category, four panels are shown depicting the F-measure, recall, precision and the number of predicted positives (true positives + false positives) for the single plant network (red line) and the sampled networks (box-and-whisker plots) for FDR thresholds ranging from $10^{-2}$ to $10^{-11}$. Very general and uninformative GO categories, such as ‘biological process’, are not plotted. In addition, GO categories for which the single plant network and over half of the sampled networks produced no predictions at $q \leq 0.01$ are omitted (see Methods). When a network produced zero predictions at a certain FDR threshold, the F-measure, recall and precision were set to zero in order to make a fair comparison to other networks that do have predictions at this FDR threshold. Boxes extend from the 25th to the 75th percentile, with the median indicated by the central black line. Whiskers extend from each end of the box to the most extreme values within 1.5 times the interquartile range from the respective end. Data points beyond this range are displayed as little black circles. The label next to the GO name on top of each plot (Very Good, Good, Average, Poor, Very Poor) indicates how well the single plant network scored compared to the sampled networks (see Methods).

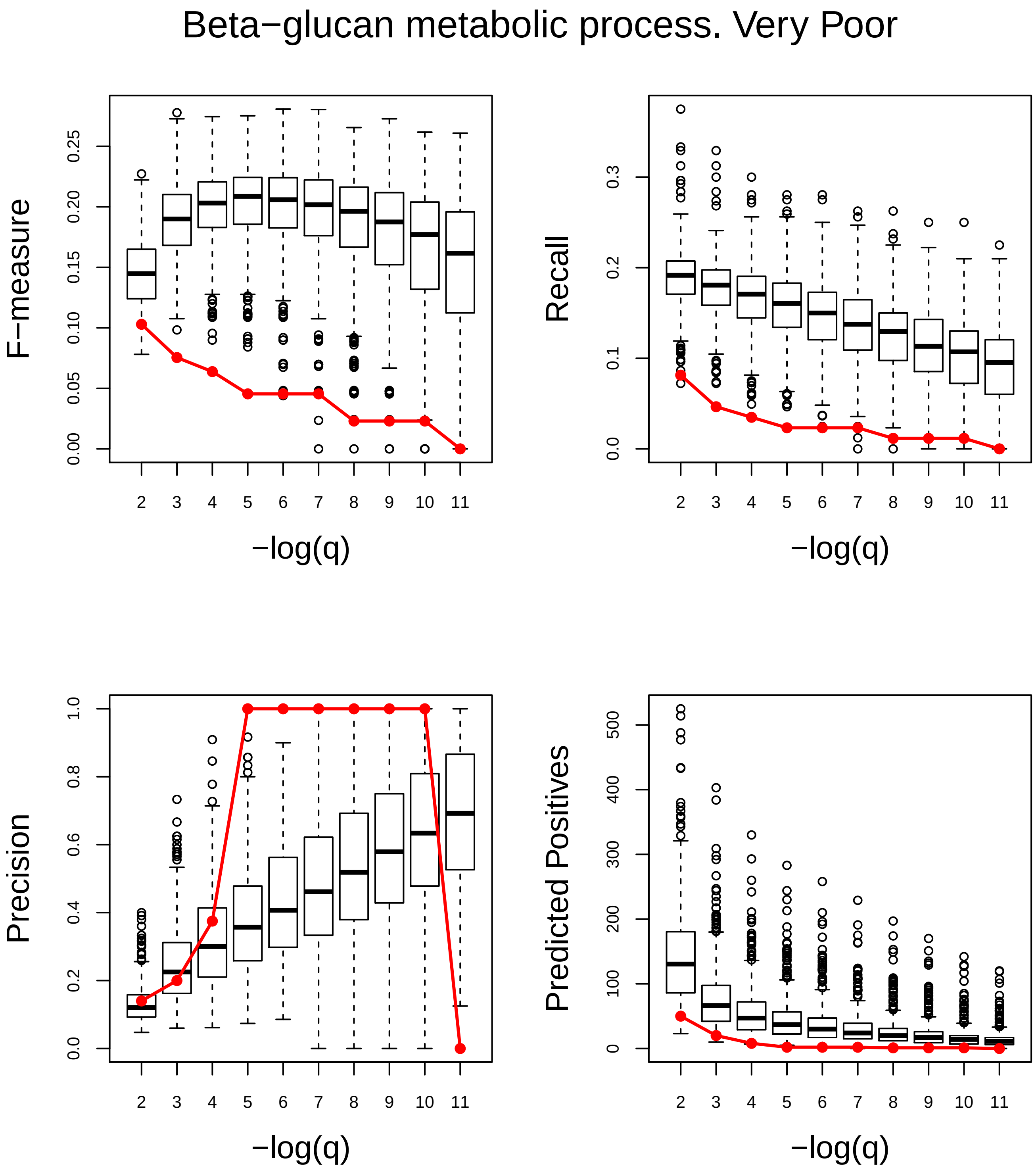
Beta−glucan metabolic process. Very Poor
F−measure
0.00
0.05
0.10
0.15
0.20
0.25
Recall
0.0
0.1
0.2
0.3
Precision
0.0
0.2
0.4
0.6
0.8
1.0
Predicted Positives
0
100
200
300
400
500
2 3 4 5 6 7 8 9 10 11
−log(q)

### Biosynthetic process. Very Good

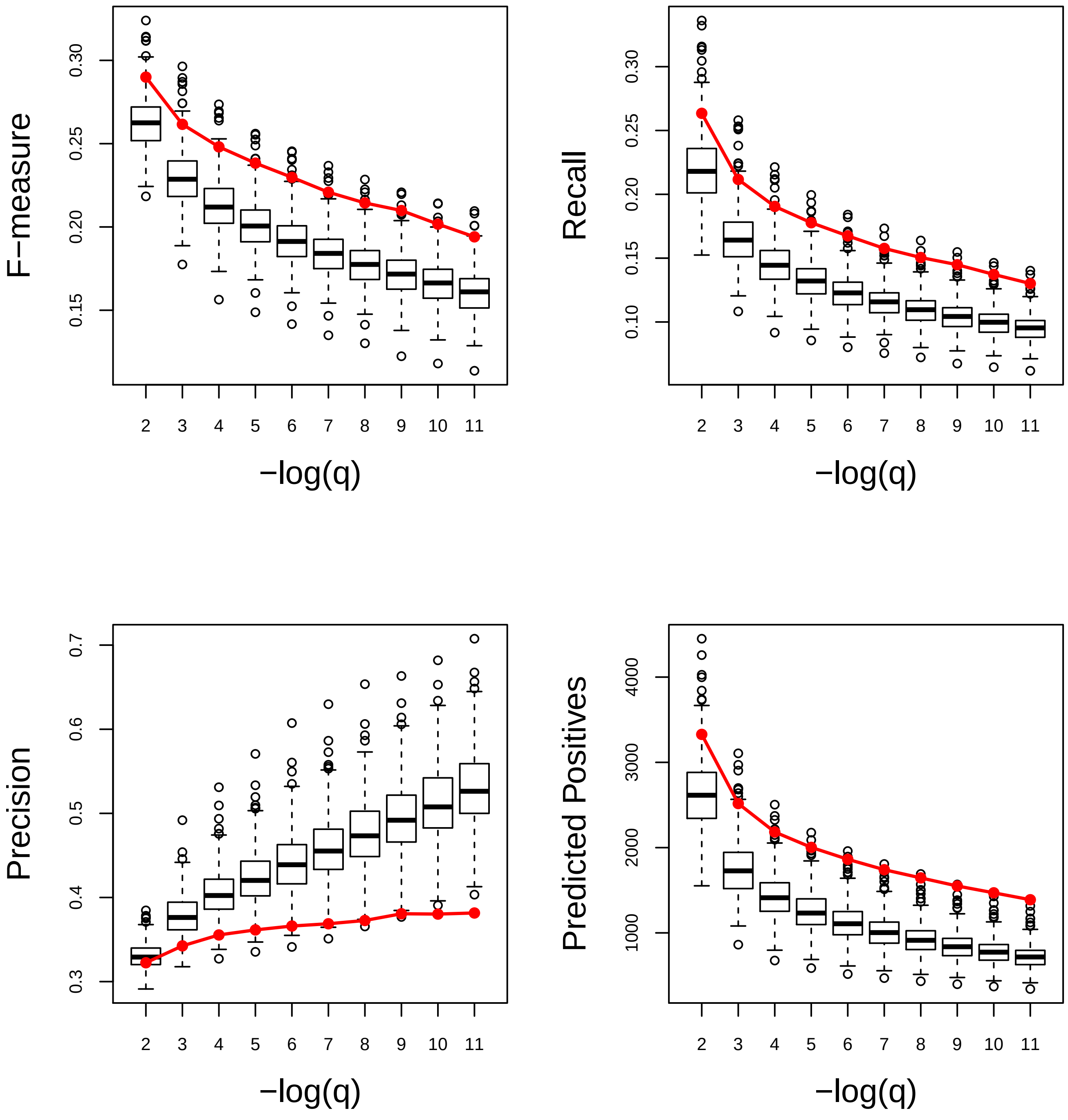

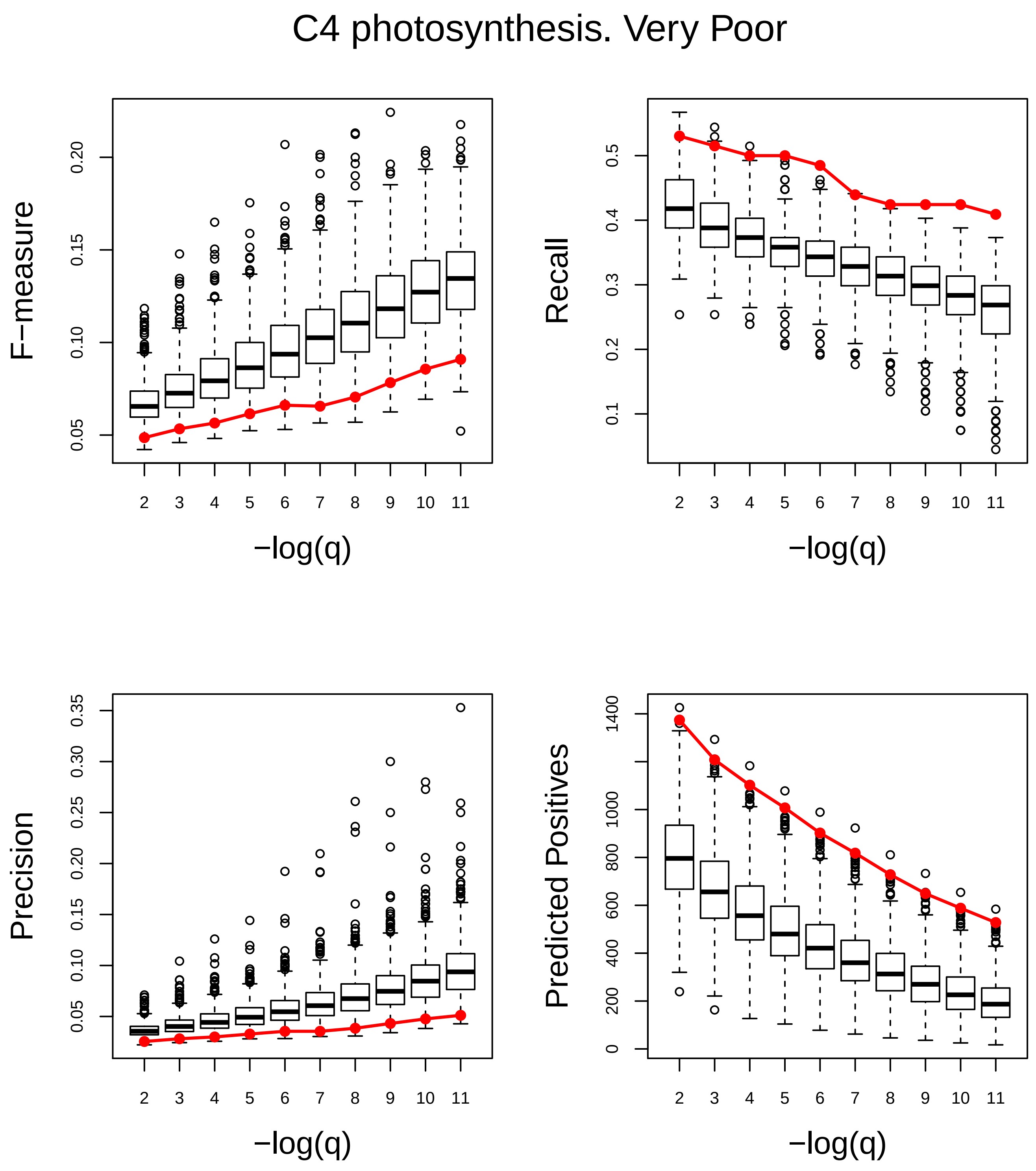
C4 photosynthesis. Very Poor
F−measure
0.05
0.10
0.15
0.20
Recall
0.1
0.2
0.3
0.4
0.5
Precision
0.05
0.10
0.15
0.20
0.25
0.30
0.35
Predicted Positives
0
200
400
600
800
1000
1400
2 3 4 5 6 7 8 9 10 11
−log(q)

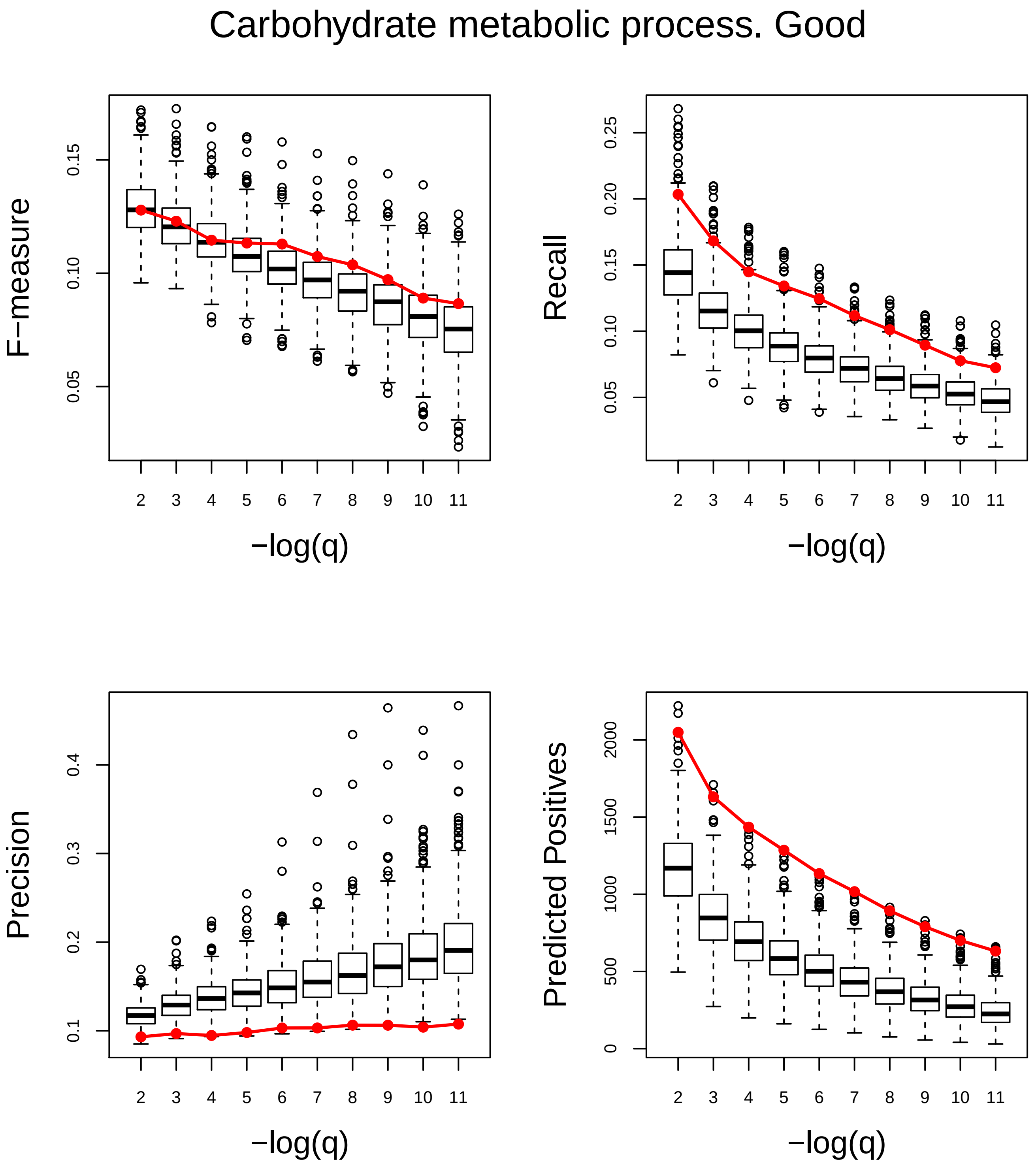
Carbohydrate metabolic process. Good
F−measure
0.05
0.10
0.15
Recall
0.05
0.10
0.15
0.20
0.25
Precision
0.1
0.2
0.3
0.4
Predicted Positives
0
500
1000
1500
2000
2 3 4 5 6 7 8 9 10 11
−log(q)

### Catabolic process. Very Good

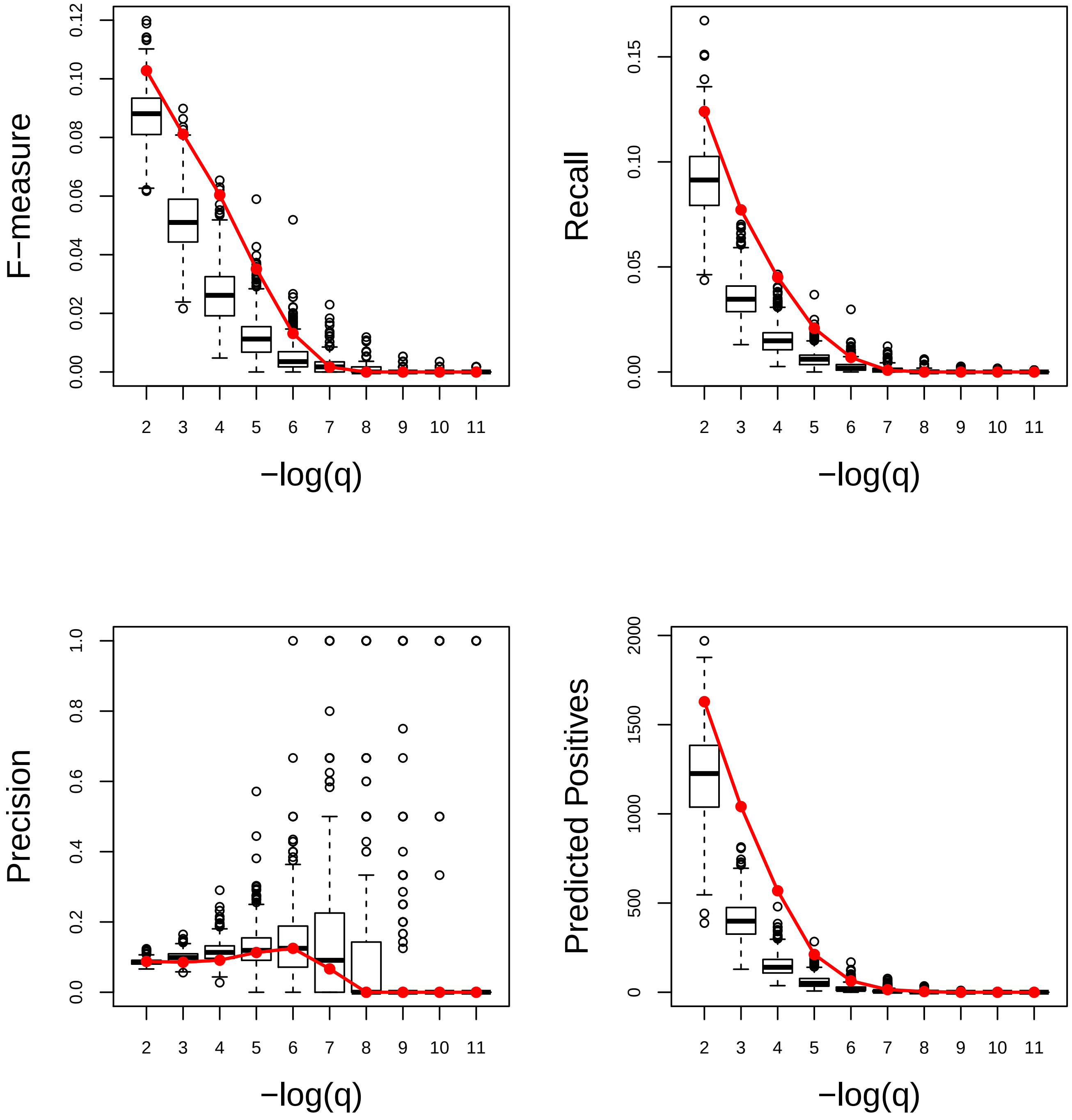

#### Cell communication. Very Good

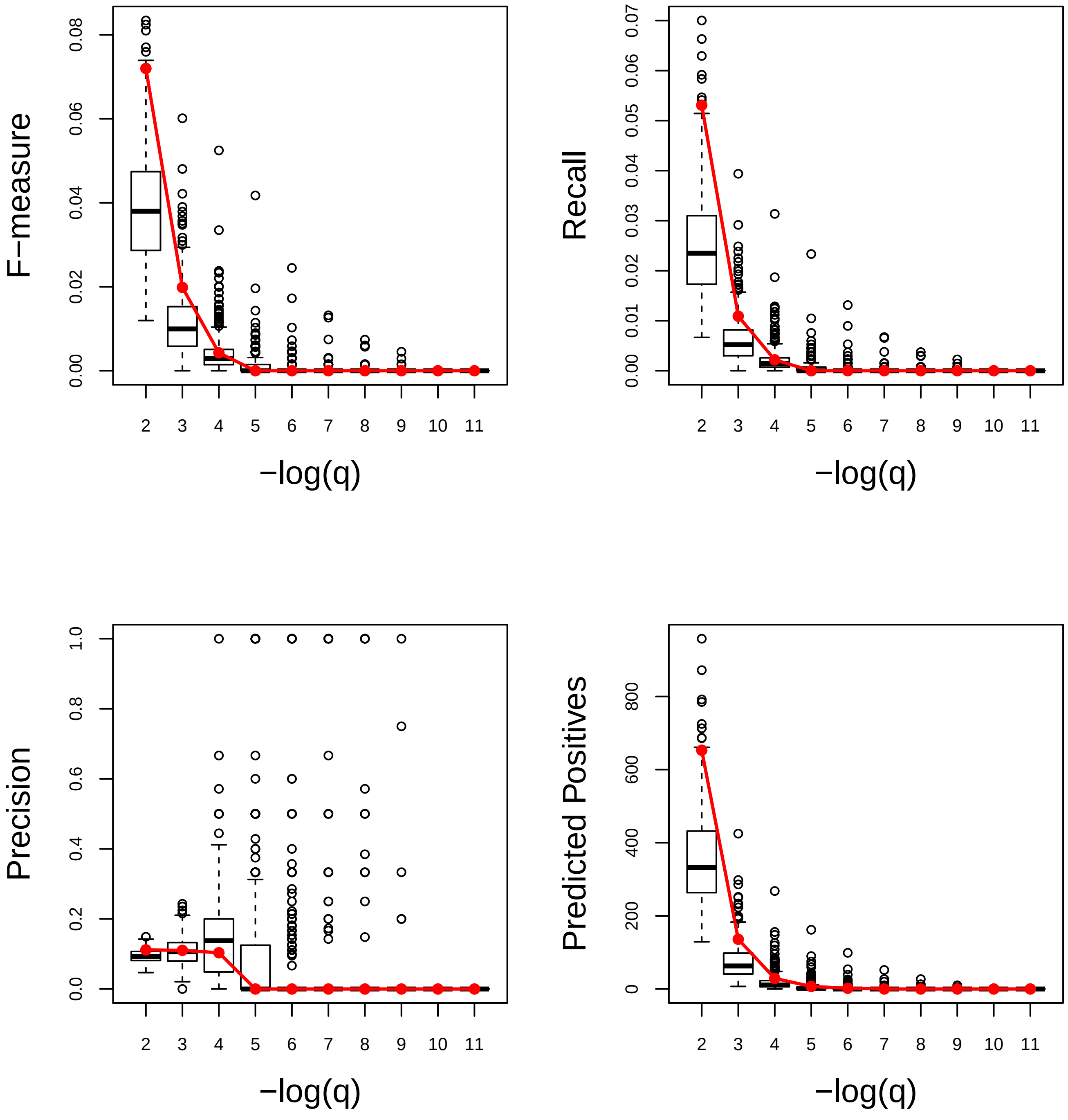

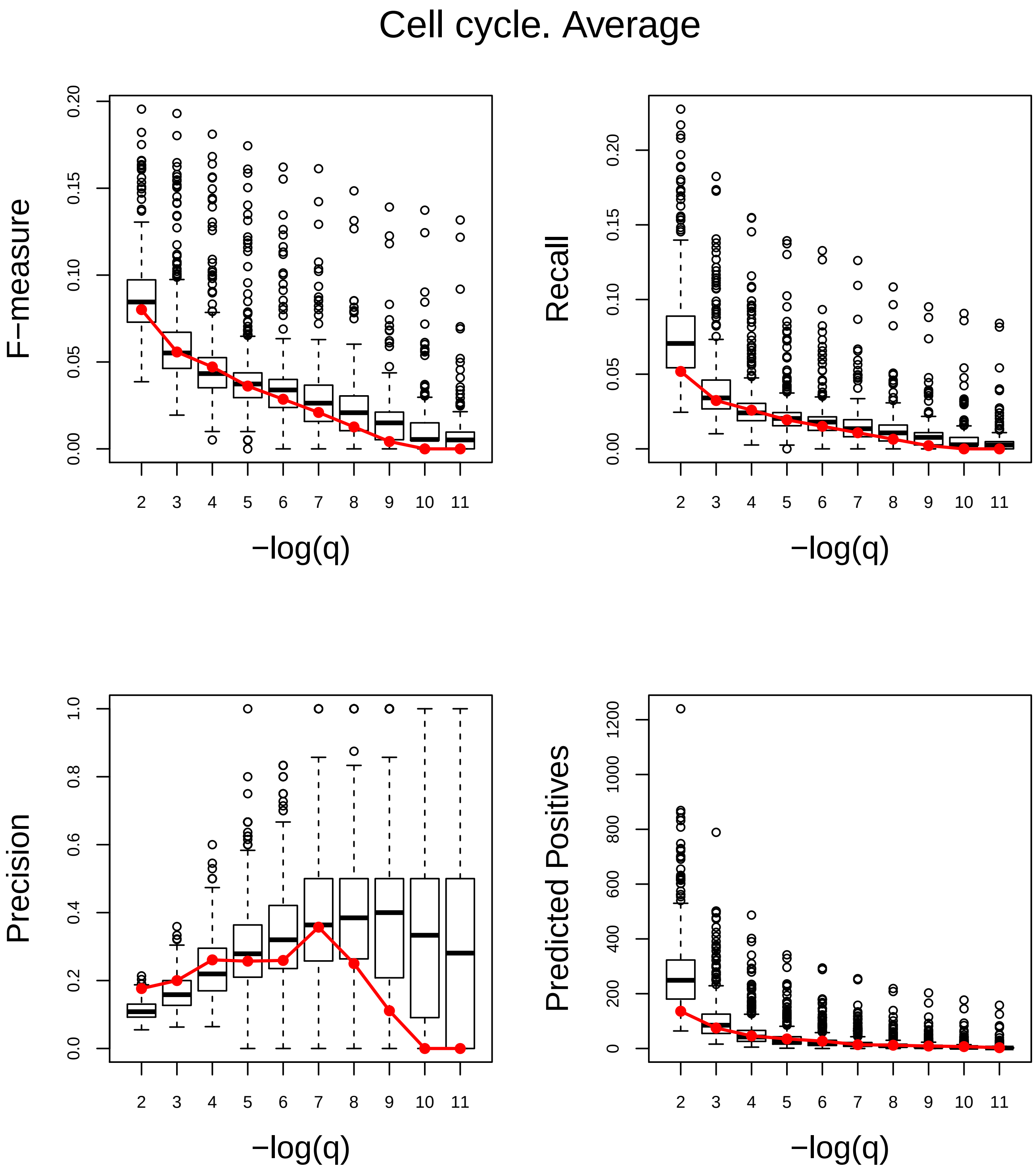
Cell cycle. Average
F−measure
Recall
Precision
Predicted Positives
−log(q)
−log(q)
−log(q)
−log(q)

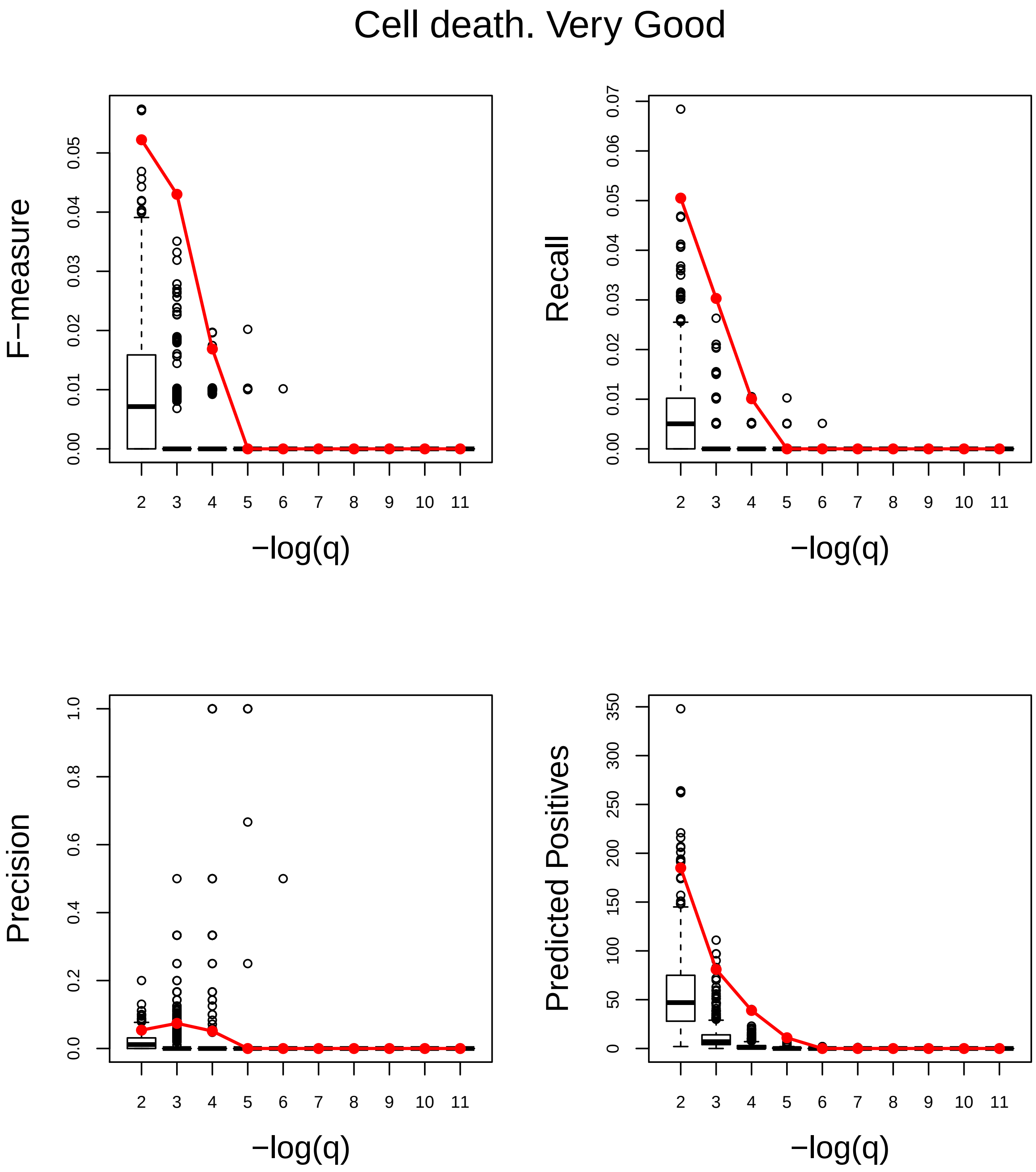
Cell death. Very Good
F−measure
Recall
Precision
Predicted Positives
−log(q)
−log(q)
−log(q)
−log(q)

### Cell differentiation. Average

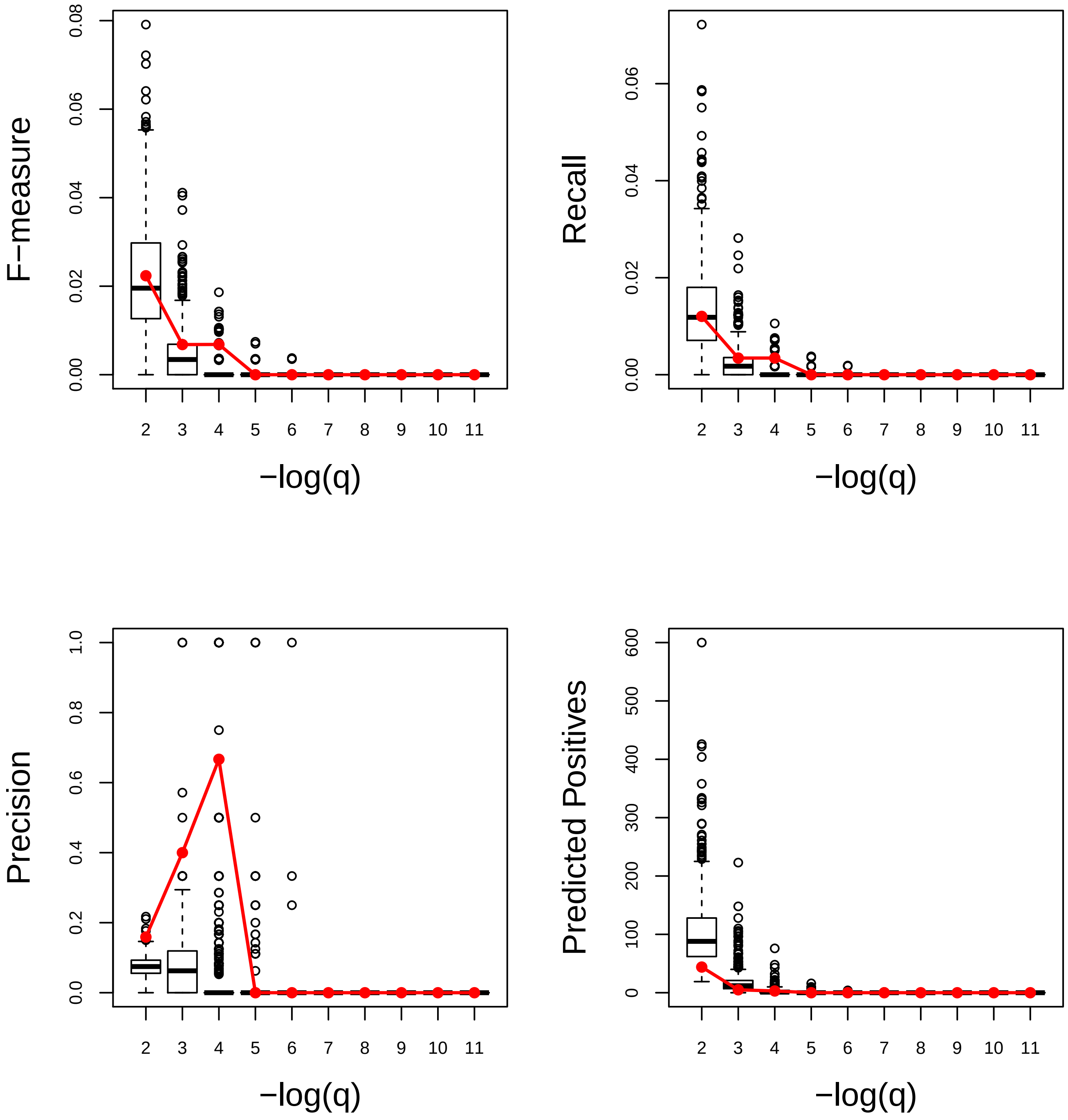

### Cell division. Very Poor

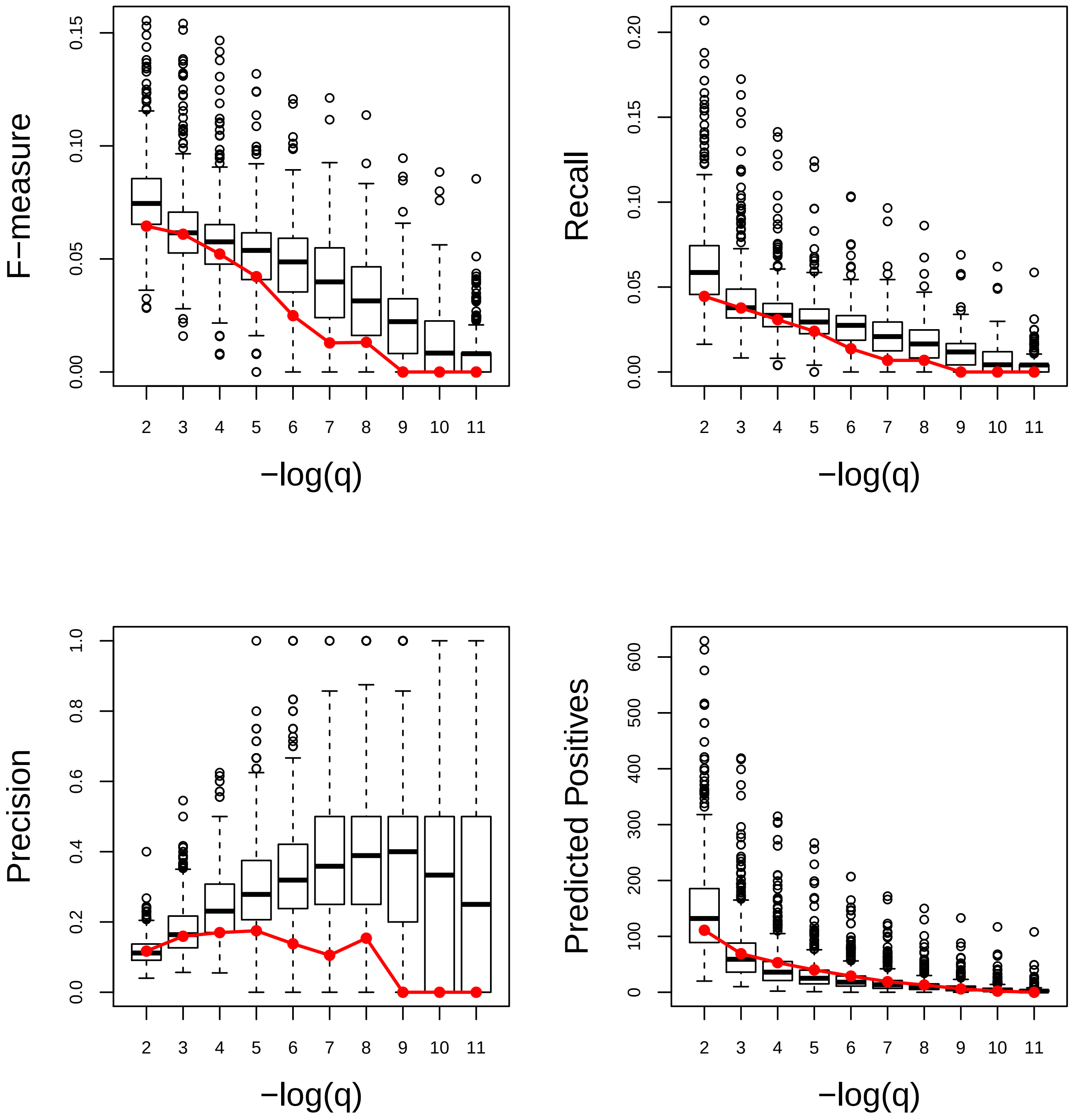

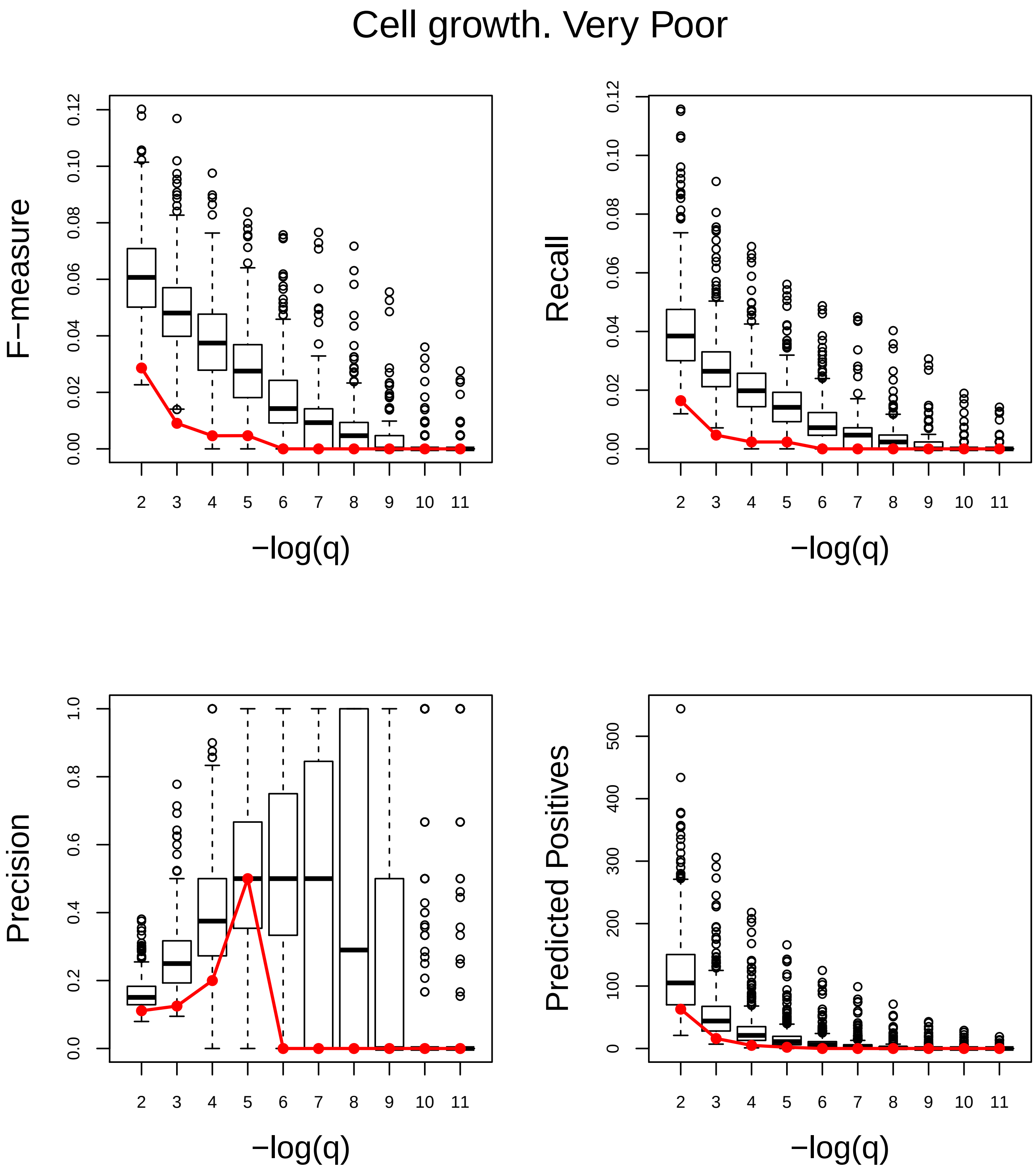
Cell growth. Very Poor
F−measure
0.00
0.02
0.04
0.06
0.08
0.10
0.12
−log(q)
Recall
Precision
0.0
0.2
0.4
0.6
0.8
1.0
Predicted Positives
0
100
200
300
400
500

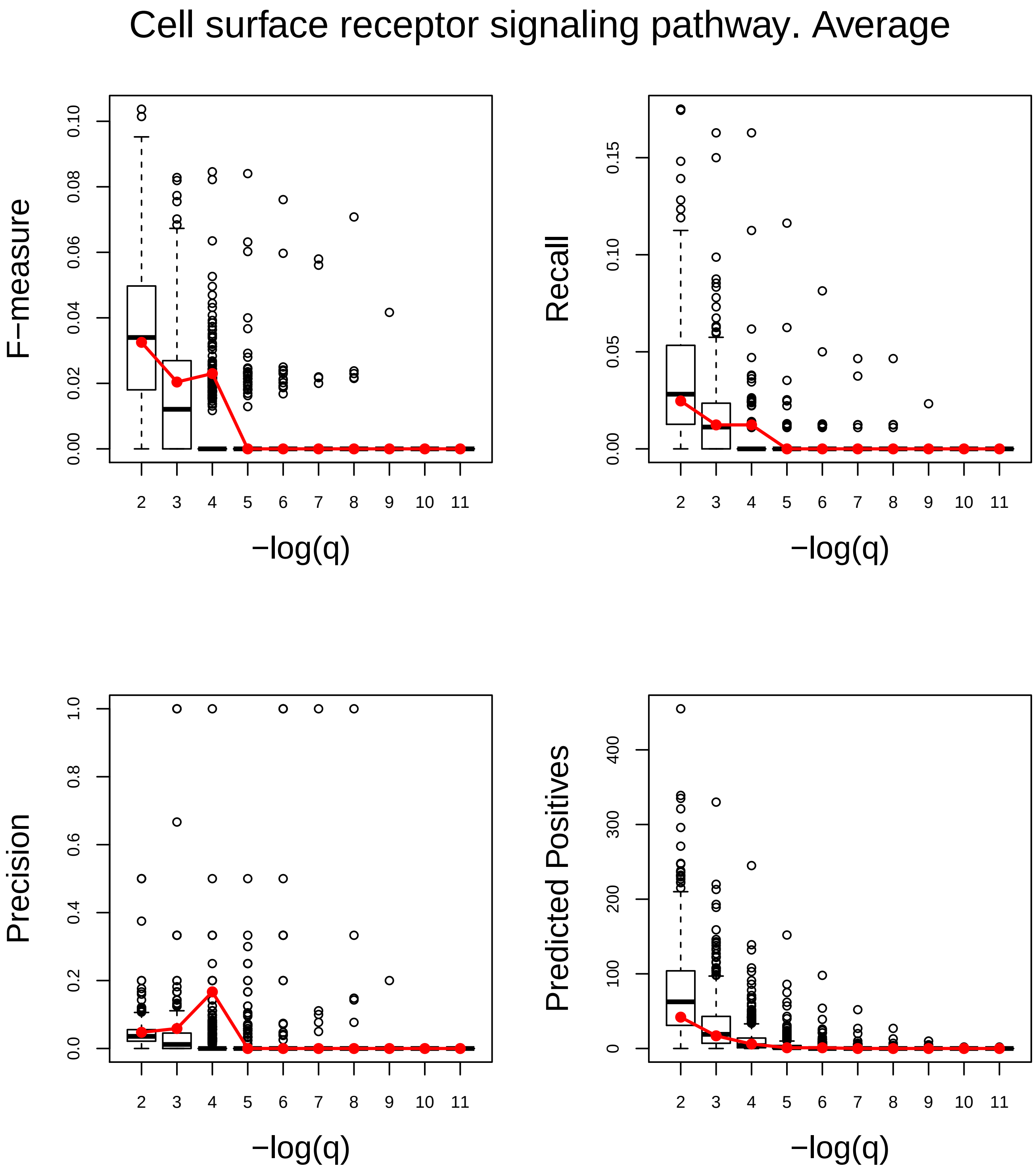
Cell surface receptor signaling pathway. Average
F−measure
Recall
Precision
Predicted Positives
−log(q)
−log(q)
−log(q)
−log(q)

### Cell wall macromolecule metabolic process. Average

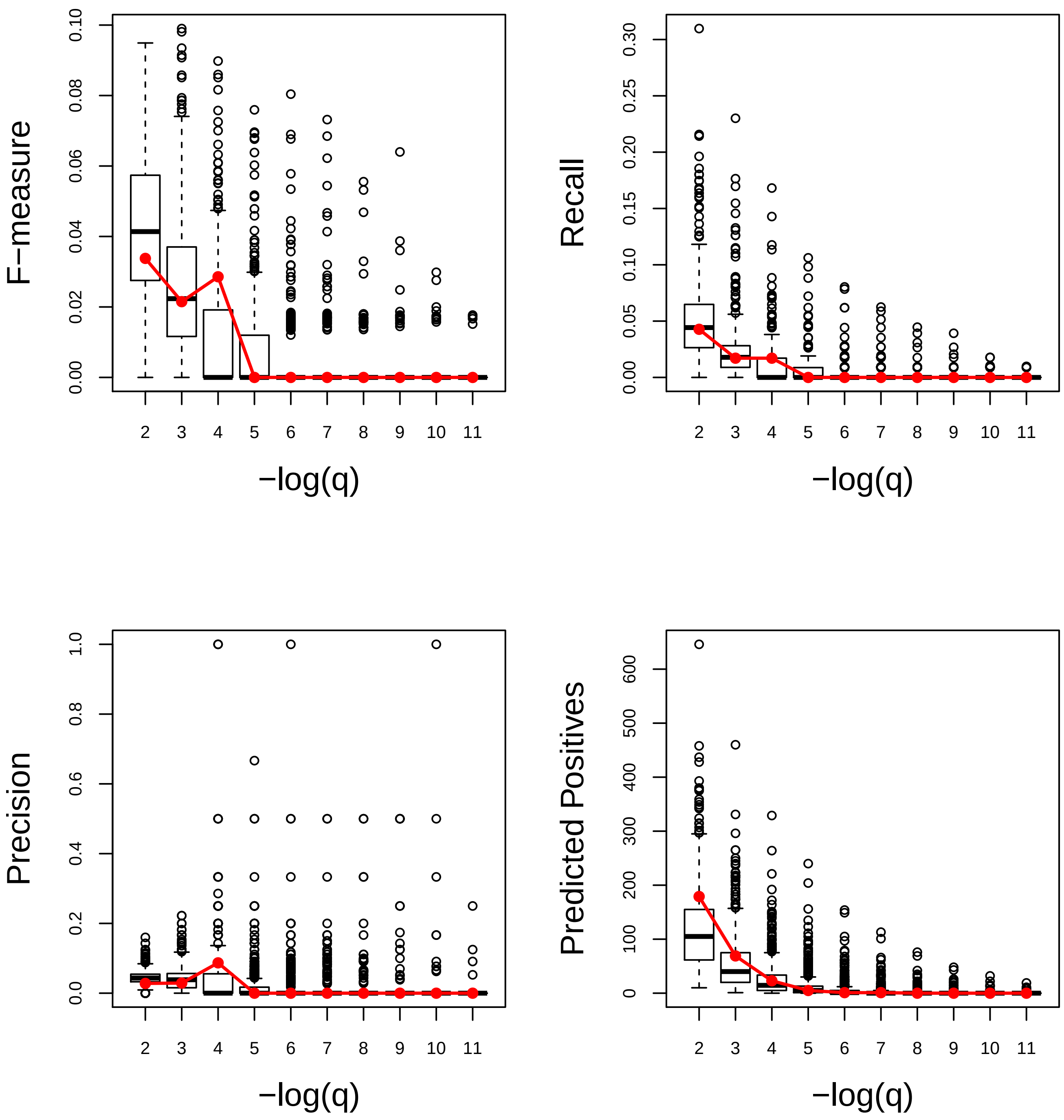

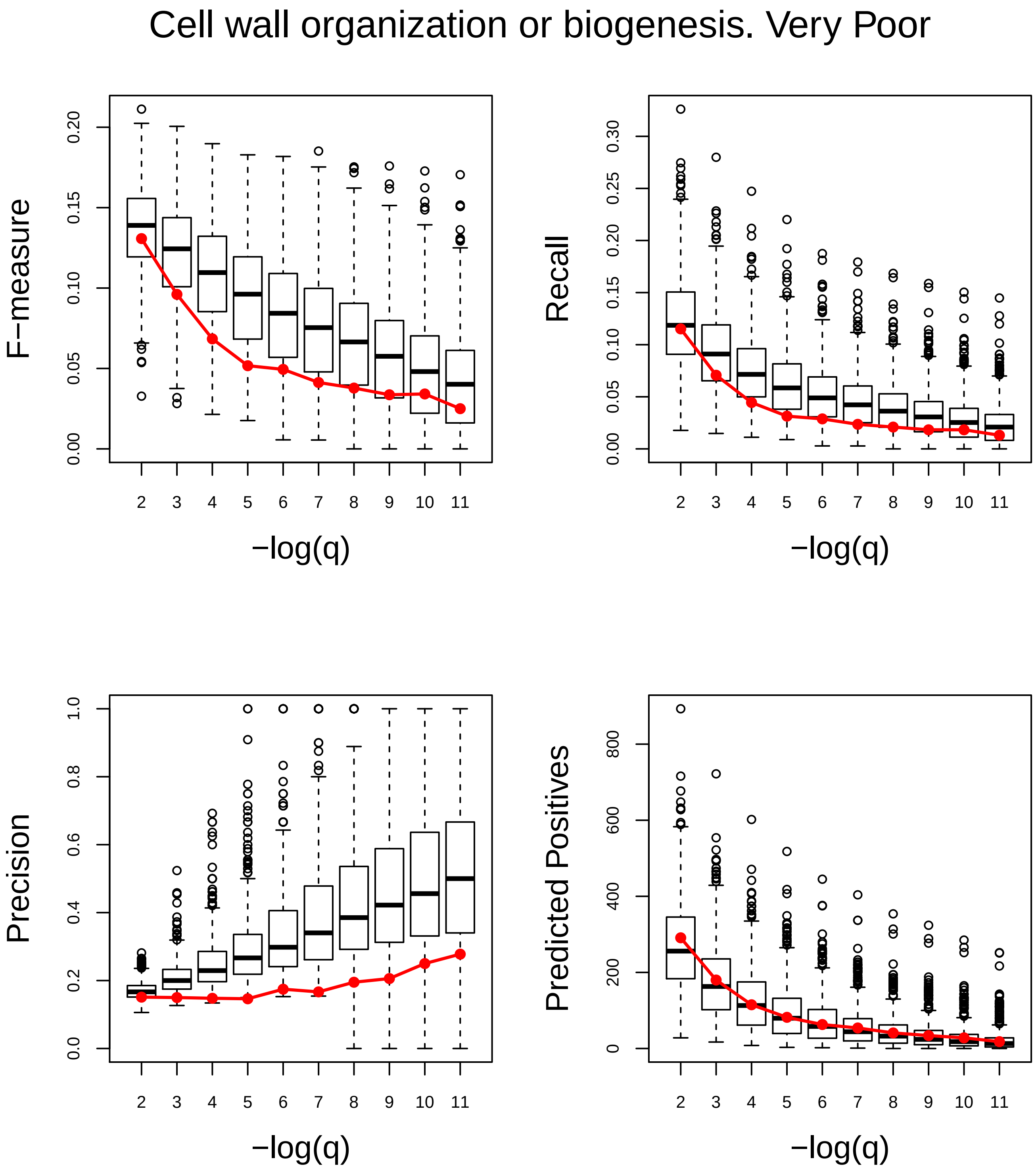
Cell wall organization or biogenesis. Very Poor
F−measure
0.00
0.05
0.10
0.15
0.20
Recall
0.00
0.05
0.10
0.15
0.20
0.25
0.30
Precision
0.0
0.2
0.4
0.6
0.8
1.0
Predicted Positives
0
200
400
600
800
2 3 4 5 6 7 8 9 10 11
−log(q)

### Cell wall polysaccharide metabolic process. Very Poor

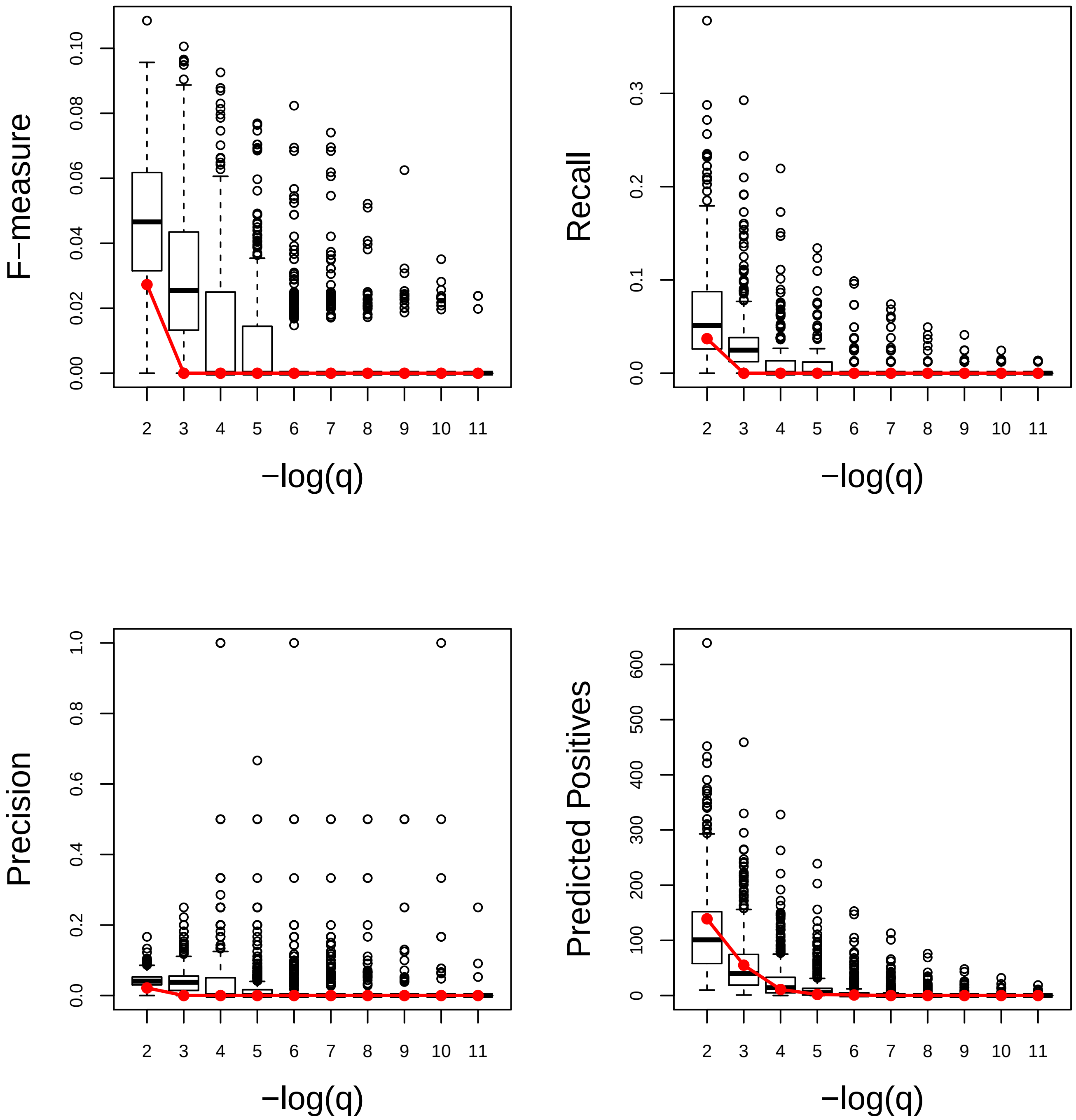

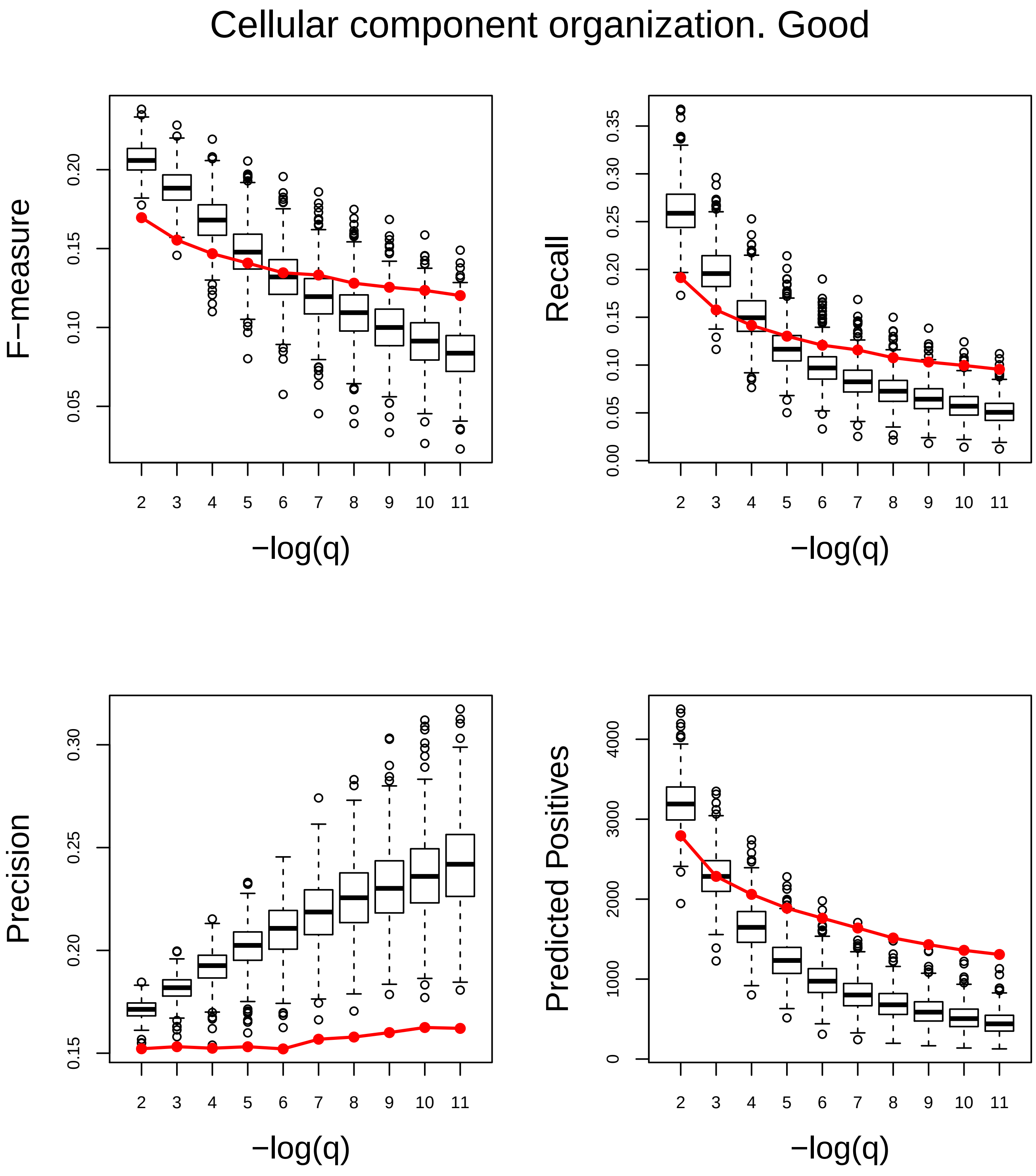
Cellular component organization. Good
F−measure
Recall
Precision
Predicted Positives
−log(q)
−log(q)
−log(q)
−log(q)
0.05
0.10
0.15
0.20
0.00
0.05
0.10
0.15
0.20
0.25
0.30
0.35
0.15
0.20
0.25
0.30
0
1000
2000
3000
4000
2 3 4 5 6 7 8 9 10 11

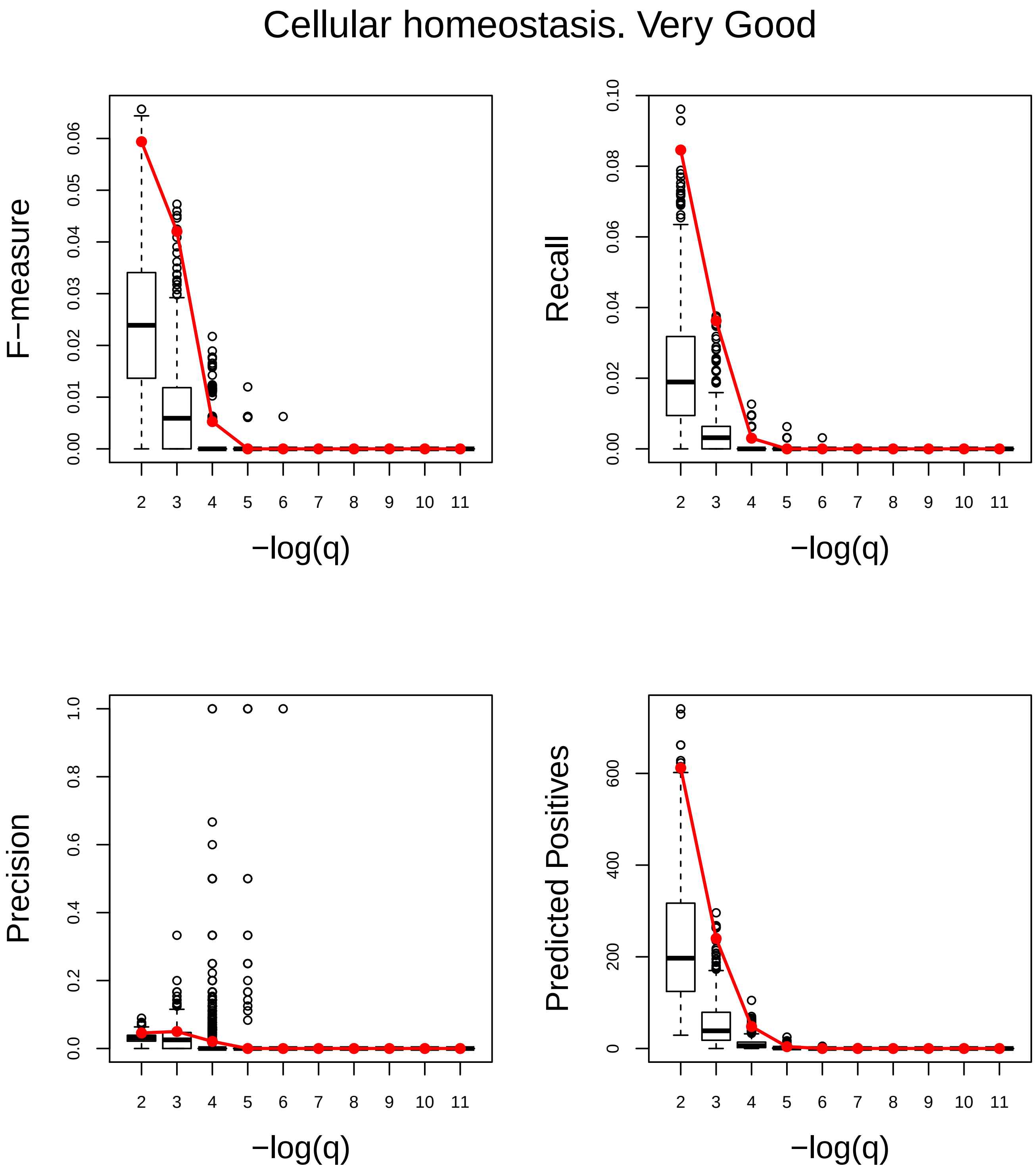
Cellular homeostasis. Very Good
F−measure
0.00
0.01
0.02
0.03
0.04
0.05
0.06
Recall
0.00
0.02
0.04
0.06
0.08
0.10
Precision
0.0
0.2
0.4
0.6
0.8
1.0
Predicted Positives
0
200
400
600
2 3 4 5 6 7 8 9 10 11
−log(q)

### Cellular protein modification process. Very Poor

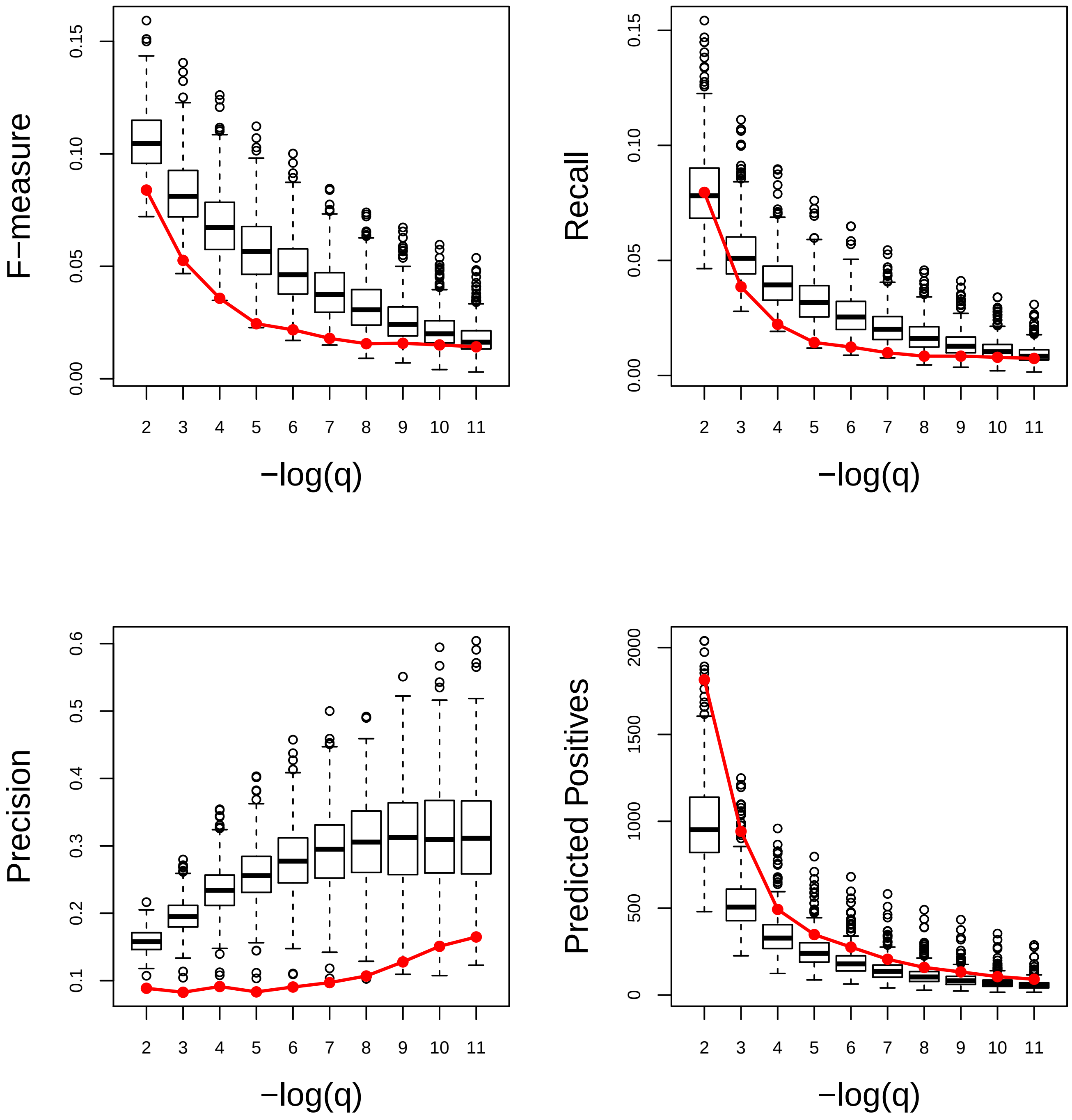

### Cellular response to DNA damage stimulus. Very Poor

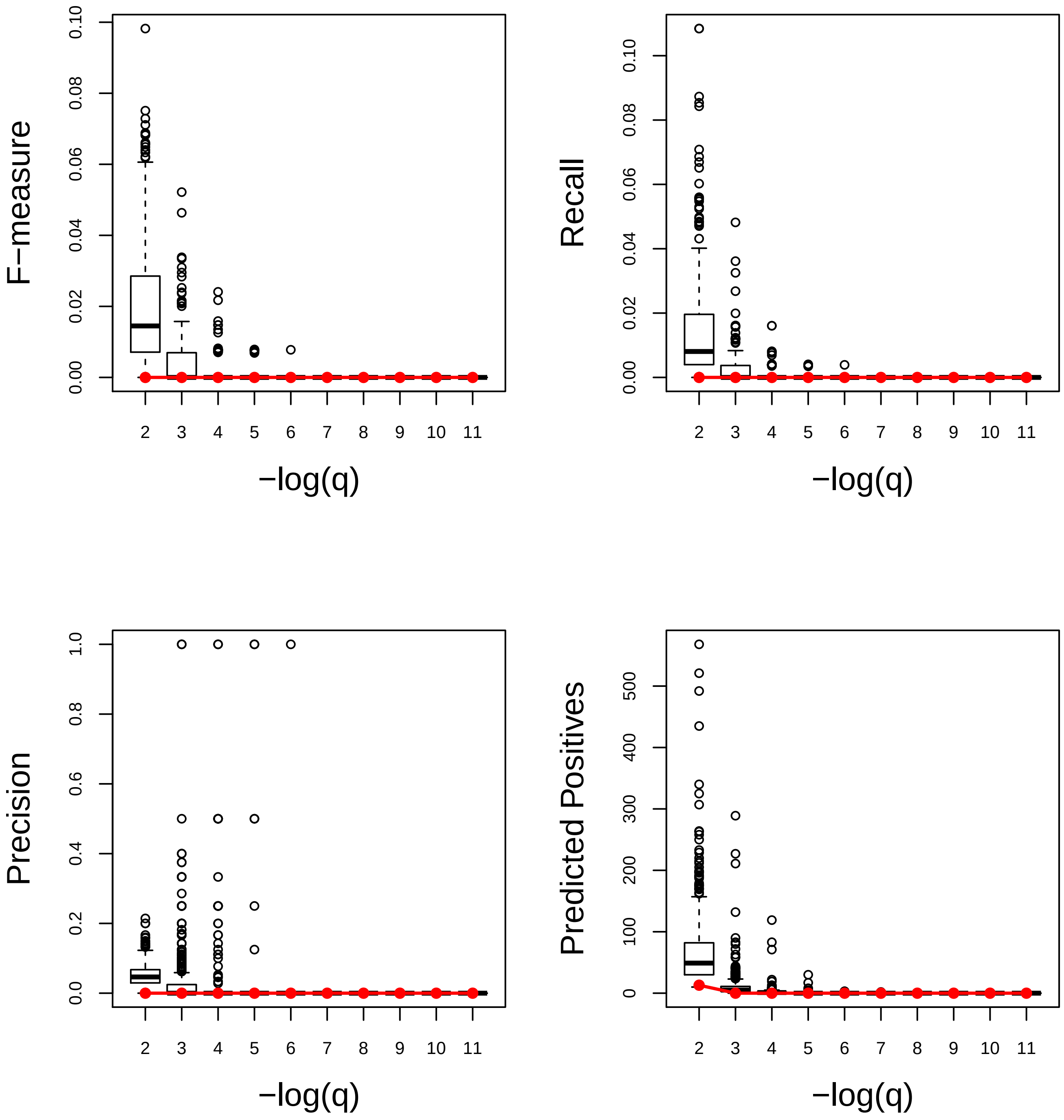

### Cellulose metabolic process. Very Poor

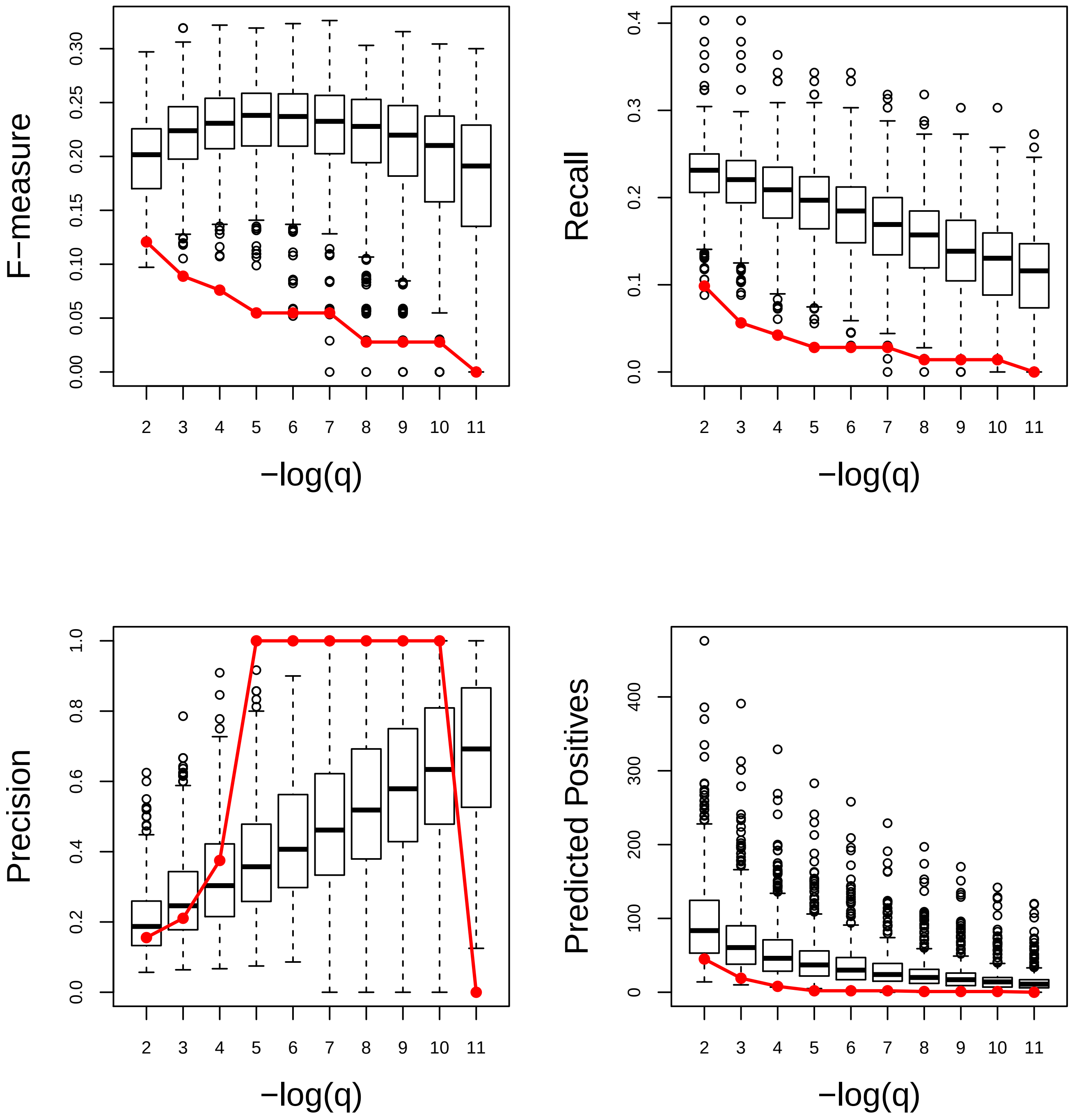

### Cofactor metabolic process. Very Good

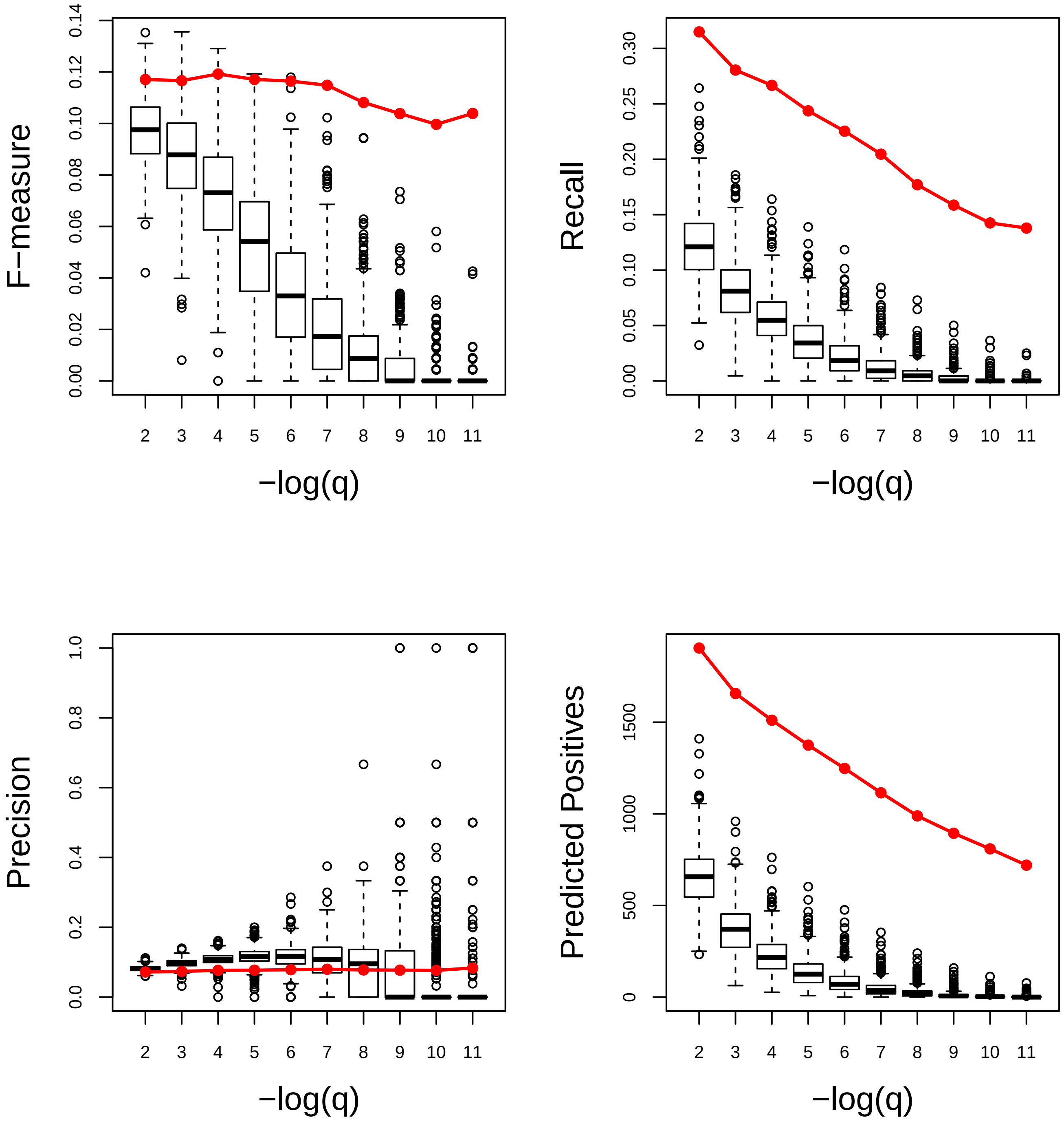

### Defense response to bacterium. Very Good

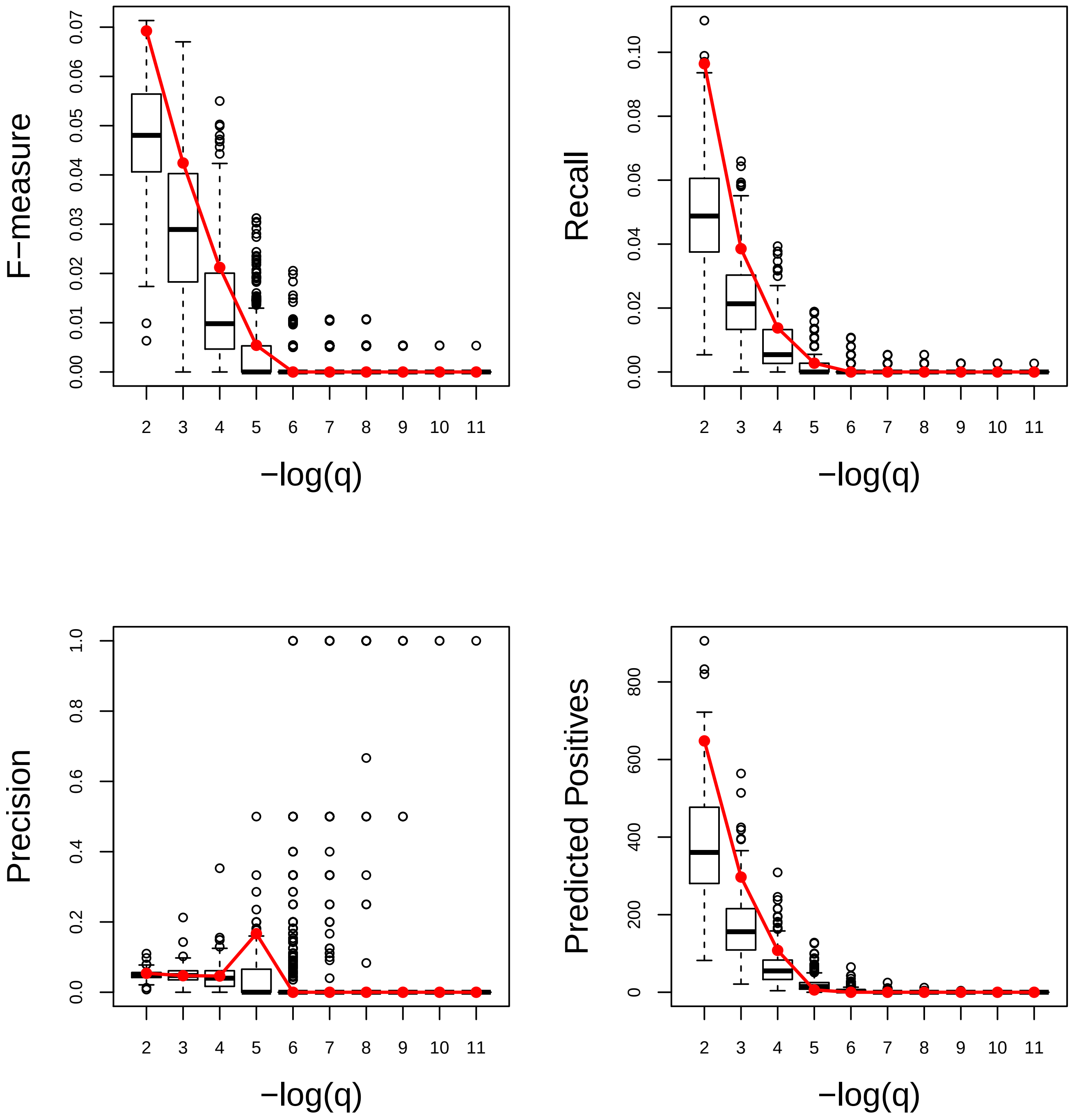

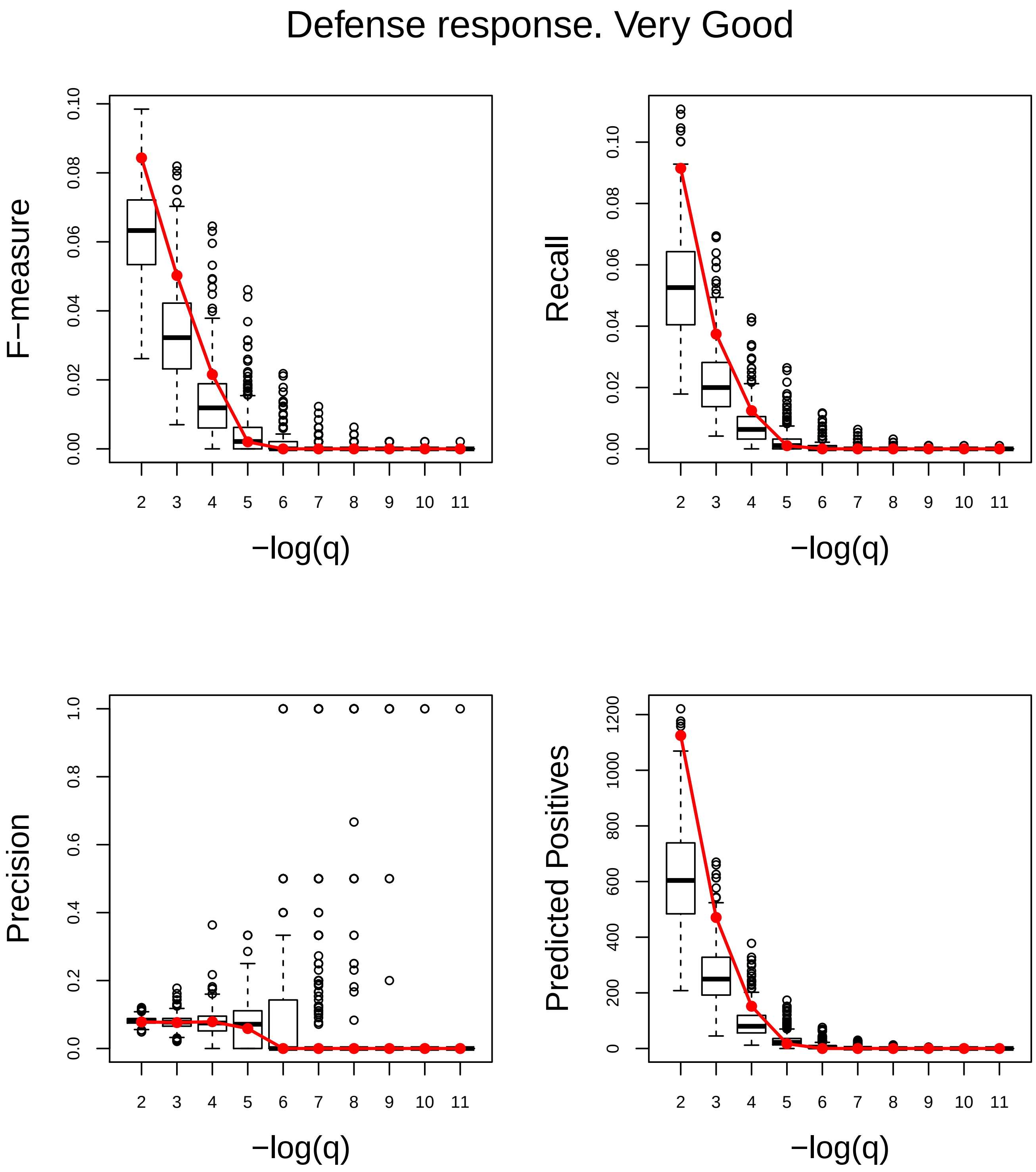
Defense response. Very Good
F−measure
Recall
Precision
Predicted Positives
−log(q)
−log(q)
−log(q)
−log(q)
0.00
0.02
0.04
0.06
0.08
0.10
0.0
0.2
0.4
0.6
0.8
1.0
0
200
400
600
800
1000
1200
2 3 4 5 6 7 8 9 10 11

### DNA metabolic process. Very Poor

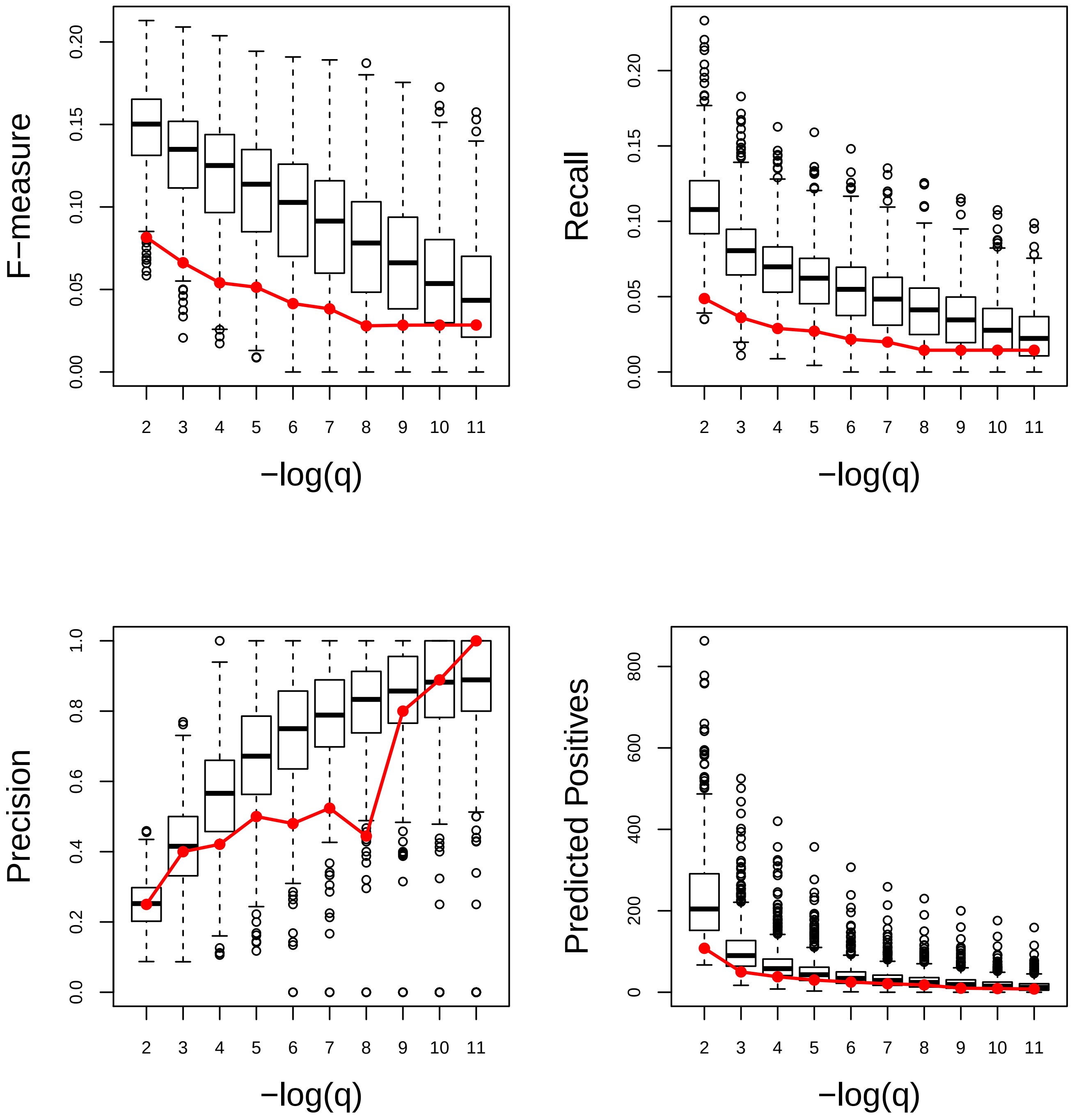

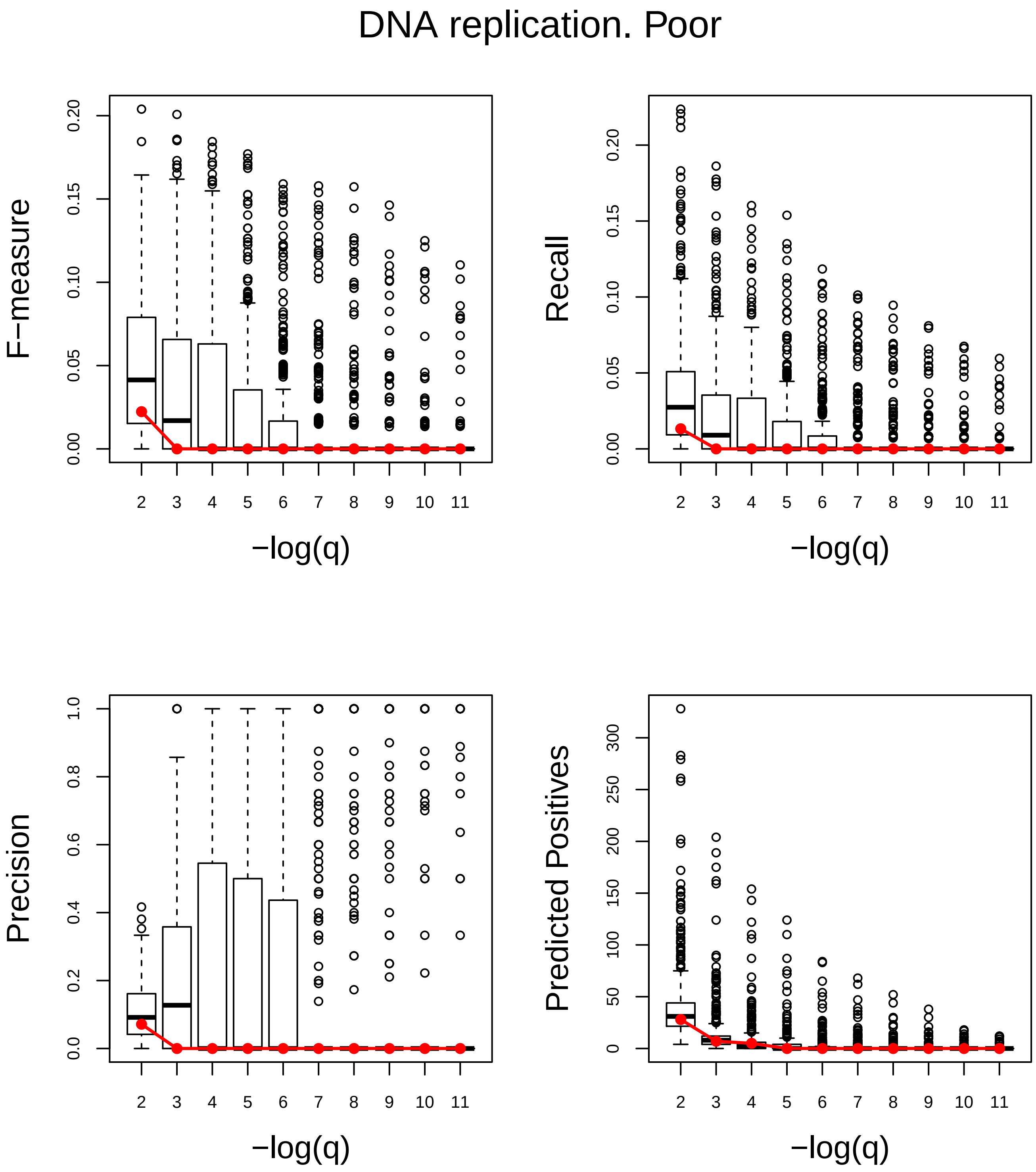
DNA replication. Poor
F−measure
Recall
Precision
Predicted Positives
−log(q)
−log(q)
−log(q)
−log(q)

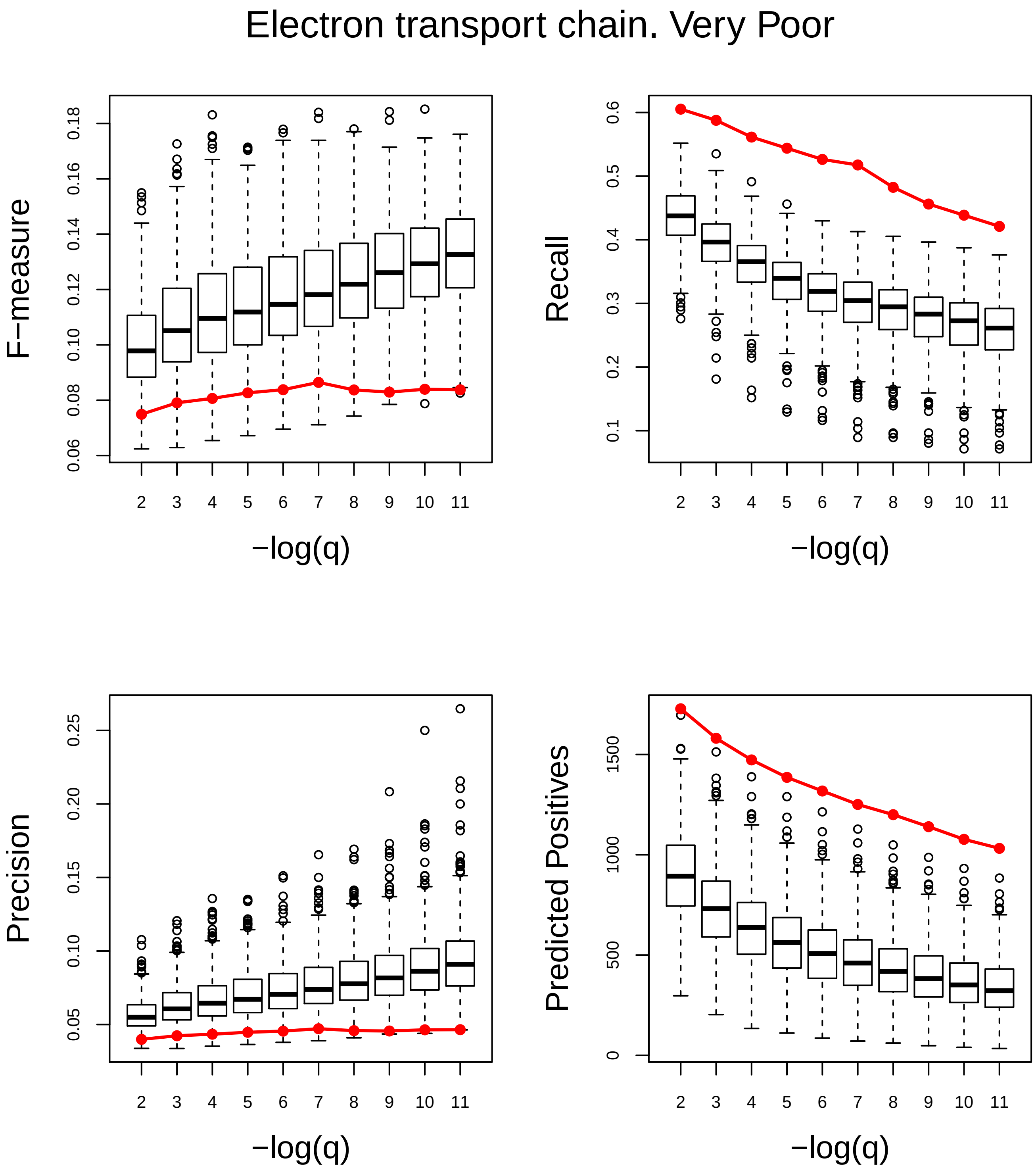
Electron transport chain. Very Poor
F−measure
Recall
Precision
Predicted Positives
−log(q)
−log(q)
−log(q)
−log(q)

### Embryo development. Very Good

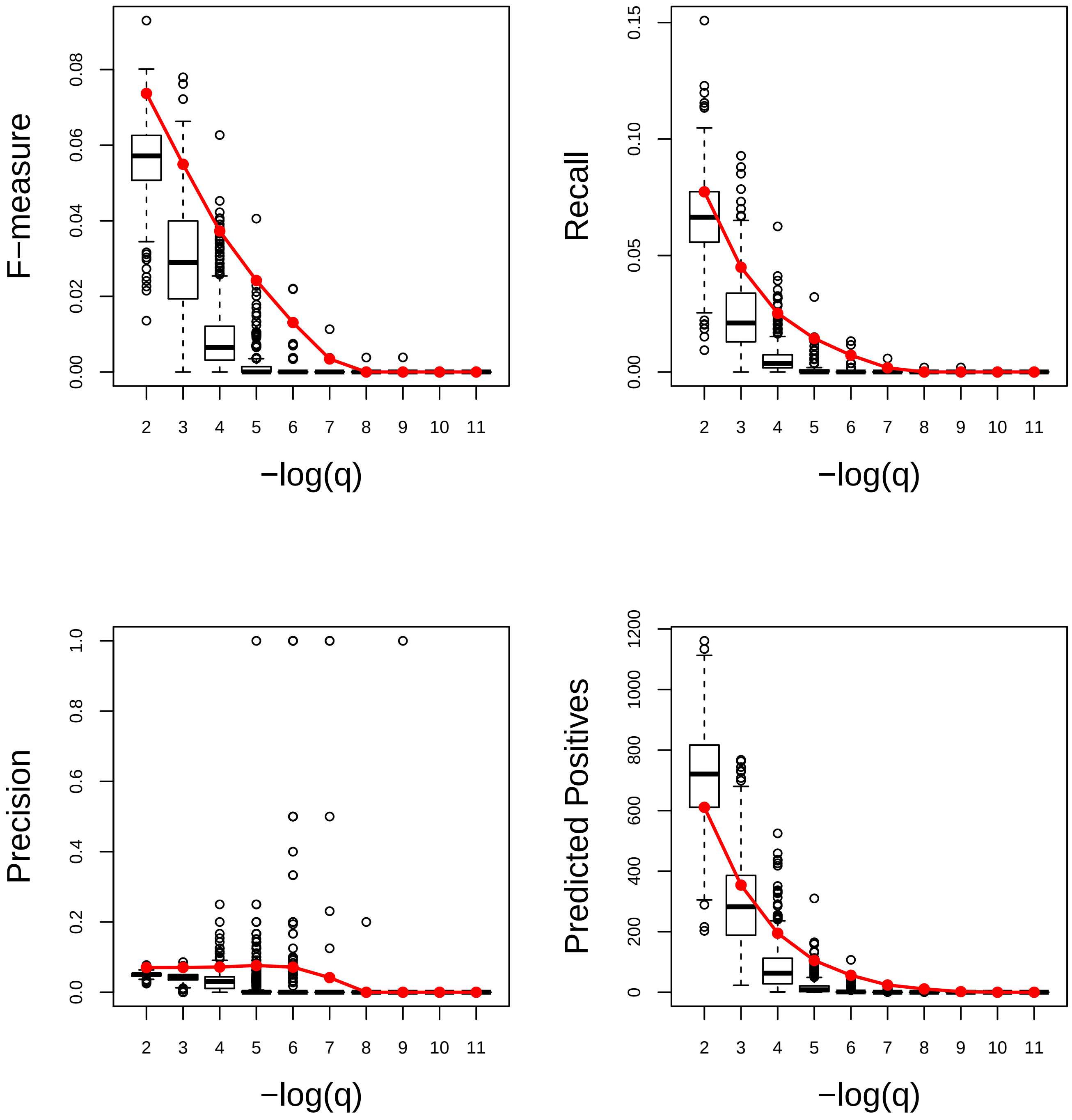

### External encapsulating structure organization. Very Poor

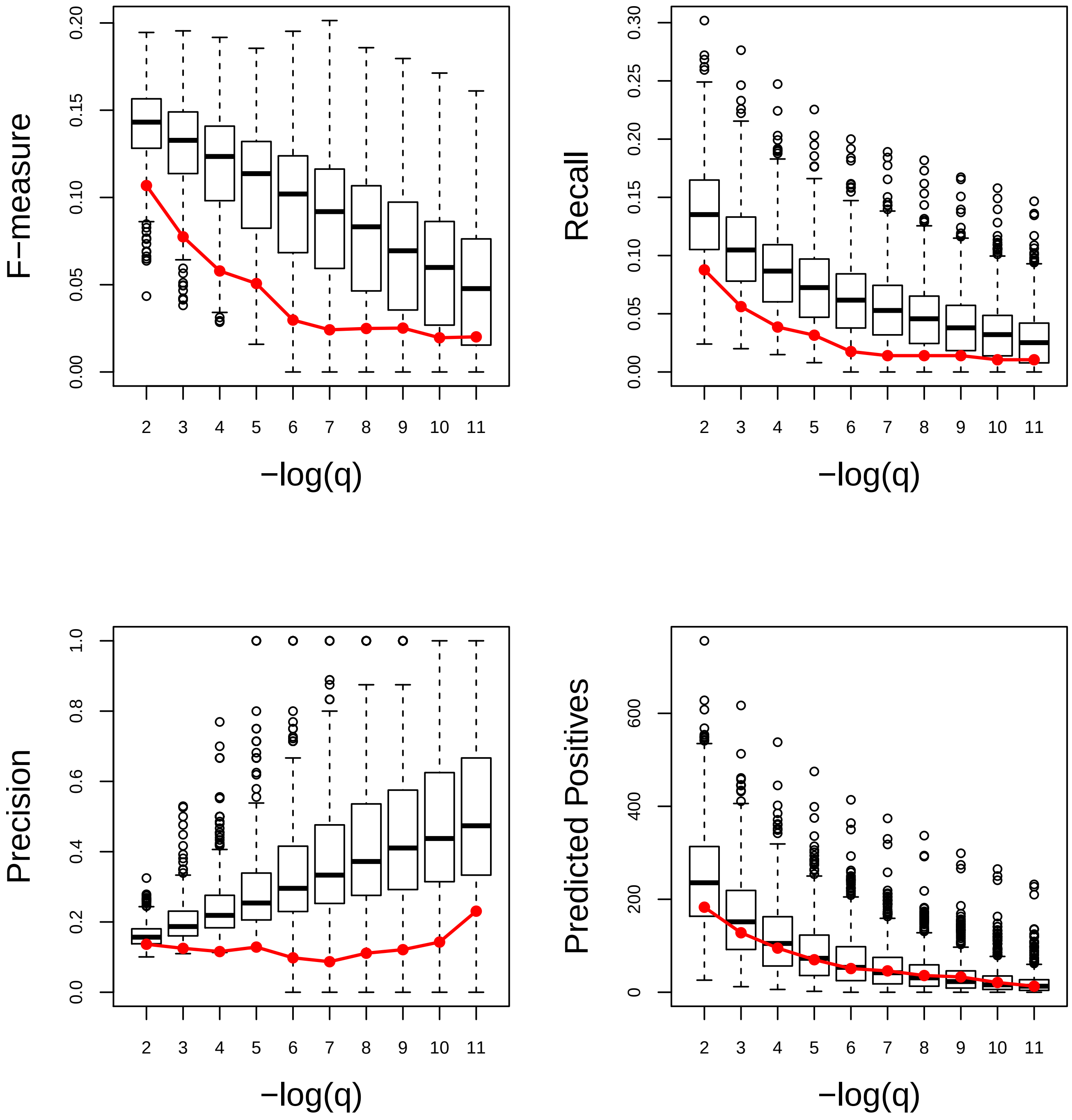

### Flower development. Very Poor

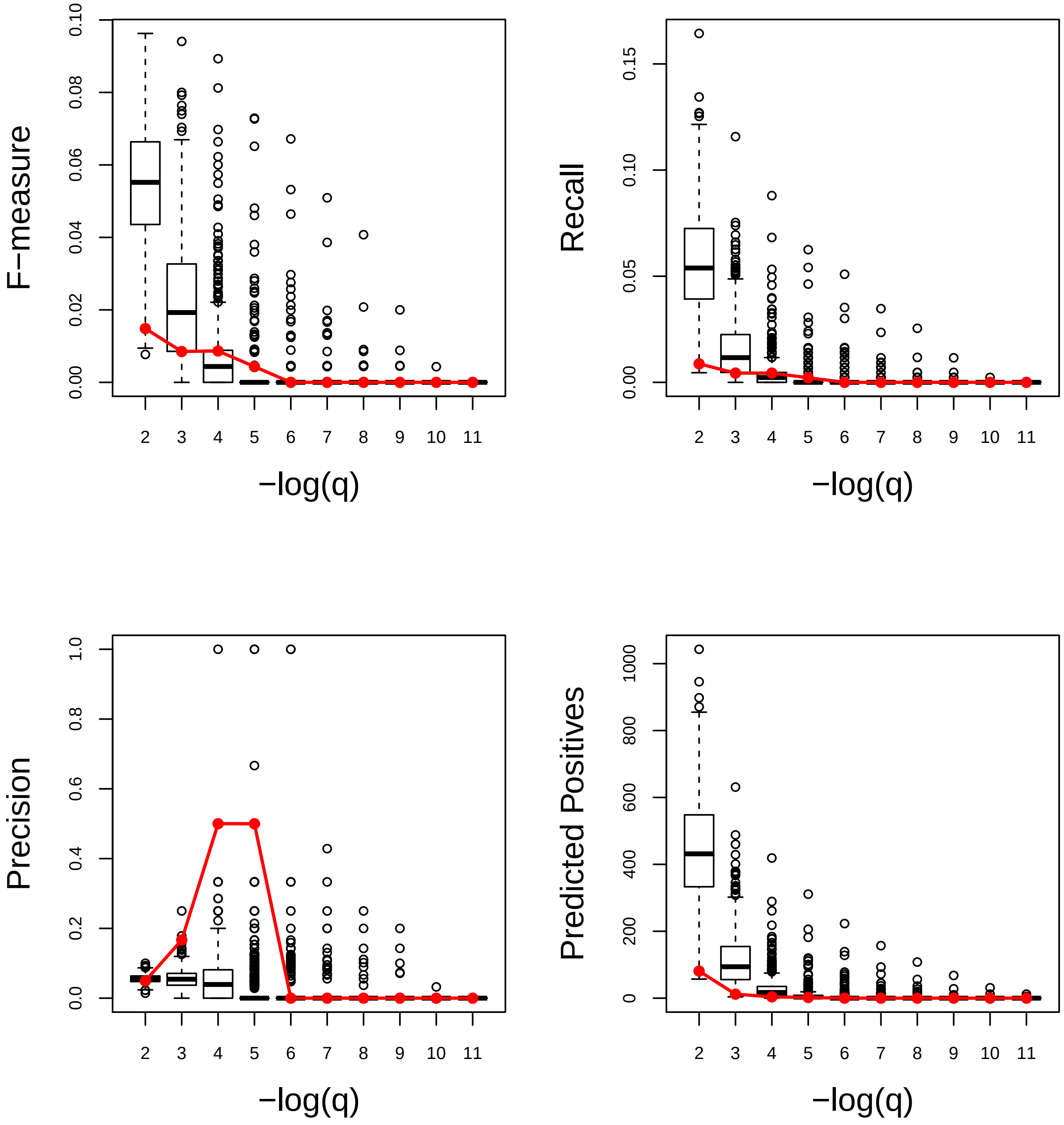

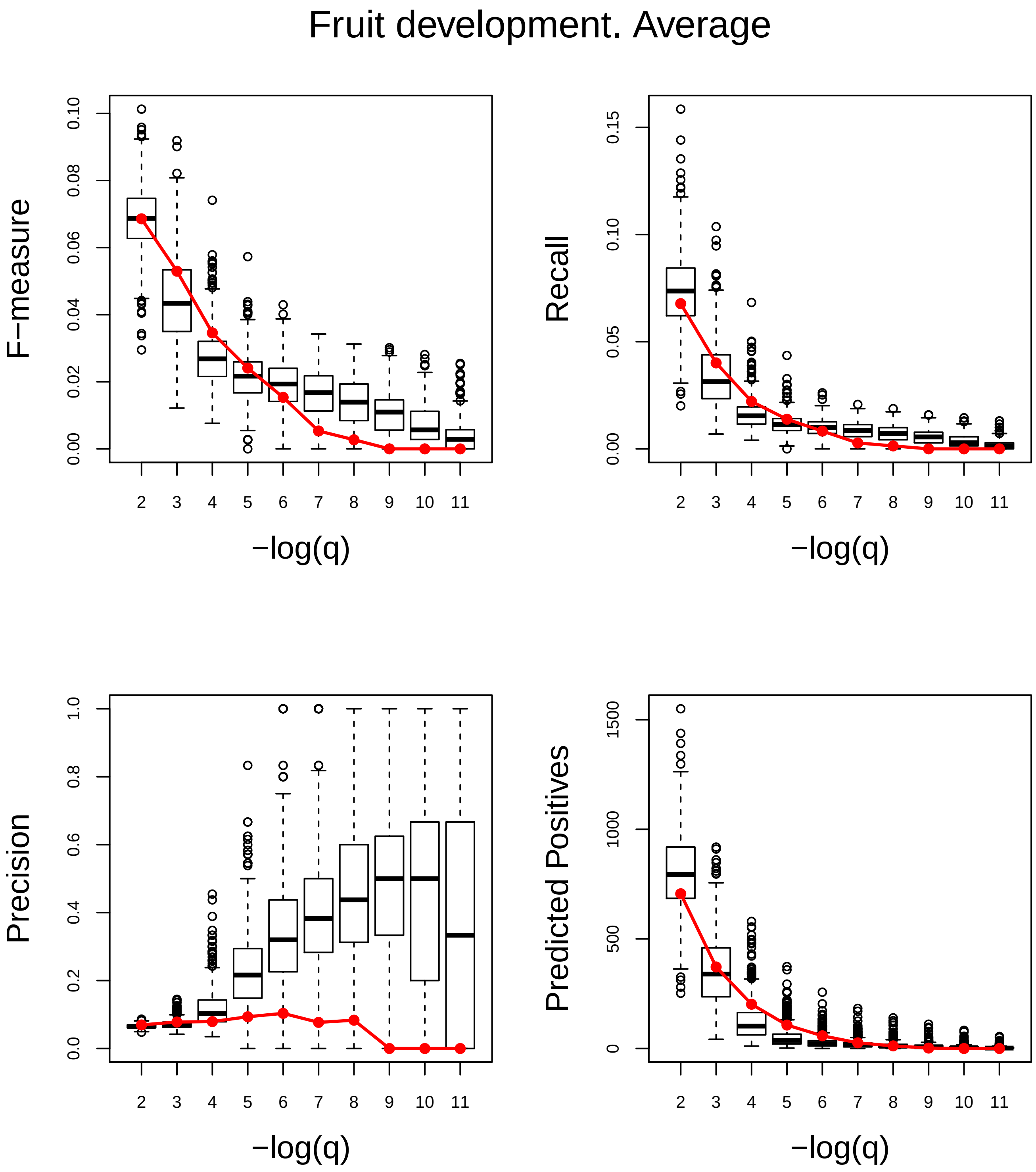
Fruit development. Average
F−measure
0.00
0.02
0.04
0.06
0.08
0.10
Recall
0.00
0.05
0.10
0.15
Precision
0.0
0.2
0.4
0.6
0.8
1.0
Predicted Positives
0
500
1000
1500
2 3 4 5 6 7 8 9 10 11
−log(q)

### Generation of precursor metabolites and energy. Very Poor

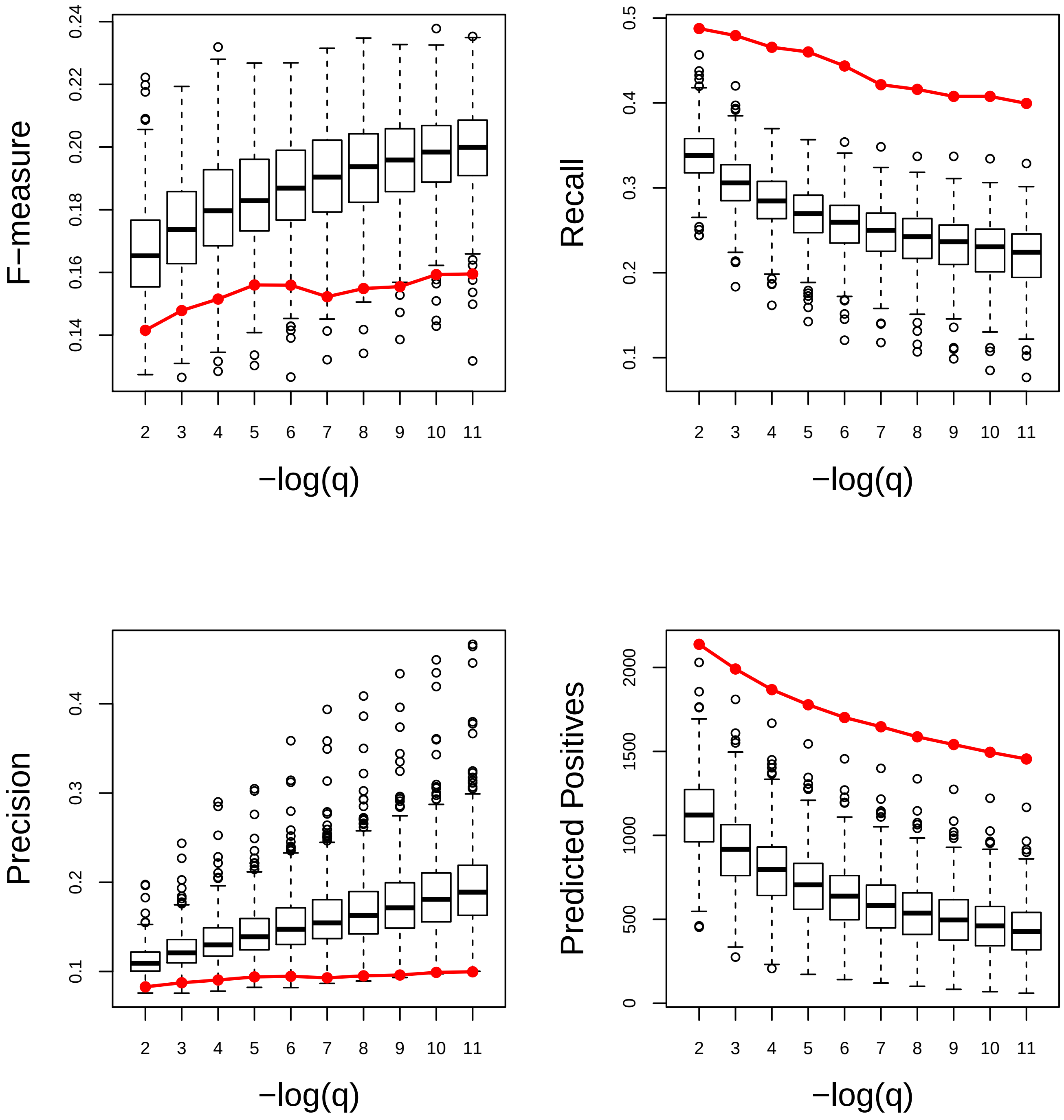

### Glucan metabolic process. Very Poor

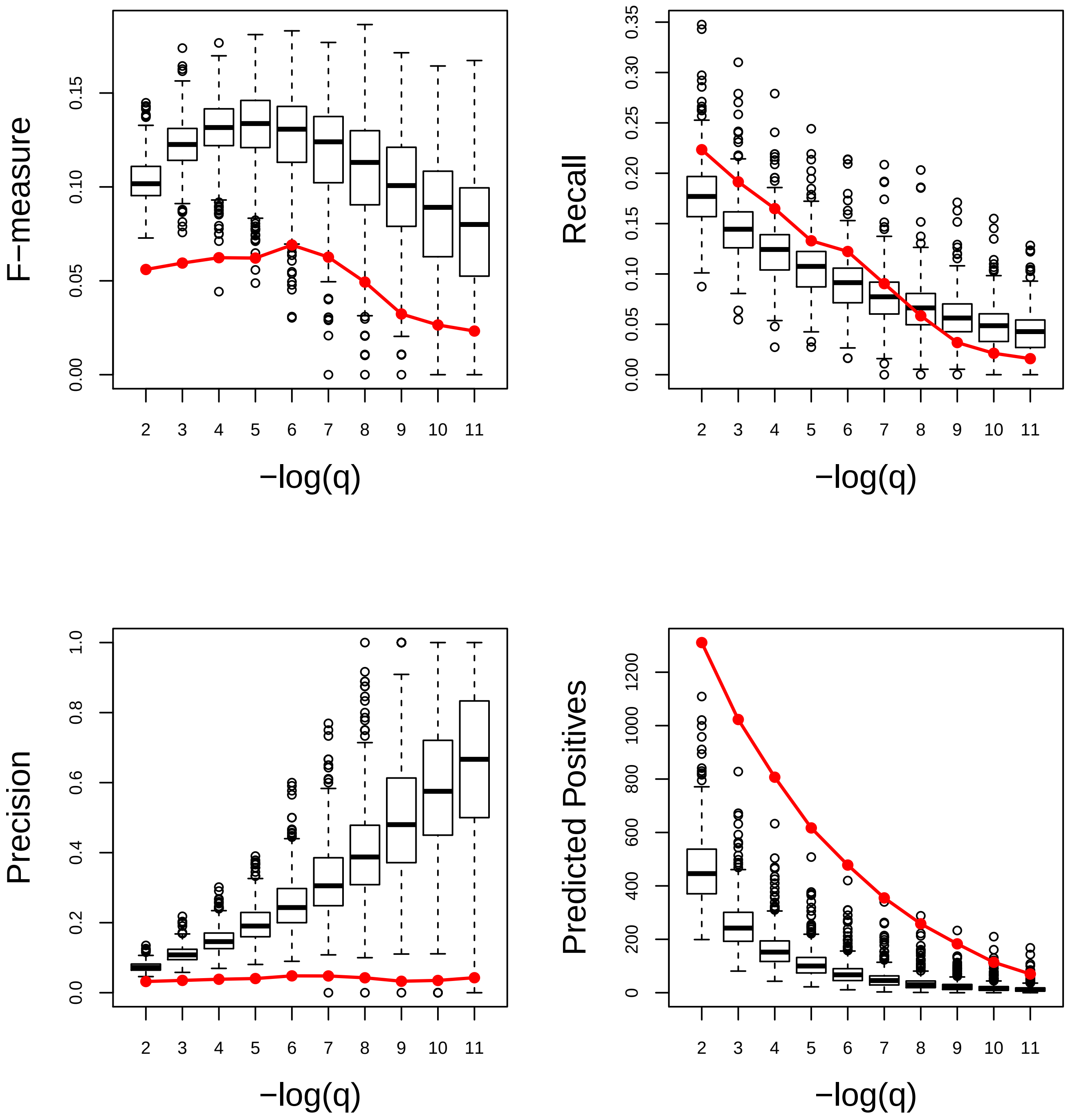

### Growth. Very Poor

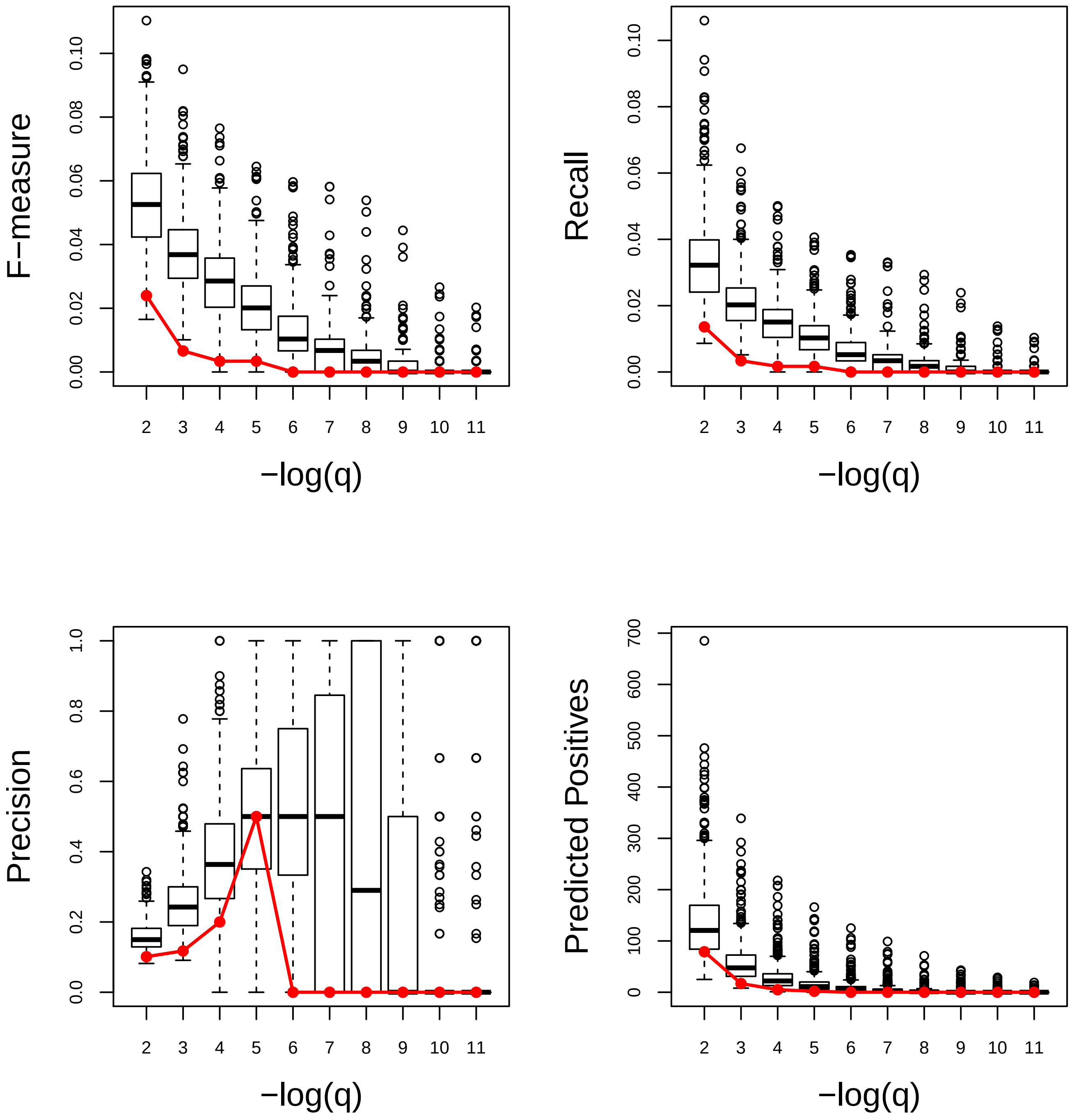

### Hemicellulose metabolic process. Poor

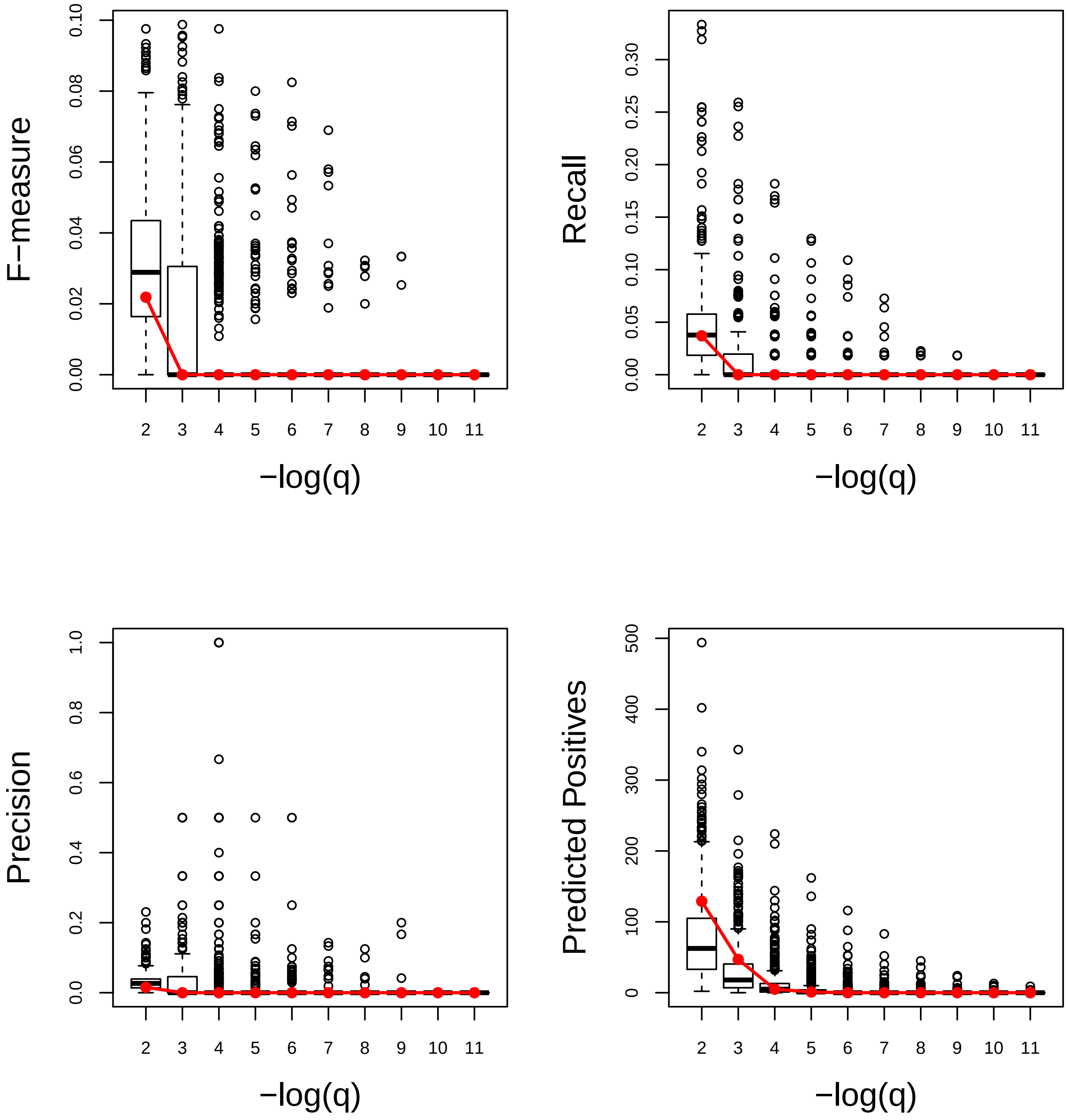

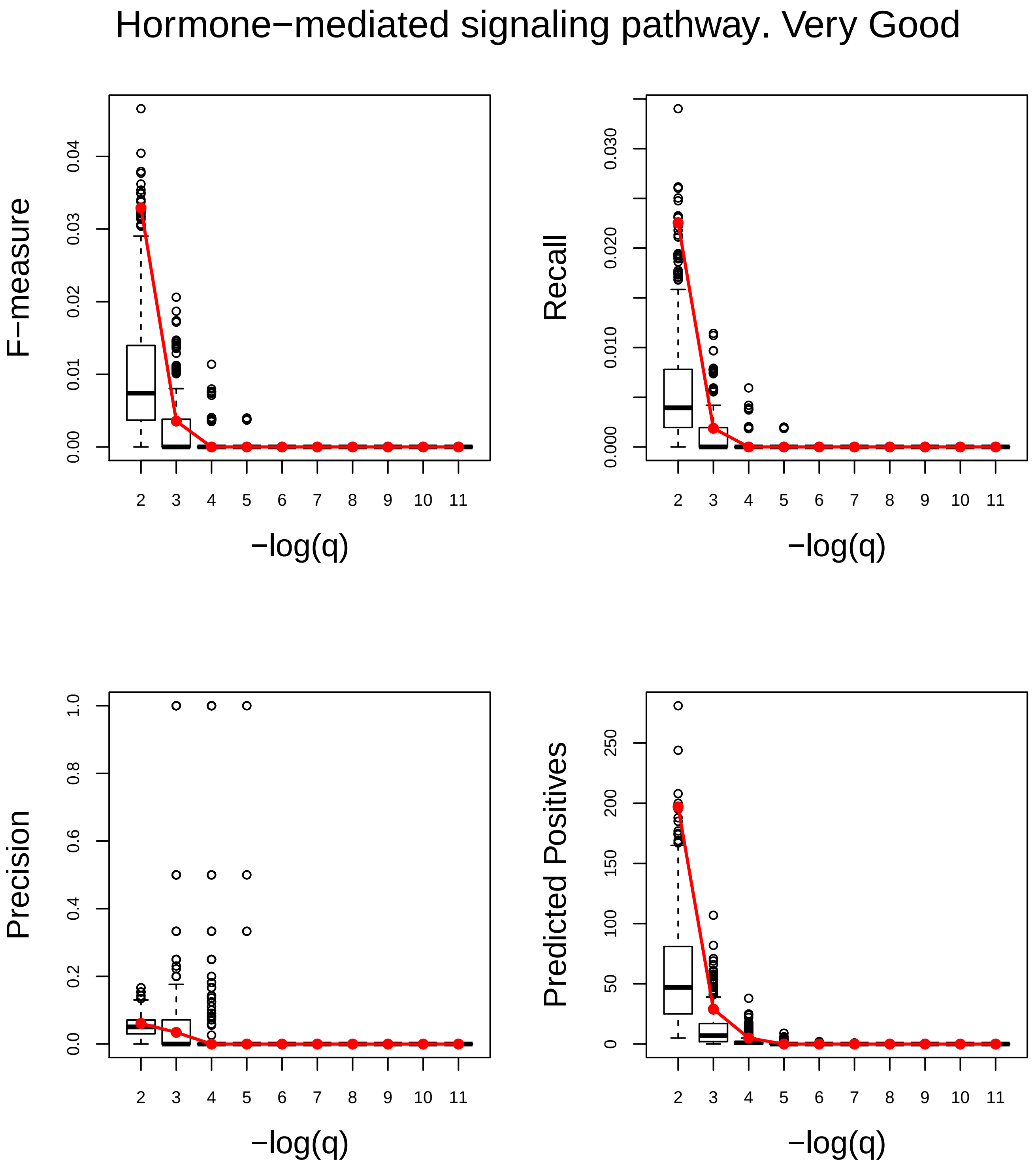
Hormone−mediated signaling pathway. Very Good
F−measure
0.00
0.01
0.02
0.03
0.04
Recall
0.000
0.010
0.020
0.030
Precision
0.0
0.2
0.4
0.6
0.8
1.0
Predicted Positives
0
50
100
150
200
250
2 3 4 5 6 7 8 9 10 11
−log(q)

### Hyperosmotic response. Very Poor

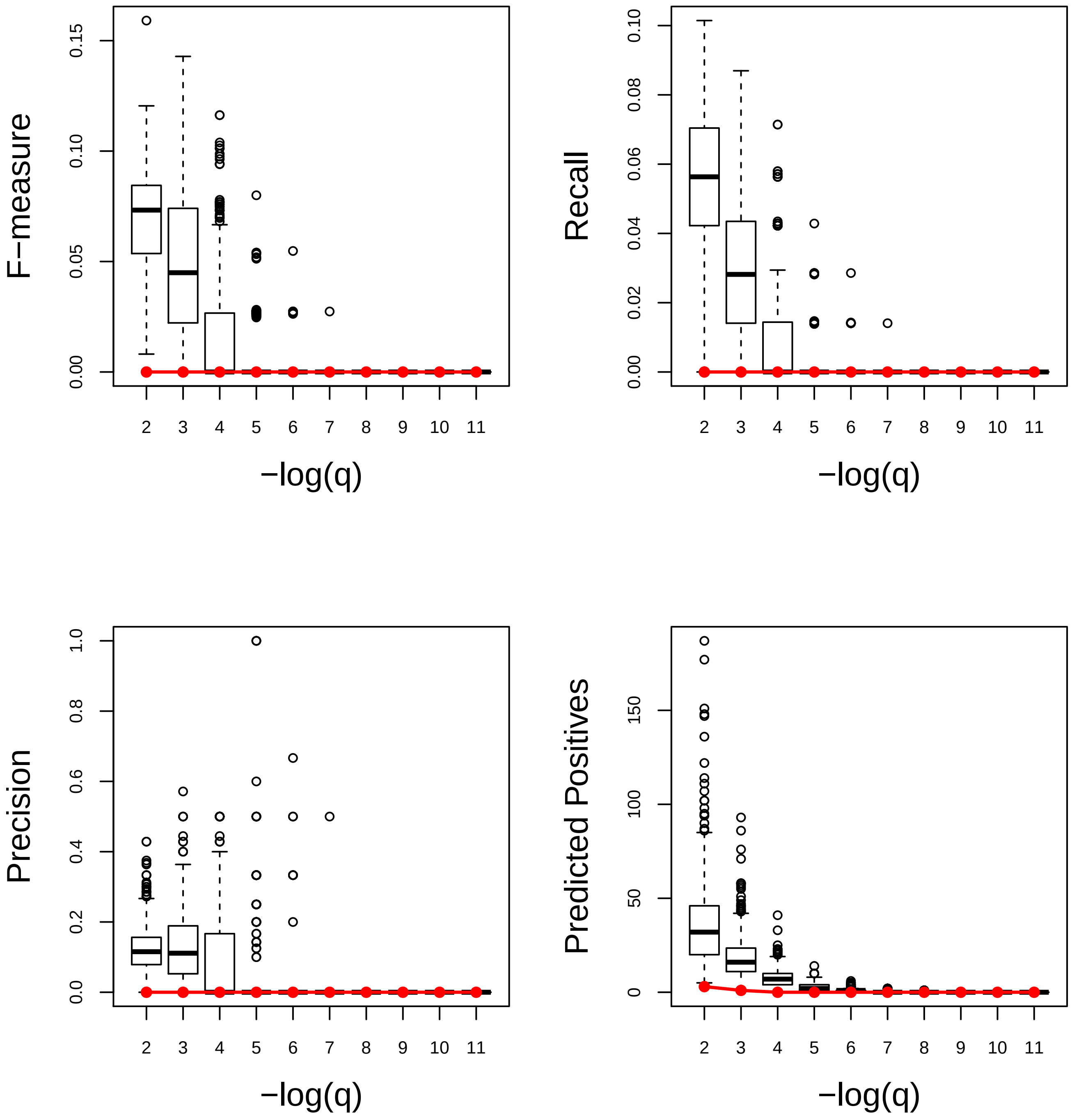

### Intracellular receptor signaling pathway. Very Good

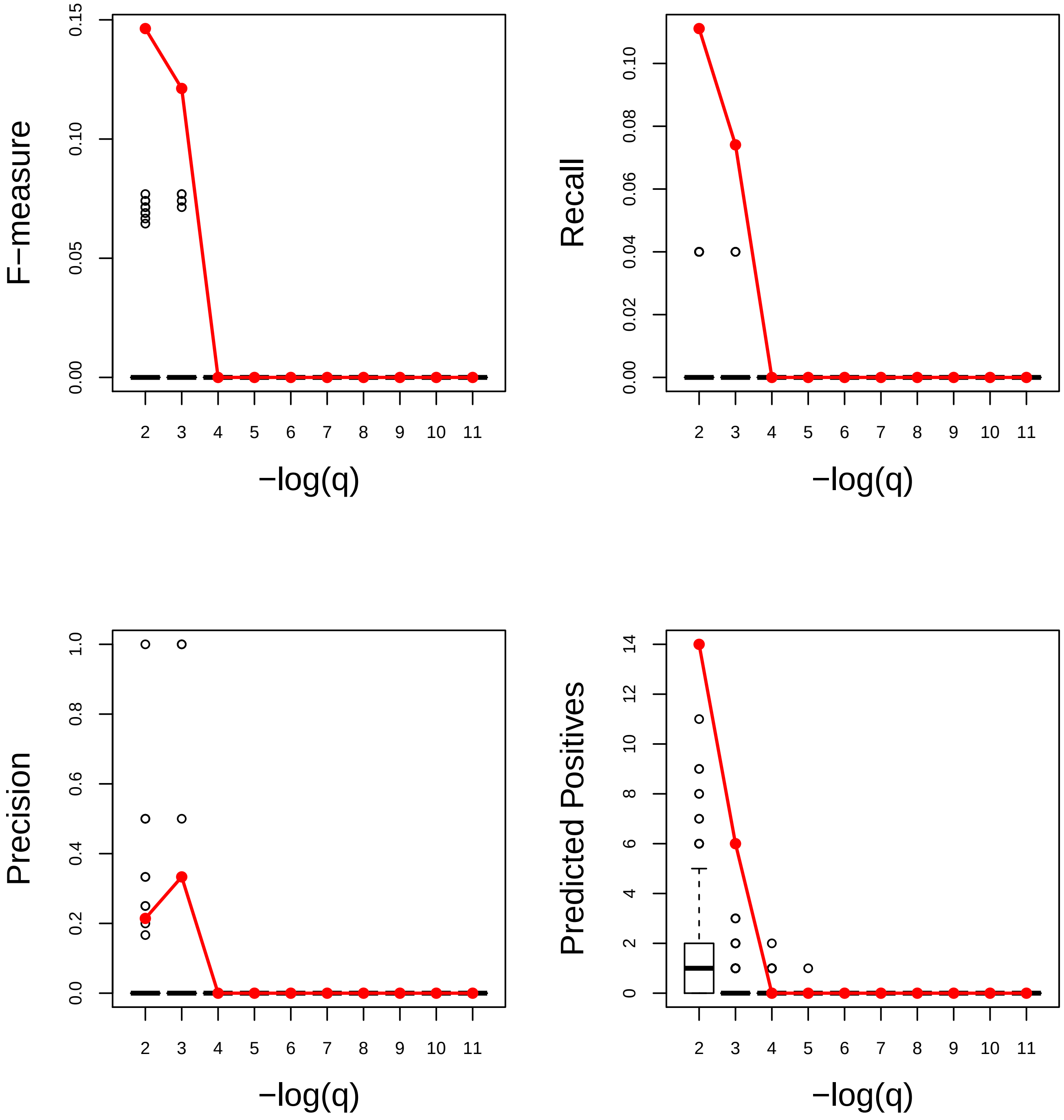

Intracellular signal transduction. Very Good

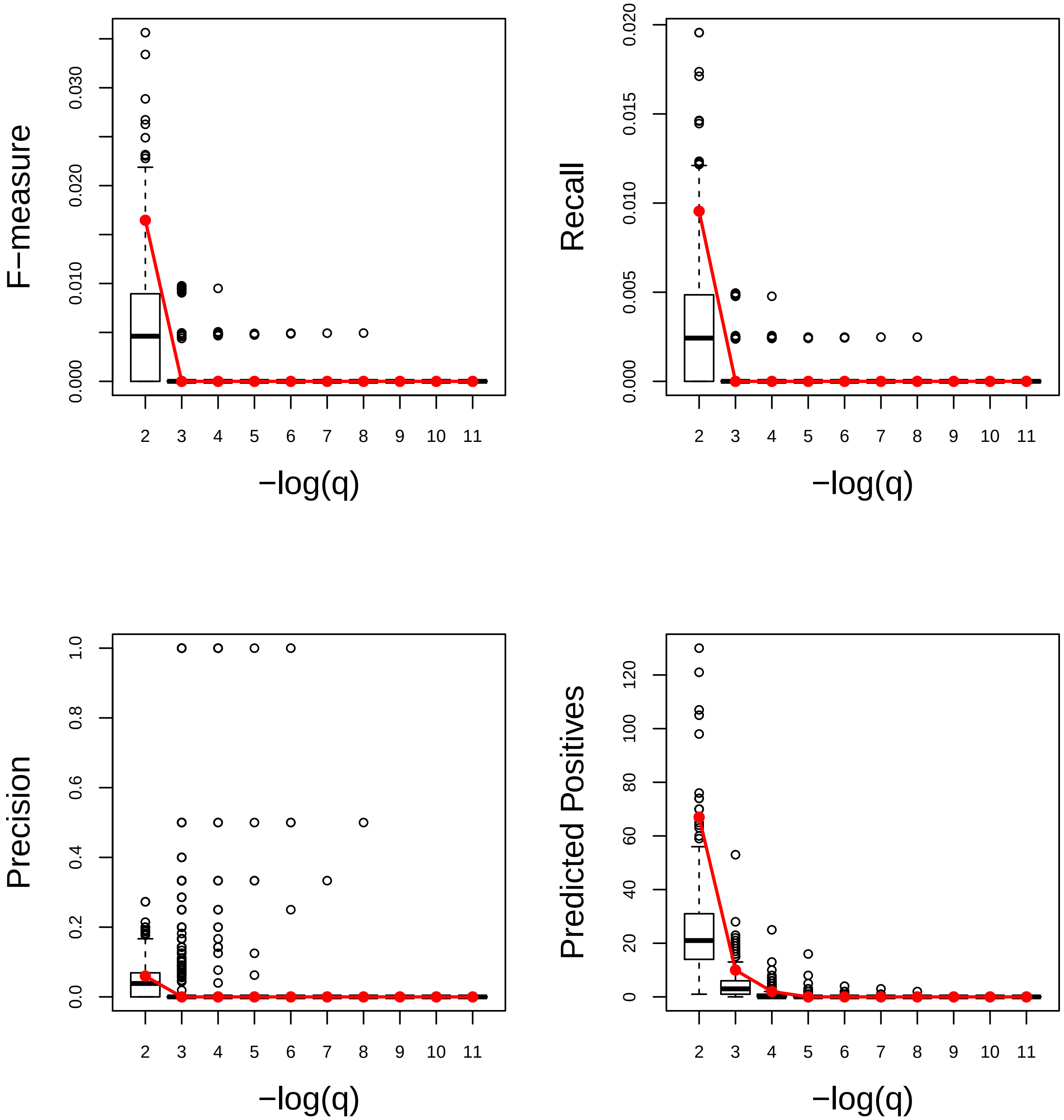

### Leaf development. Very Good

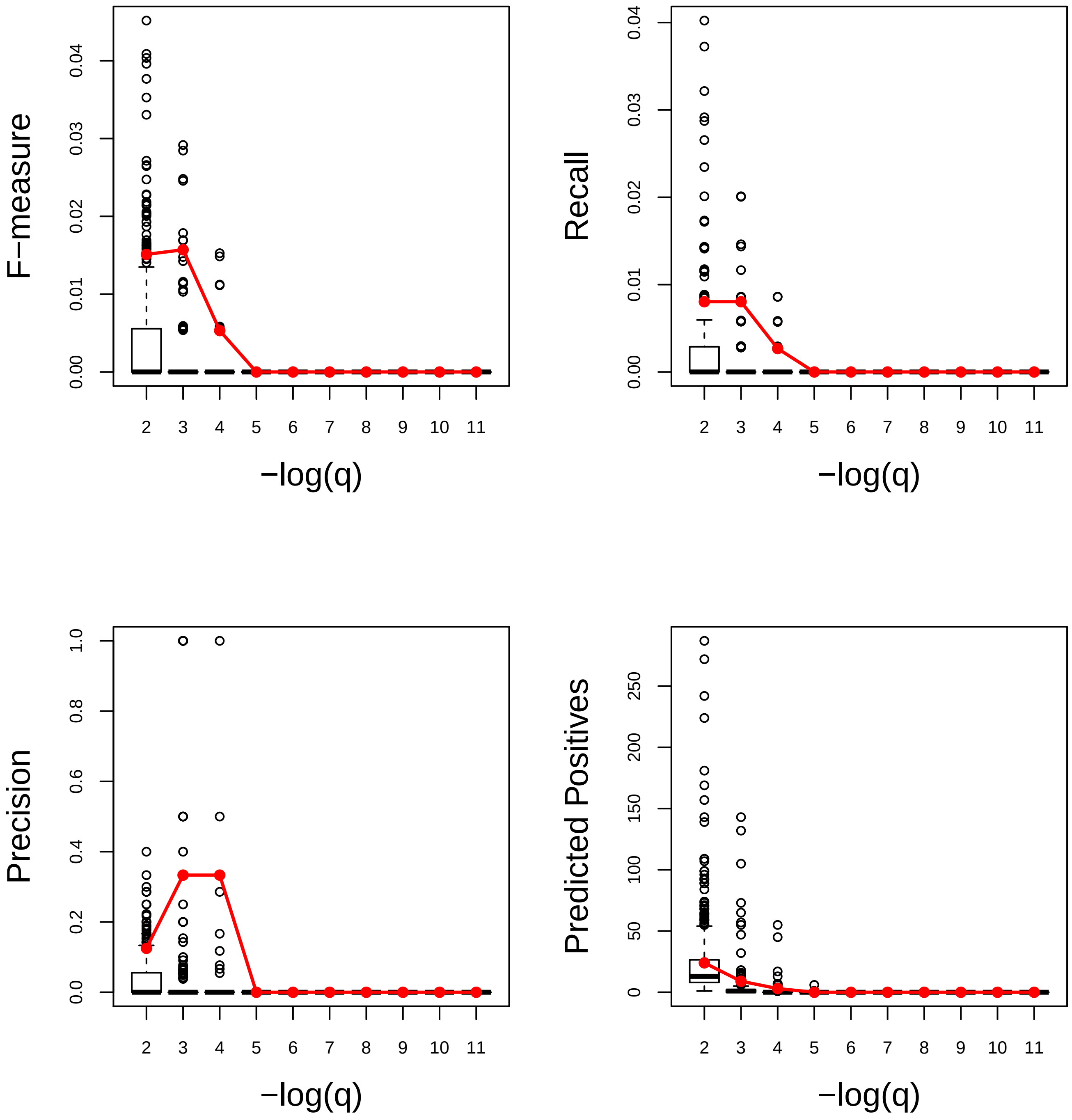

Lipid metabolic process. Very Good

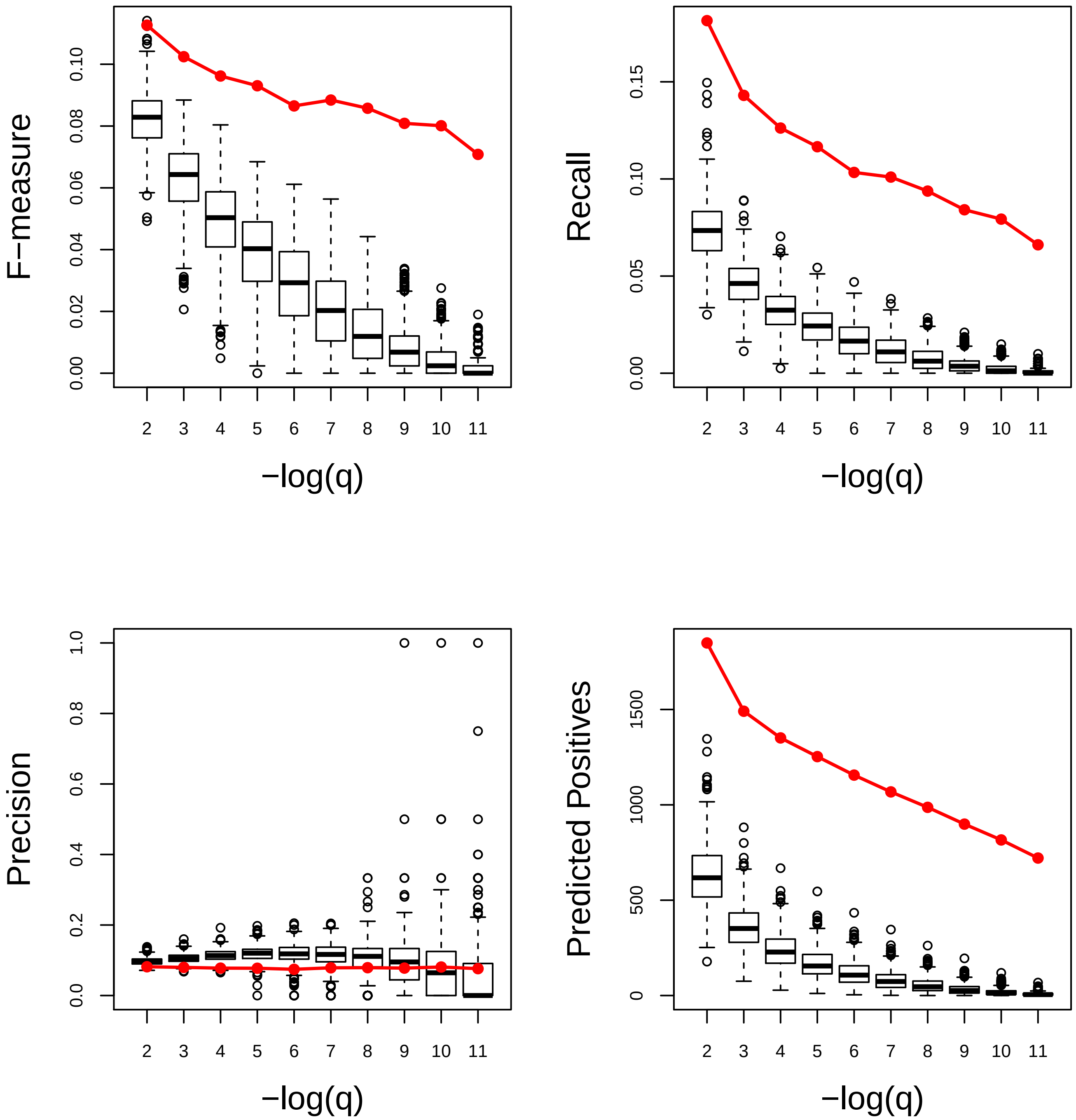

### Nonphotochemical quenching. Very Good

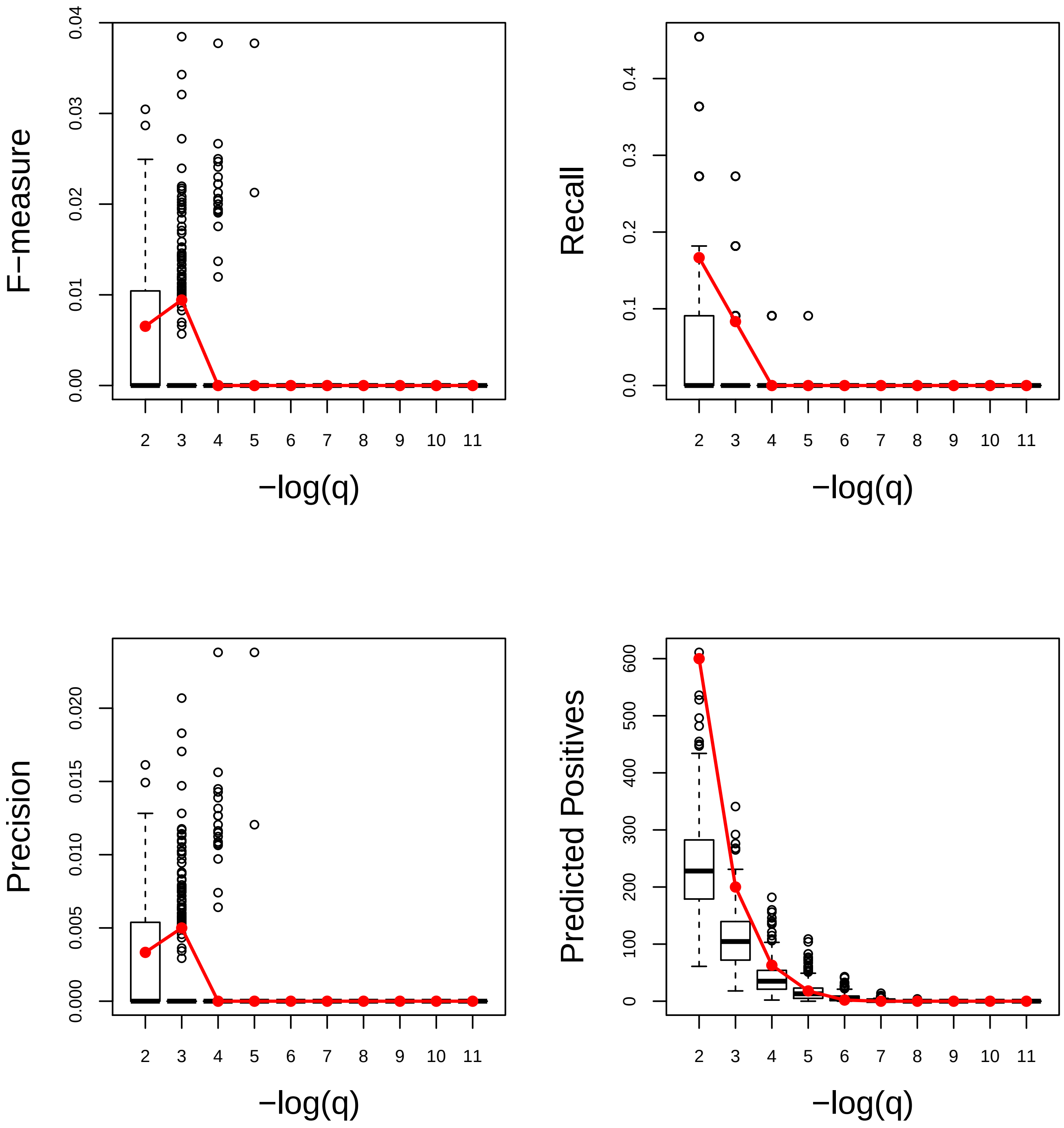

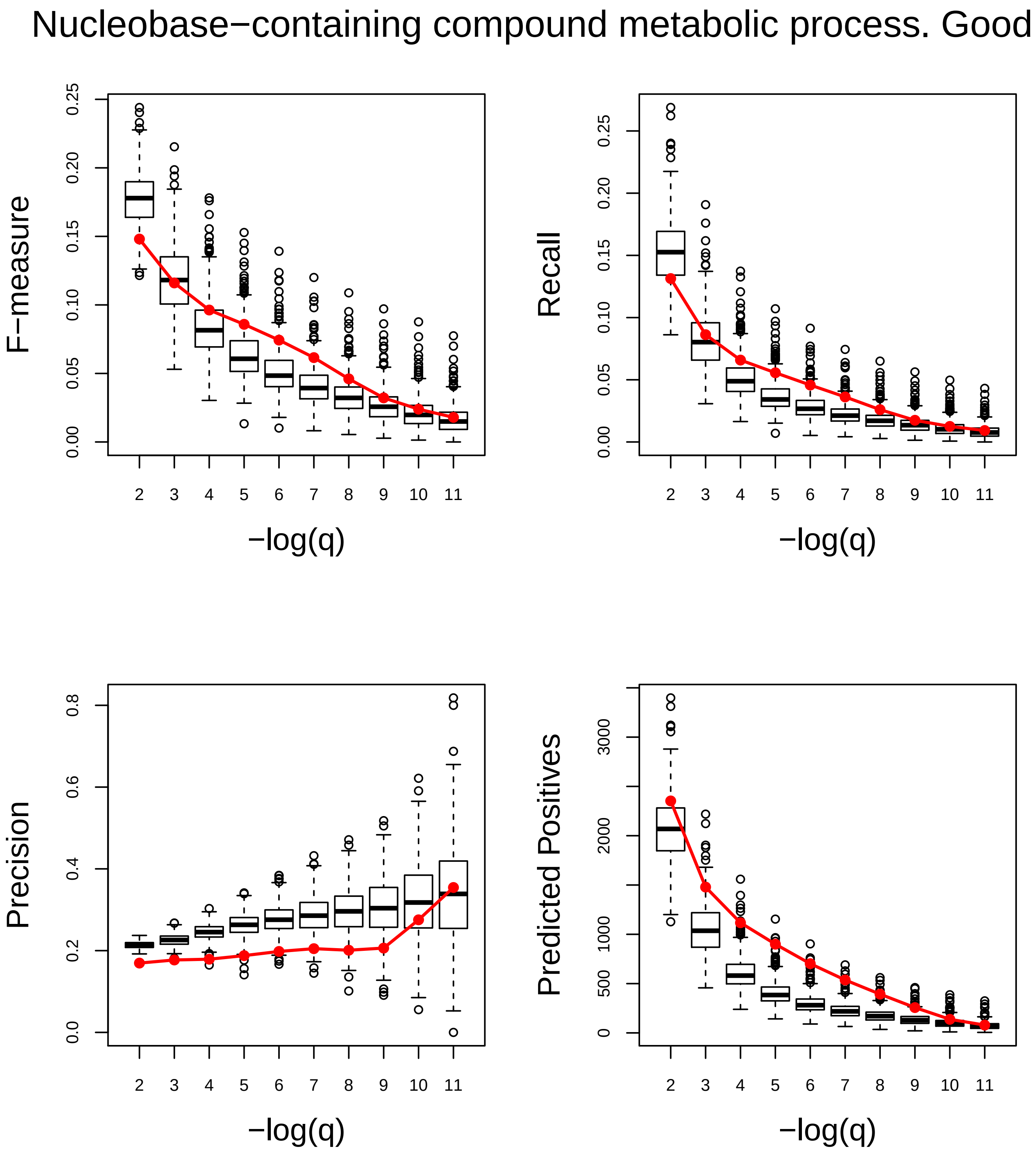
Nucleobase−containing compound metabolic process. Good
F−measure
Recall
Precision
Predicted Positives
−log(q)
−log(q)
−log(q)
−log(q)

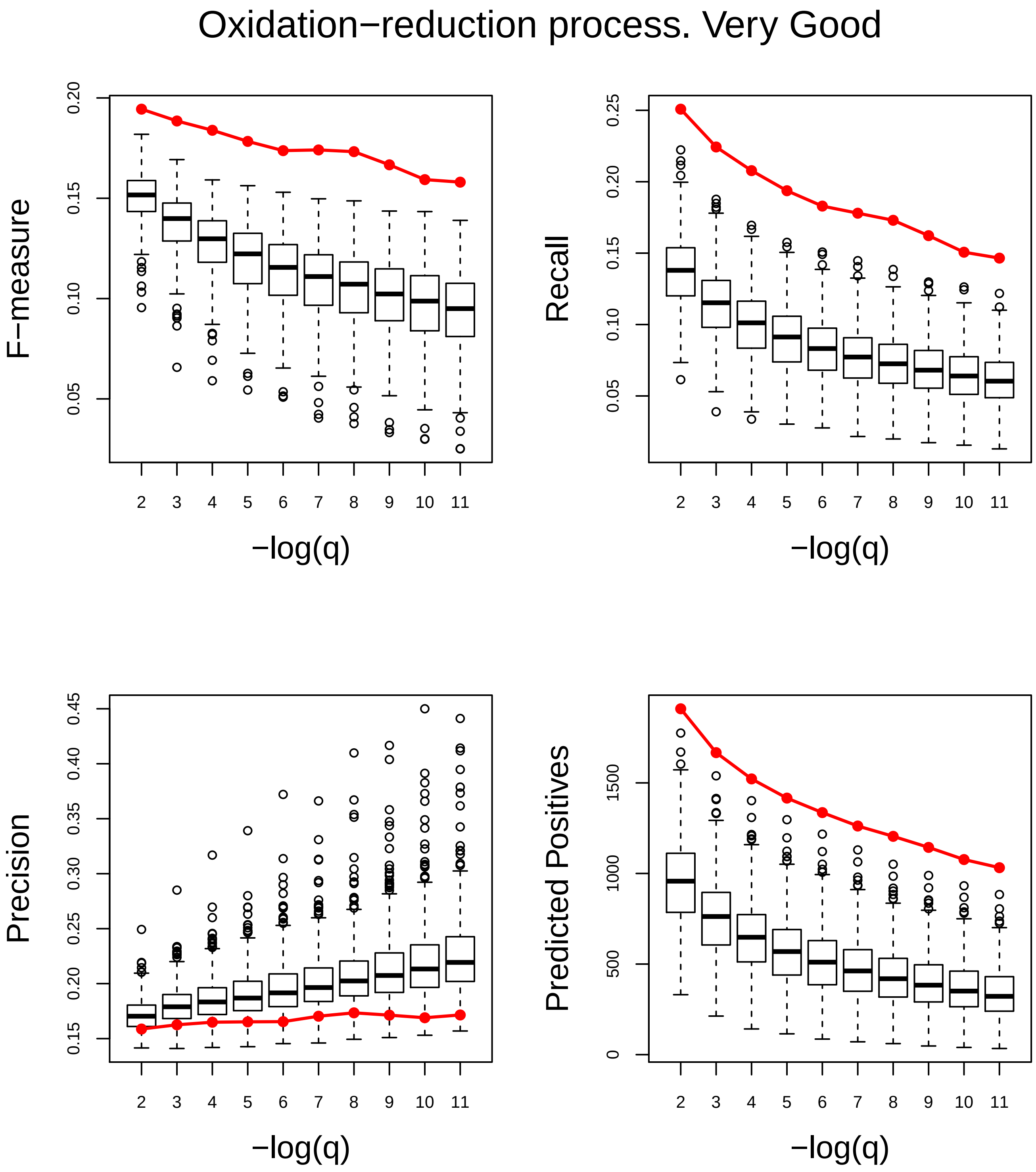

Oxidation−reduction process. Very Good
F−measure
Recall
Precision
Predicted Positives
−log(q)
−log(q)
−log(q)
−log(q)

### Peptide metabolic process. Very Poor

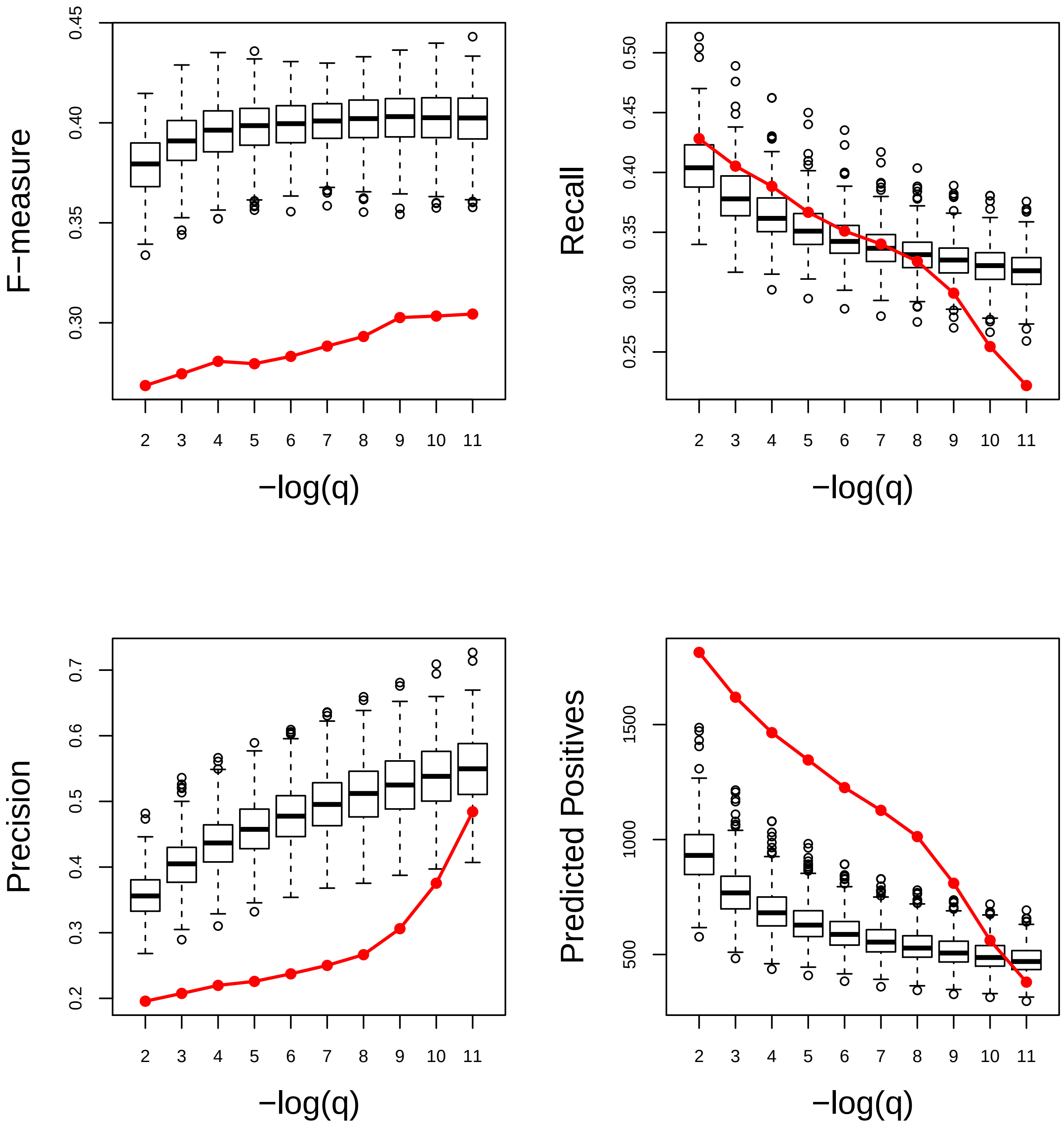

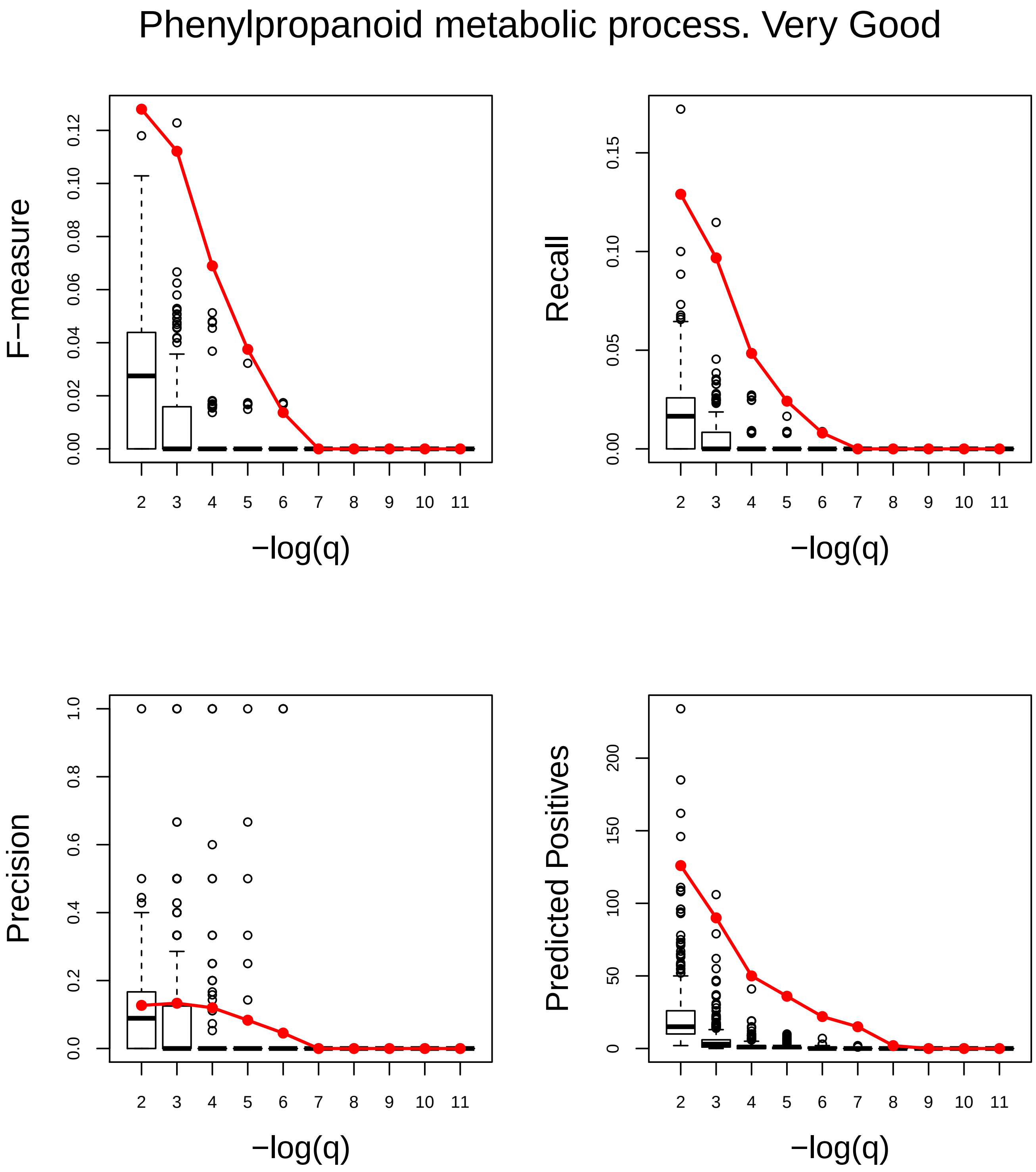
Phenylpropanoid metabolic process. Very Good
F−measure
0.00
0.02
0.04
0.06
0.08
0.10
0.12
Recall
0.00
0.05
0.10
0.15
Precision
0.0
0.2
0.4
0.6
0.8
1.0
Predicted Positives
0
50
100
150
200
2 3 4 5 6 7 8 9 10 11
−log(q)

### Phosphorelay signal transduction system. Very Good

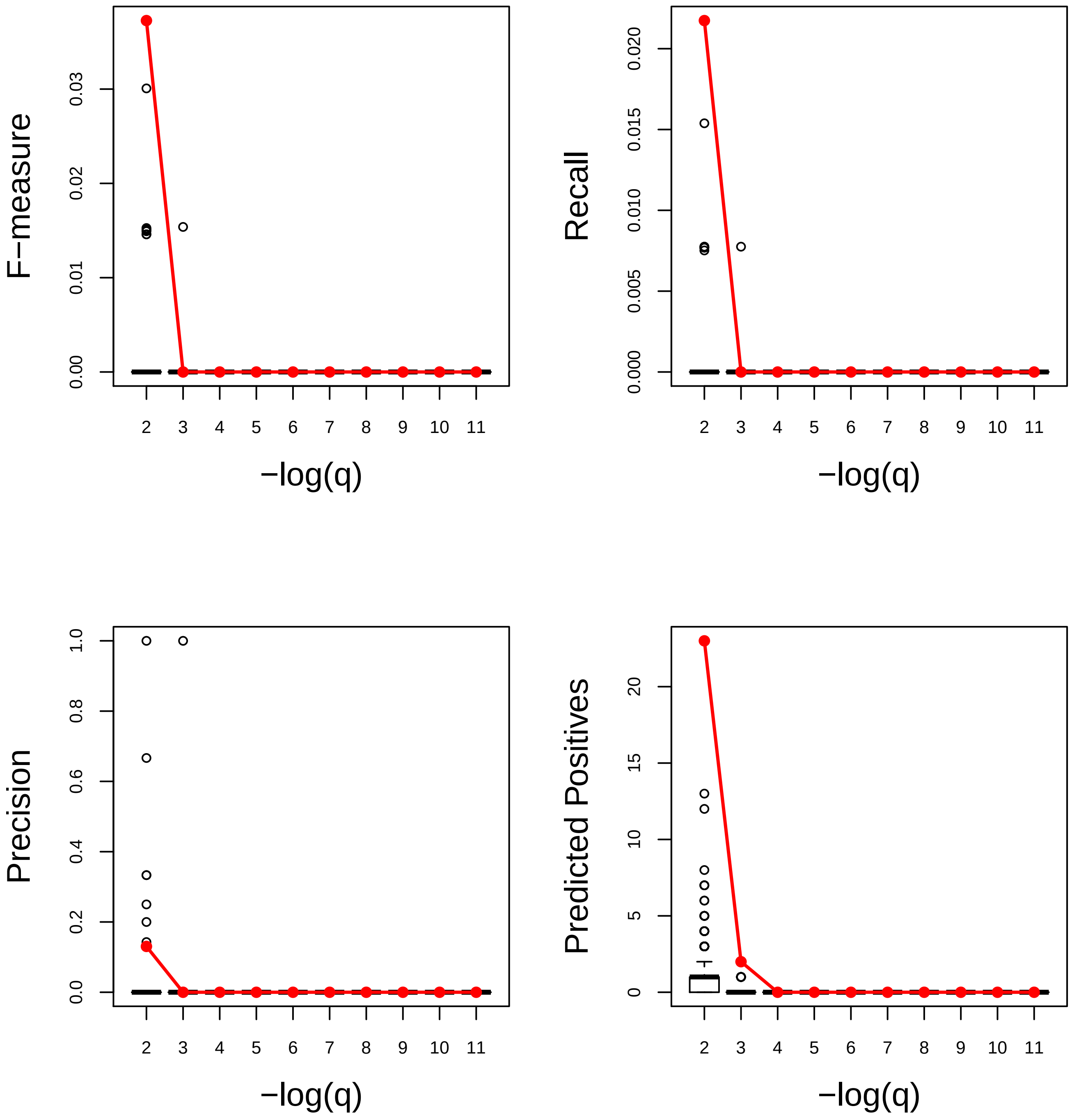

#### Photoperiodism. Very Poor

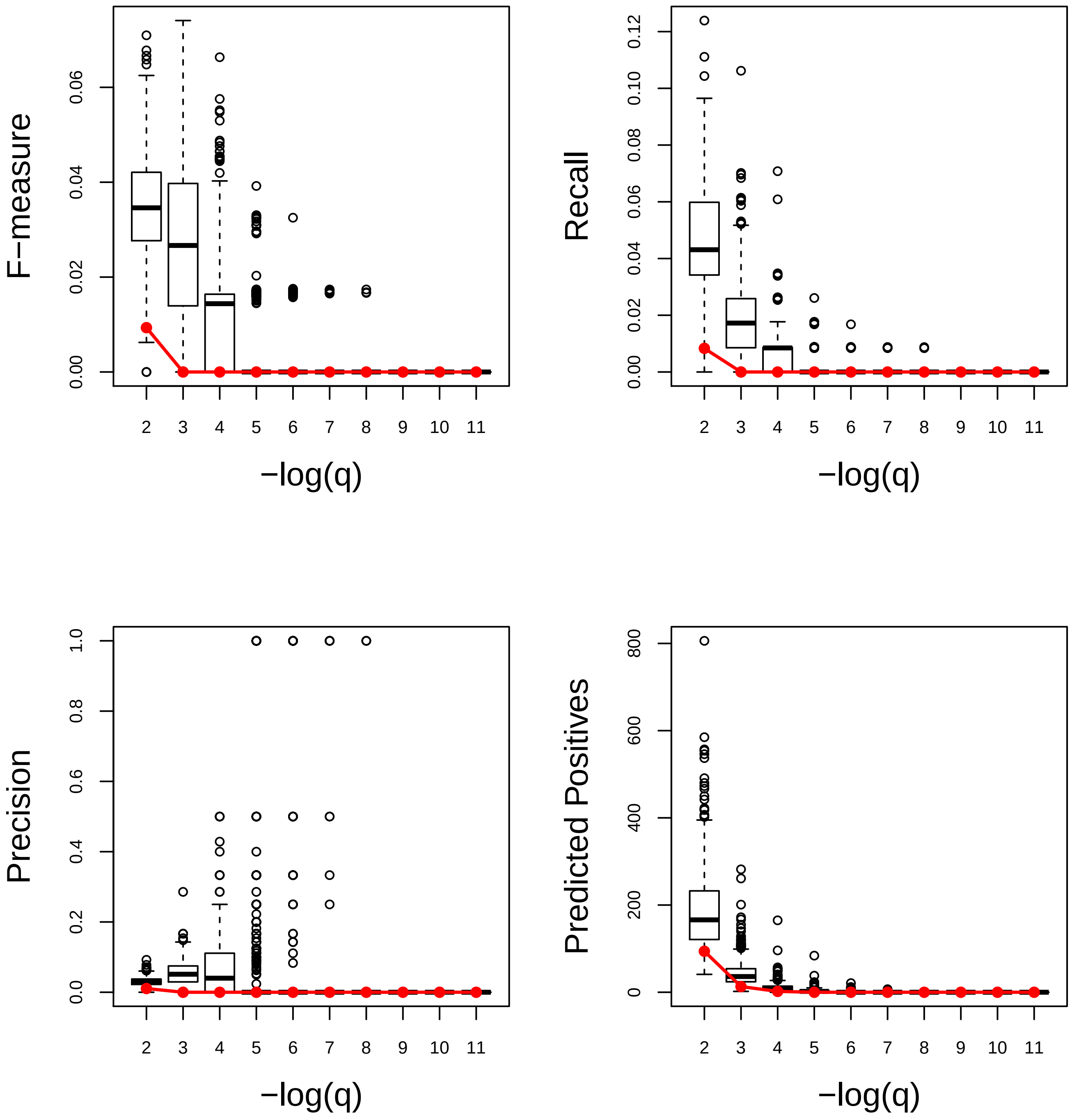

### Photosynthesis, light reaction. Very Poor

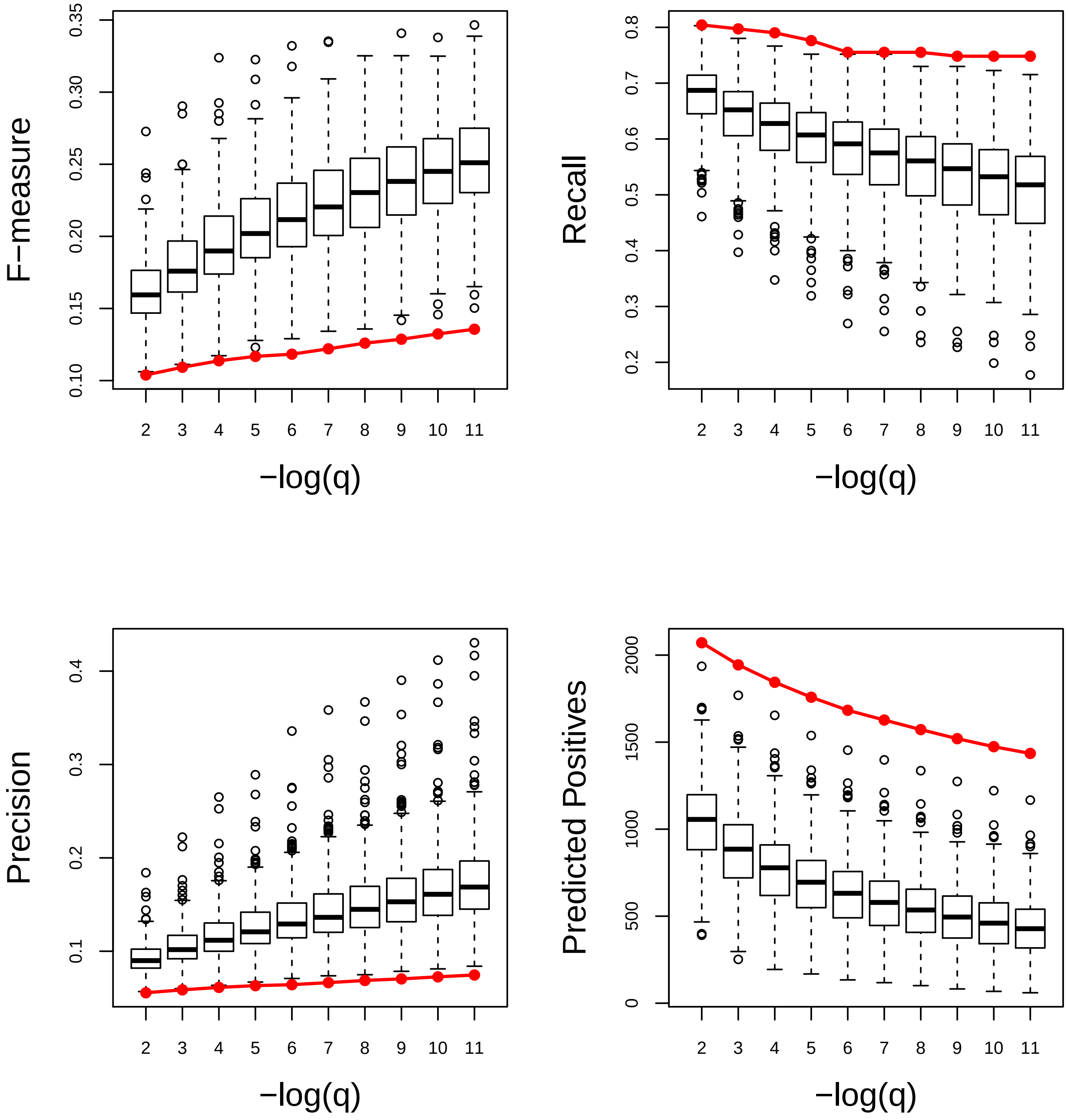

### Photosynthesis. Very Poor

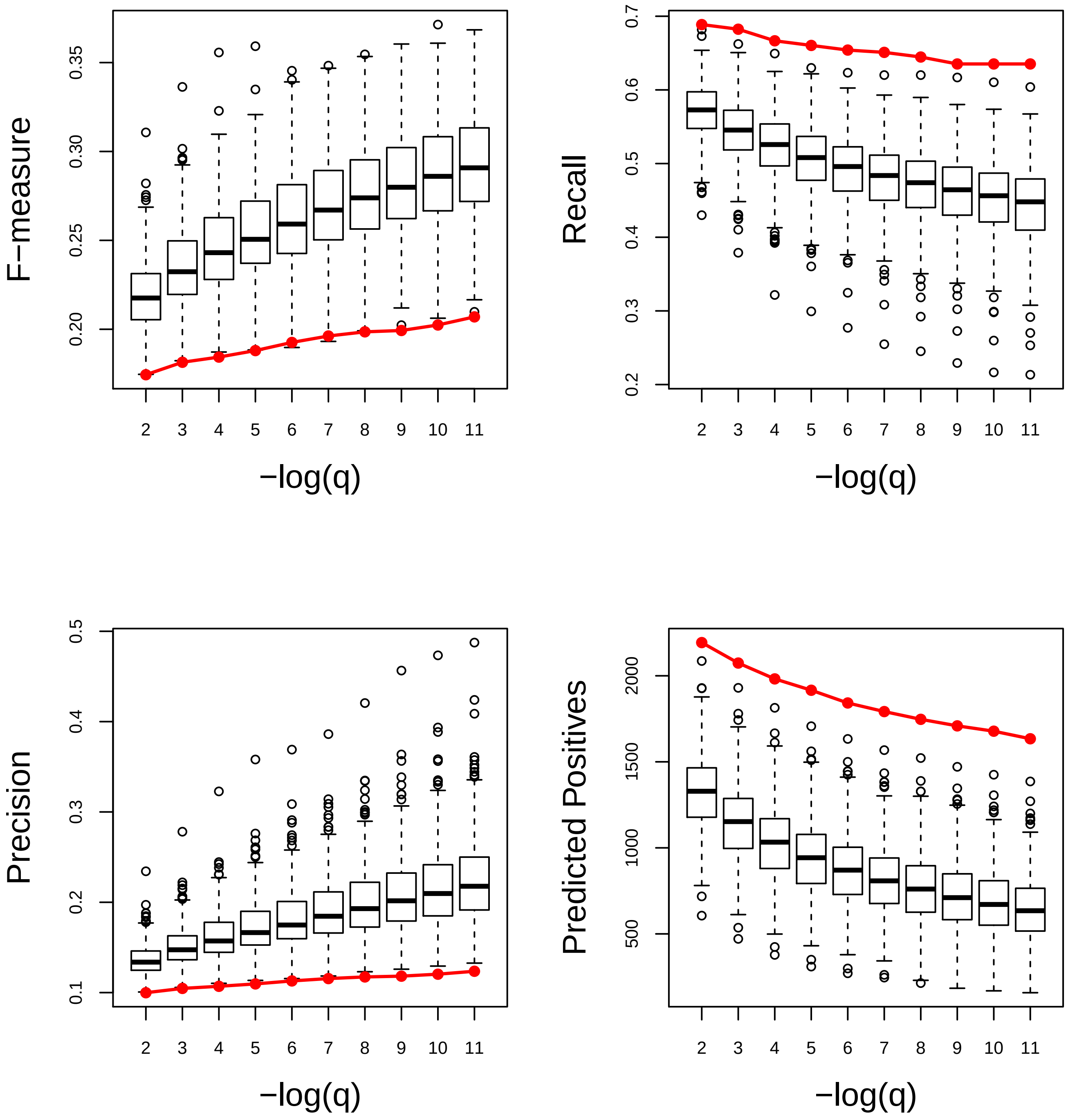

### Photosynthetic electron transport chain. Very Poor

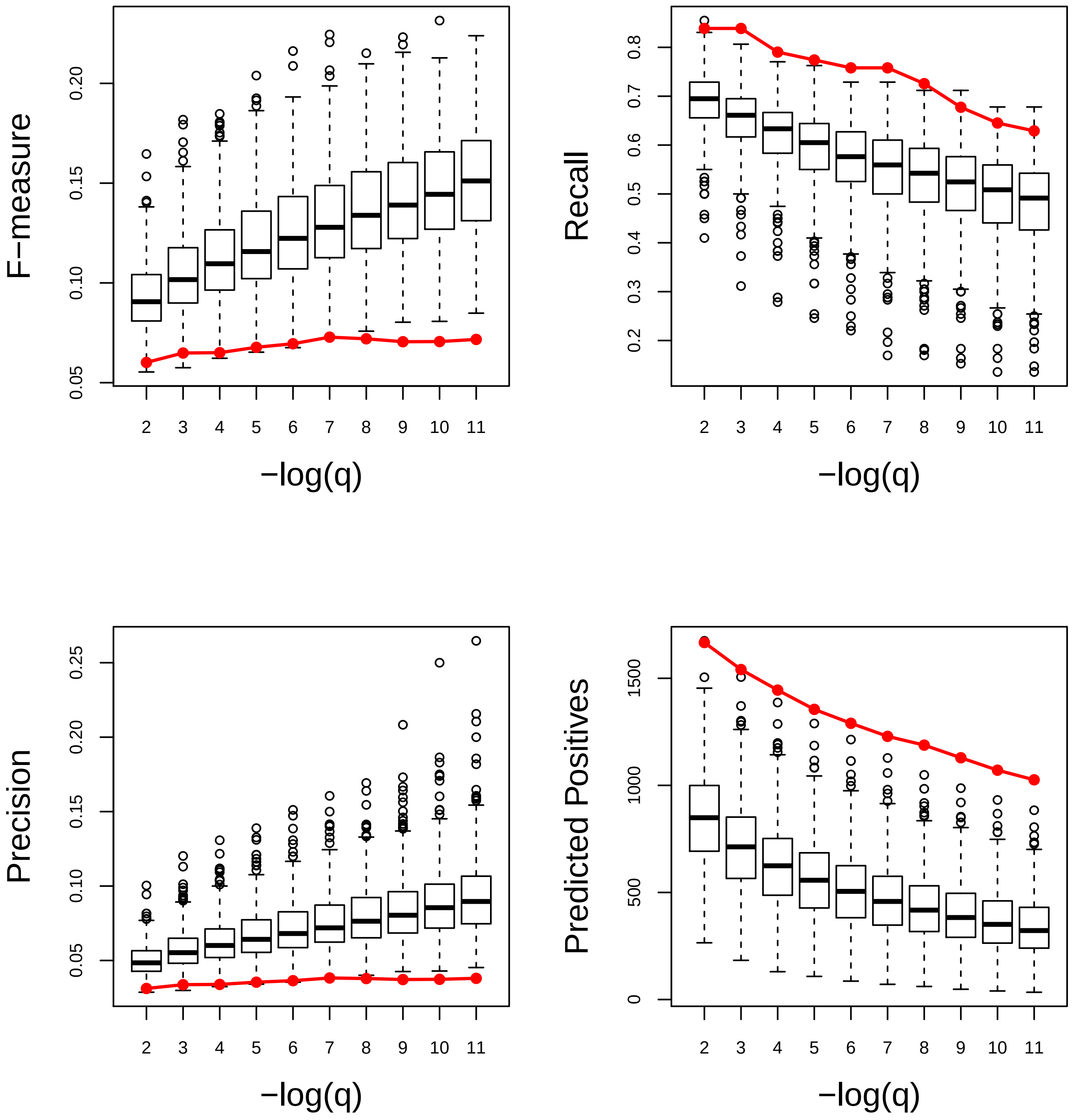

### Phototransduction. Good

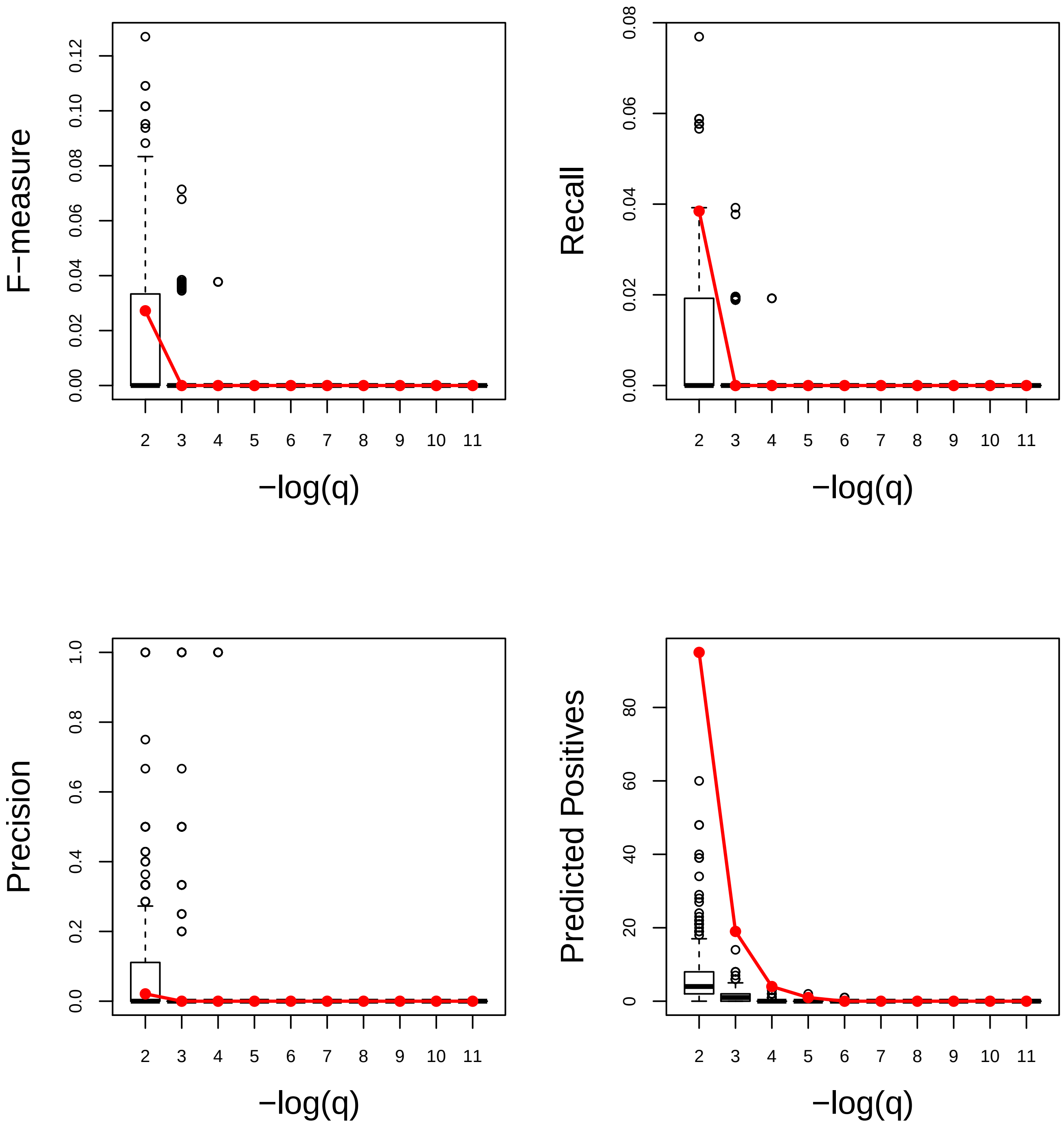

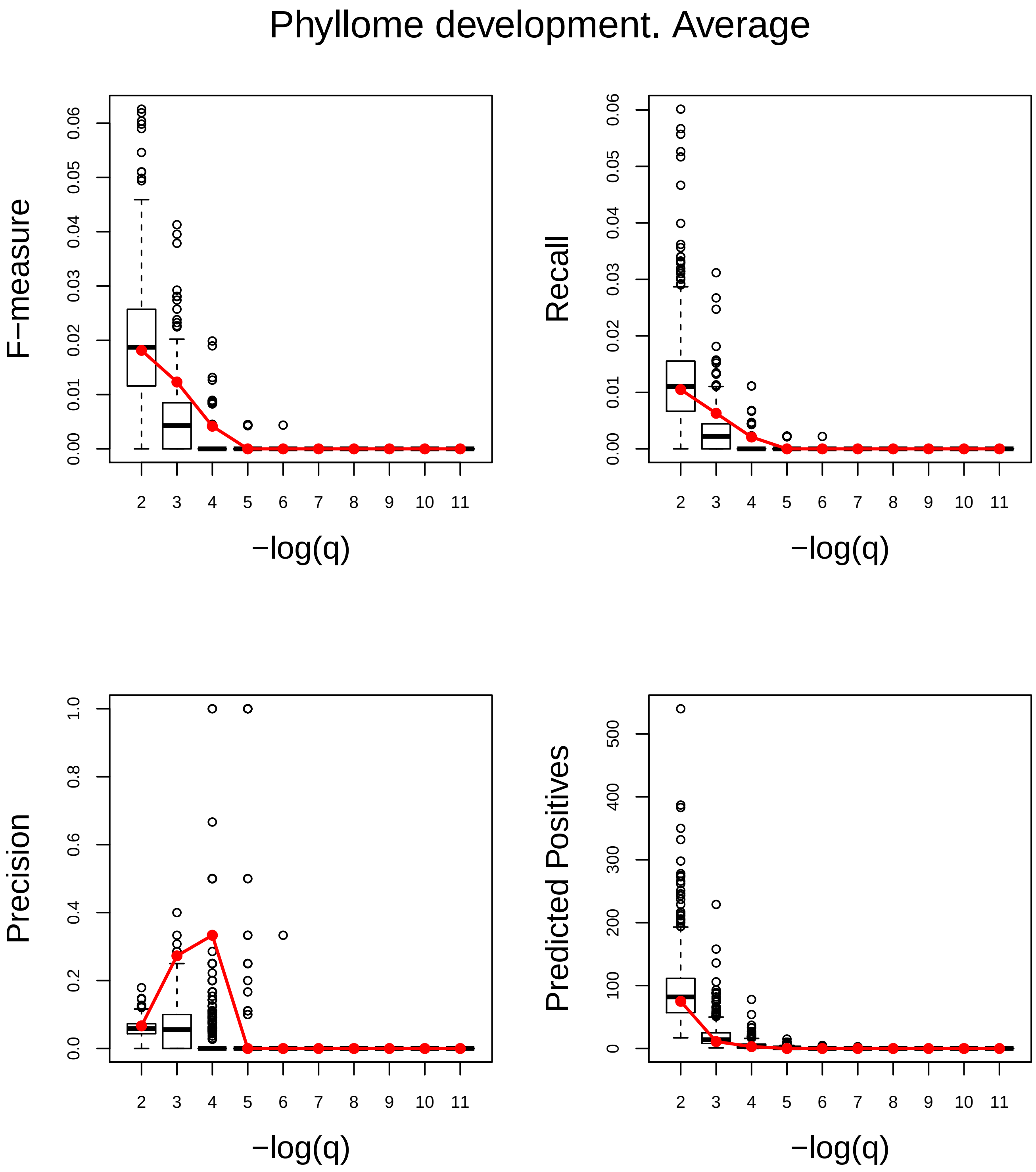
Phyllome development. Average
F−measure
0.00
0.01
0.02
0.03
0.04
0.05
0.06
Recall
0.00
0.01
0.02
0.03
0.04
0.05
0.06
Precision
0.0
0.2
0.4
0.6
0.8
1.0
Predicted Positives
0
100
200
300
400
500
2 3 4 5 6 7 8 9 10 11
−log(q)

### Plant−type cell wall organization or biogenesis. Very Poor

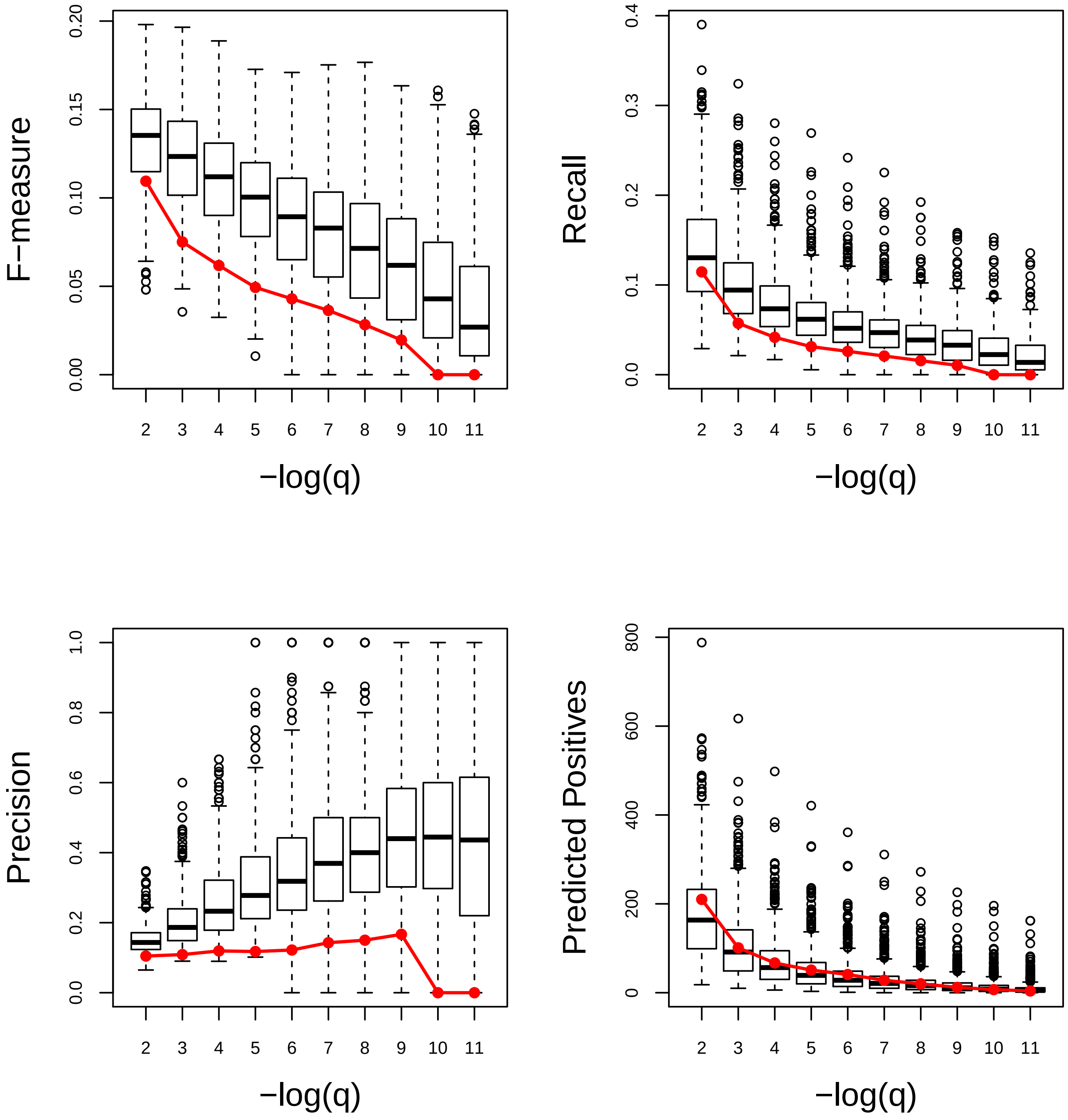

### Plant−type secondary cell wall biogenesis. Average

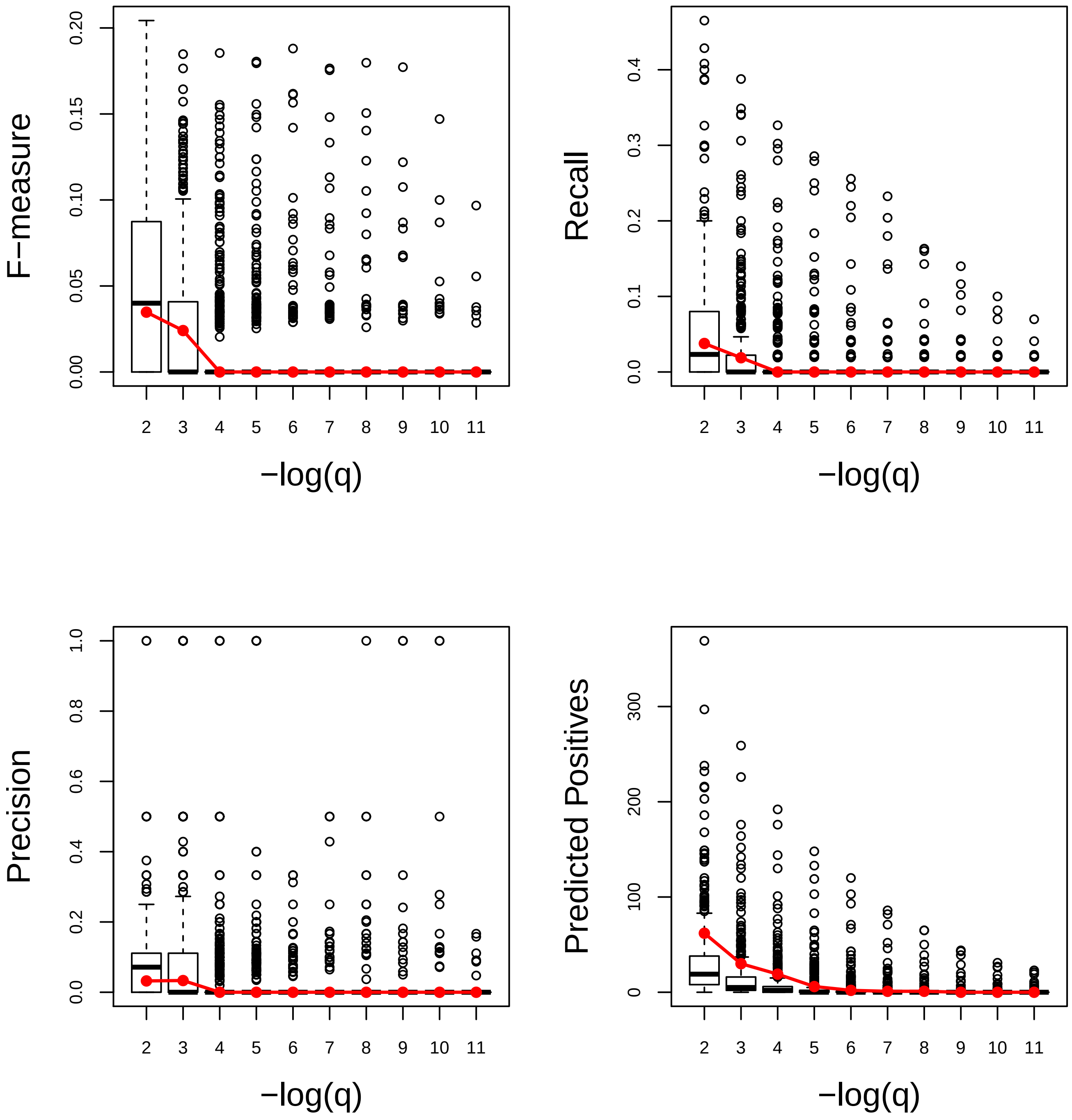

### Plastid organization. Good

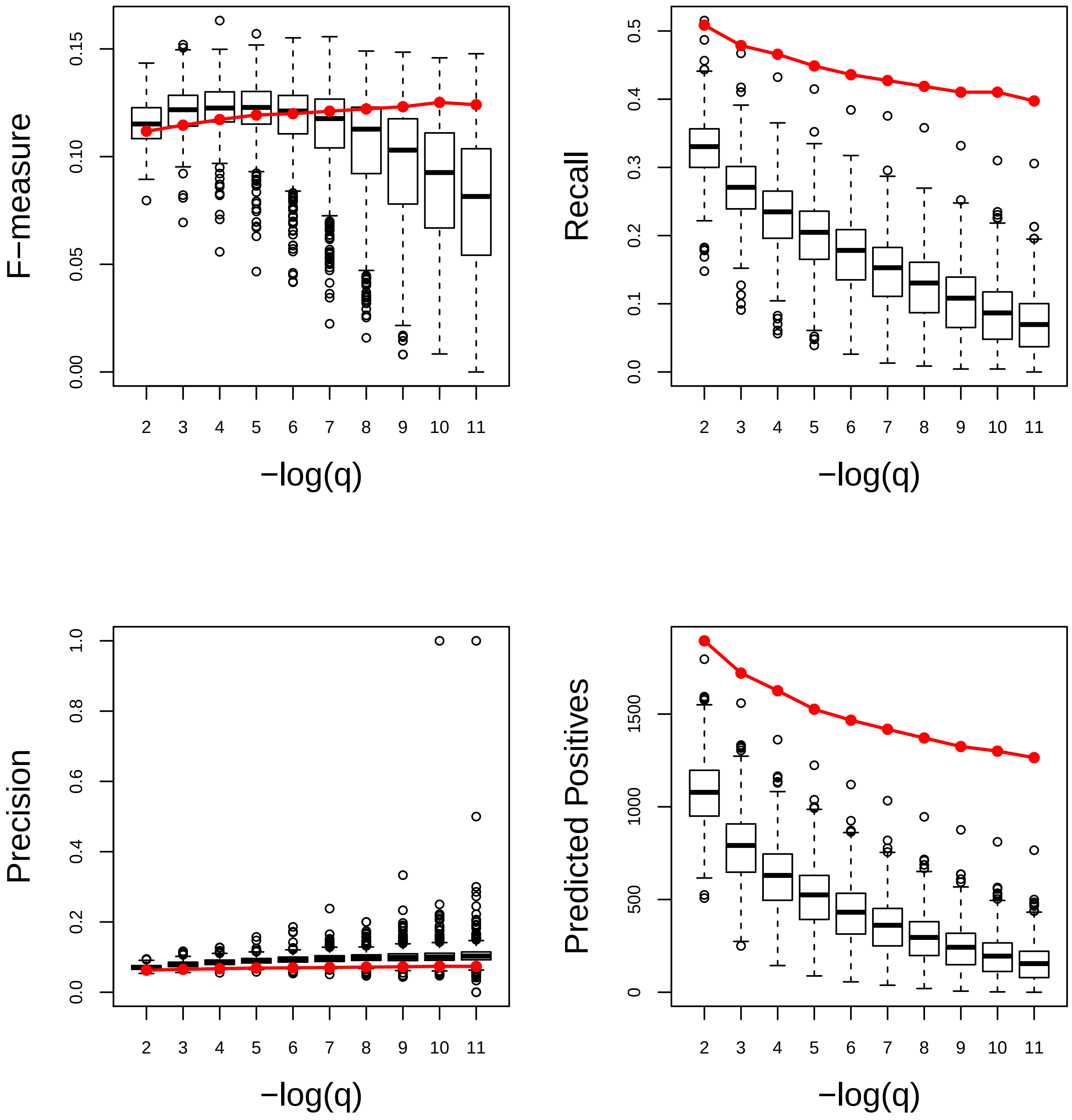

### Pollination. Poor

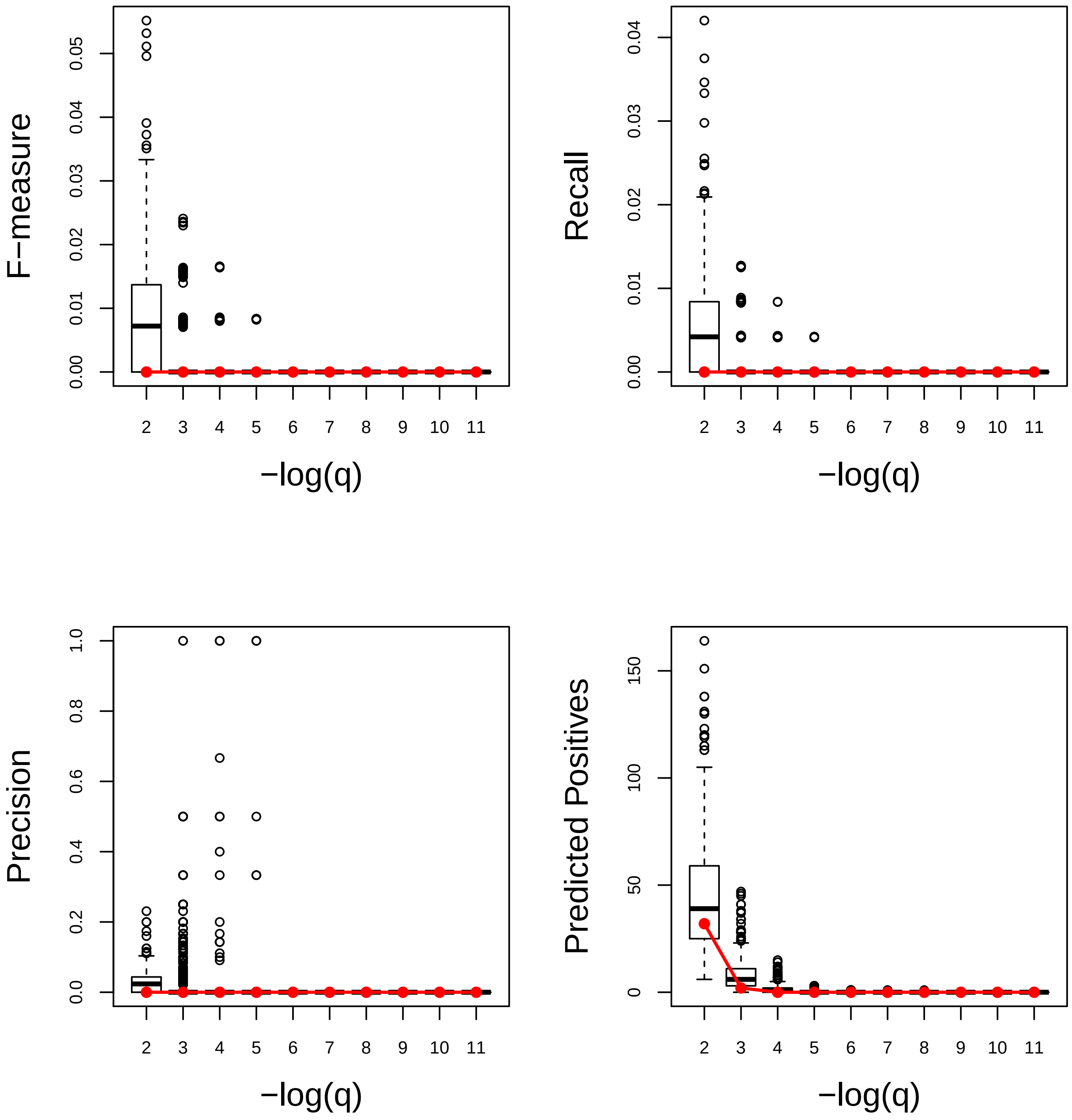

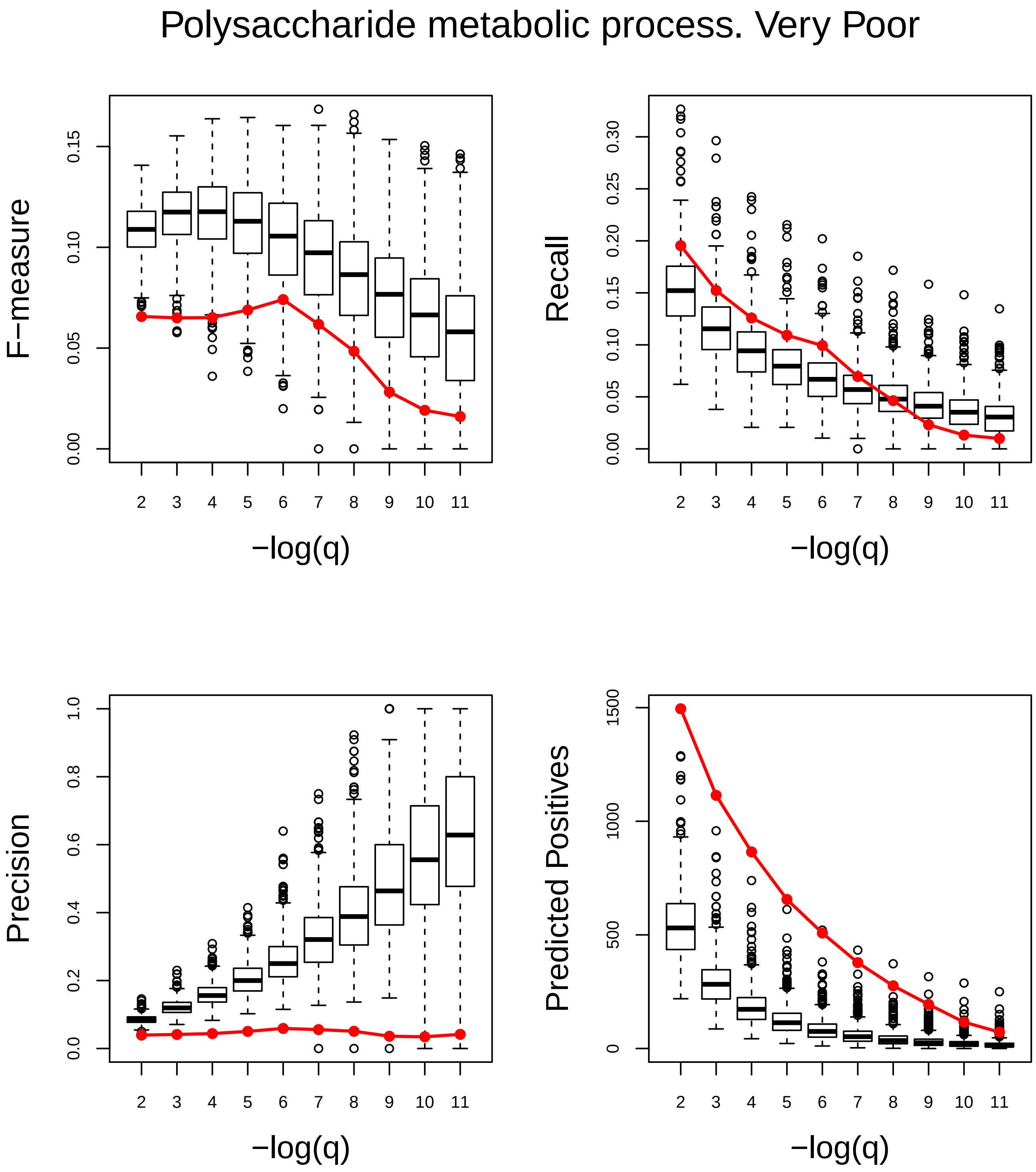
Polysaccharide metabolic process. Very Poor
F−measure
Recall
Precision
Predicted Positives
−log(q)
0.00
0.05
0.10
0.15
0.20
0.25
0.30
0.0
0.2
0.4
0.6
0.8
1.0
0
500
1000
1500

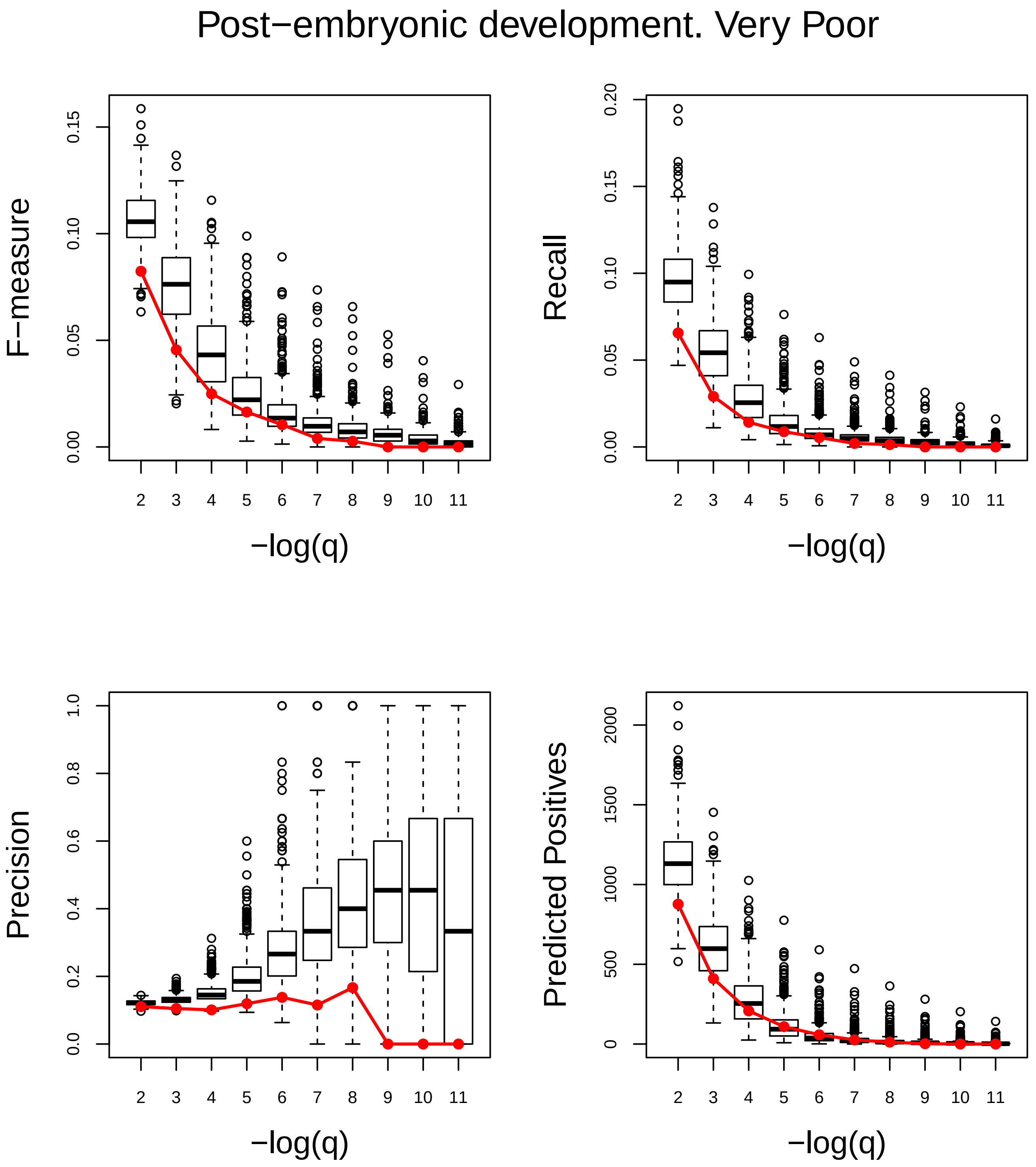
Post−embryonic development. Very Poor
F−measure
0.00
0.05
0.10
0.15
Recall
0.00
0.05
0.10
0.15
0.20
Precision
0.0
0.2
0.4
0.6
0.8
1.0
Predicted Positives
0
500
1000
1500
2000
2 3 4 5 6 7 8 9 10 11
−log(q)

### Protein metabolic process. Poor

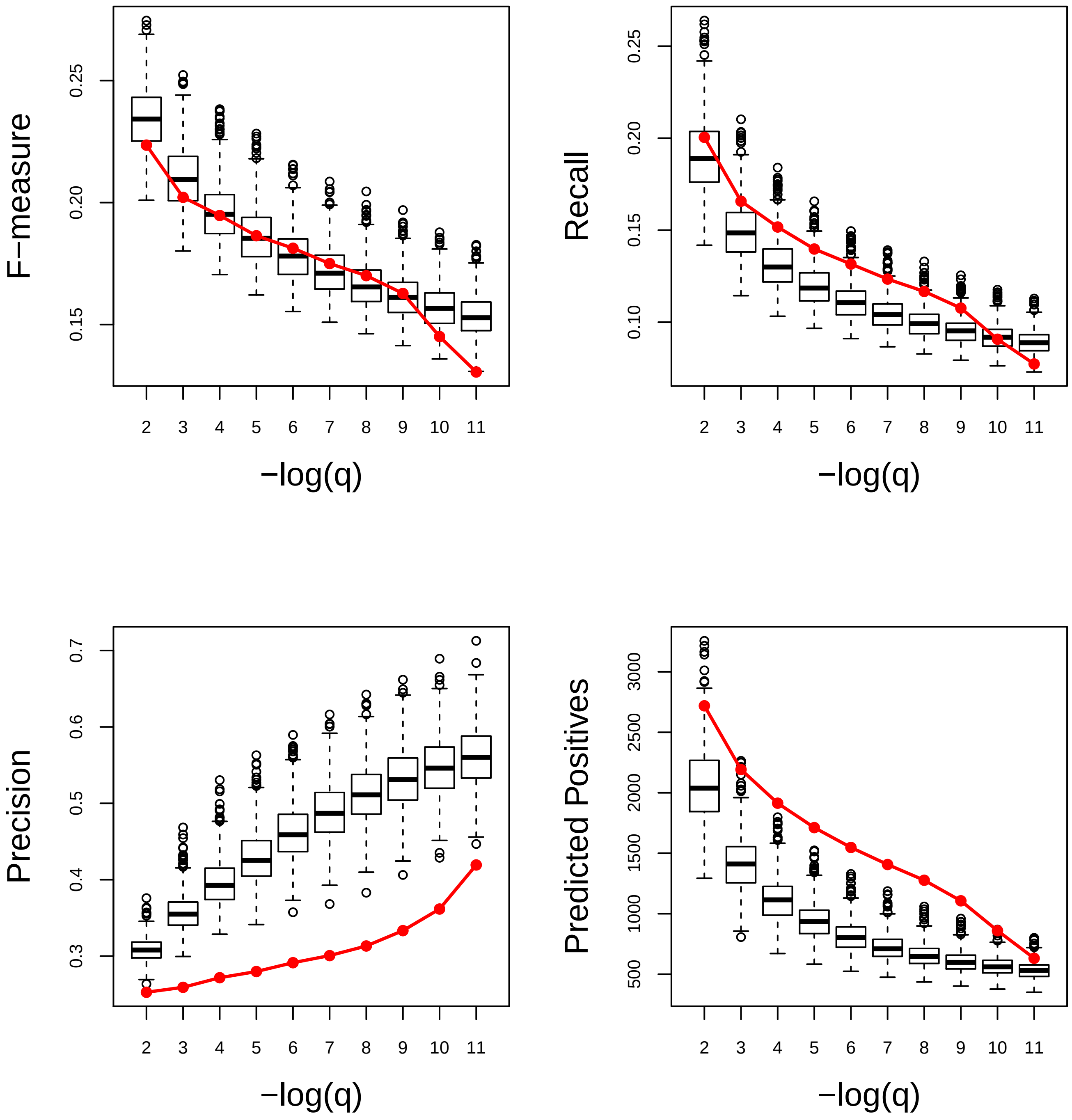

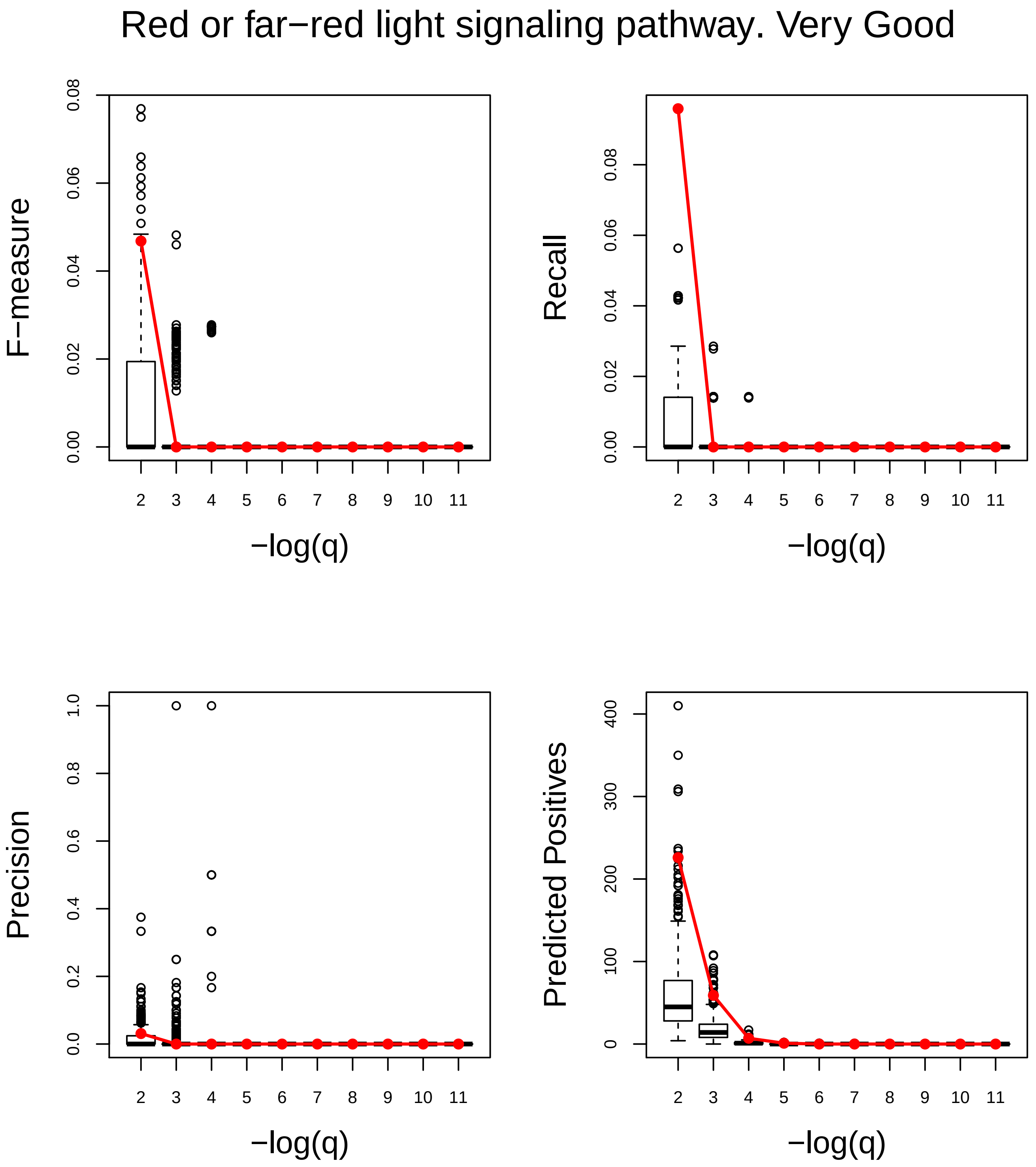
Red or far−red light signaling pathway. Very Good
F−measure
0.00
0.02
0.04
0.06
0.08
Recall
0.00
0.02
0.04
0.06
0.08
Precision
0.0
0.2
0.4
0.6
0.8
1.0
Predicted Positives
0
100
200
300
400
2 3 4 5 6 7 8 9 10 11
−log(q)

#### Regulation of defense response. Very Good

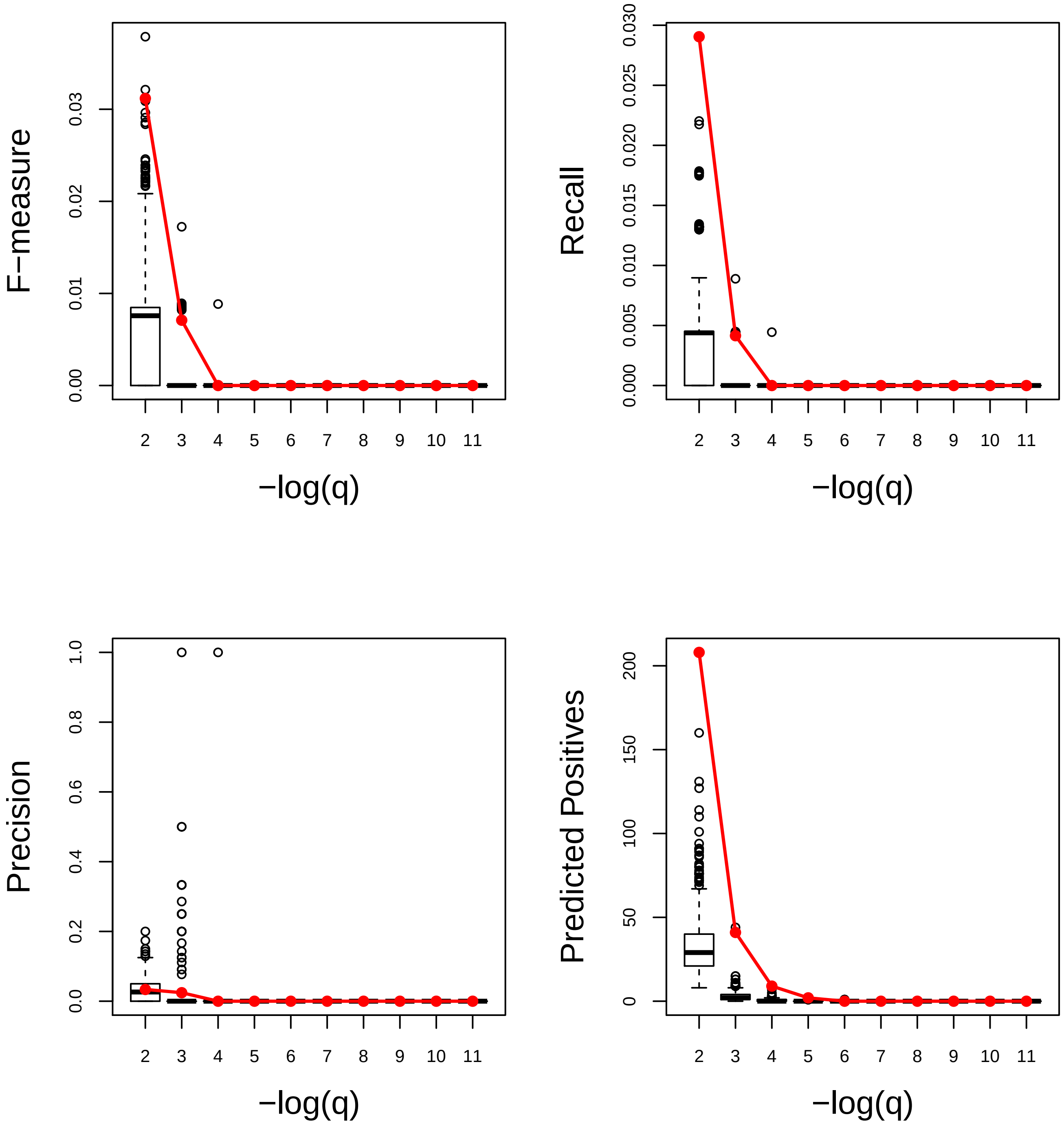

### Regulation of gene expression, epigenetic. Very Poor

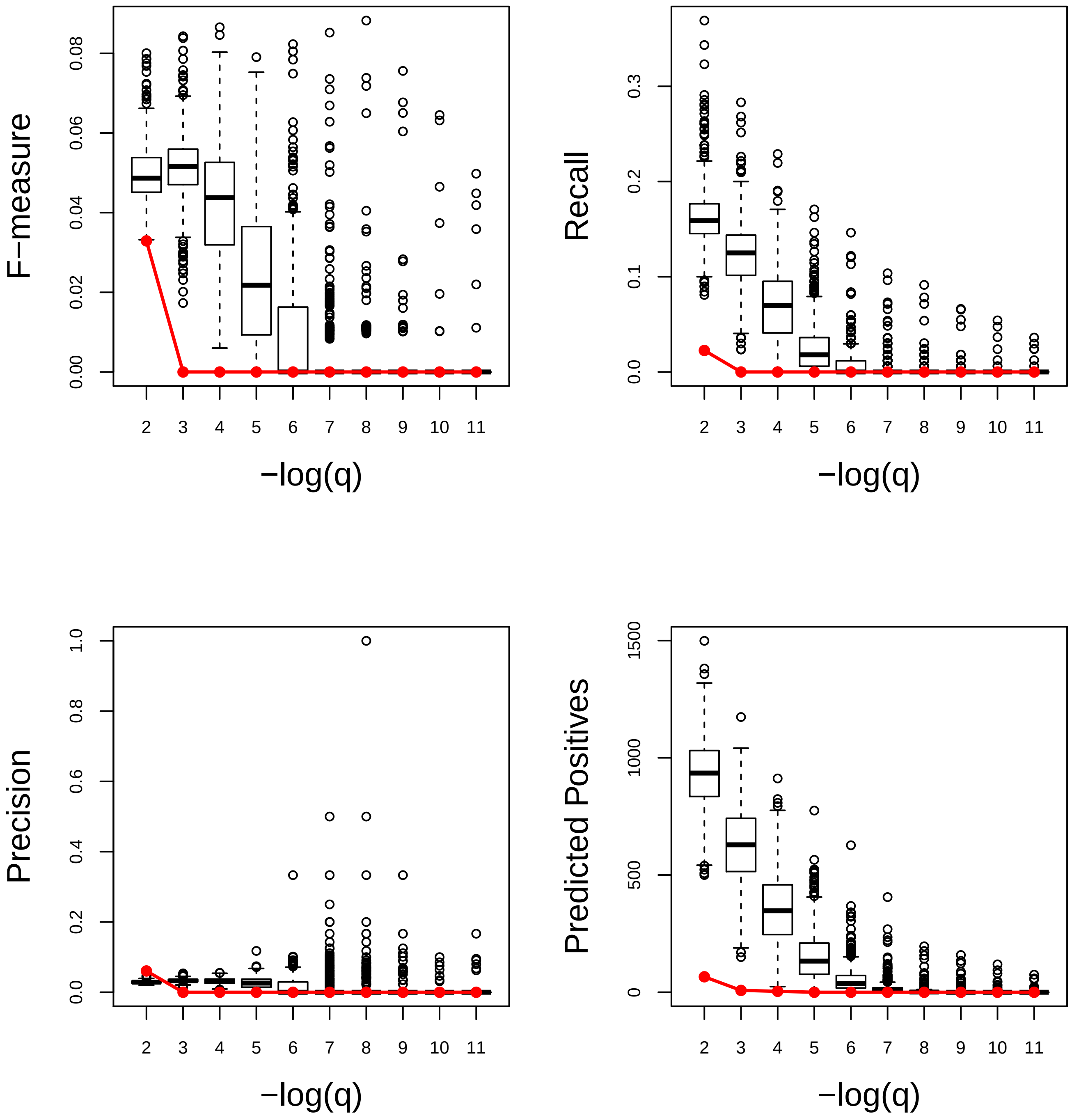

### Regulation of response to osmotic stress. Very Good

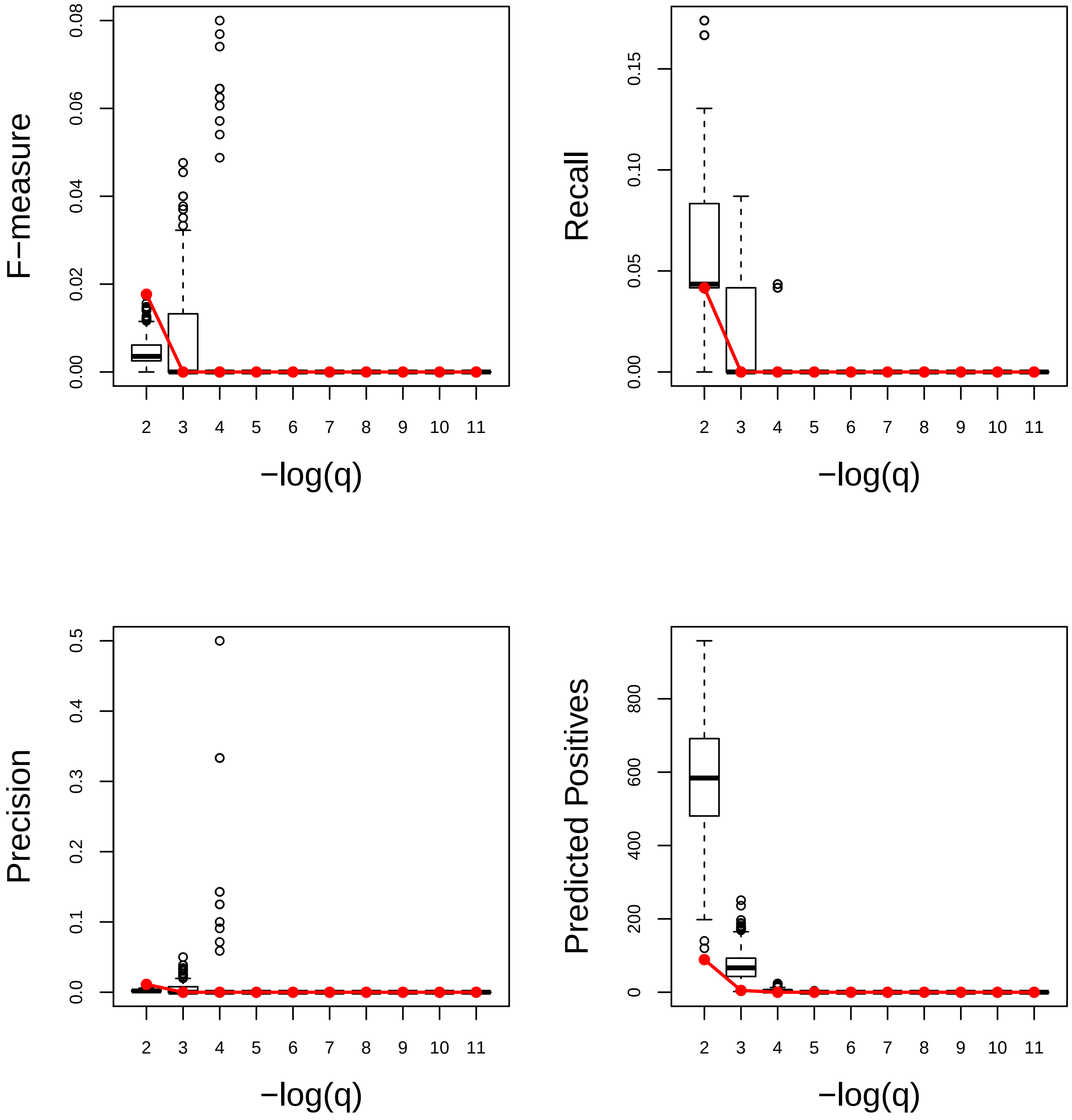

### Regulation of response to stress. Very Good

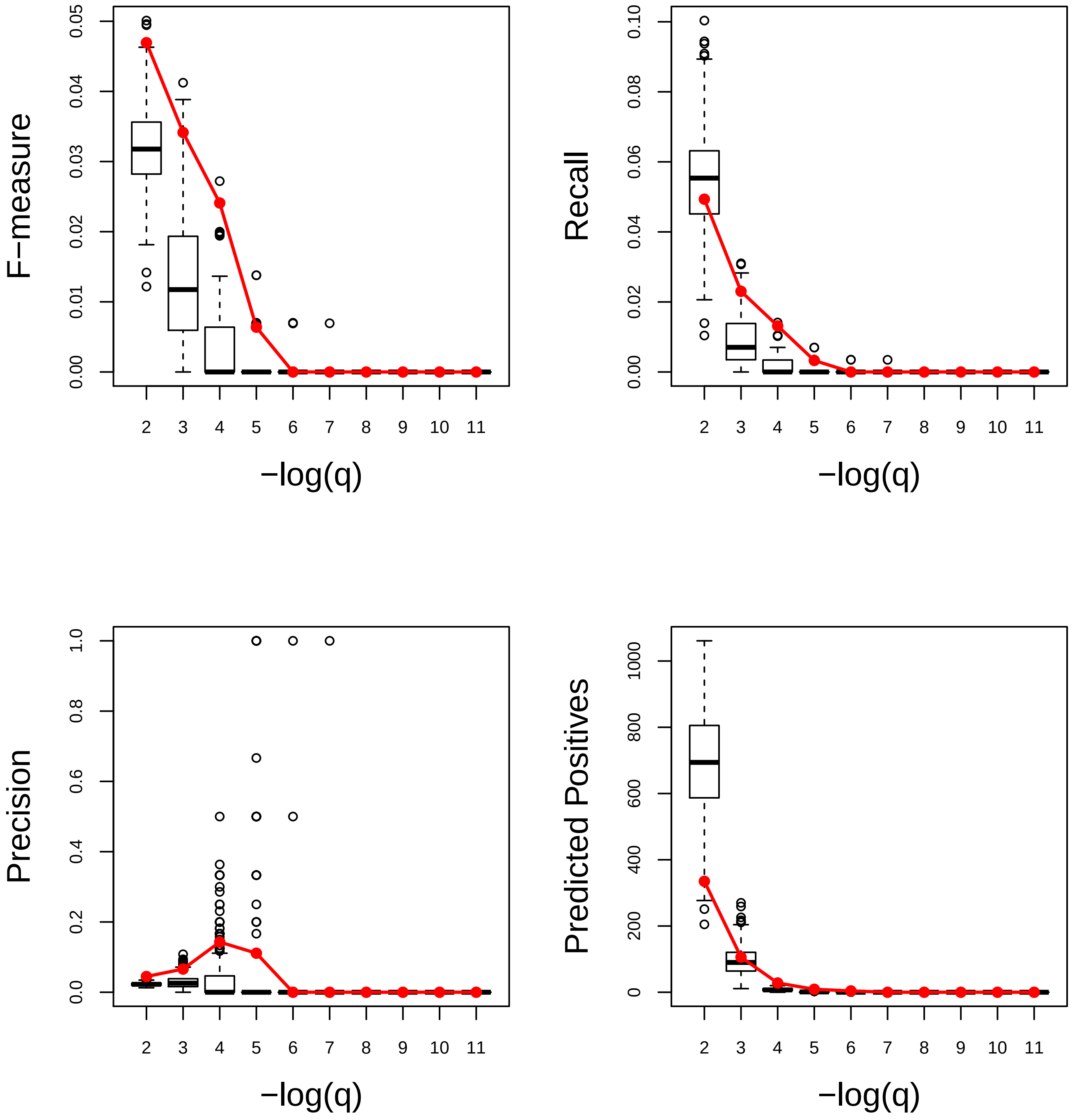

### Regulation of signal transduction. Very Good

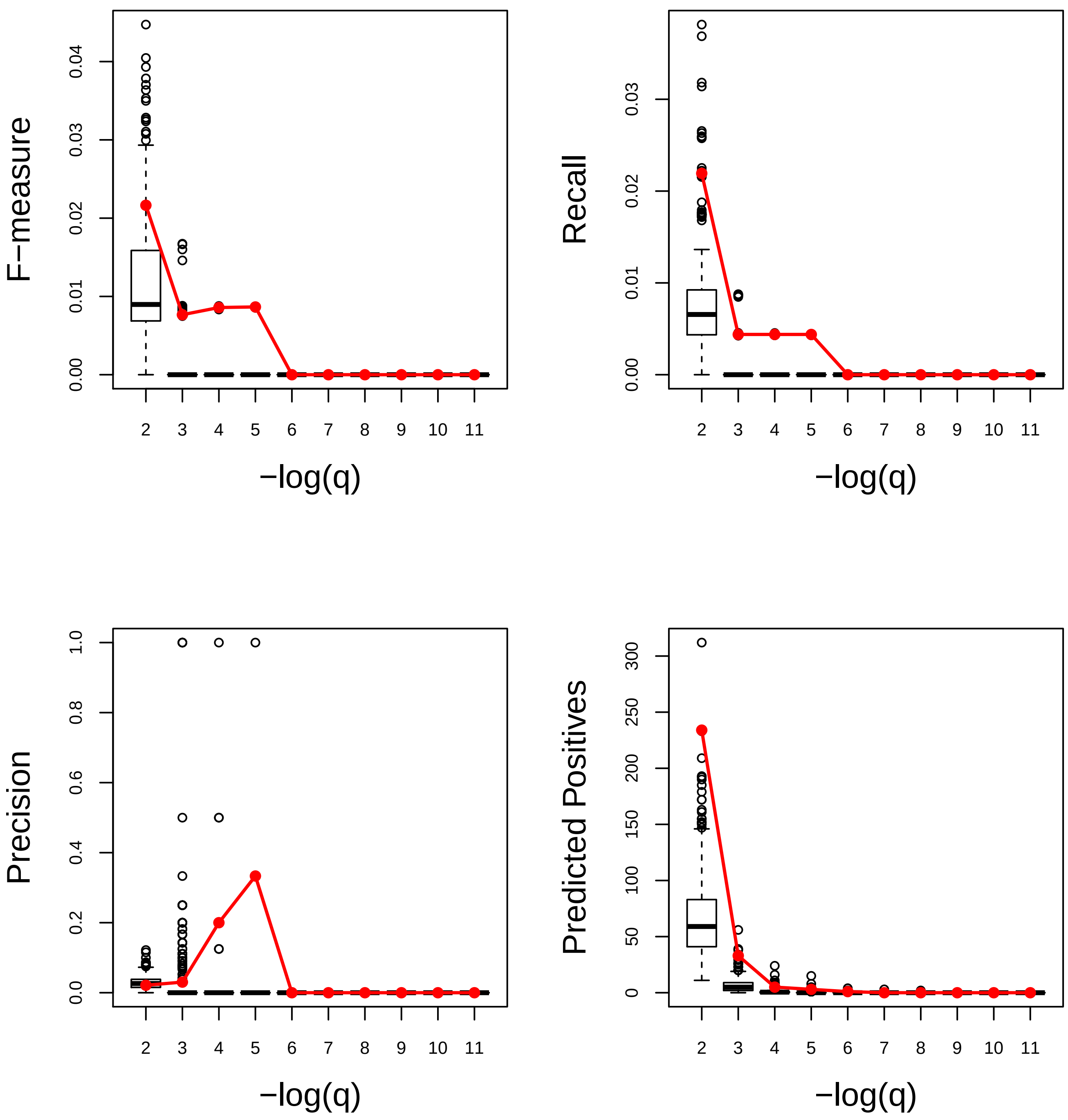

Reproduction. Very Poor

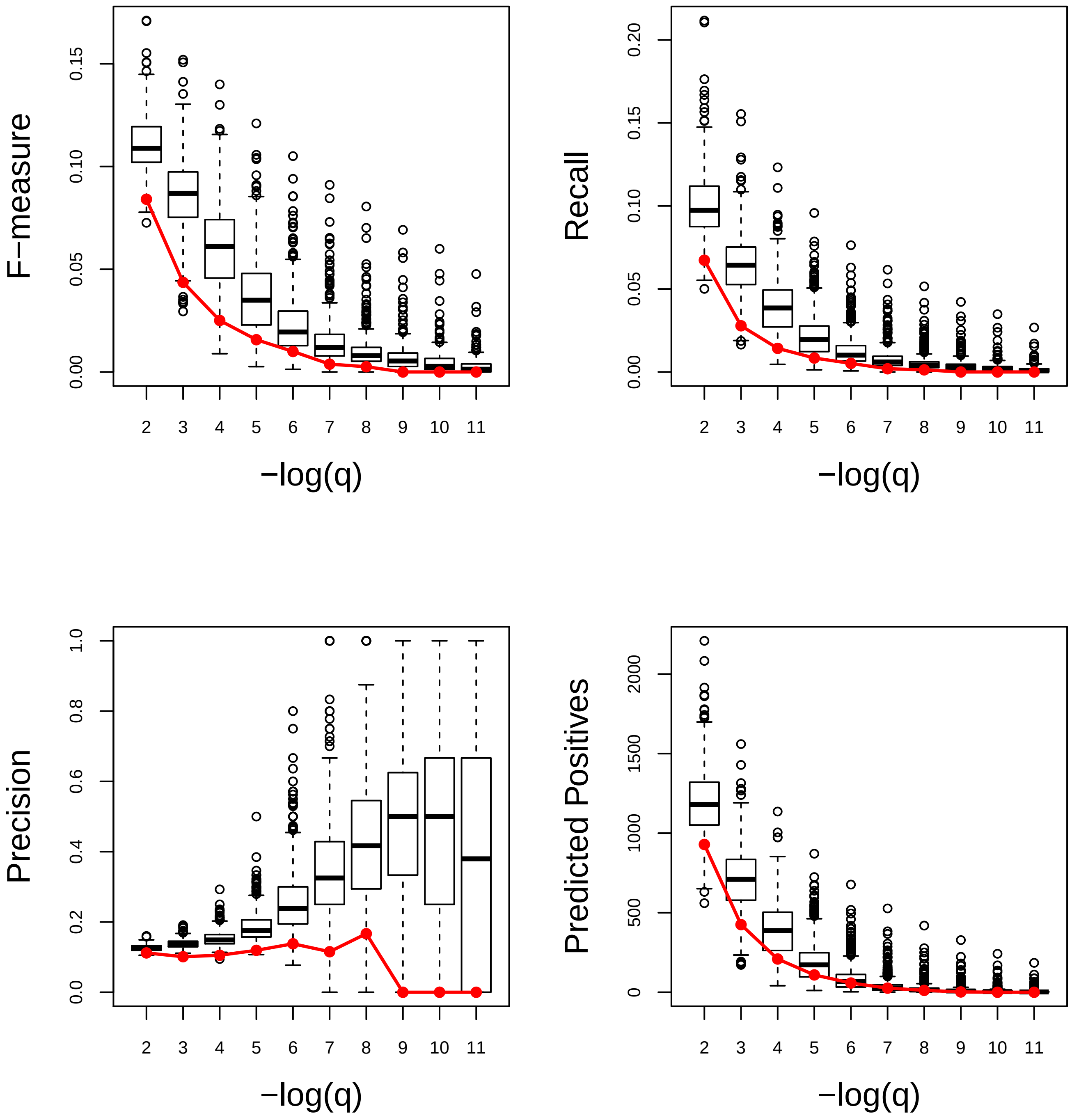

### Response to abiotic stimulus. Very Good

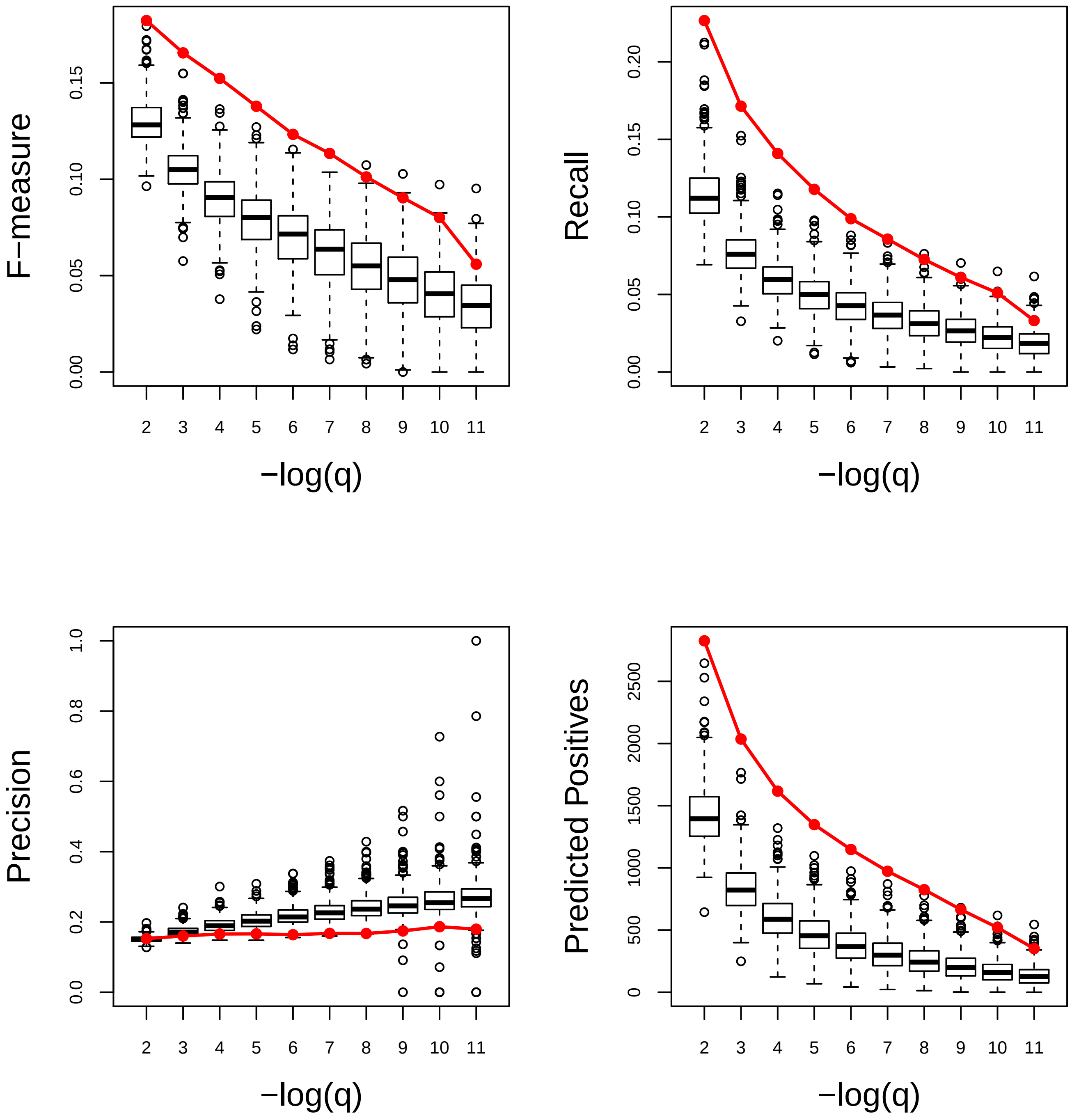

### Response to abscisic acid. Very Good

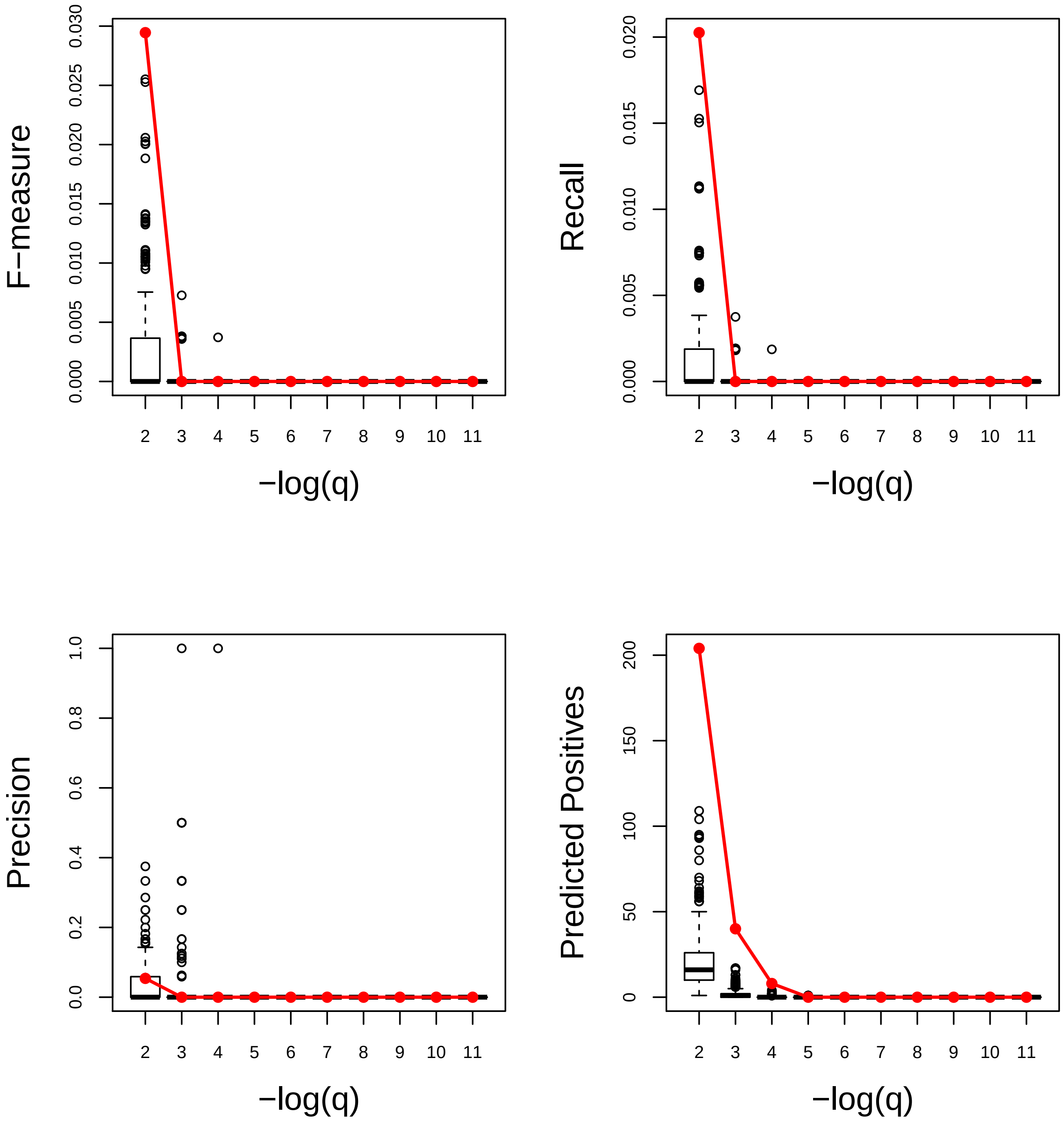

### Response to acid chemical. Very Good

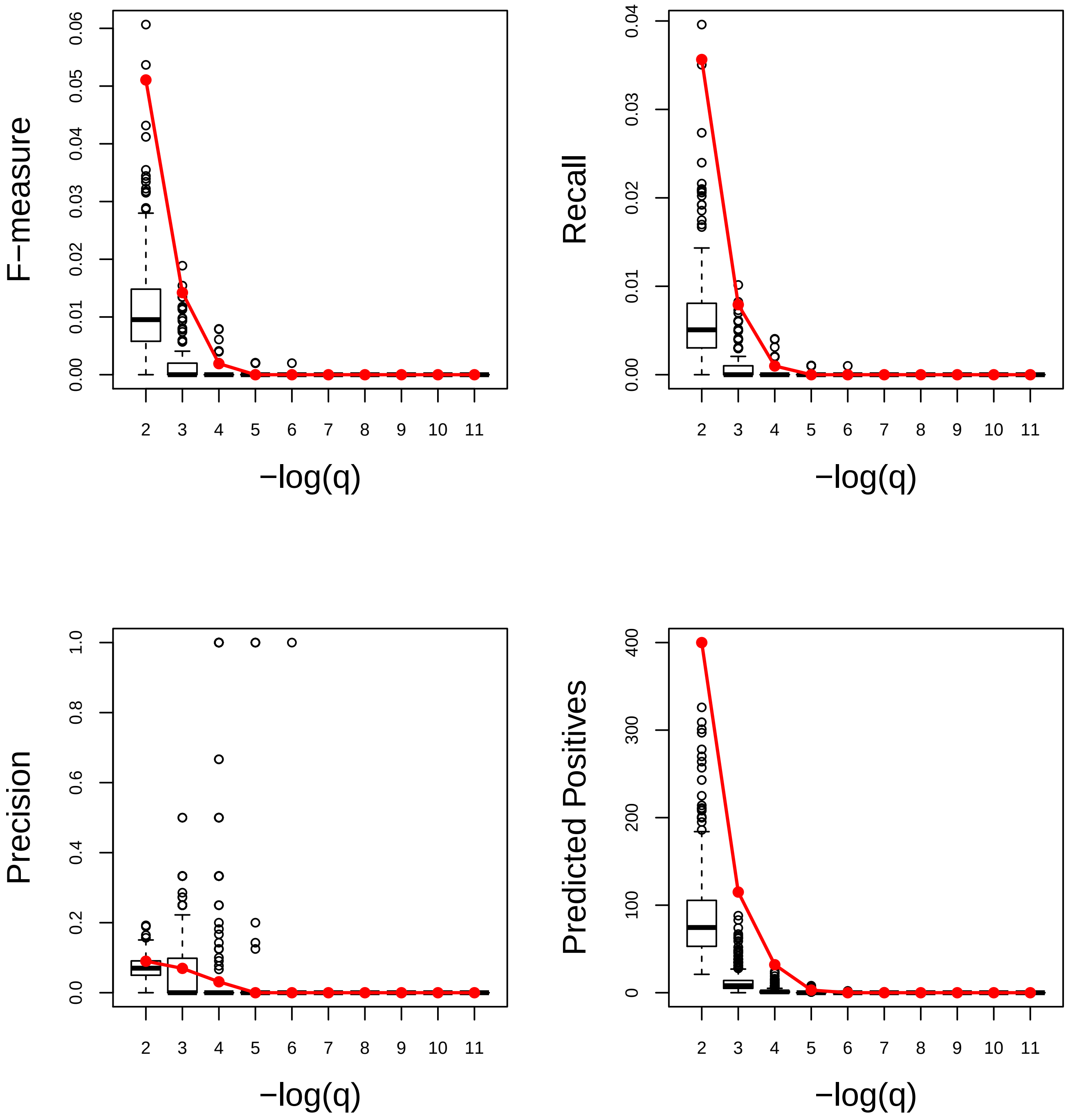

### Response to auxin. Very Poor

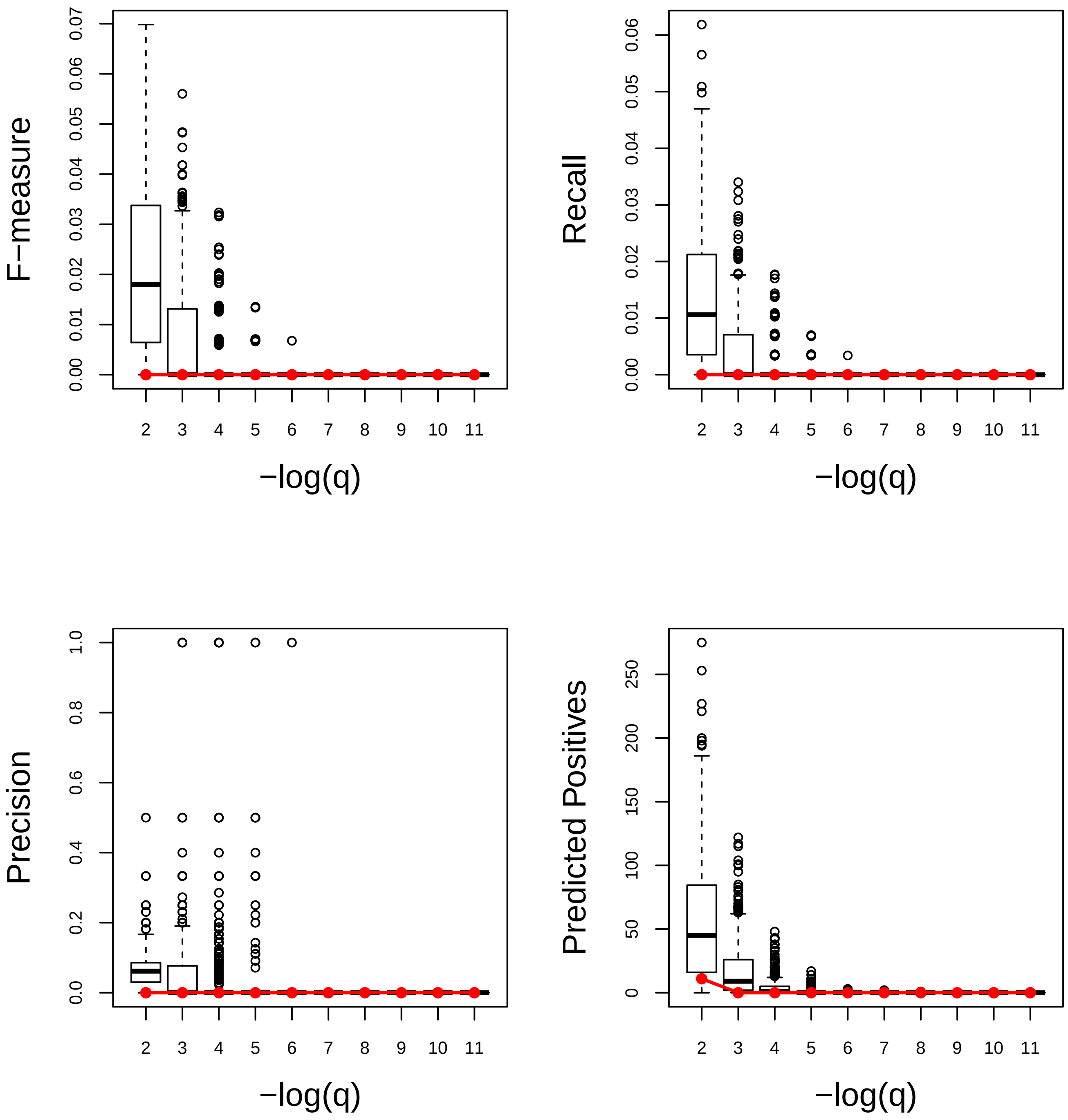

### Response to bacterium. Very Good

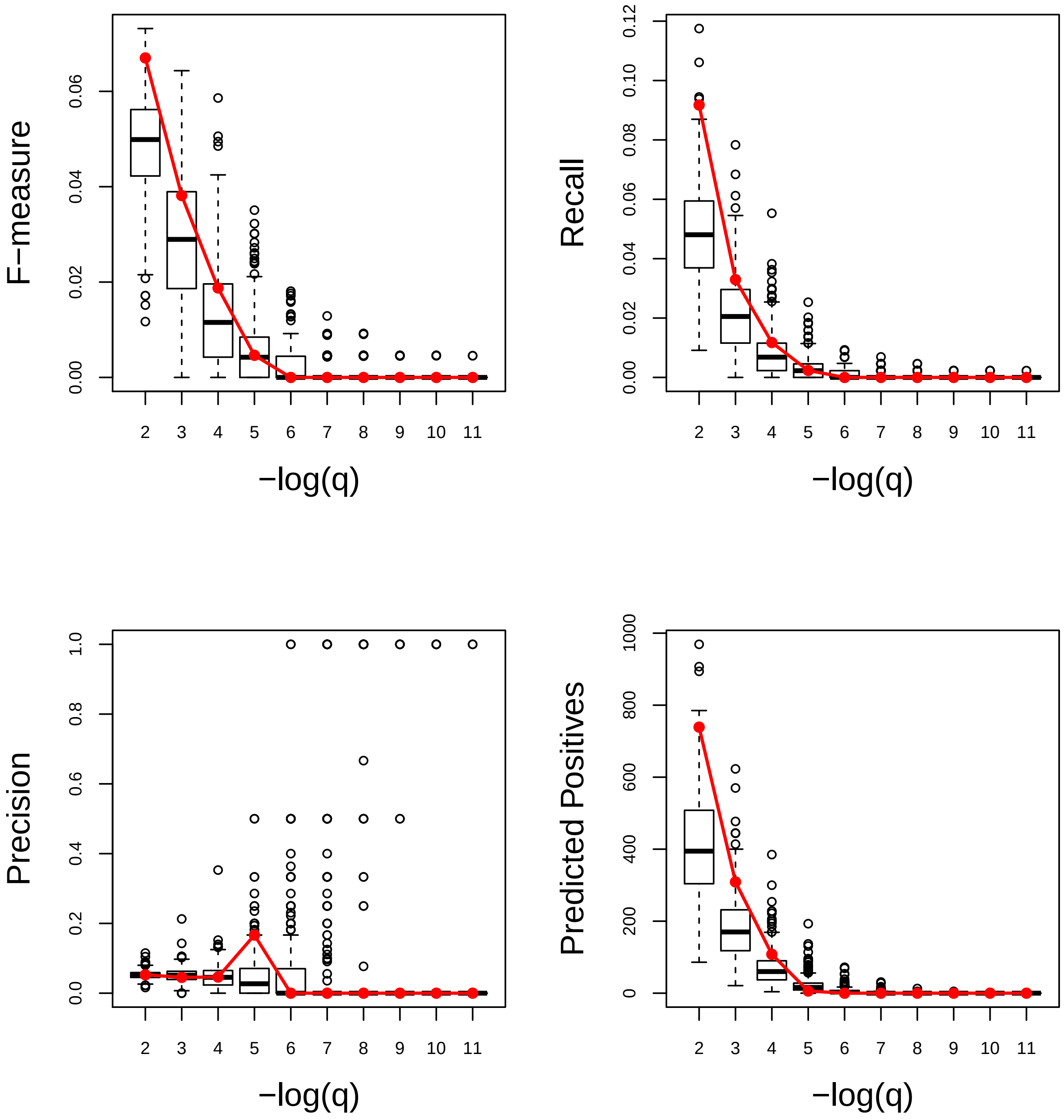

### Response to biotic stimulus. Average

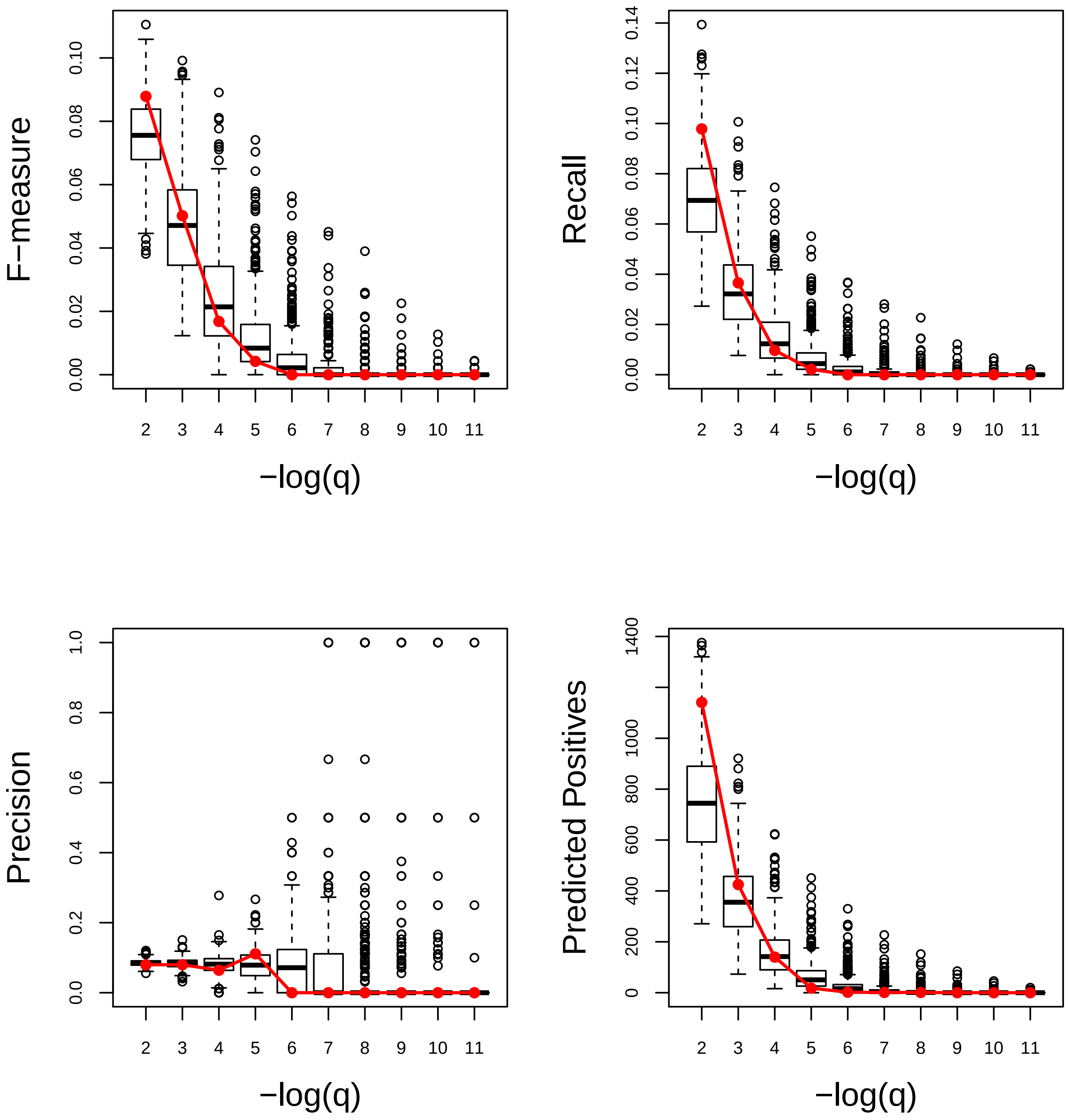

### Response to blue light. Very Good

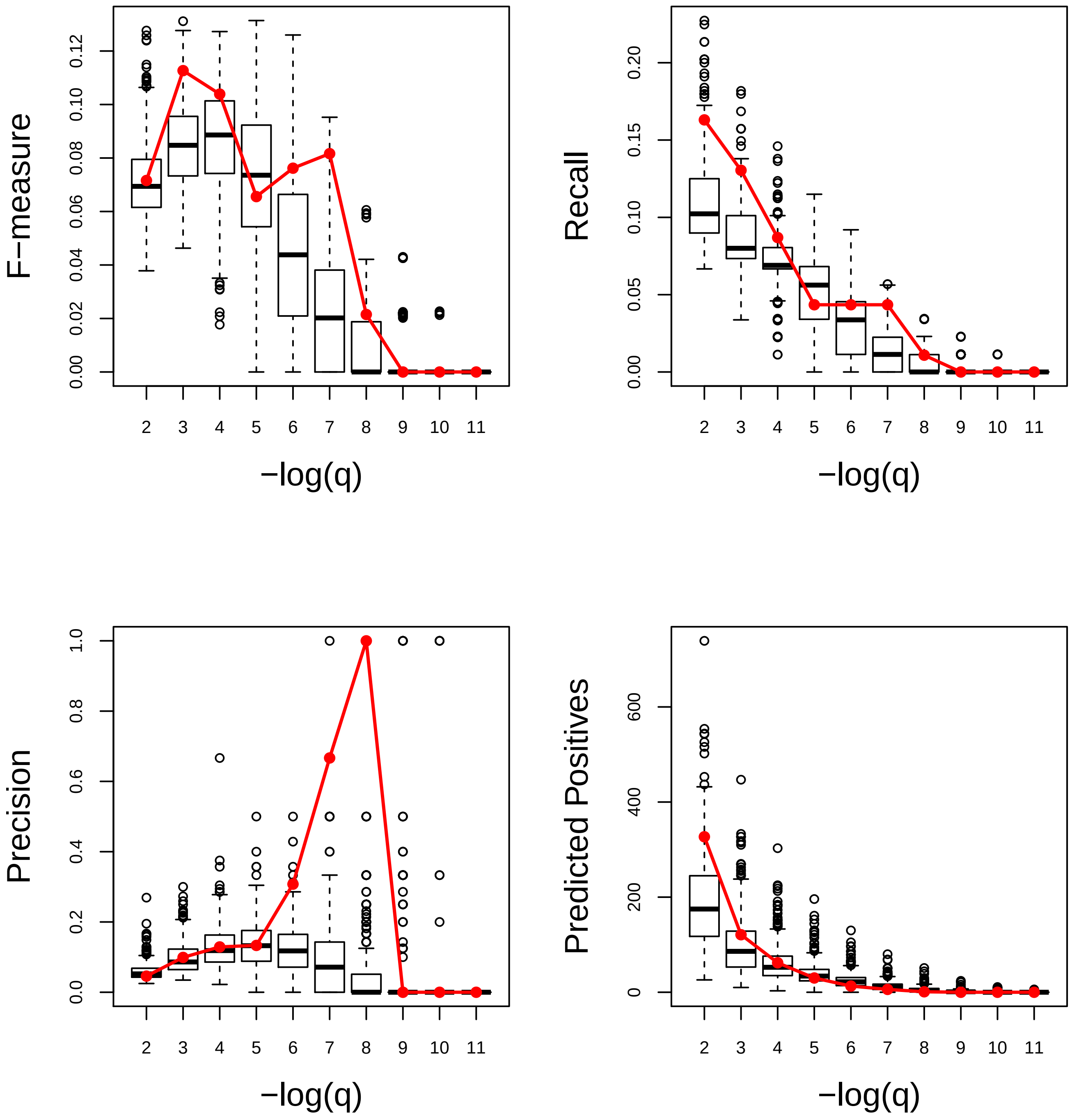

### Response to brassinosteroid. Average

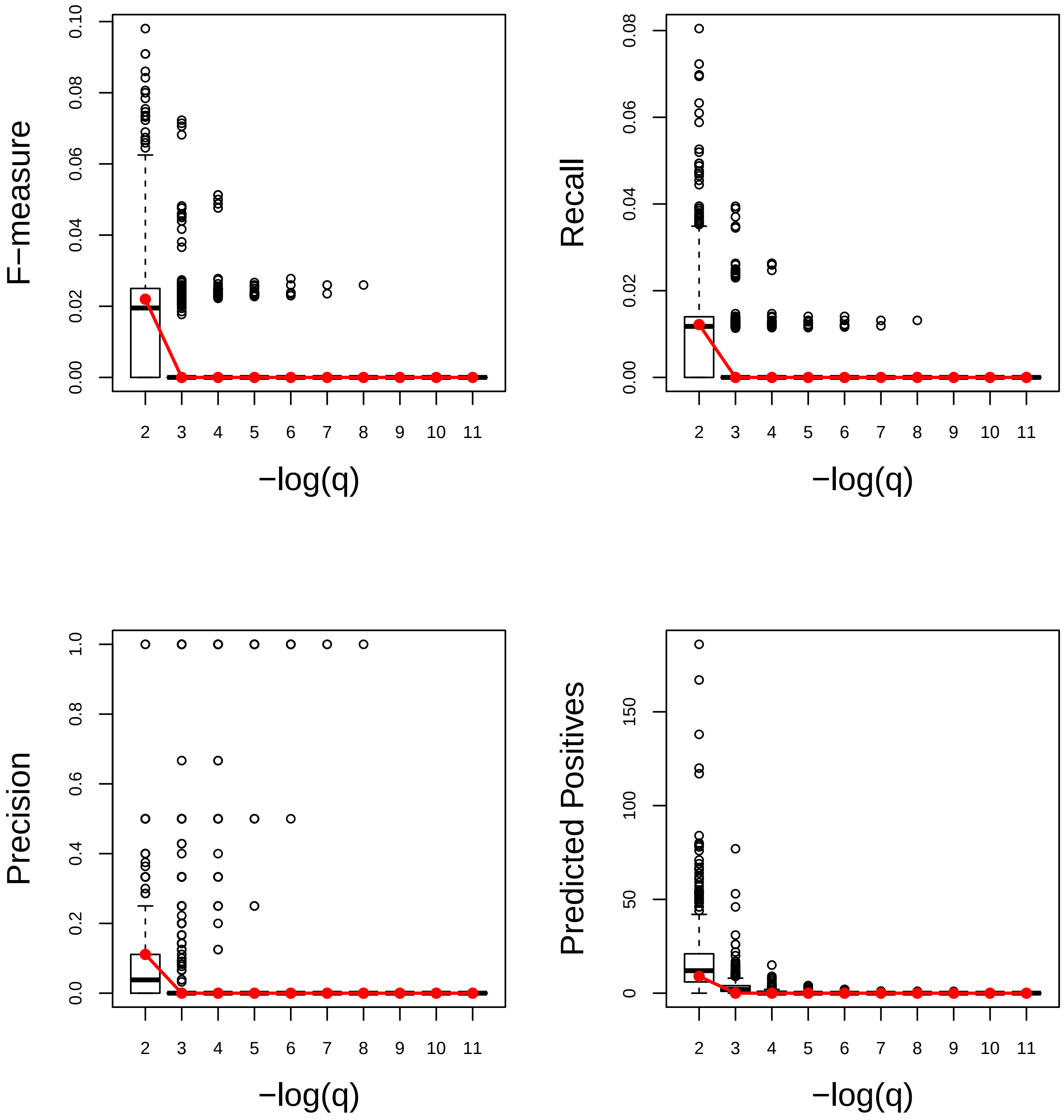

### Response to chitin. Very Good

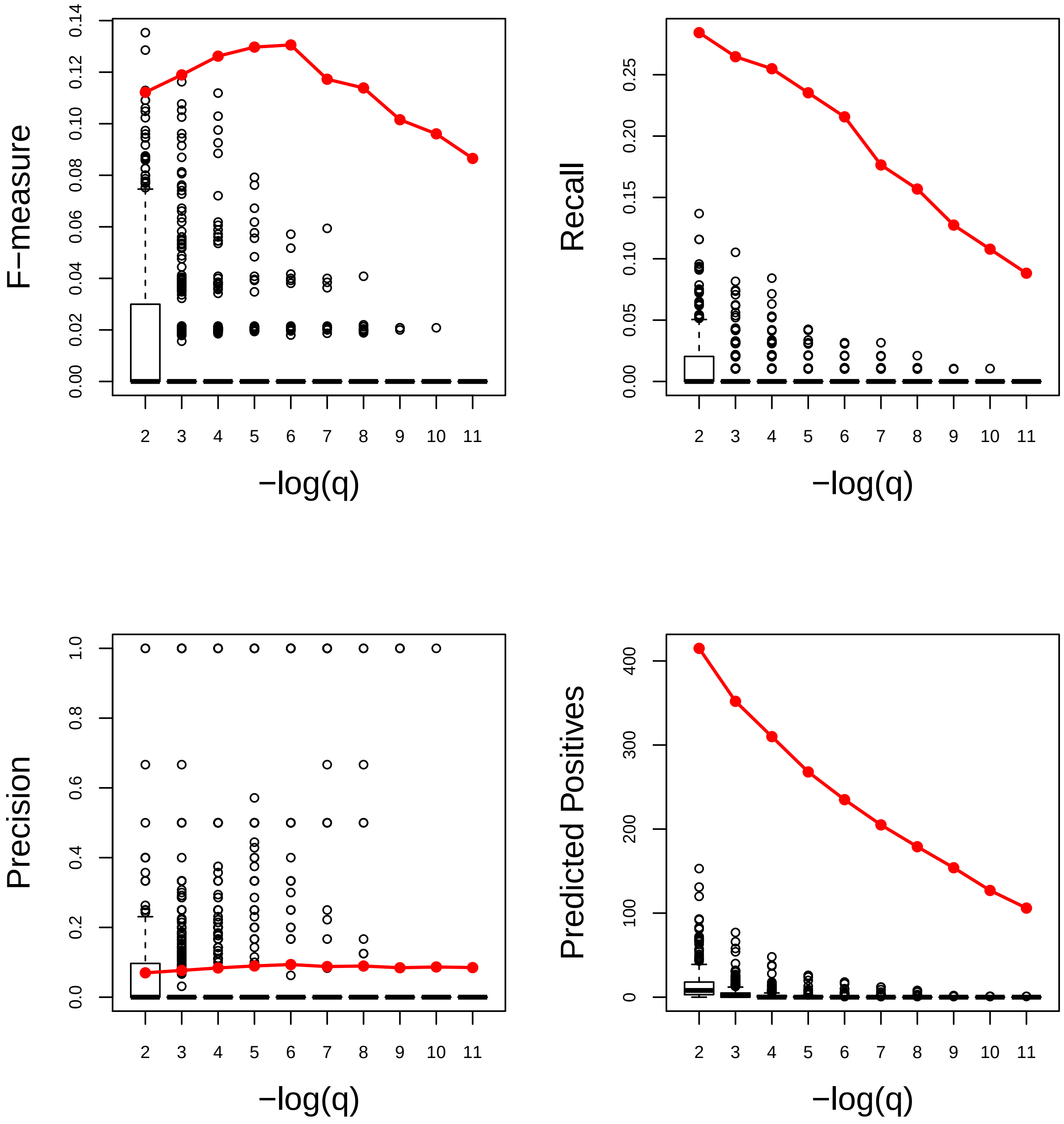

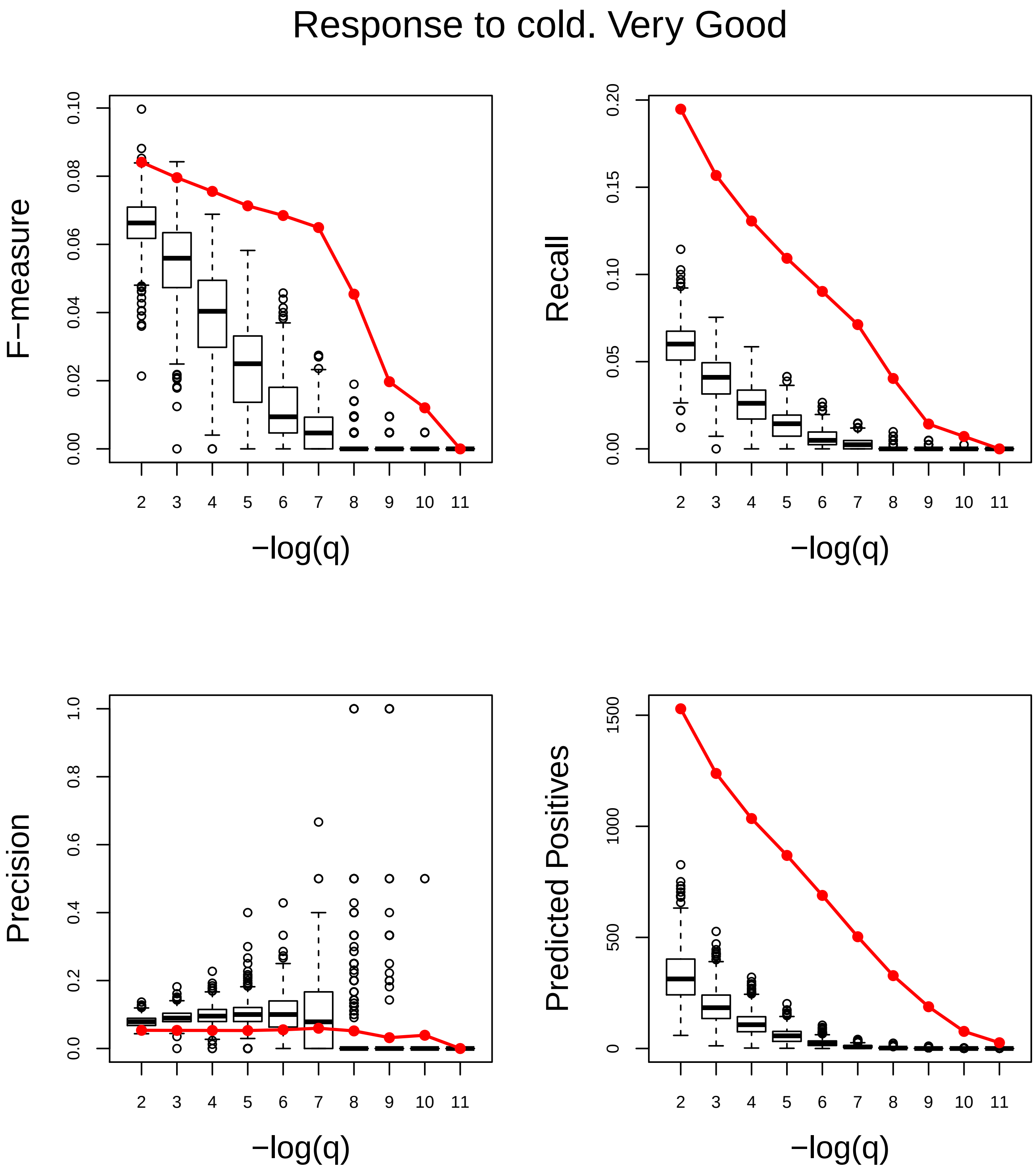
Response to cold. Very Good
F−measure
Recall
Precision
Predicted Positives
−log(q)
−log(q)
−log(q)
−log(q)

### Response to cytokinin. Very Good

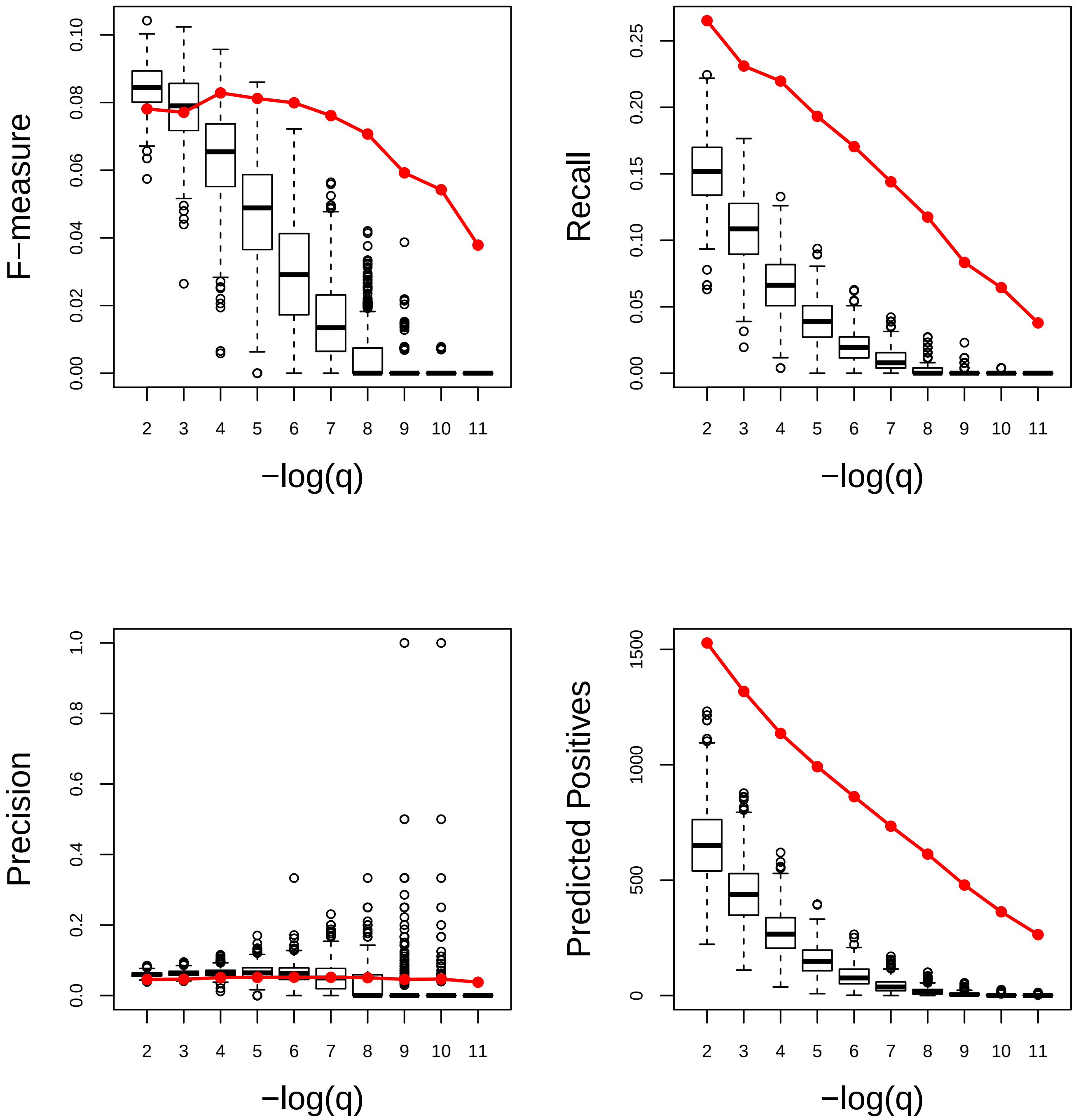

### Response to endogenous stimulus. Very Good

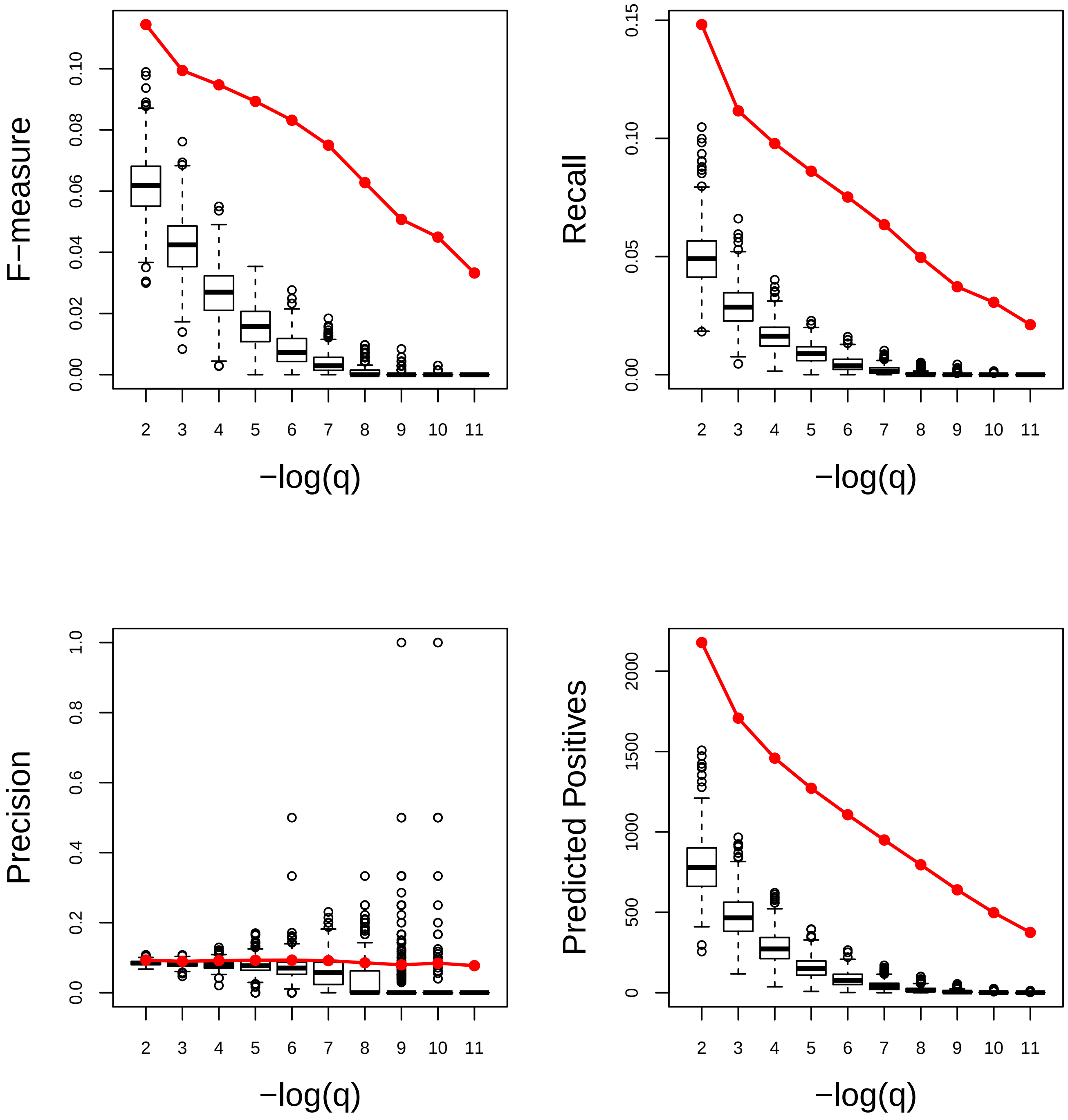

### Response to endoplasmic reticulum stress. Very Good

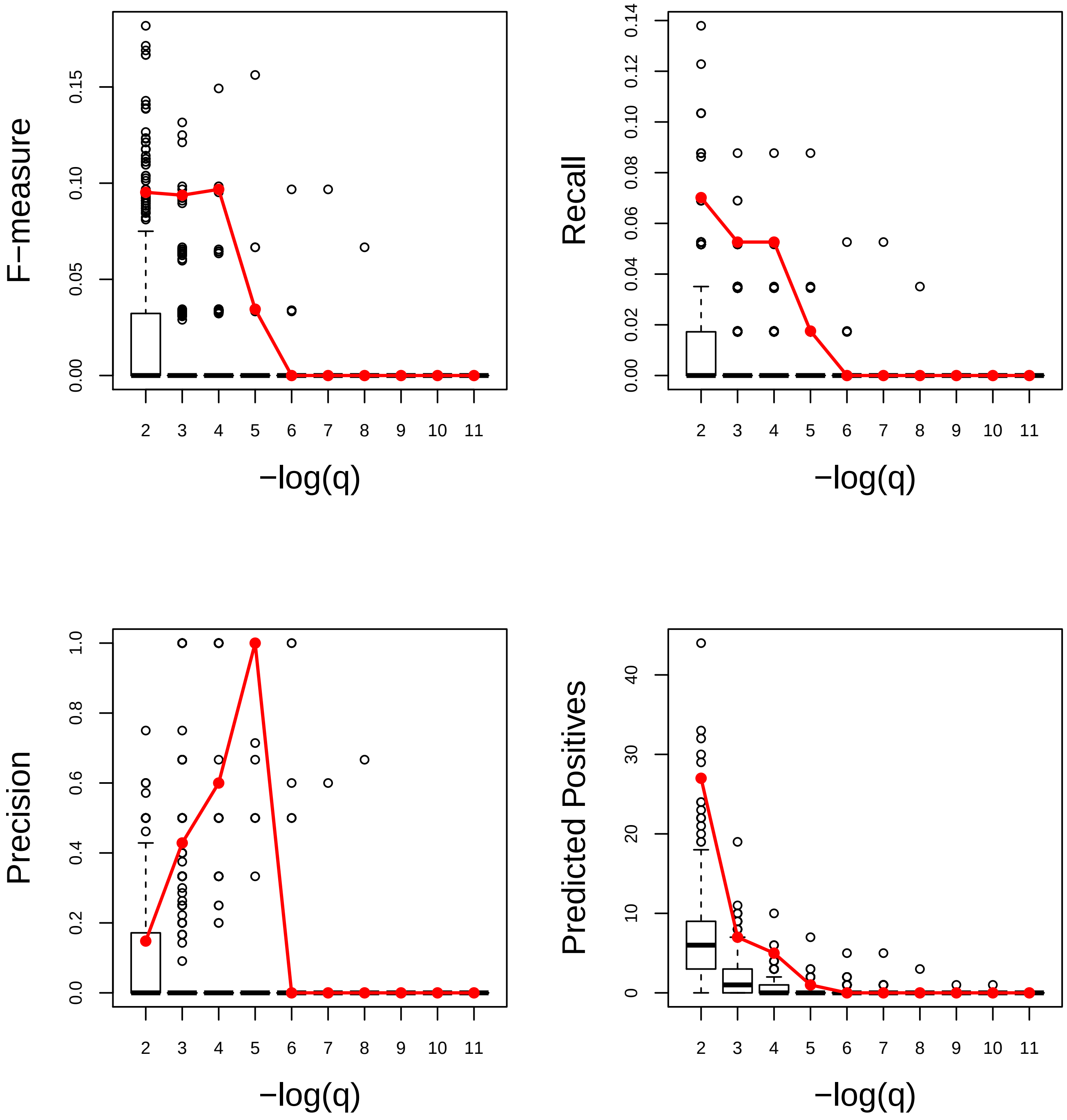

### Response to ethylene. Very Good

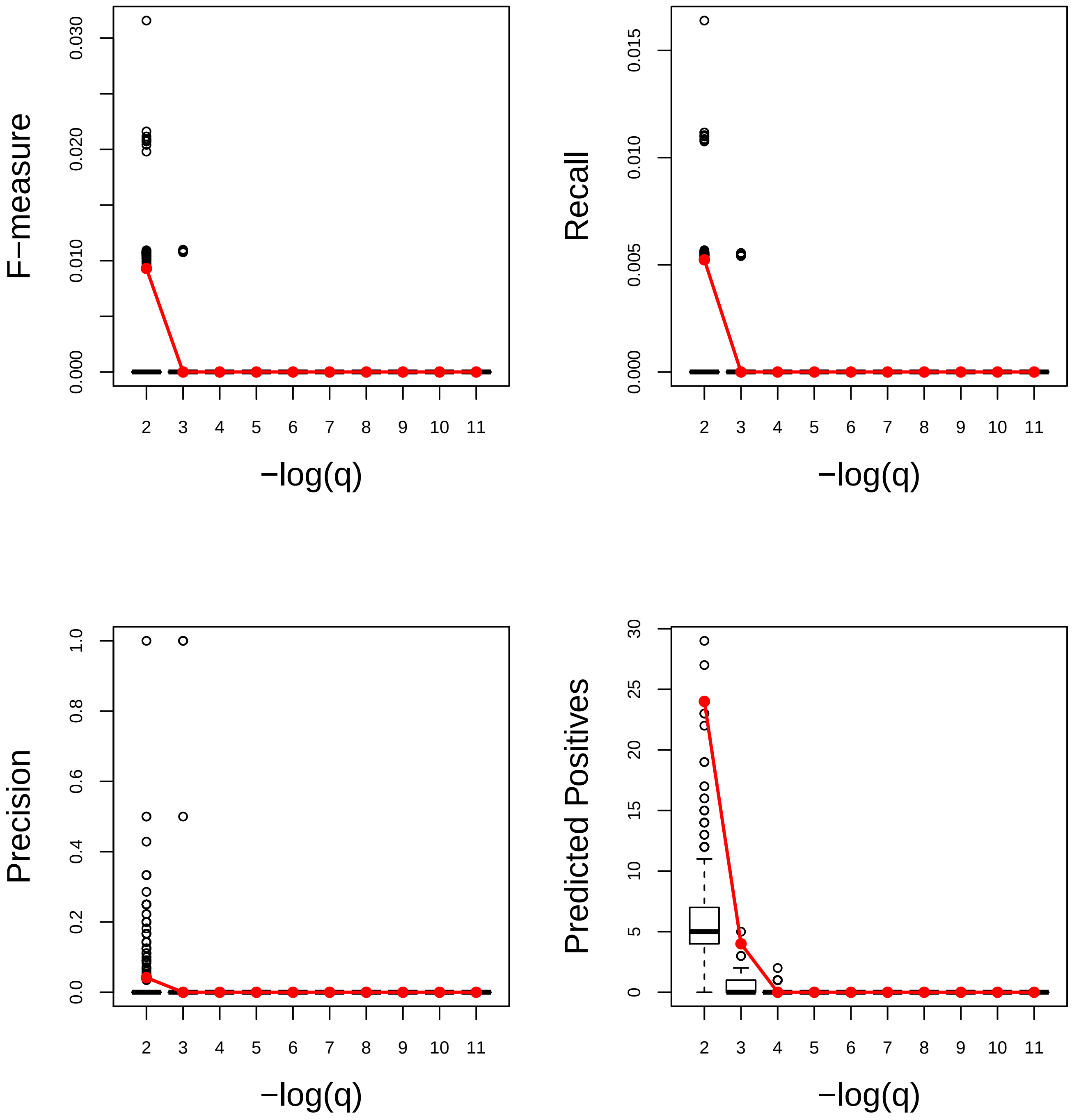

### Response to external stimulus. Average

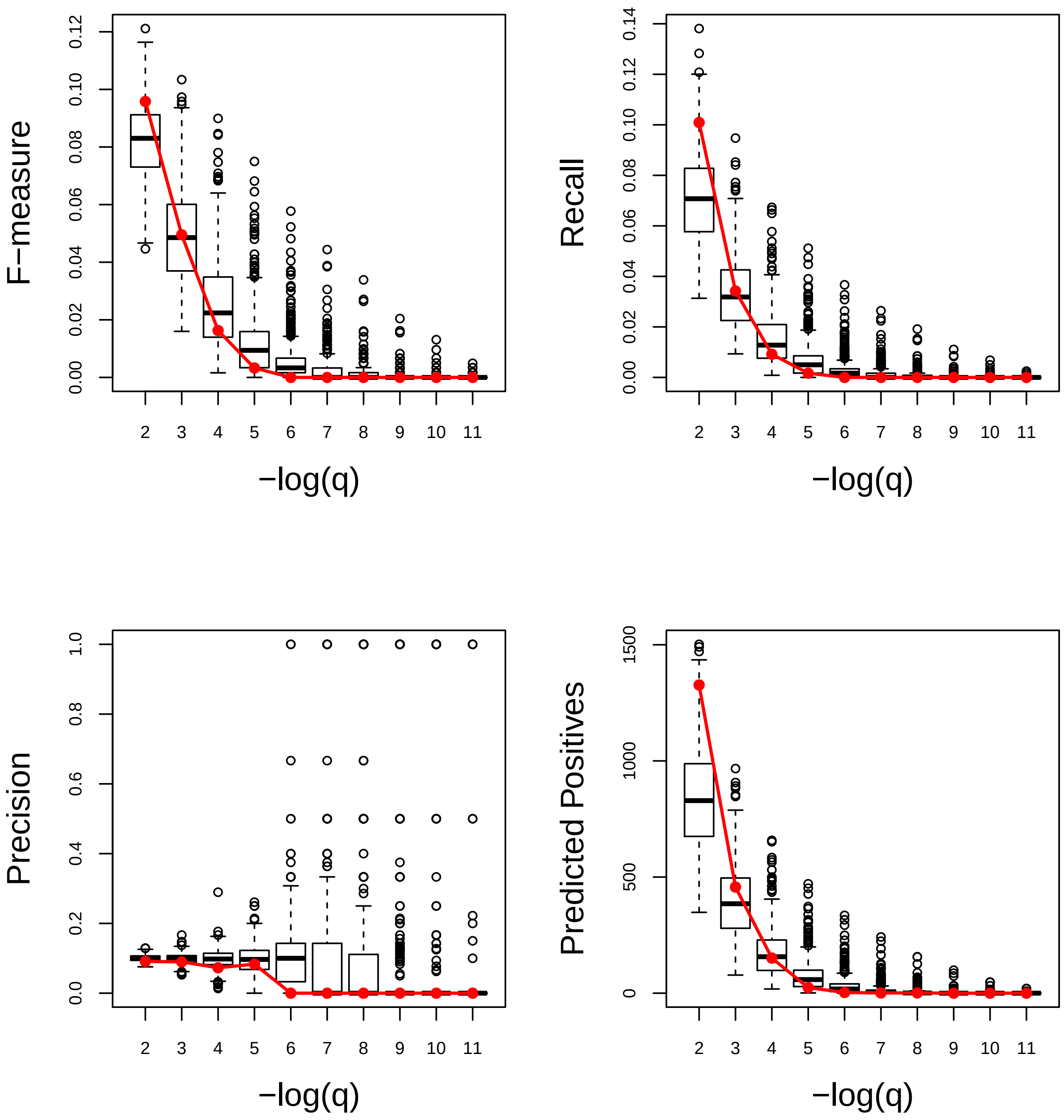

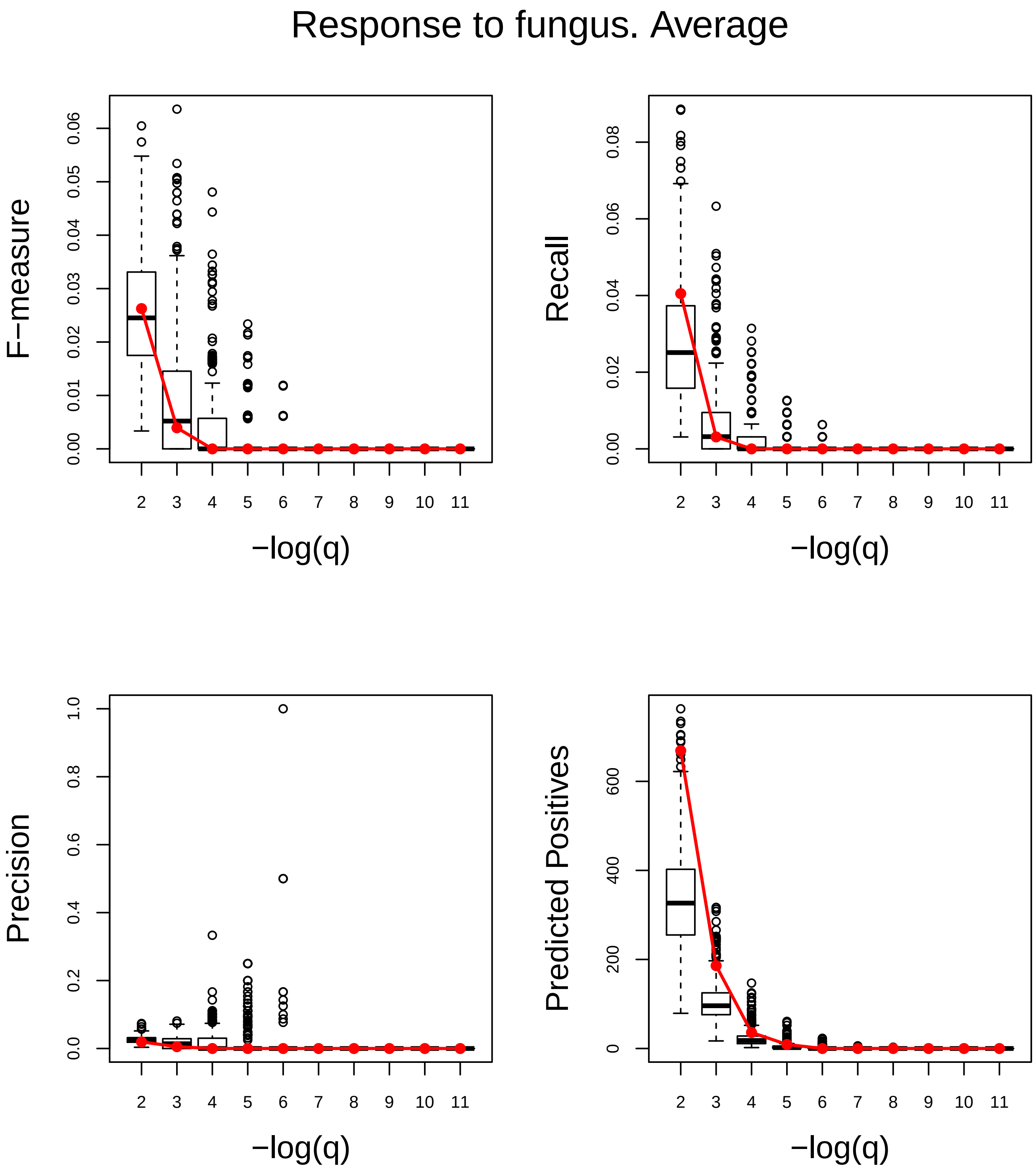
Response to fungus. Average
F−measure
0.00
0.01
0.02
0.03
0.04
0.05
0.06
Recall
0.00
0.02
0.04
0.06
0.08
Precision
0.0
0.2
0.4
0.6
0.8
1.0
Predicted Positives
0
200
400
600
2 3 4 5 6 7 8 9 10 11
−log(q)

### Response to gravity. Very Good

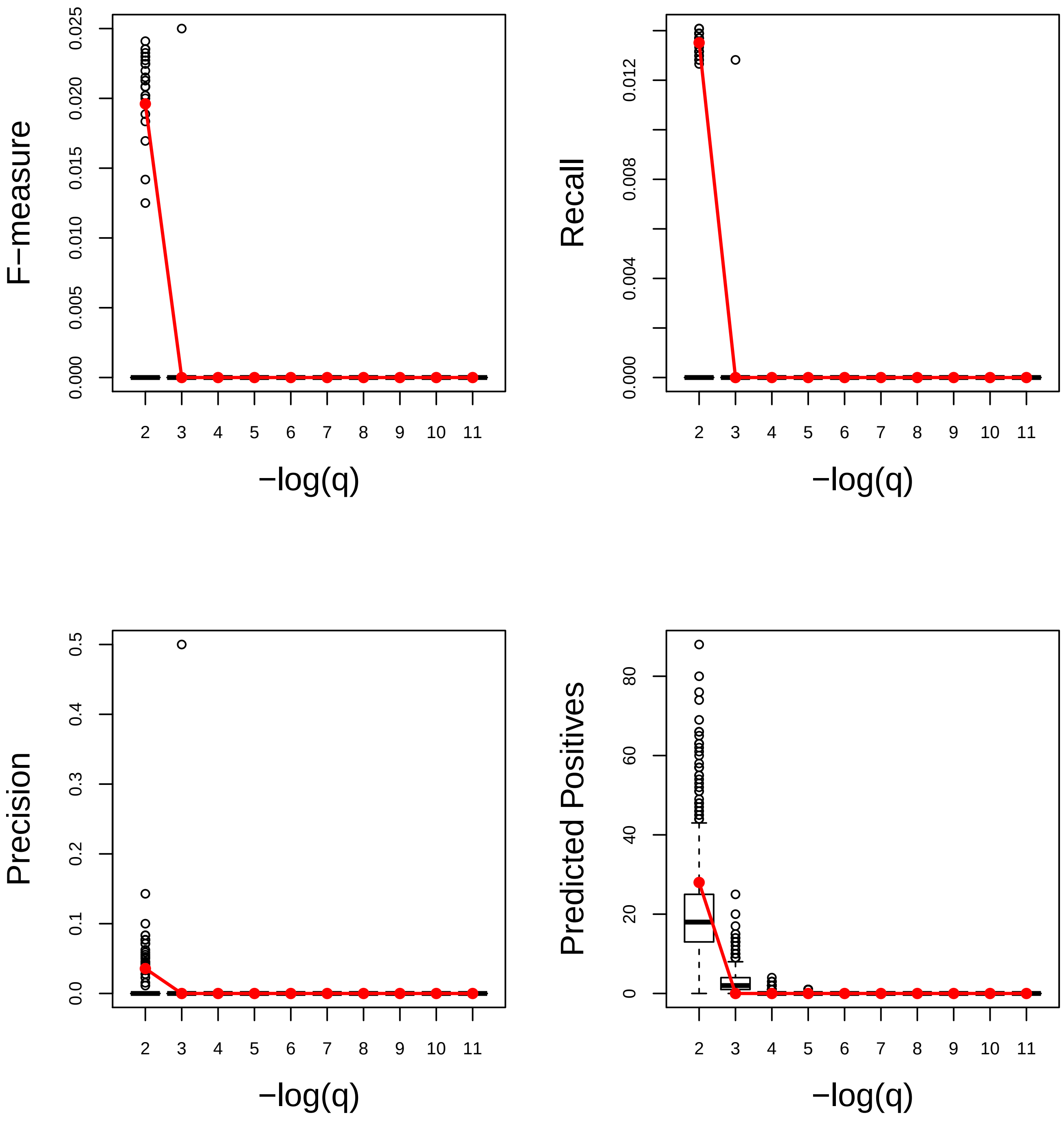

### Response to heat. Very Good

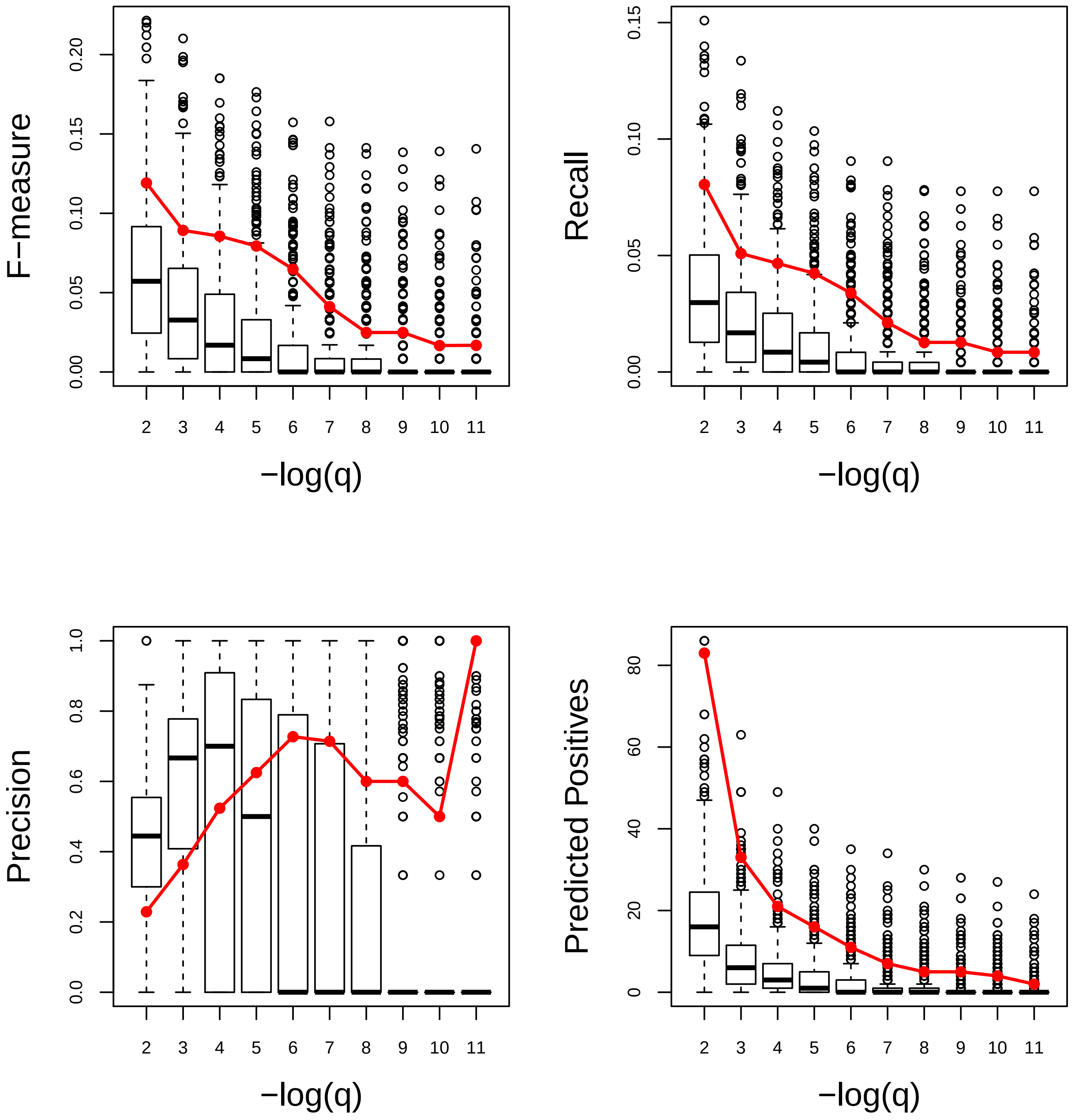

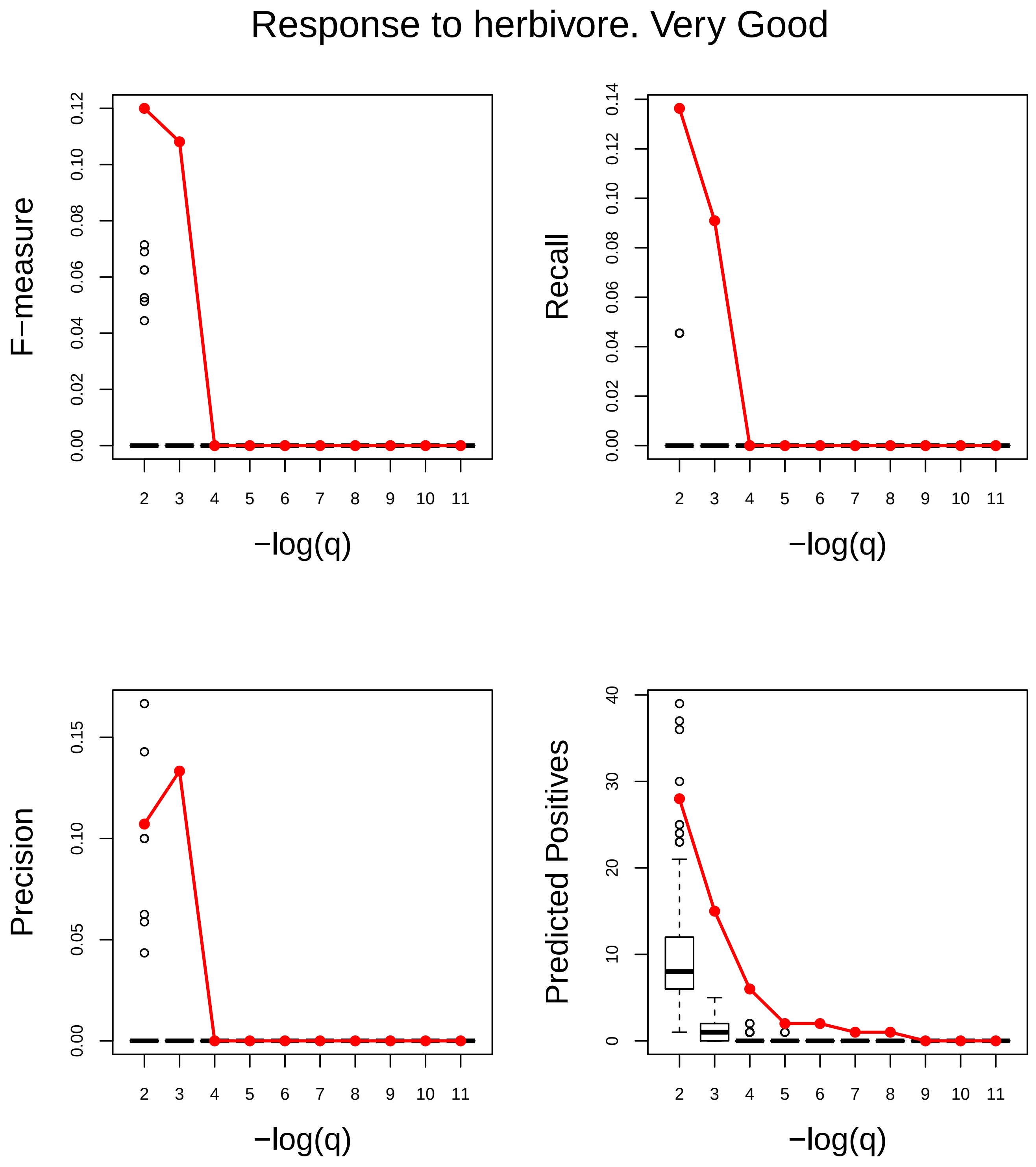
Response to herbivore. Very Good
F−measure
Recall
Precision
Predicted Positives
−log(q)
−log(q)
−log(q)
−log(q)

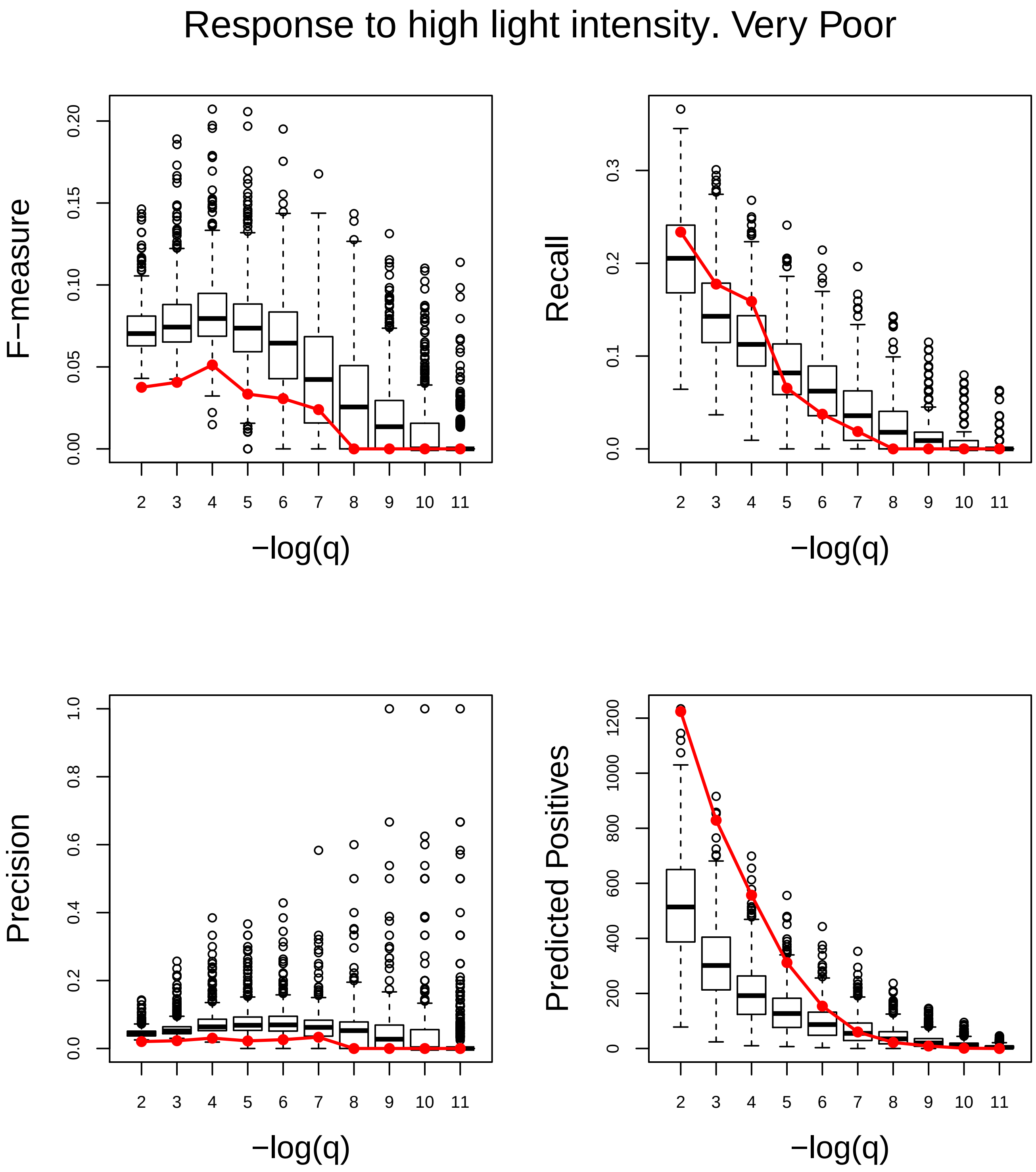
Response to high light intensity. Very Poor
F−measure
Recall
Precision
Predicted Positives
−log(q)
−log(q)
−log(q)
−log(q)
0.00
0.05
0.10
0.15
0.20
0.0
0.1
0.2
0.3
0.0
0.2
0.4
0.6
0.8
1.0
0
200
400
600
800
1000
1200

Response to hormone. Very Good

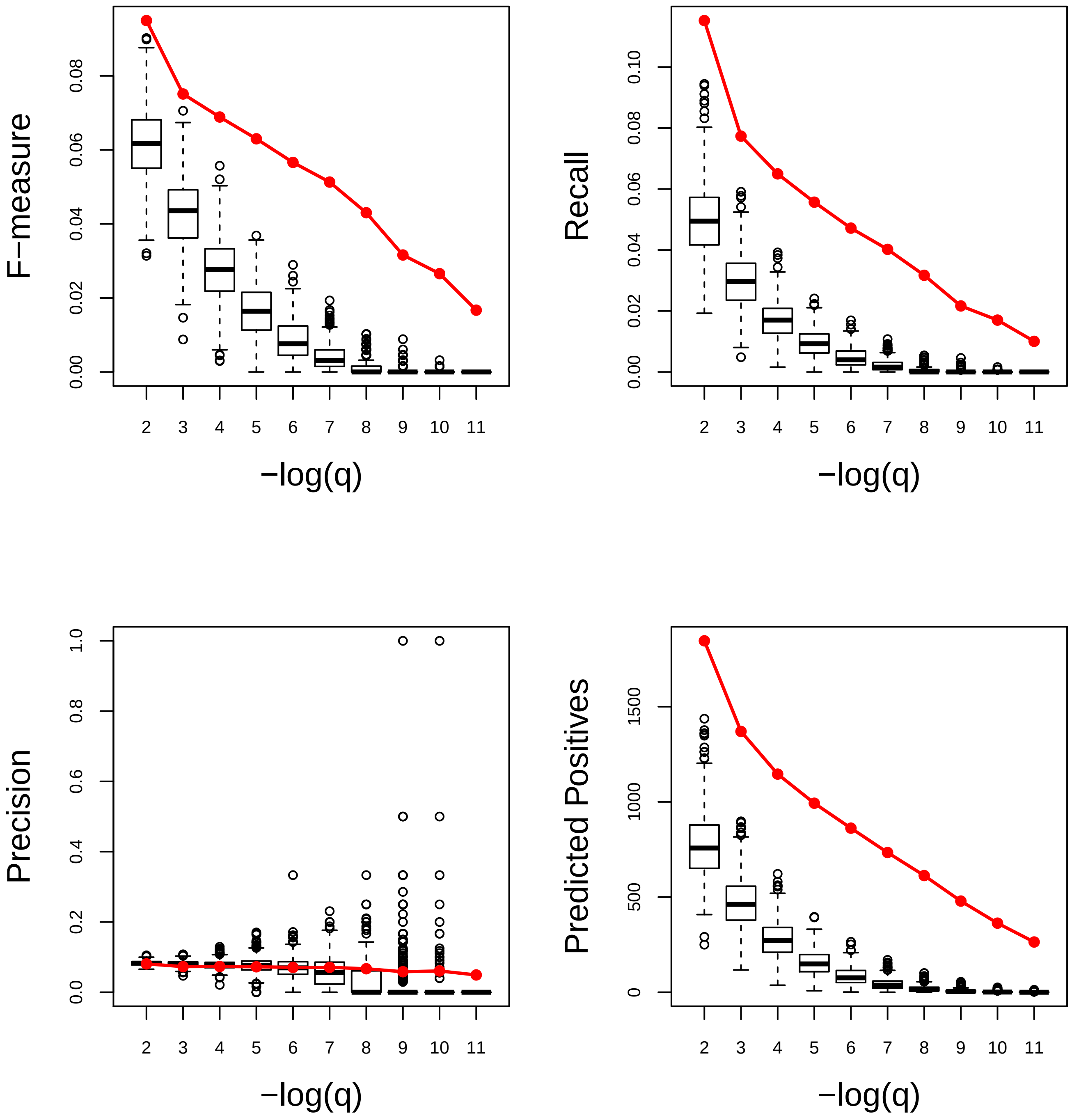

### Response to karrikin. Average

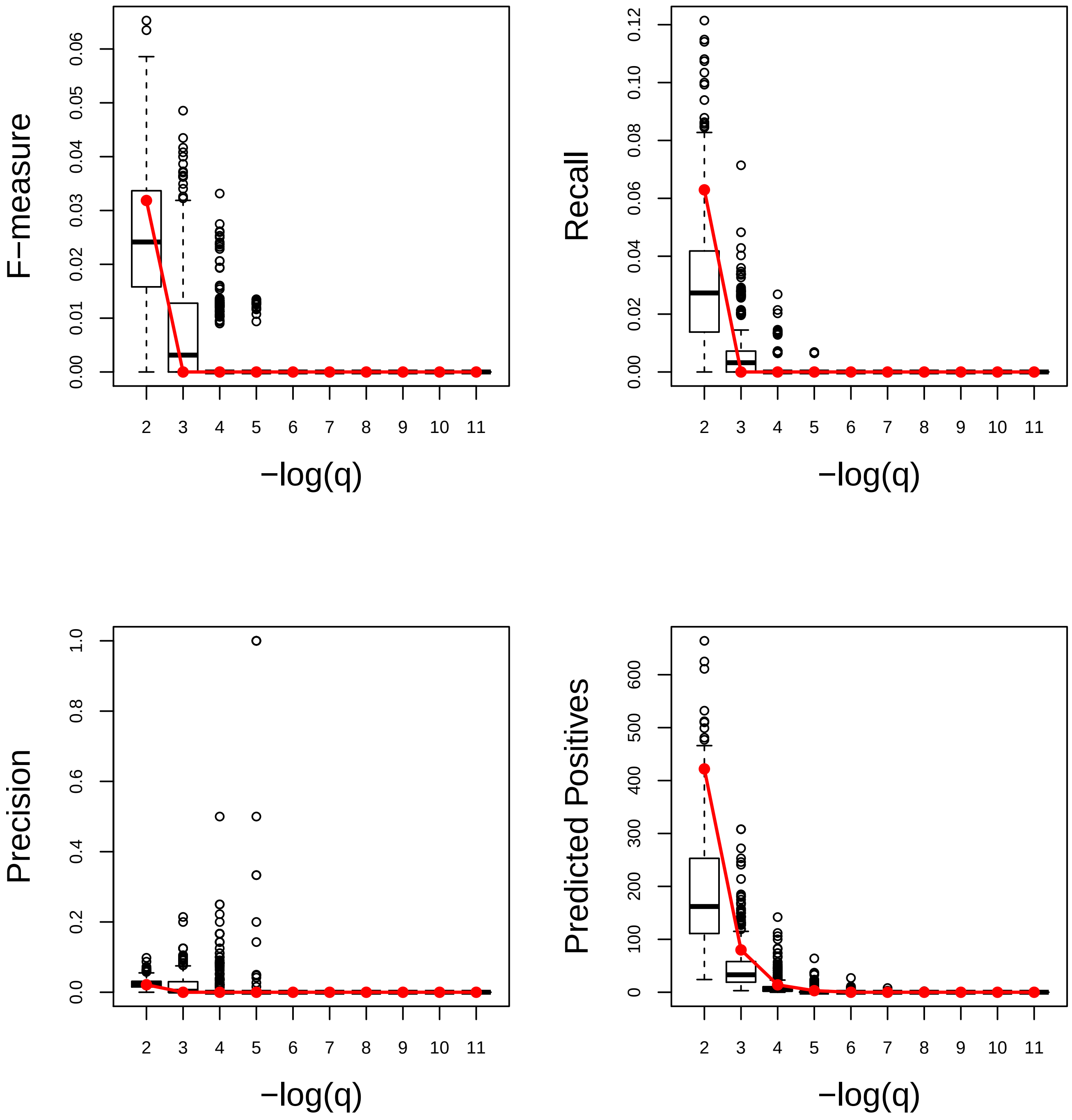

### Response to light intensity. Very Poor

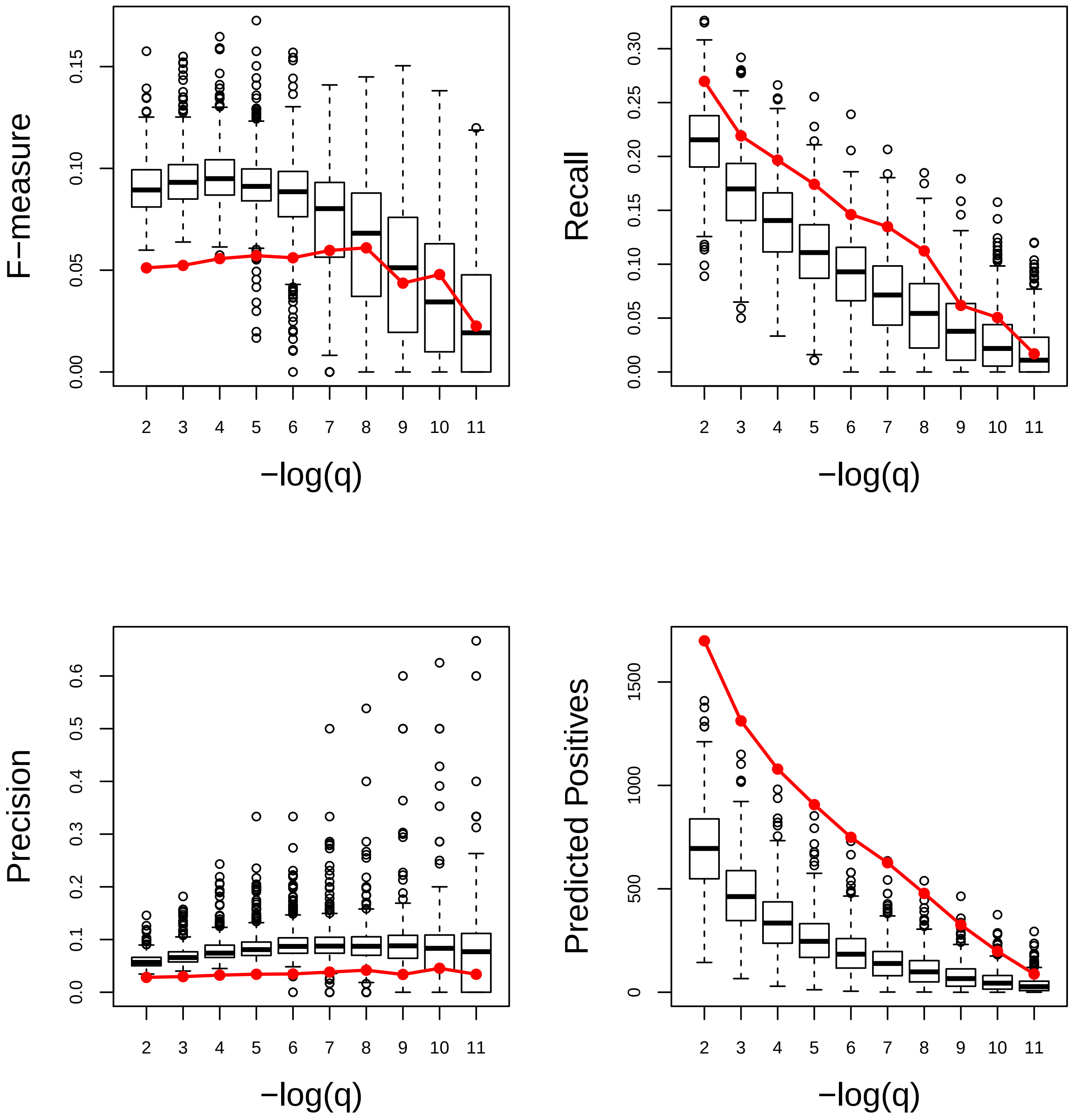

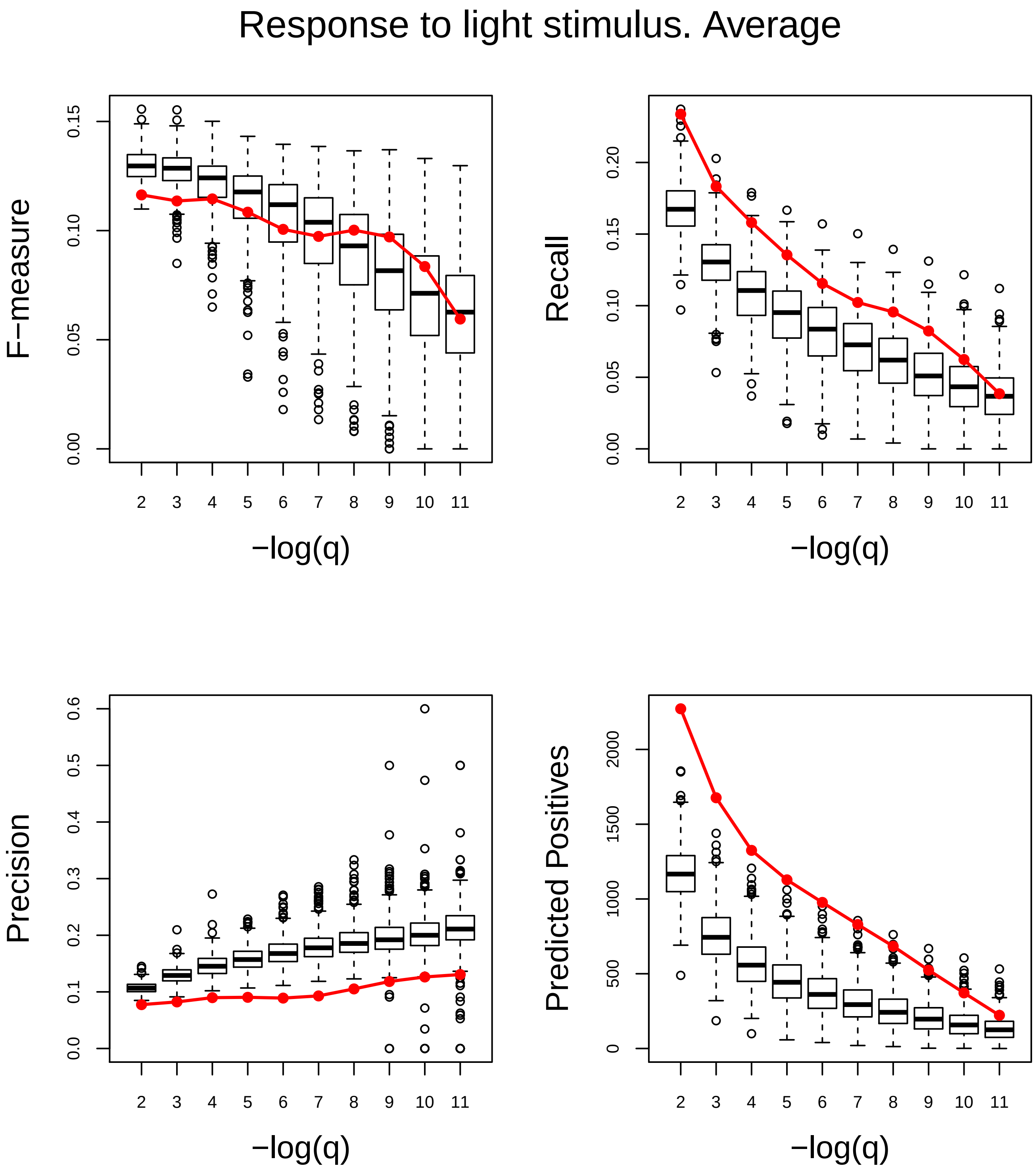
Response to light stimulus. Average
F−measure
0.00
0.05
0.10
0.15
Recall
0.00
0.05
0.10
0.15
0.20
Precision
0.0
0.1
0.2
0.3
0.4
0.5
0.6
Predicted Positives
0
500
1000
1500
2000
2 3 4 5 6 7 8 9 10 11
−log(q)

### Response to nematode. Poor

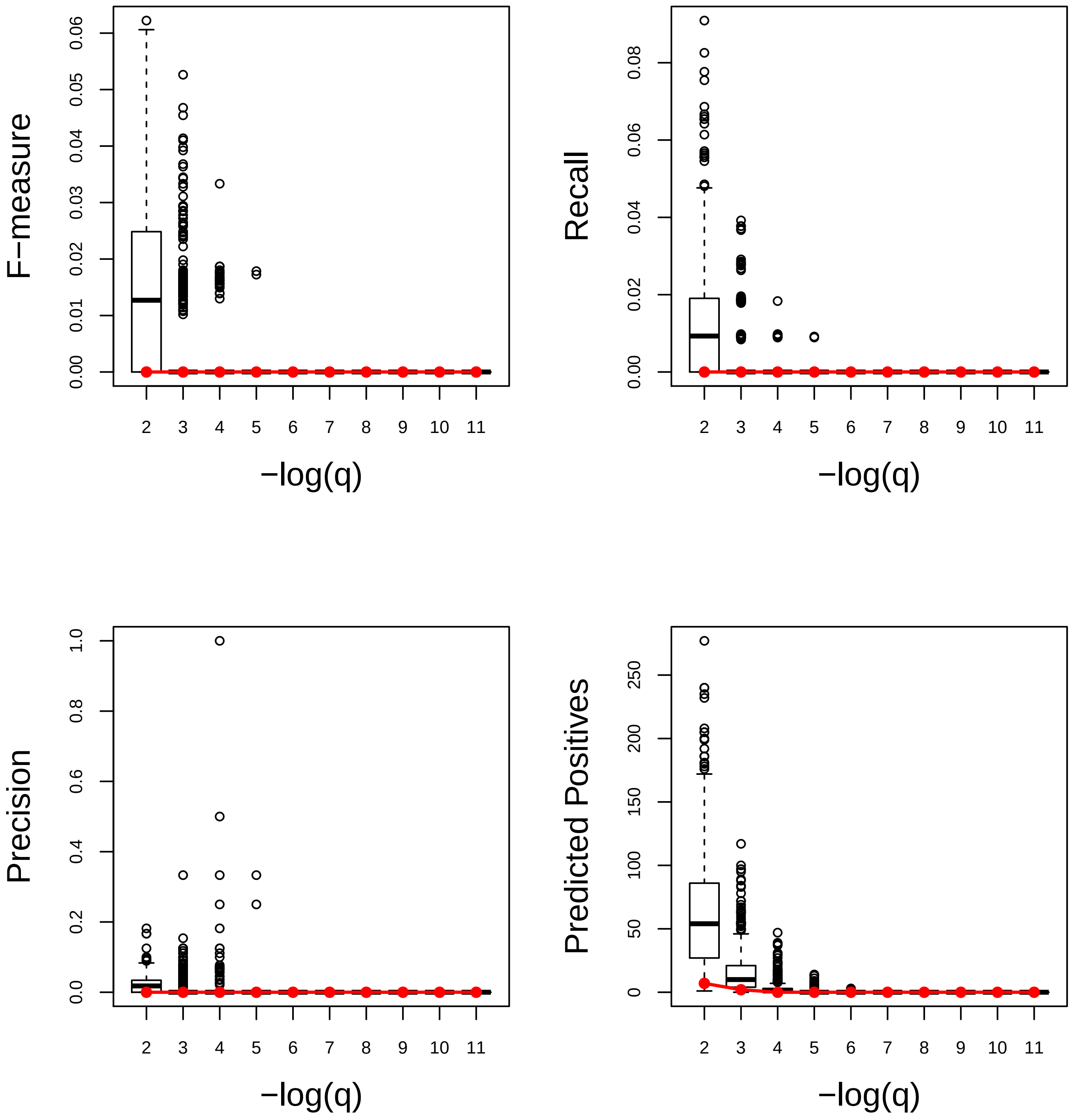

### Response to nitrogen compound. Very Good

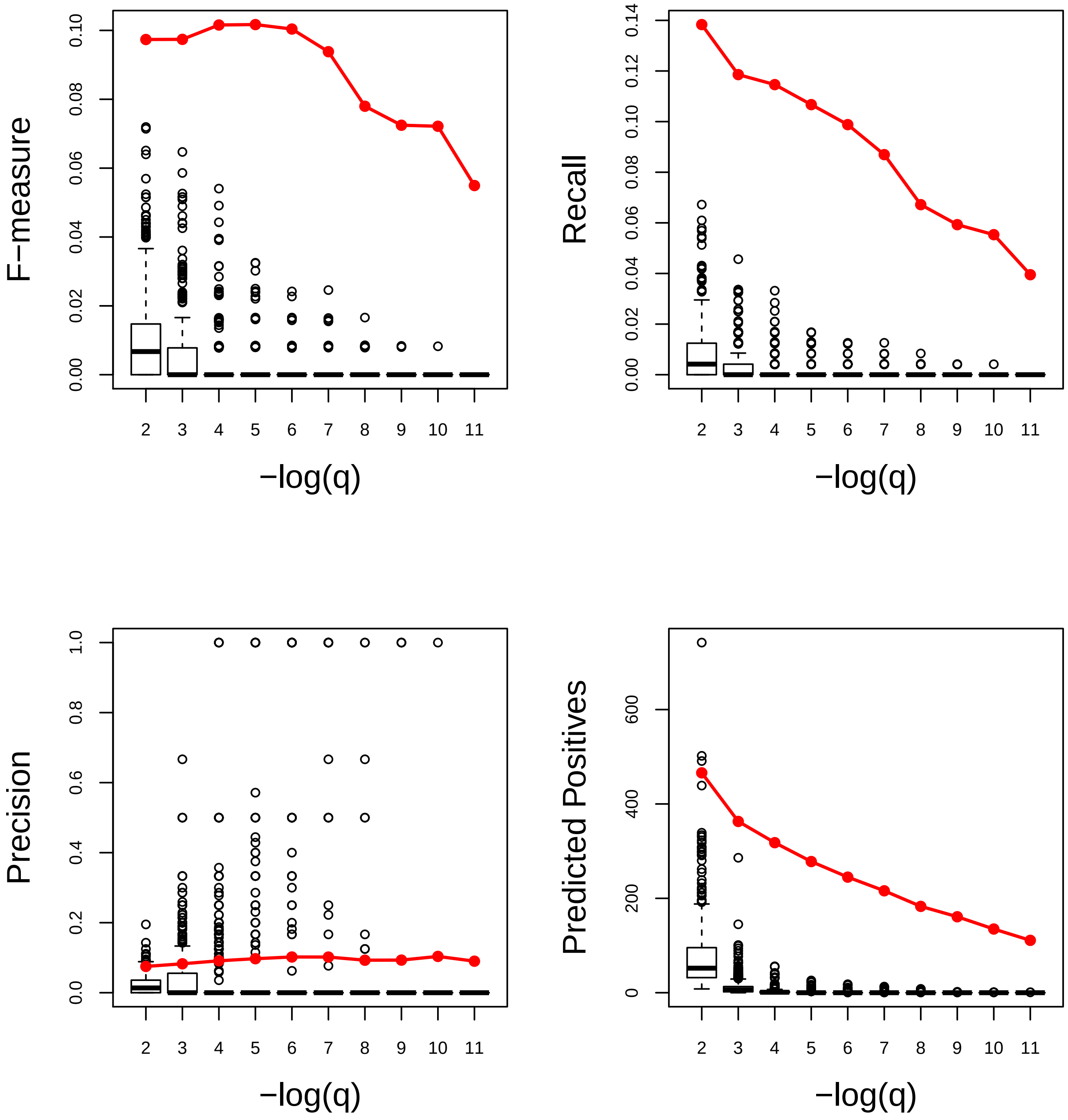

### Response to organonitrogen compound. Very Good

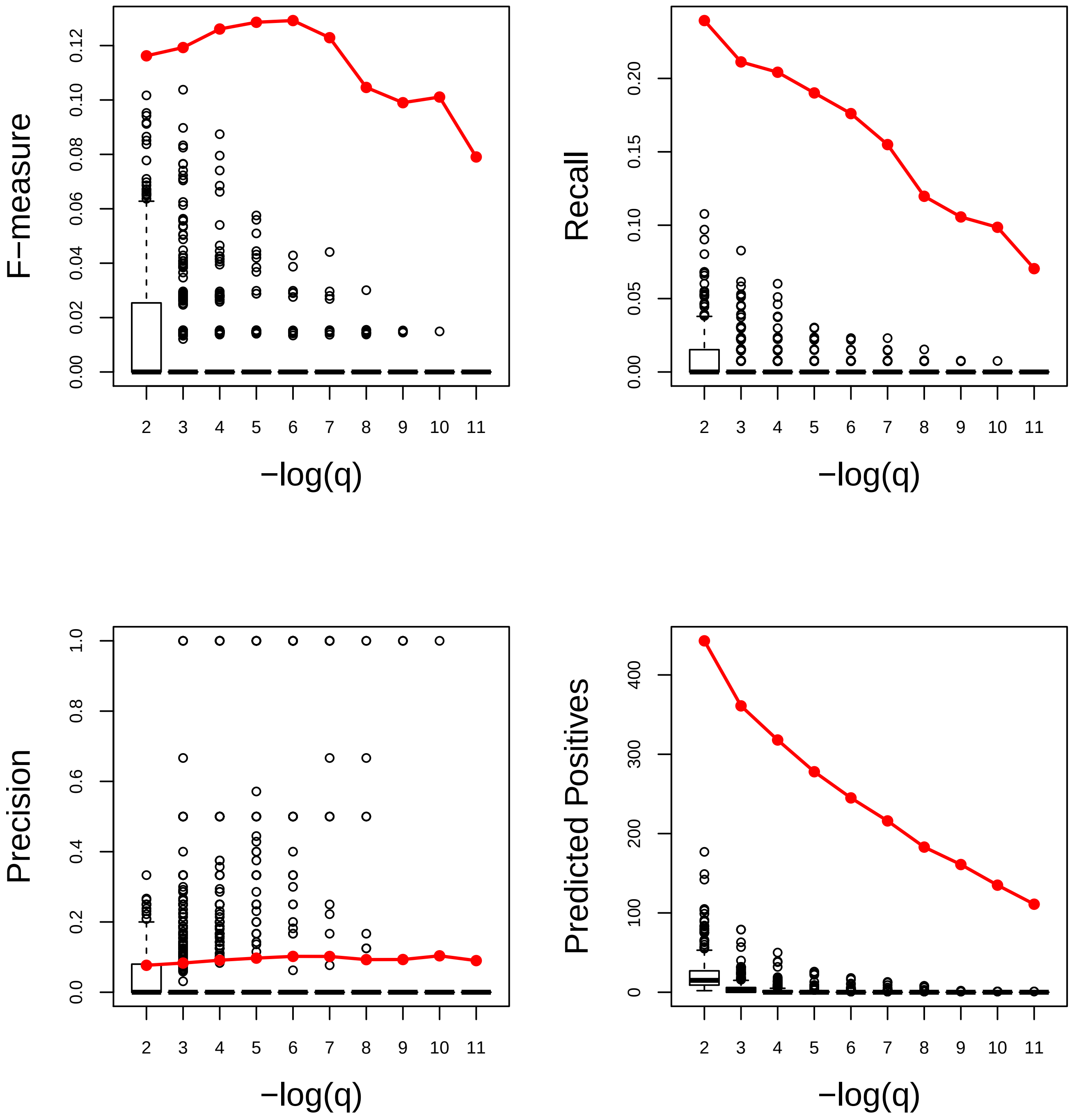

### Response to osmotic stress. Very Good

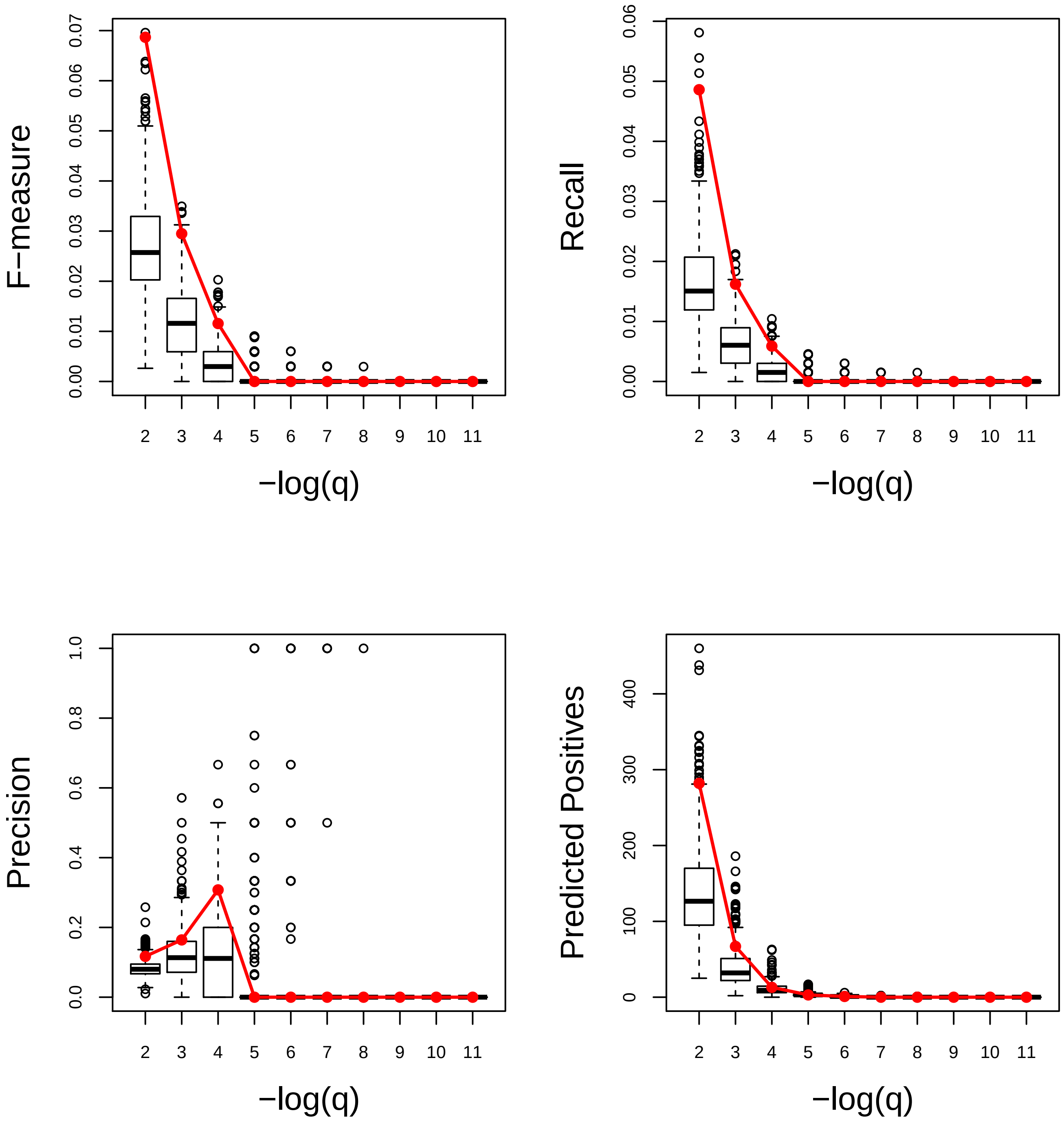

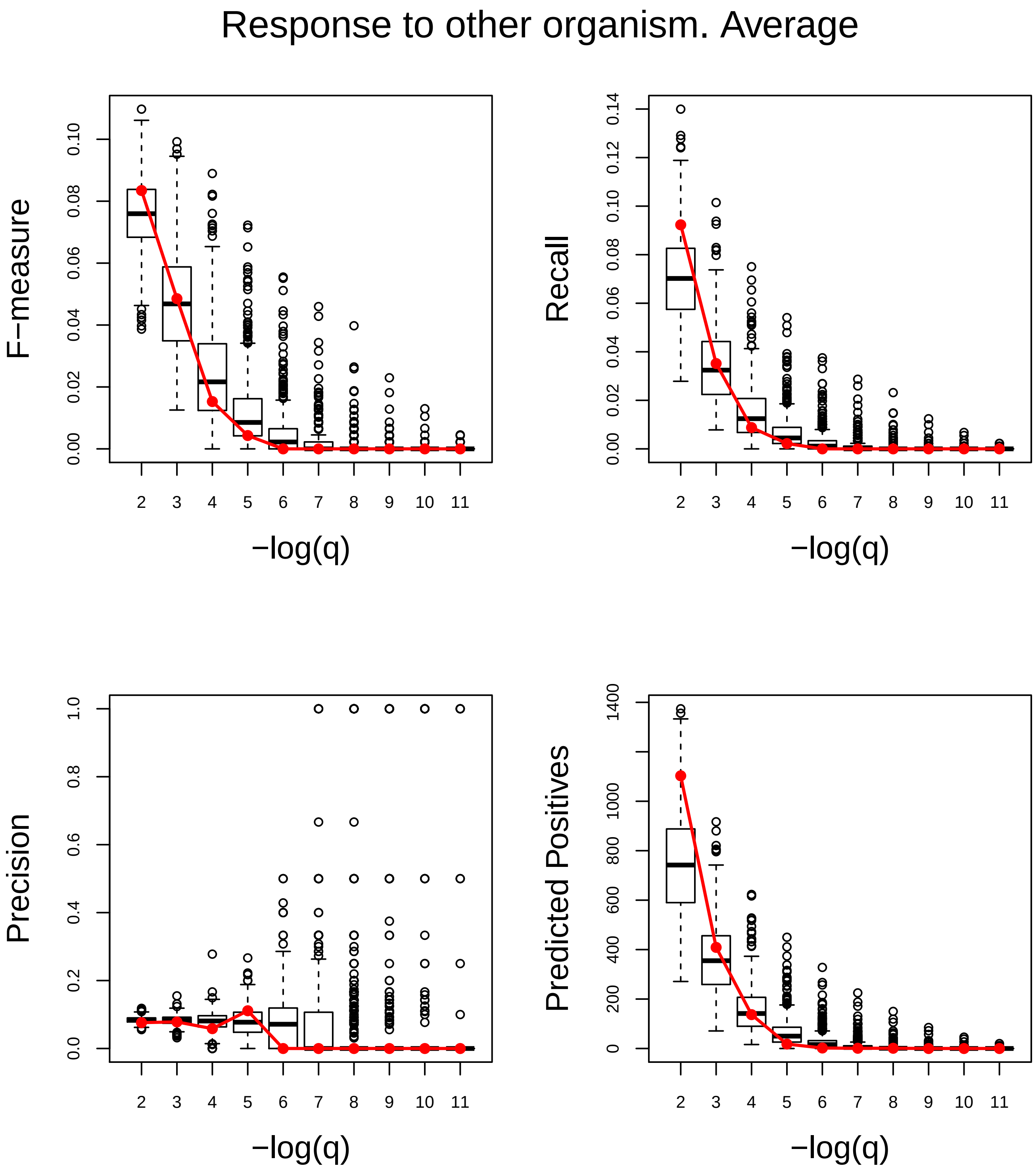
Response to other organism. Average
F−measure
0.00
0.02
0.04
0.06
0.08
0.10
Recall
0.00
0.02
0.04
0.06
0.08
0.10
0.12
0.14
Precision
0.0
0.2
0.4
0.6
0.8
1.0
Predicted Positives
0
200
400
600
800
1000
1400
2 3 4 5 6 7 8 9 10 11
−log(q)

### Response to oxidative stress. Good

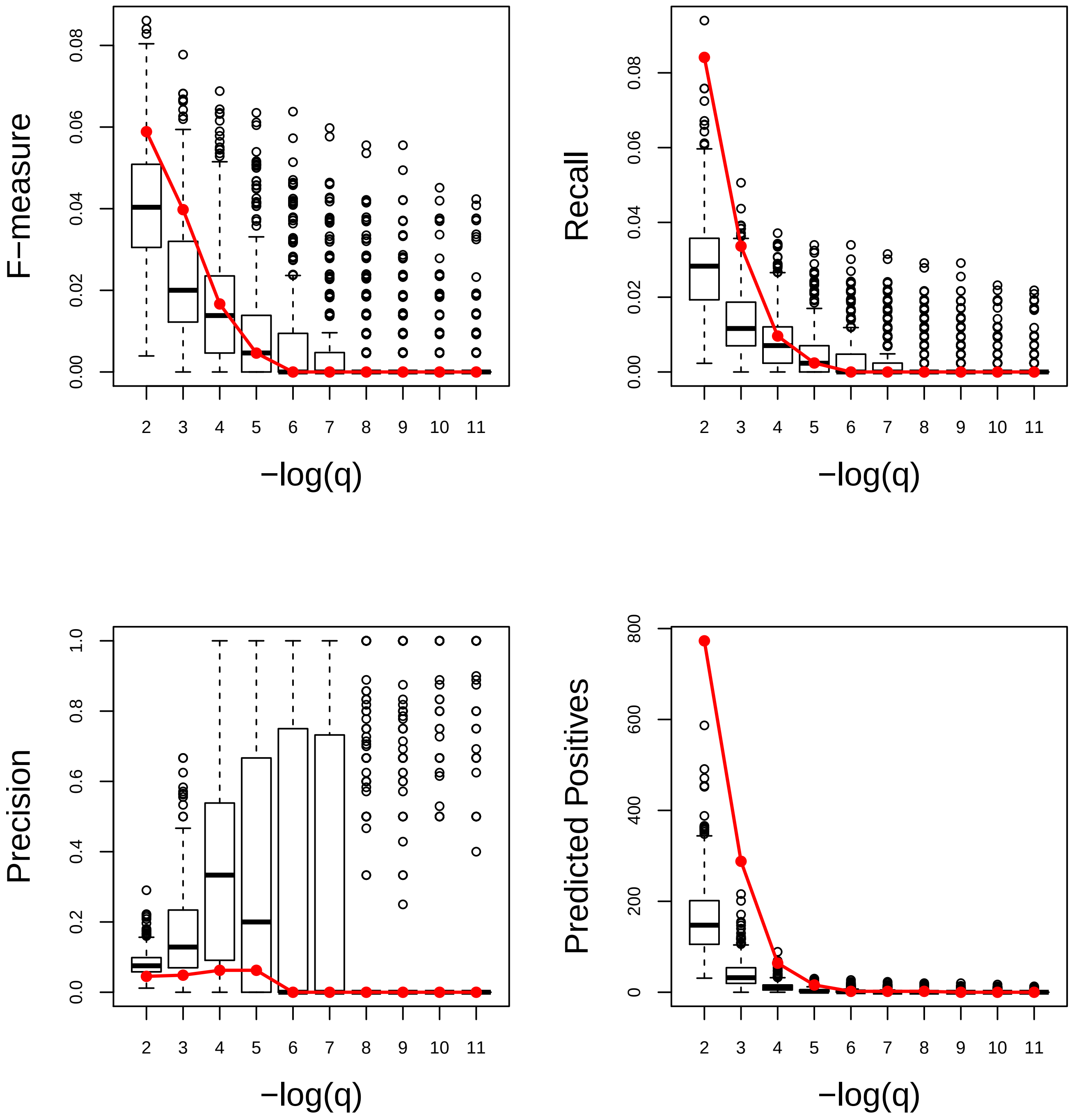

### Response to oxygen−containing compound. Very Good

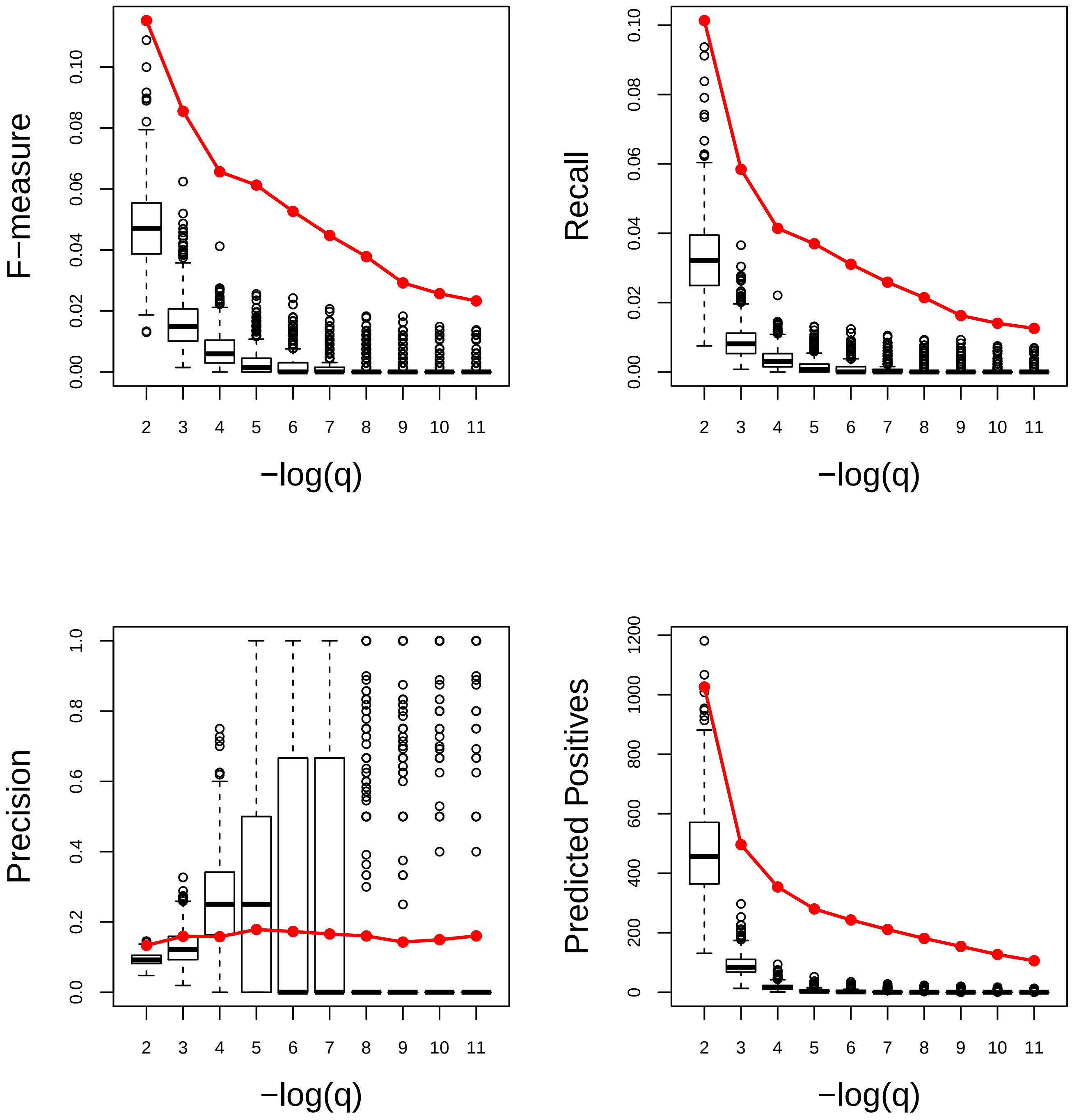

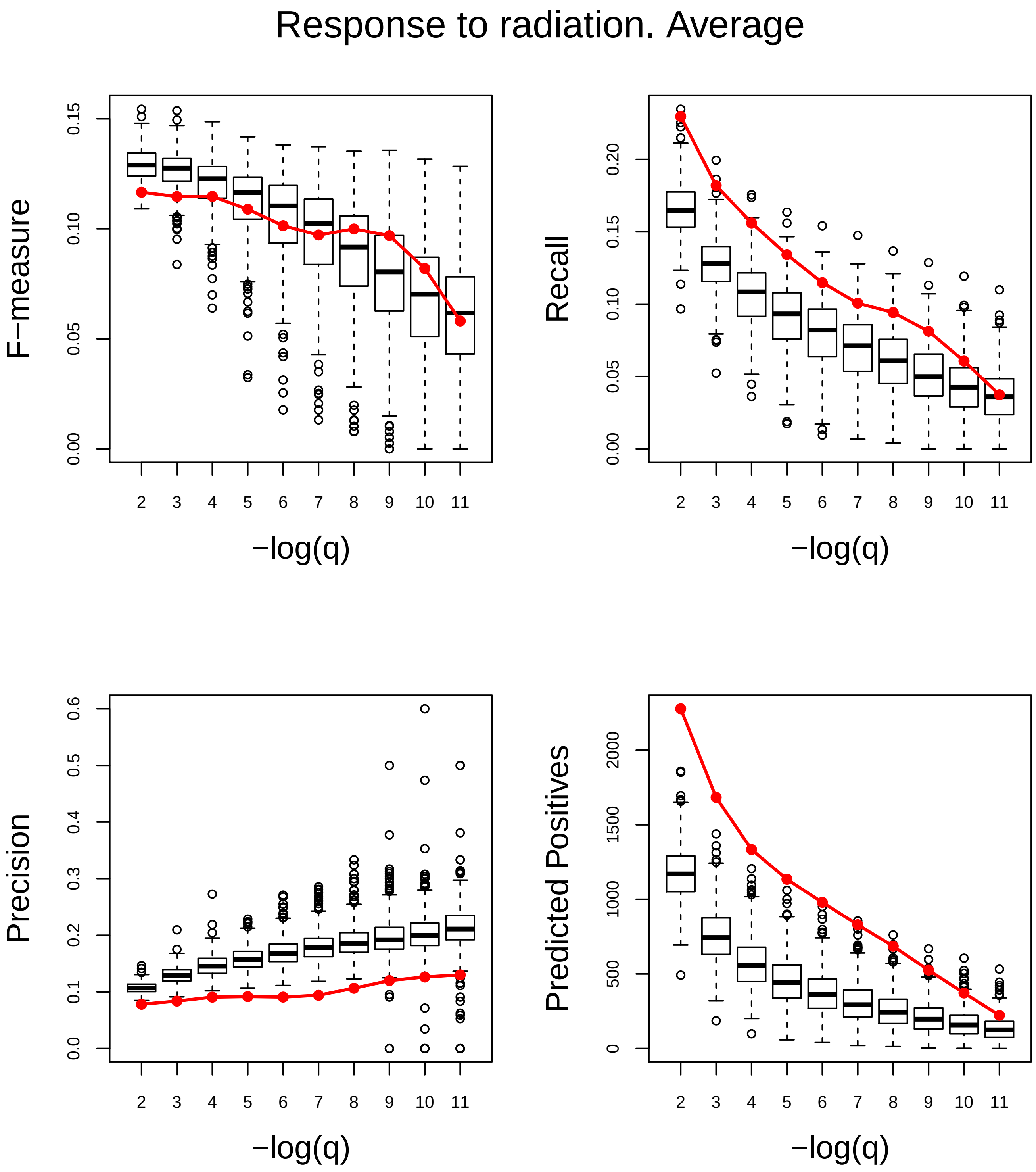
Response to radiation. Average
F−measure
0.00
0.05
0.10
0.15
Recall
0.00
0.05
0.10
0.15
0.20
Precision
0.0
0.1
0.2
0.3
0.4
0.5
0.6
Predicted Positives
0
500
1000
1500
2000
2 3 4 5 6 7 8 9 10 11
−log(q)

### Response to reactive oxygen species. Poor

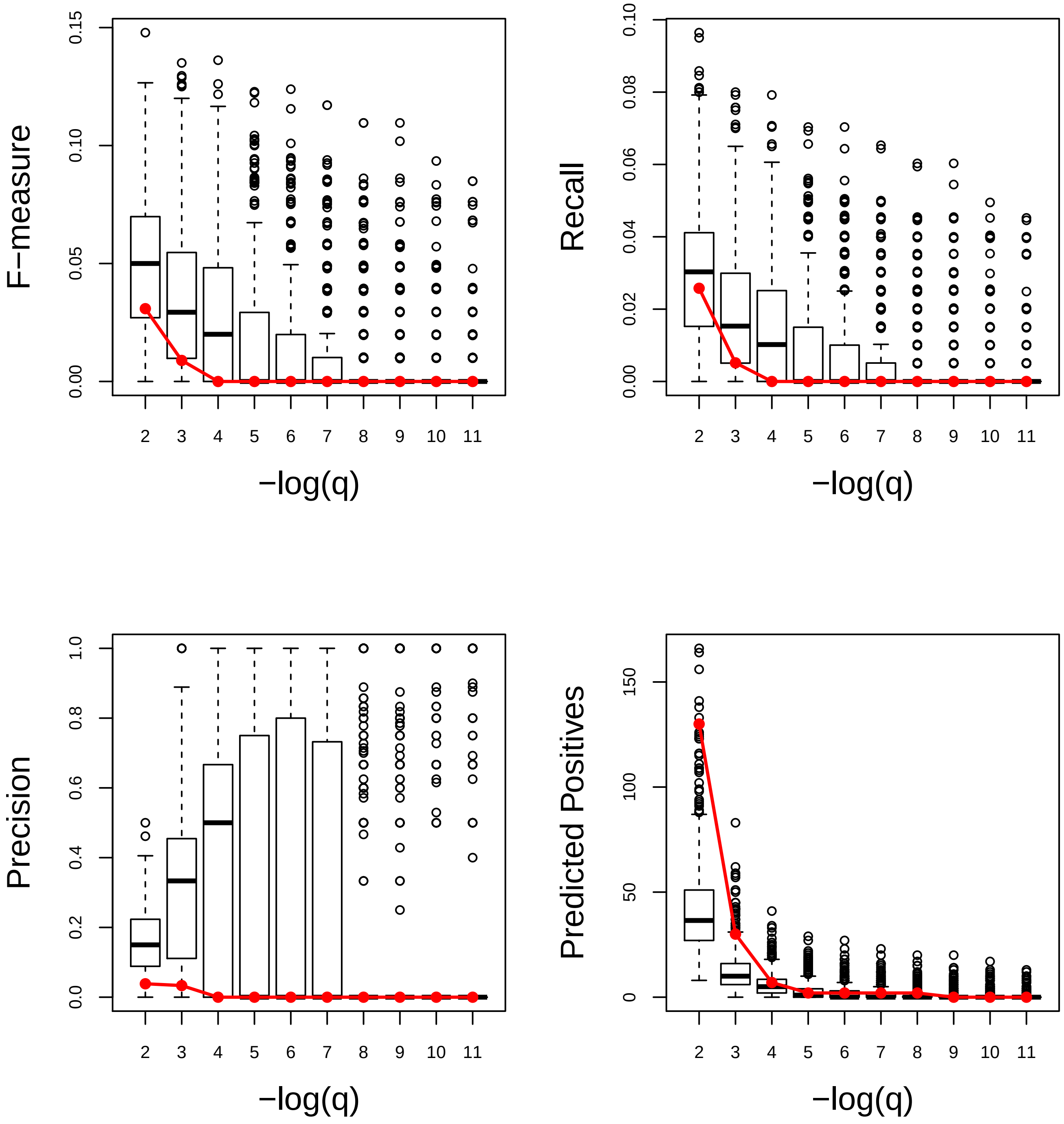

### Response to red or far red light. Poor

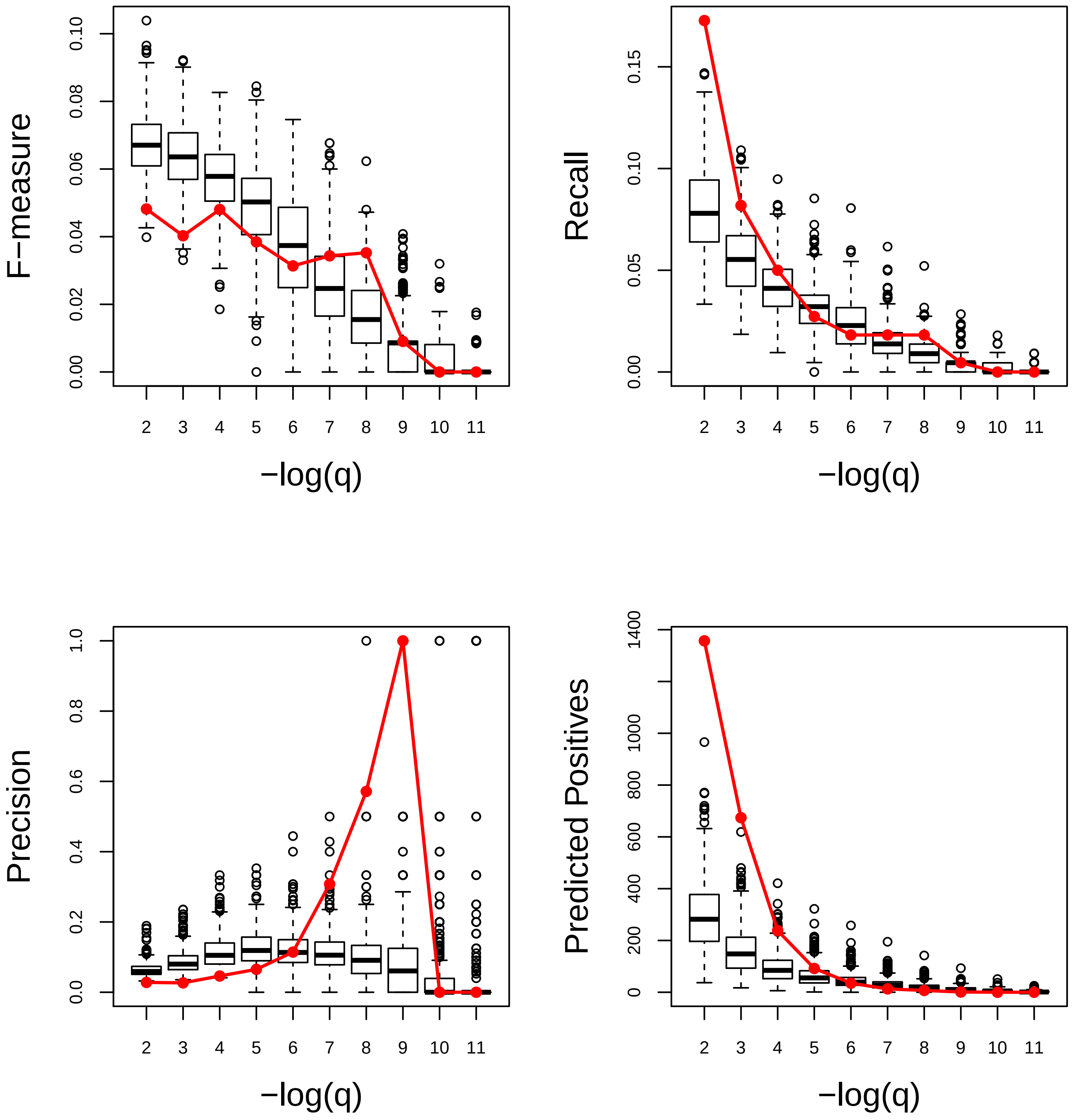

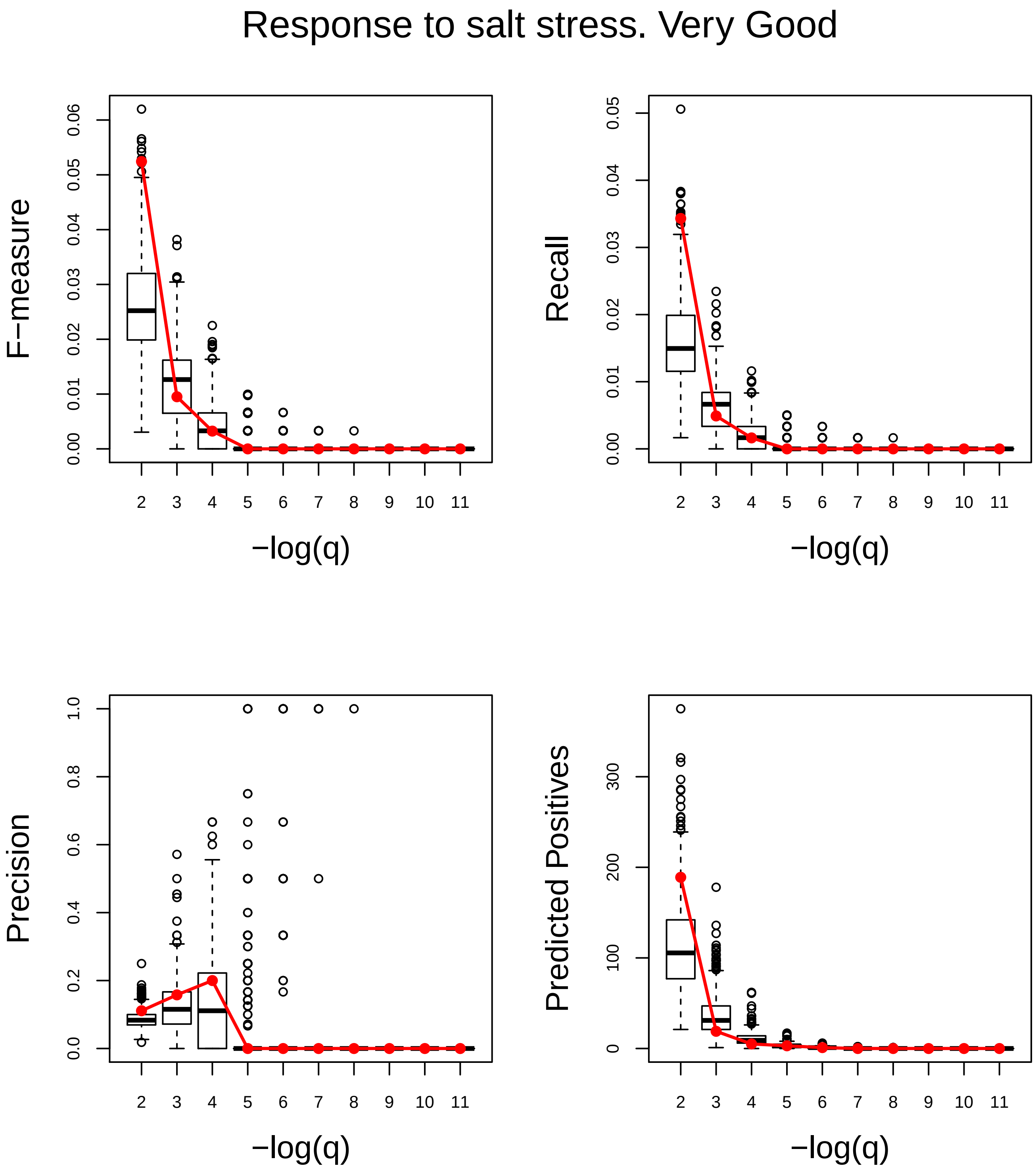
Response to salt stress. Very Good
F−measure
Recall
Precision
Predicted Positives
−log(q)
−log(q)
−log(q)
−log(q)

### Response to starvation. Very Good

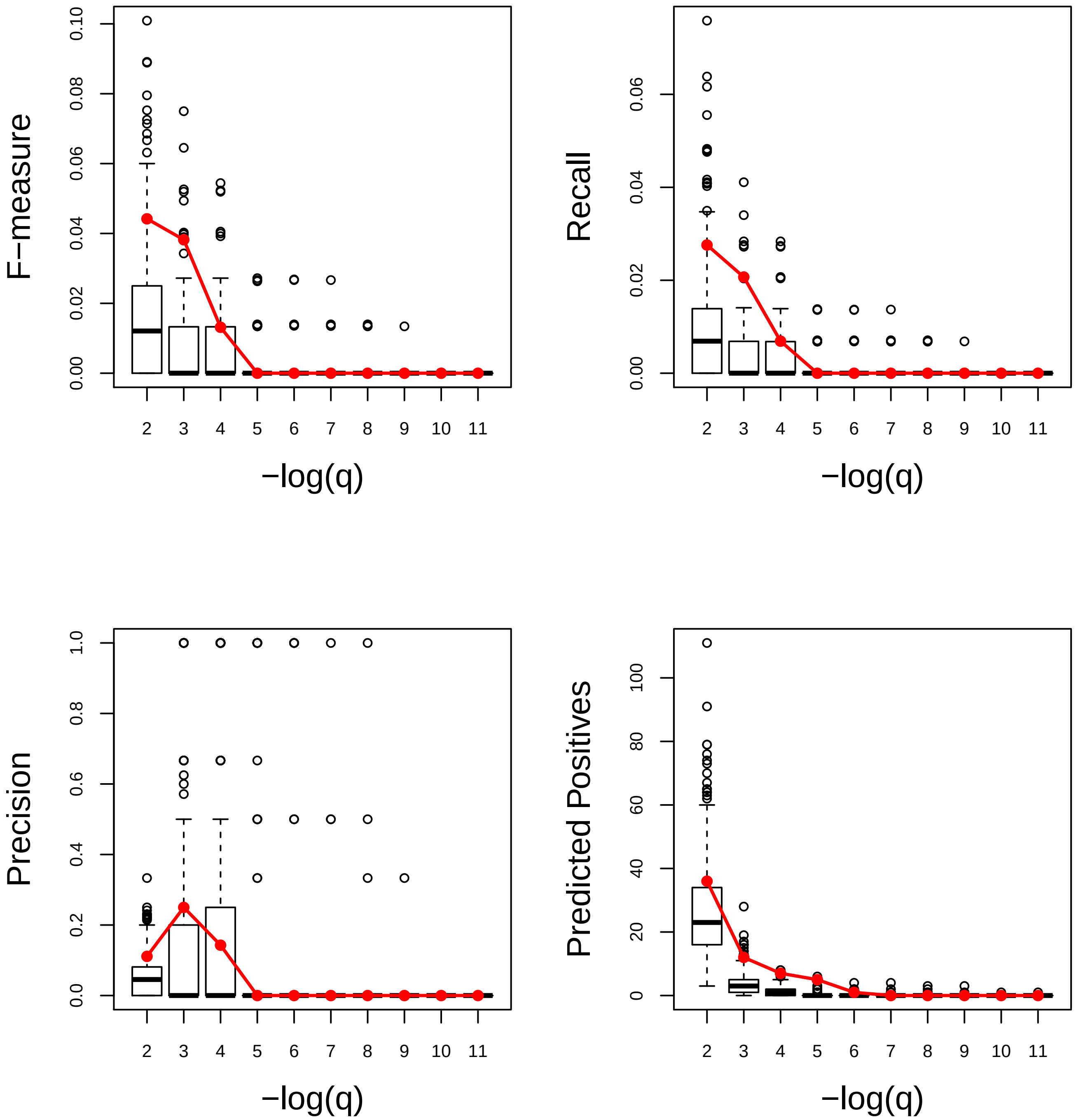

### Response to stress. Very Good

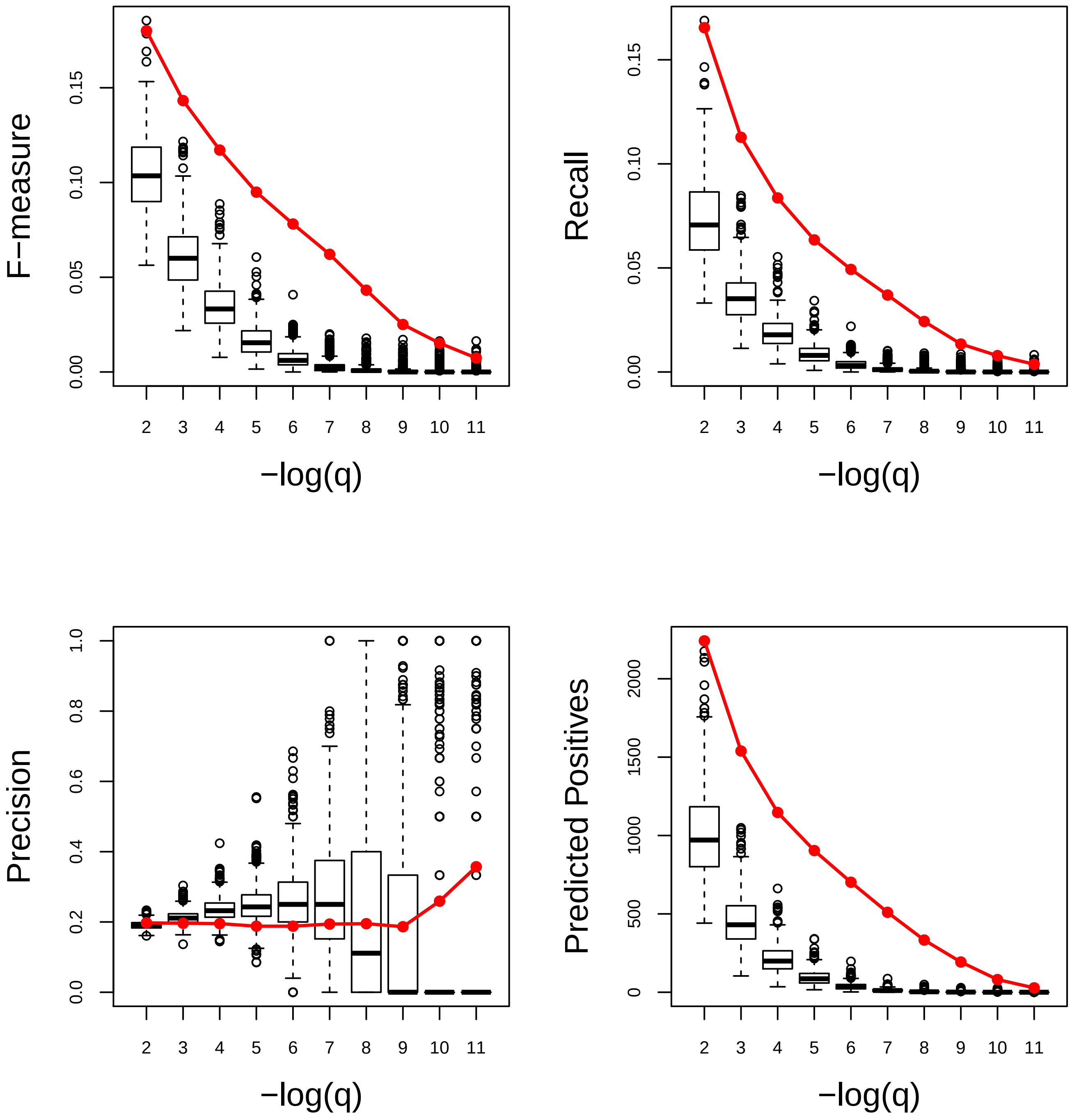

### Response to symbiont. Poor

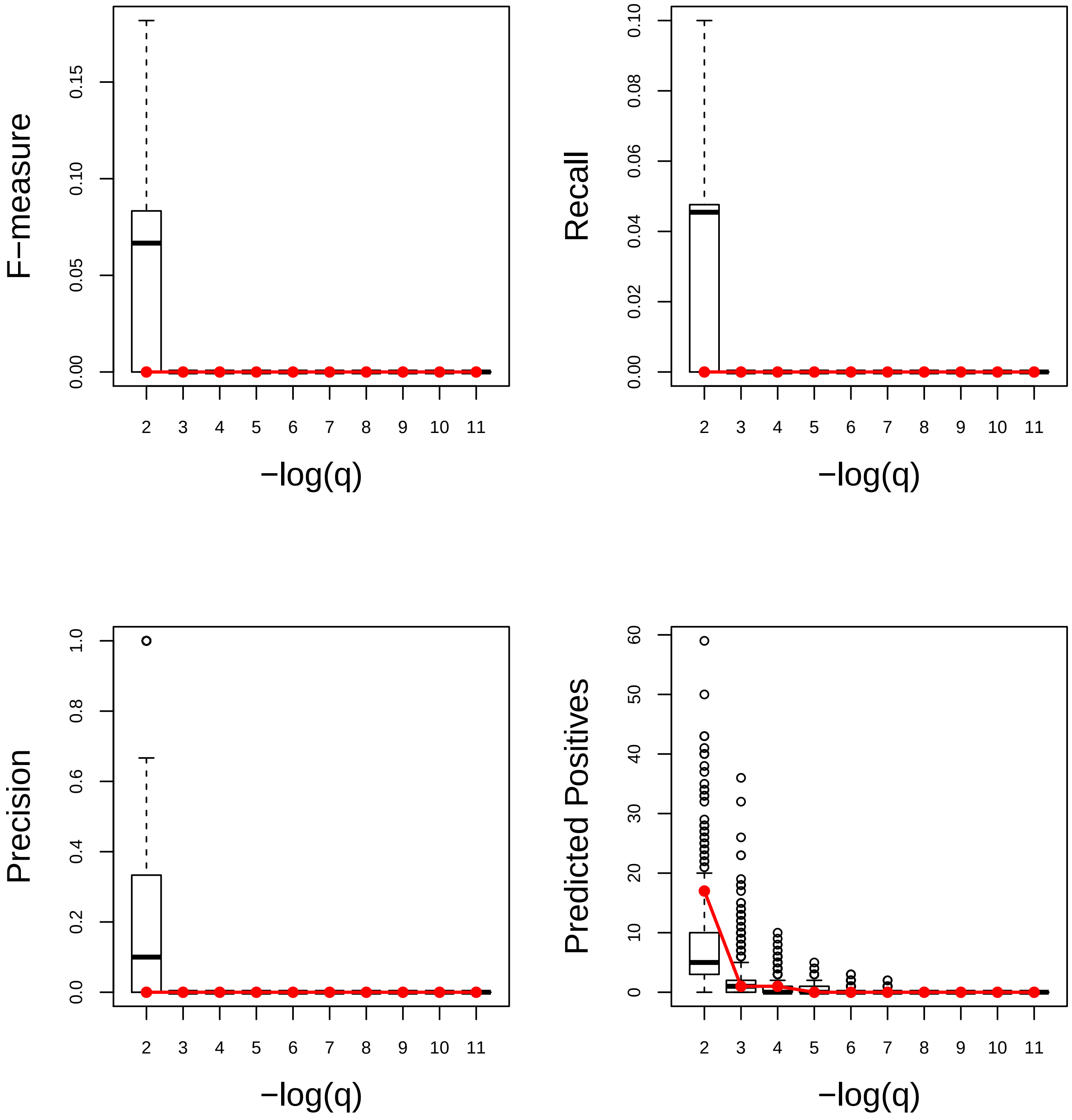

### Response to temperature stimulus. Very Good

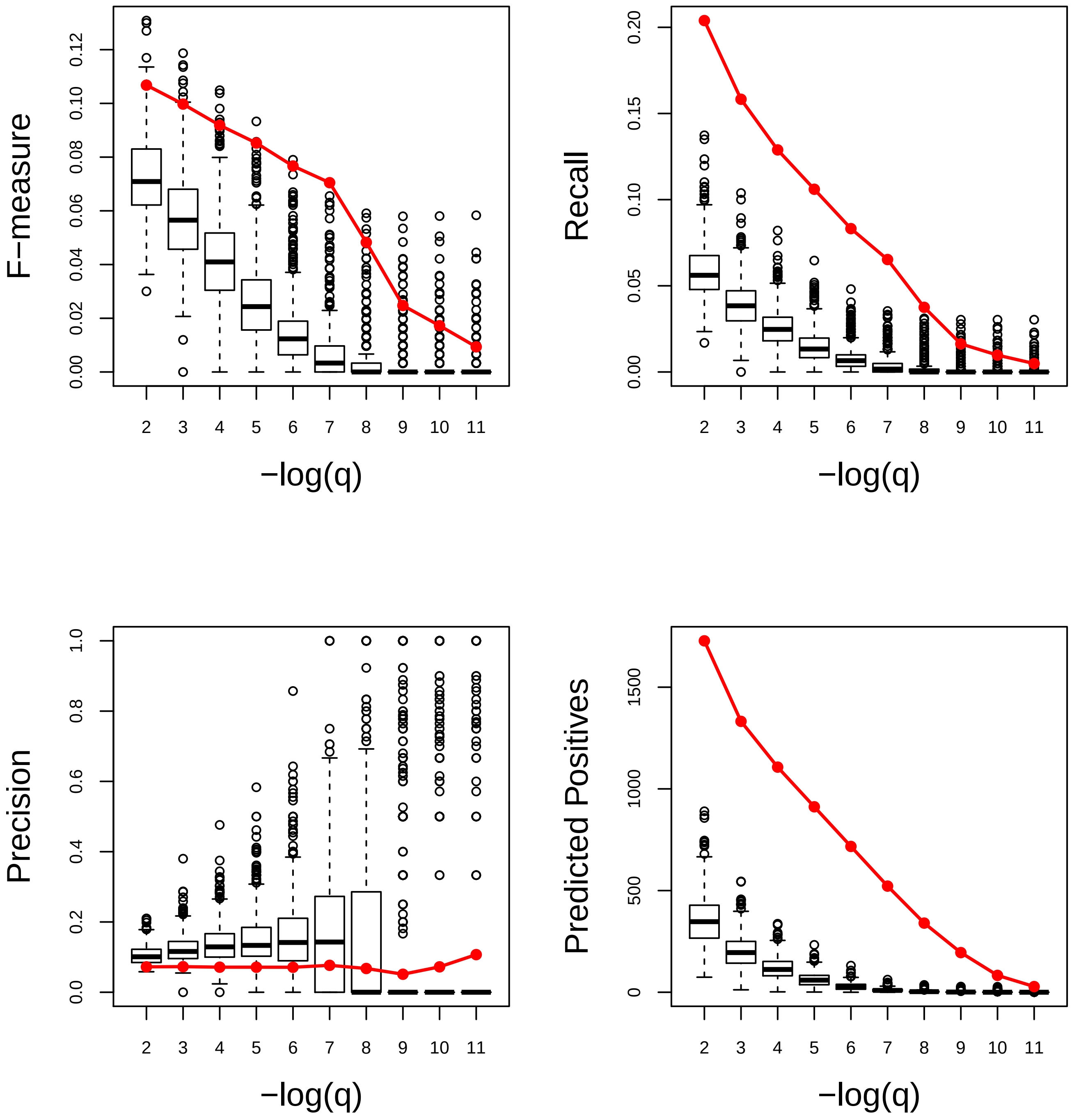

### Response to UV. Very Good

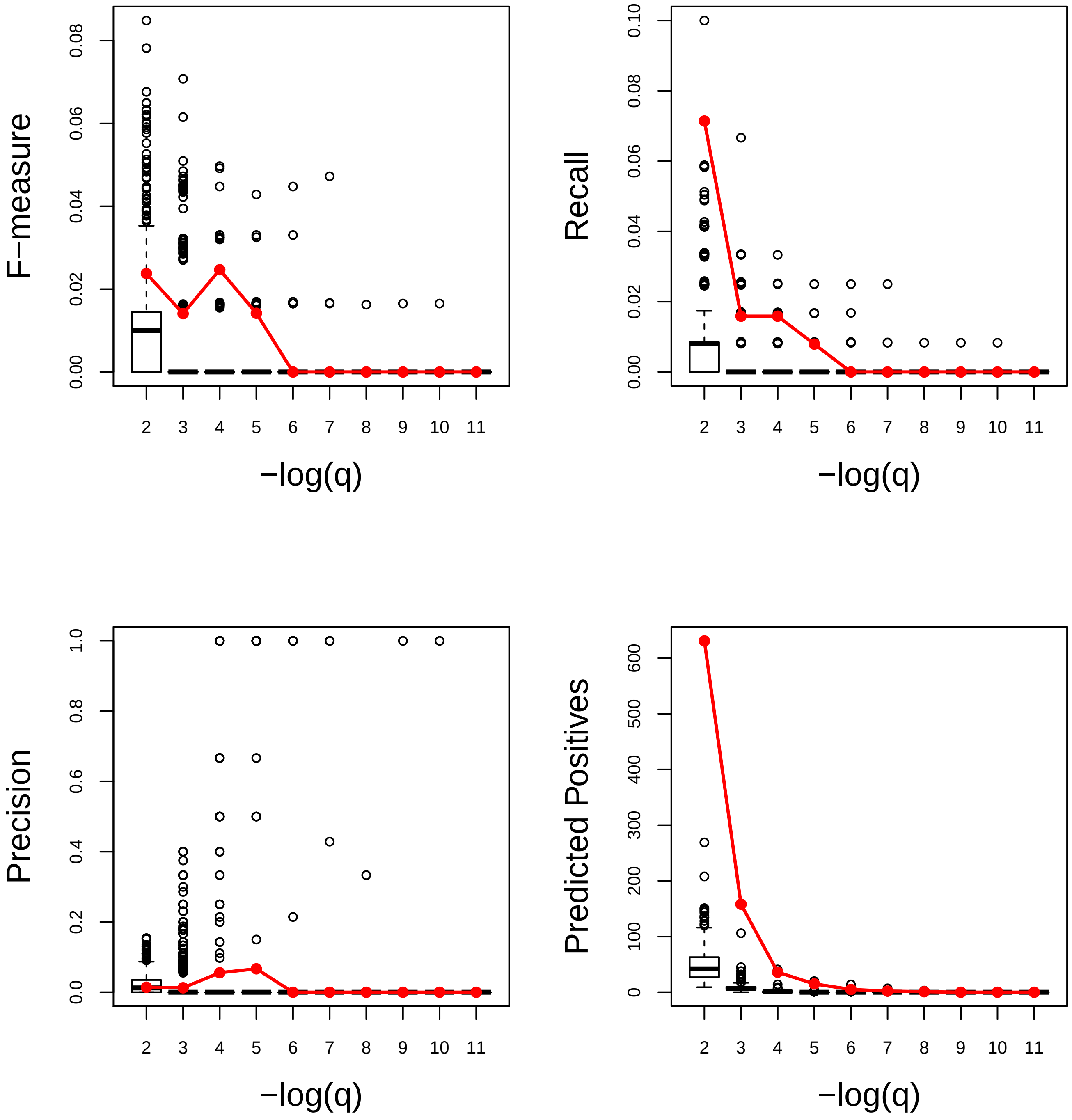

### Response to water deprivation. Very Good

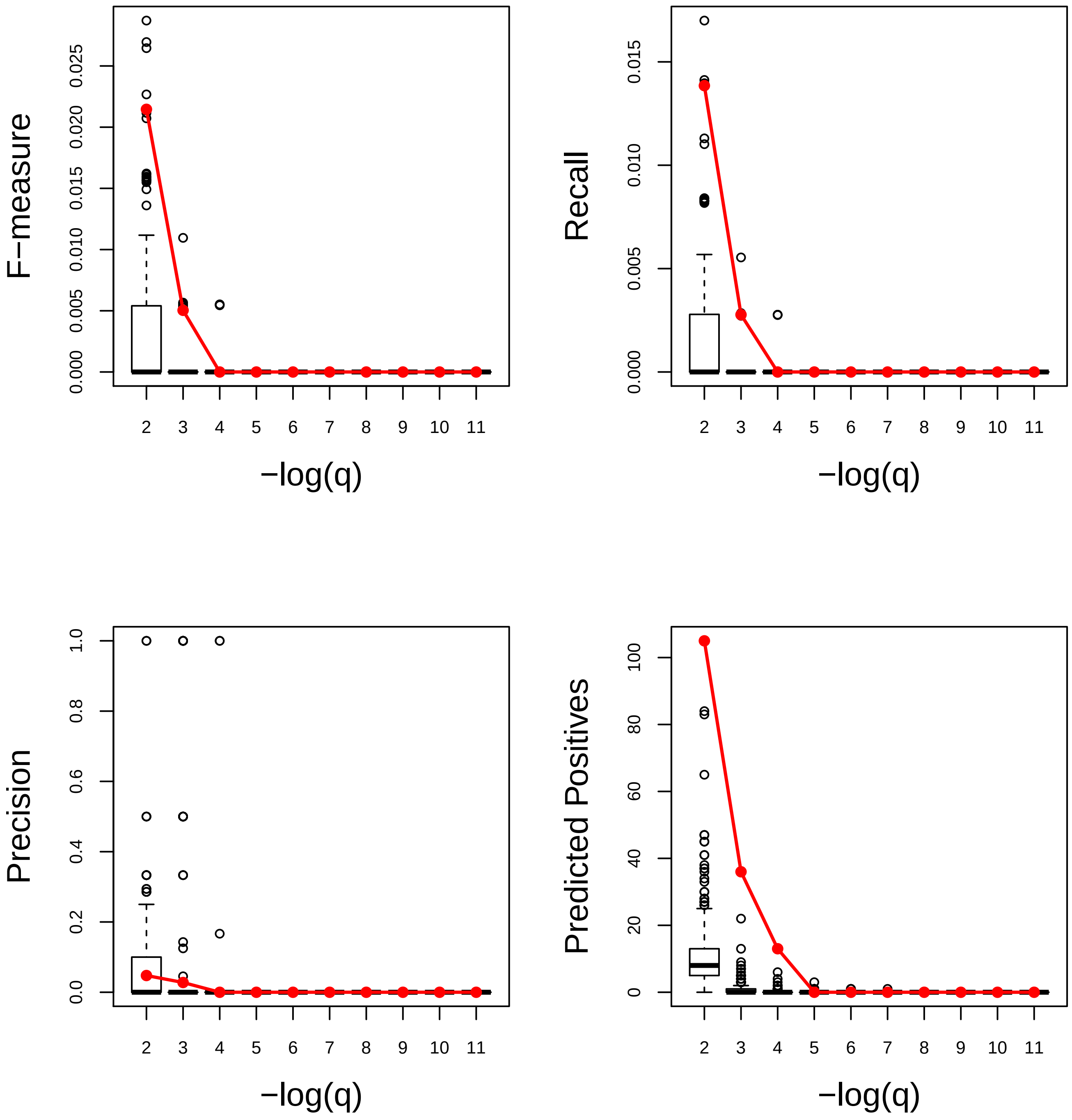

### Response to water. Very Good

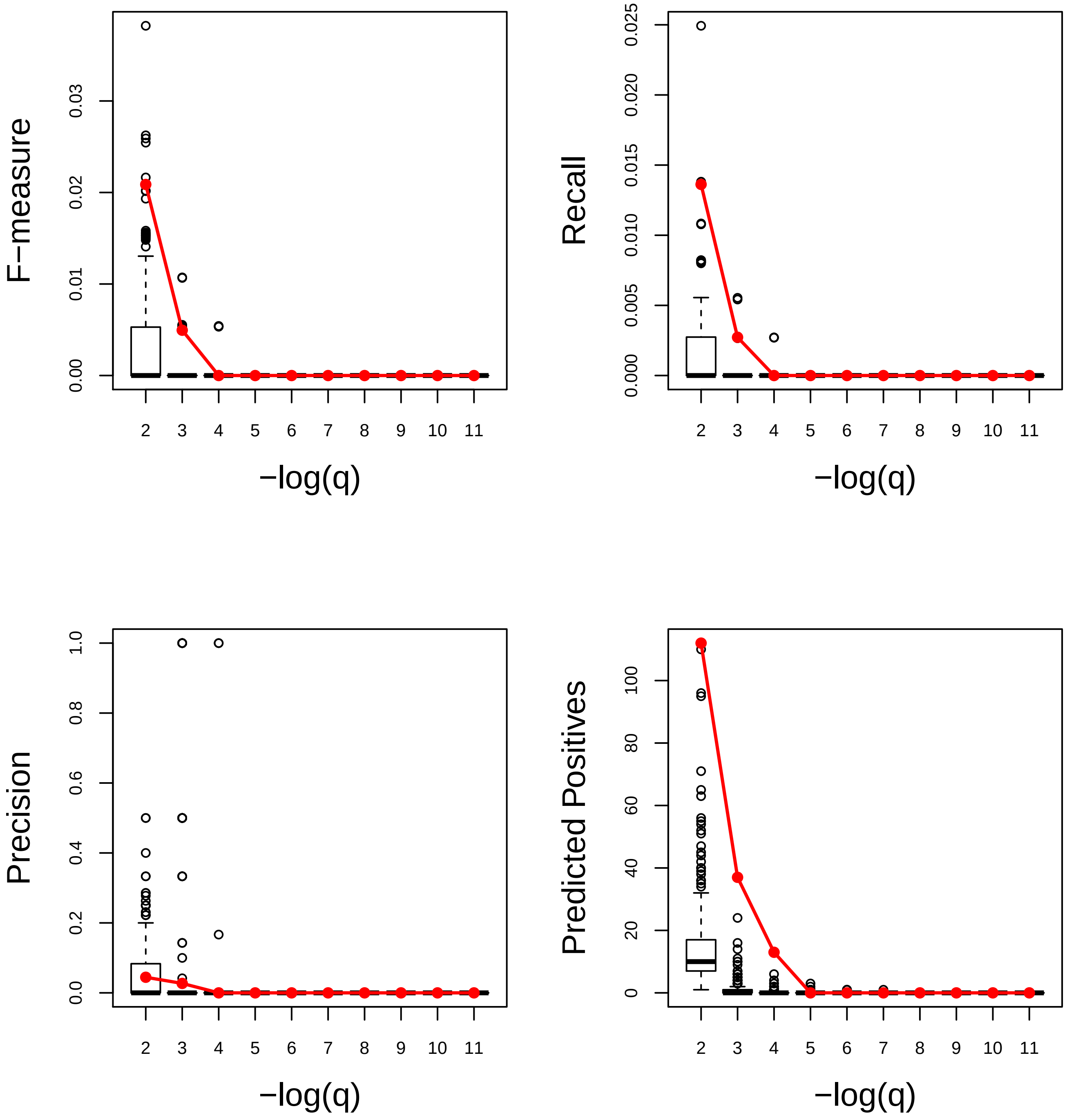

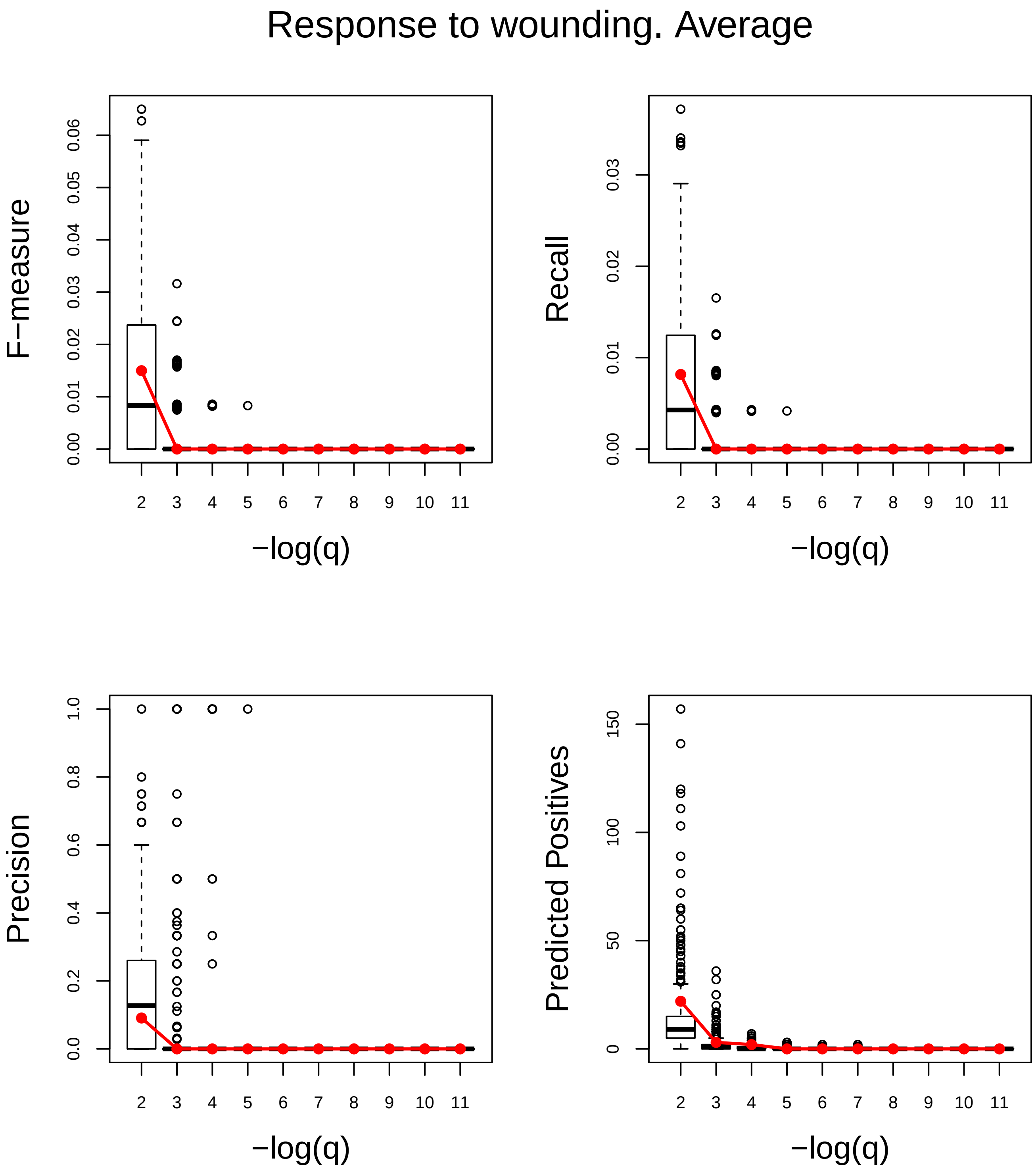
Response to wounding. Average
F−measure
0.00
0.01
0.02
0.03
0.04
0.05
0.06
Recall
0.00
0.01
0.02
0.03
Precision
0.0
0.2
0.4
0.6
0.8
1.0
Predicted Positives
0
50
100
150
2 3 4 5 6 7 8 9 10 11
−log(q)

### Root development. Good

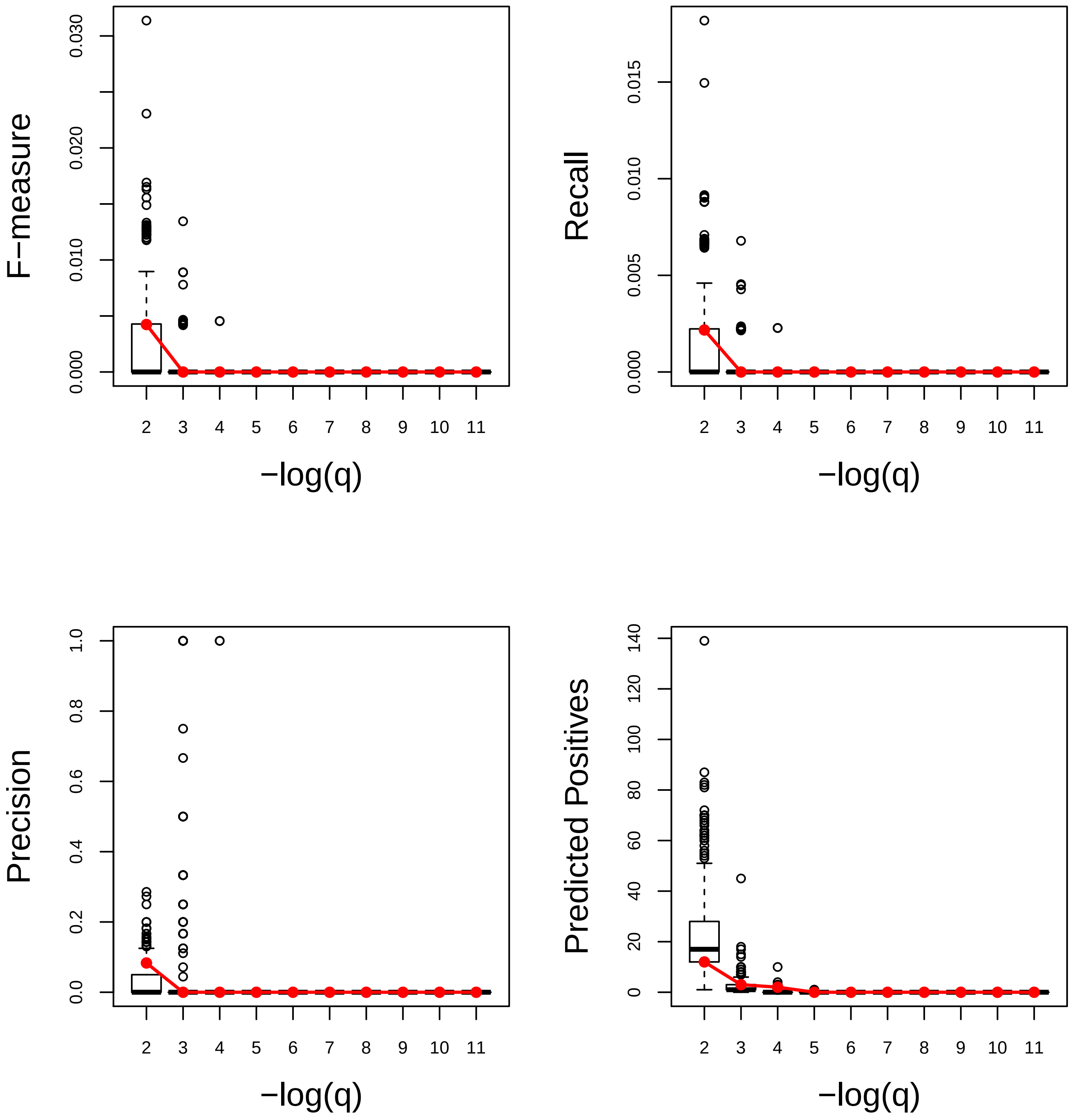

### Secondary metabolic process. Very Good

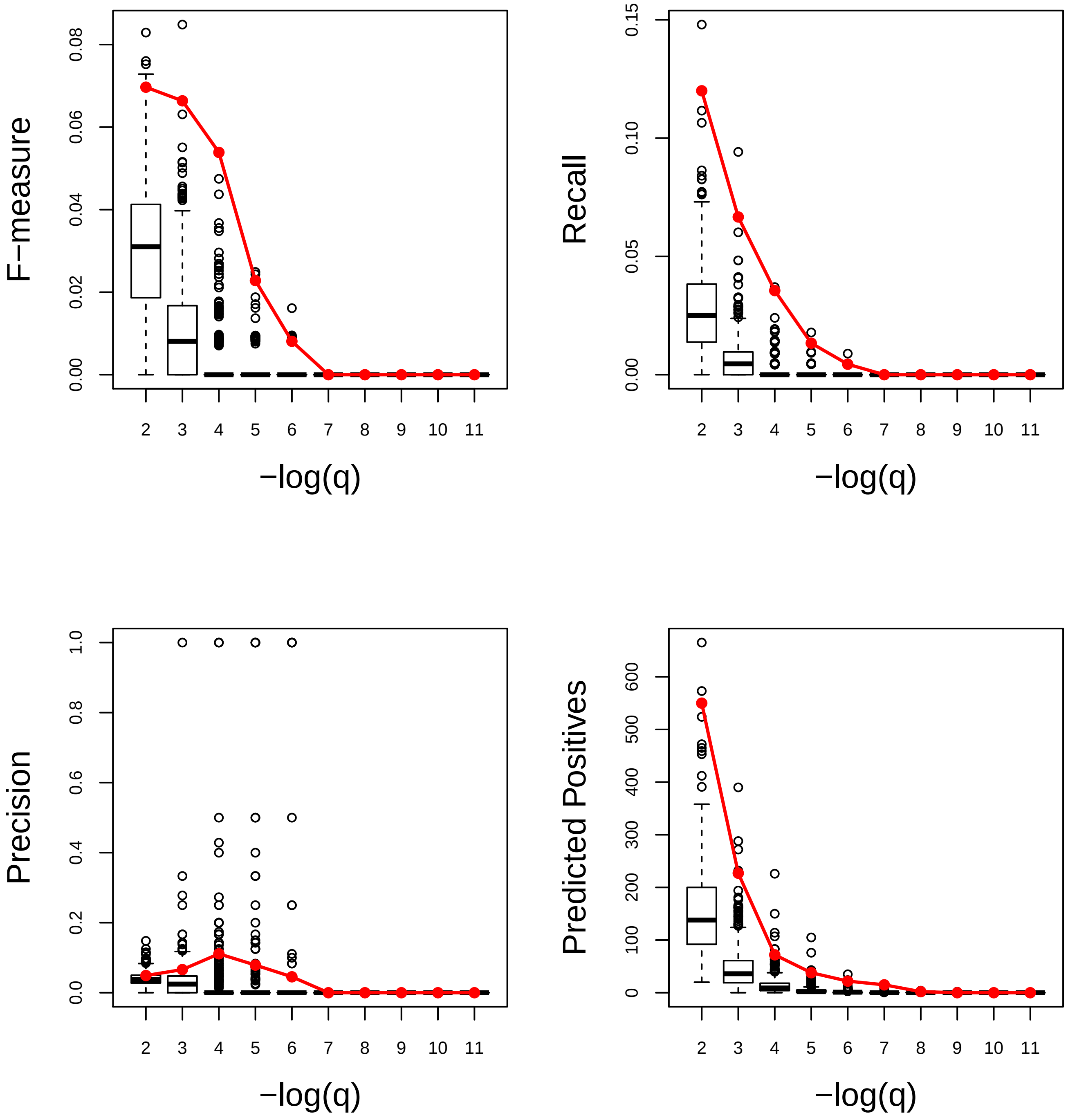

### Seed development. Average

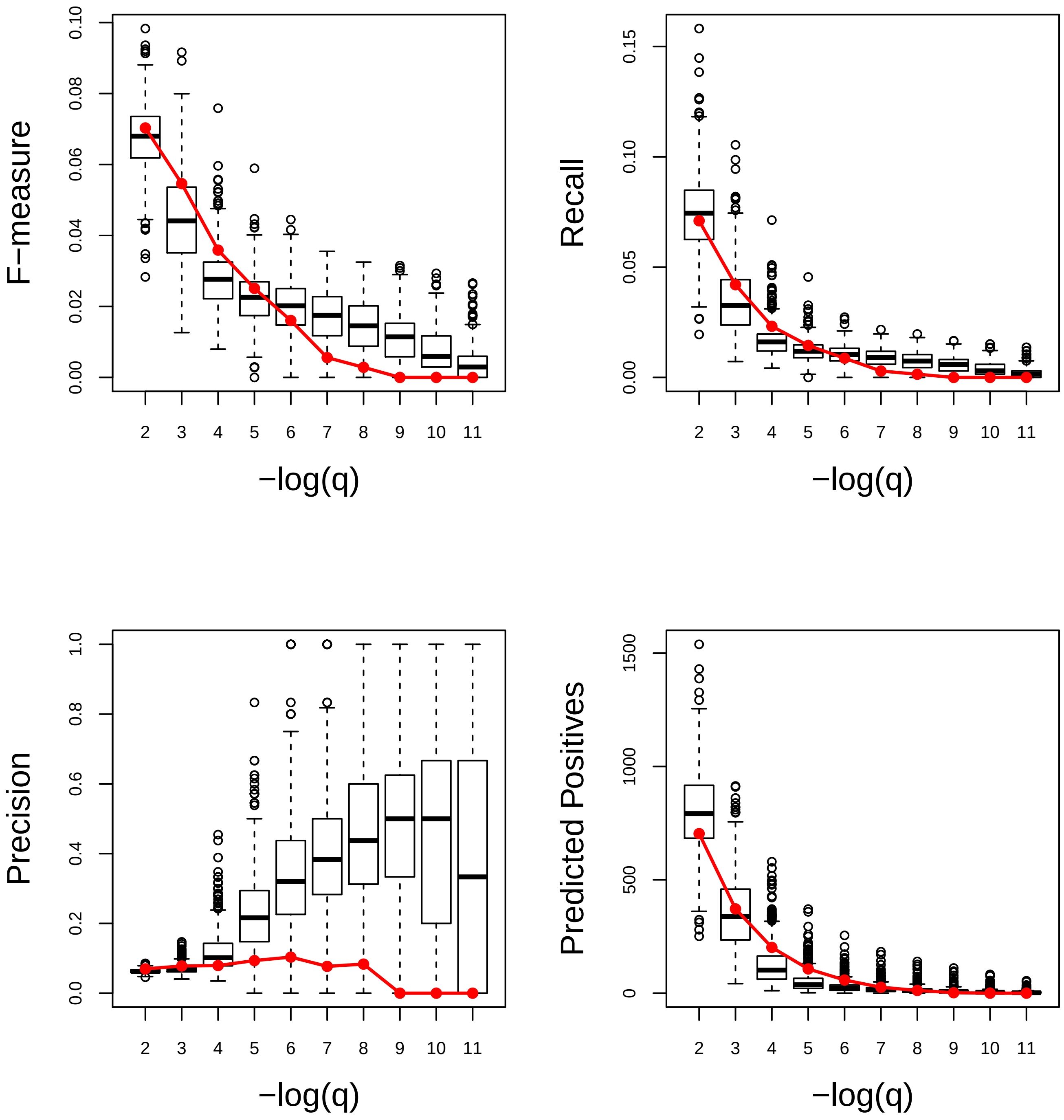

### Shoot system development. Very Poor

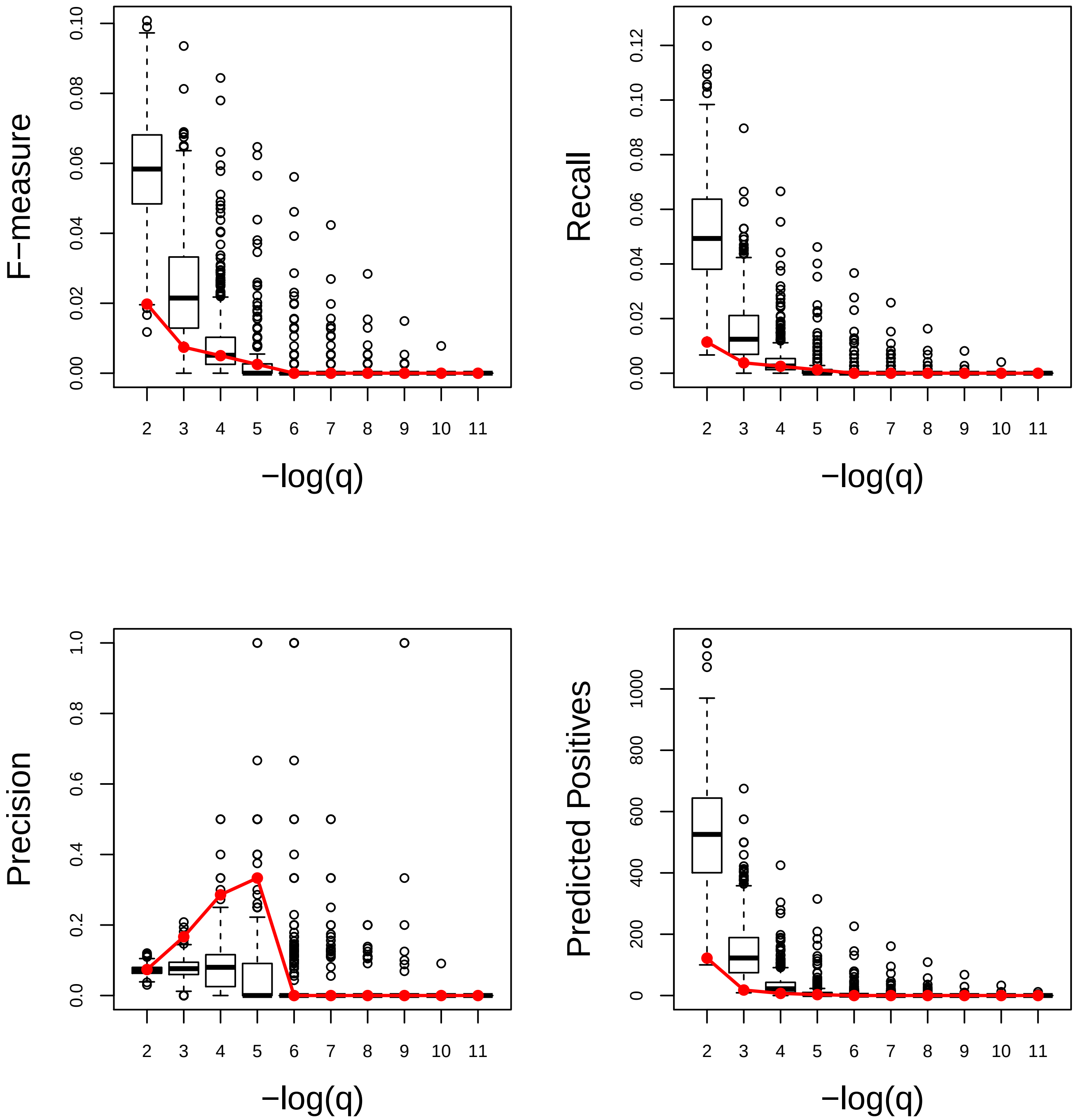

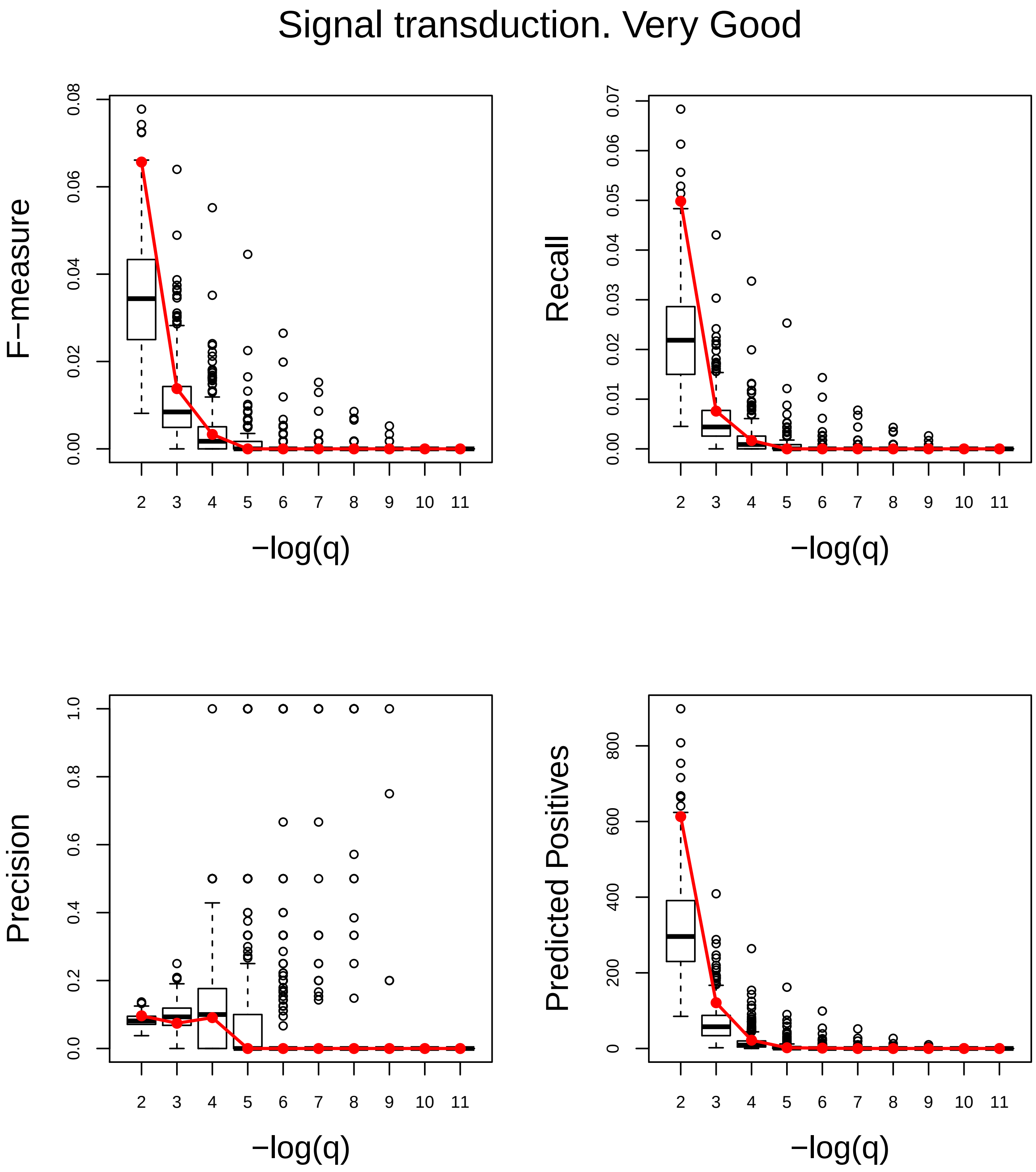
Signal transduction. Very Good
F−measure
0.00
0.02
0.04
0.06
0.08
Recall
0.00
0.01
0.02
0.03
0.04
0.05
0.06
0.07
Precision
0.0
0.2
0.4
0.6
0.8
1.0
Predicted Positives
0
200
400
600
800
2 3 4 5 6 7 8 9 10 11
−log(q)

### Translation. Very Poor

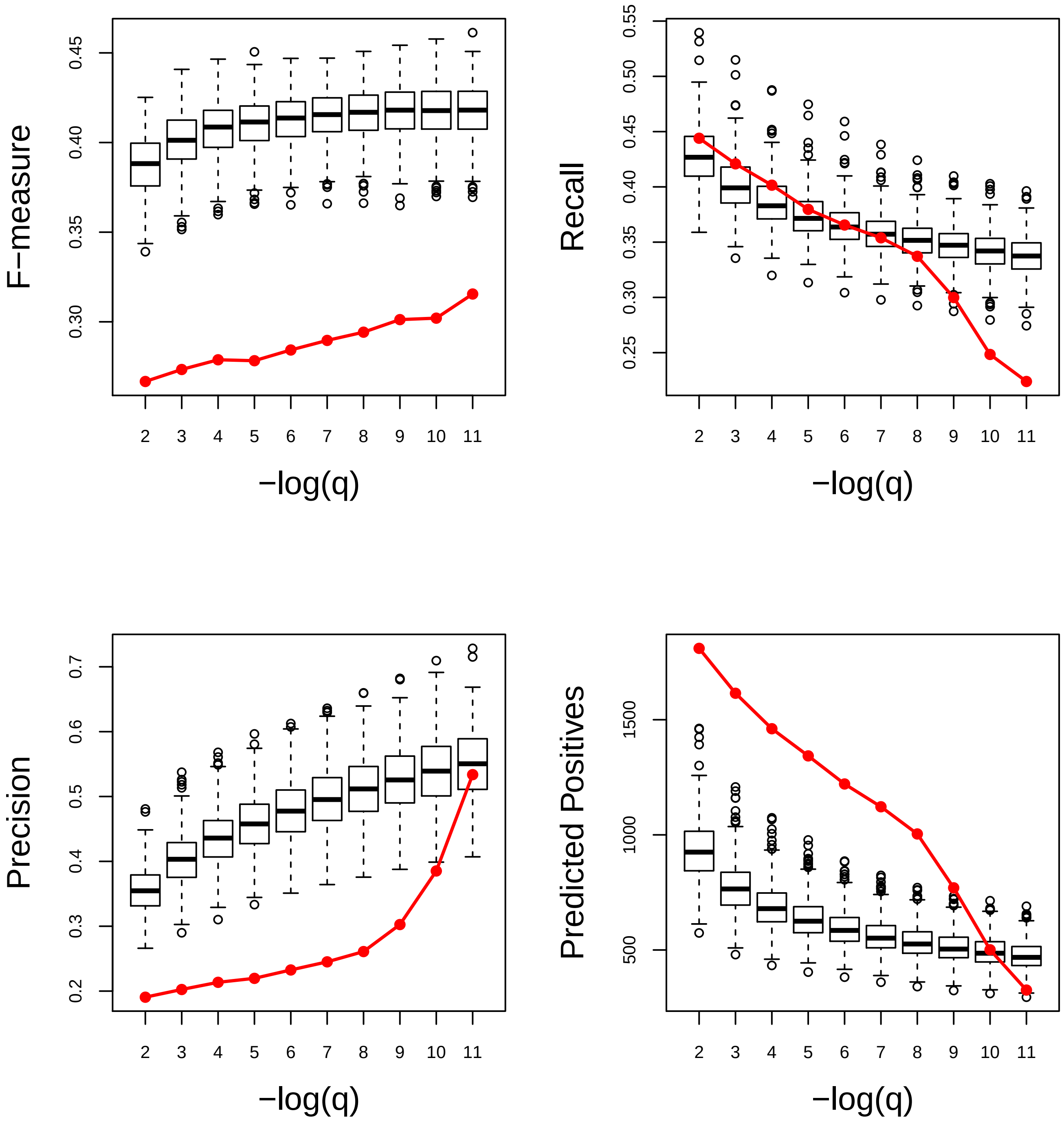

### Transport. Very Poor

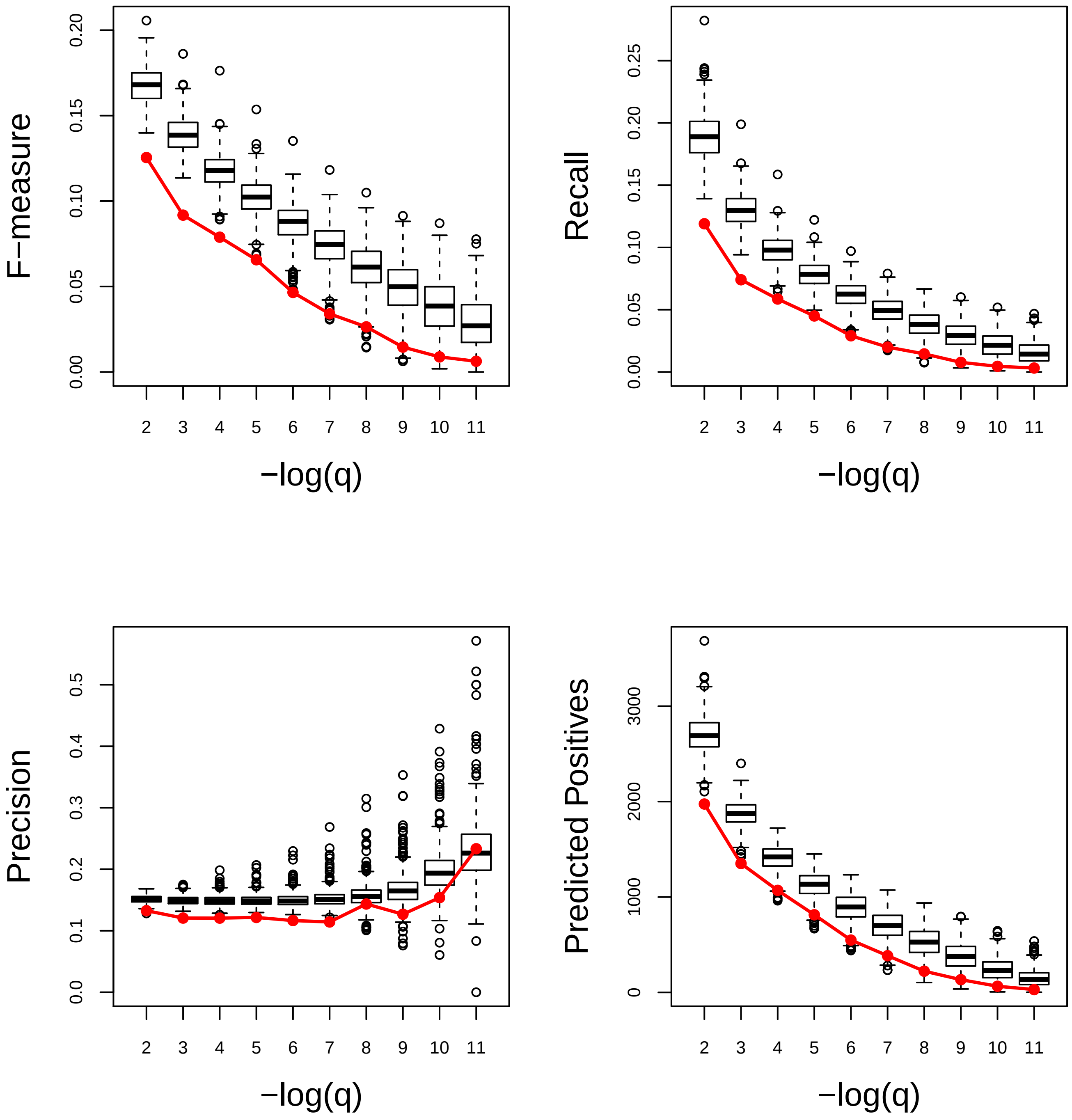

### Xylan metabolic process. Good

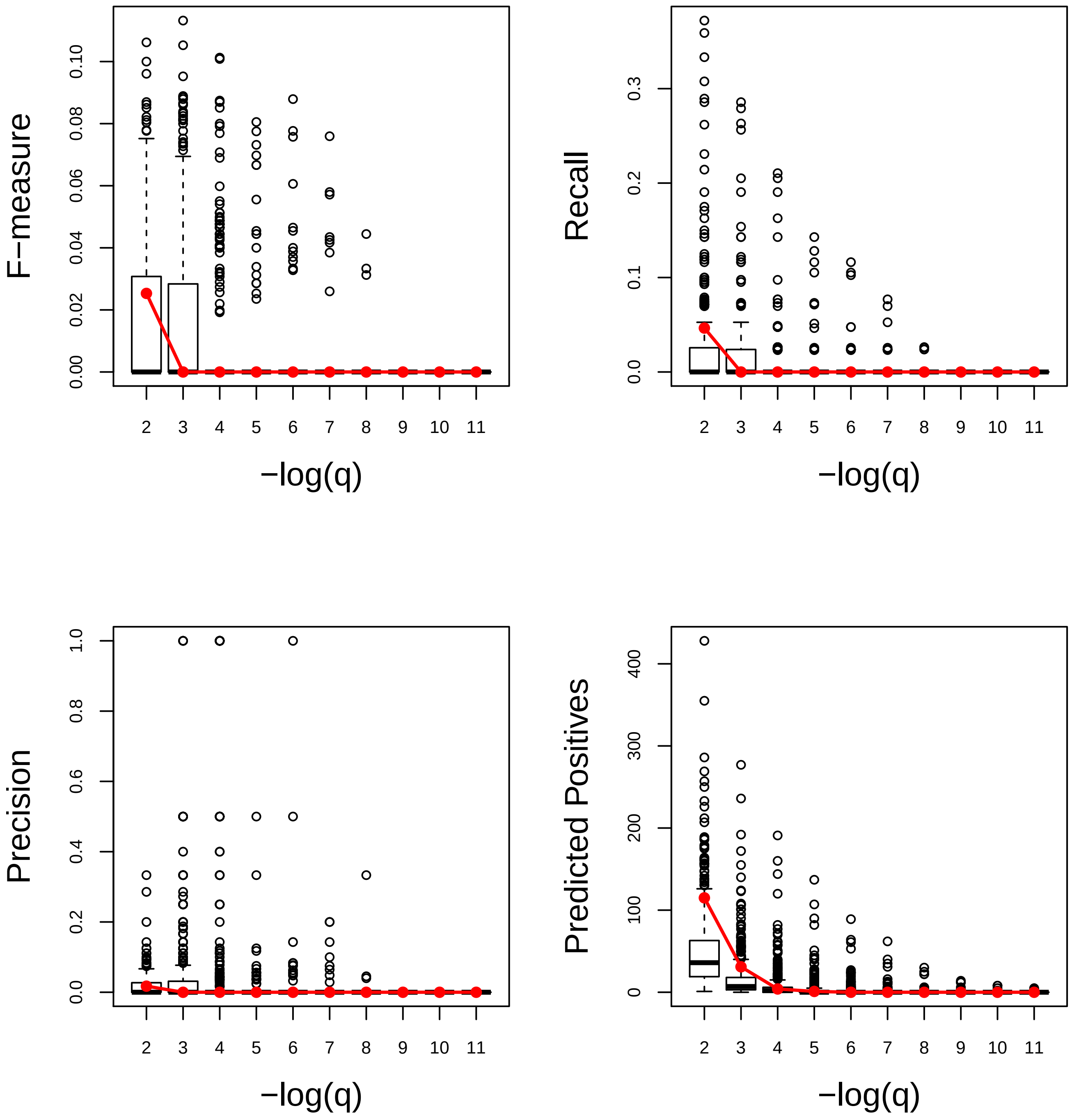
